# Reduced cue-induced reinstatement of cocaine-seeking behavior in Plcb1+/- mice

**DOI:** 10.1101/2020.06.18.158964

**Authors:** Judit Cabana-Domínguez, Elena Martín-García, Ana Gallego-Roman, Rafael Maldonado, Noèlia Fernàndez-Castillo, Bru Cormand

**Author notes:** These authors contributed equally (Judit Cabana-Domínguez, Elena Martín-García). These authors equally supervised this work (Rafael Maldonado, Noèlia Fernàndez-Castillo, Bru Cormand). Address corresponding to Bru Cormand, Ph.D., and Noèlia Fernàndez-Castillo, Ph.D., Departament de Genètica, Microbiologia i Estadística, Facultat de Biologia, Universitat de Barcelona, Avinguda Diagonal 643, edifici Prevosti, 3^<u>a</u>^ planta, 08028 Barcelona, Catalonia, Spain.

## Abstract

**Background and Purpose:** Cocaine addiction causes serious health problems and no effective treatment is available yet. We previously identified a genetic risk variant for cocaine addiction in the *PLCB1* gene and found this gene upregulated in postmortem brains of cocaine abusers and in human dopaminergic neuron-like cells after an acute cocaine exposure. Here, we functionally tested the contribution of *PLCB1* gene to cocaine addictive properties in mice.

**Experimental approach:** We used heterozygous *Plcb1* knockout mice (*Plcb1*+/-) and characterized their behavioral phenotype. Subsequently, mice were trained for operant conditioning and self-administered cocaine for 10 days. *Plcb1*+/- mice were assessed for cocaine motivation, followed by 26 days of extinction and finally evaluated for cue-induced reinstatement of cocaine seeking. Gene expression alterations after reinstatement were assessed in medial prefrontal cortex (mPFC) and hippocampus (HPC) by RNAseq.

**Key Results:** *Plcb1*+/- mice showed normal behavior, although they had increased anxiety and impaired short-term memory. Importantly, after cocaine self-administration and extinction, we found a reduction in the cue-induced reinstatement of cocaine-seeking behavior in *Plcb1*+/- mice. After reinstatement, we identified transcriptomic alterations in the medial prefrontal cortex of *Plcb1*+/- mice, mostly related to pathways relevant to addiction like the dopaminergic synapse and long-term potentiation.

**Conclusions and Implications:** To conclude, we found that heterozygous deletion of the *Plcb1* gene decreases cue-induced reinstatement of cocaine seeking, pointing at PLCB1 as a possible therapeutic target for preventing relapse and treating cocaine addiction.

## INTRODUCTION

Cocaine is the most used psychostimulant illicit drug worldwide (UNODC, 2019), causing severe health problems that include the development of cocaine addiction in around 15-16% of cocaine users (O’Brien and Anthony, 2005). Cocaine addiction is a complex psychiatric disorder that results from the interaction of genetic, epigenetic, and environmental risk factors (Pierce et al., 2018). The heritability of cocaine addiction is one of the highest among psychiatric disorders, estimated around 65% for women (Kendler et al., 2000) and 79% for men (Kendler and Prescott, 1998). However, the genetic factors and mechanisms that underlie the transition from drug use to addiction and its establishment remain unknown.

In a previous study, we identified a single nucleotide polymorphism in the *PLCB1* gene associated with drug dependence (rs1047383), and especially with a subgroup of cocaine-addicted patients, that was replicated in an independent clinical sample (Cabana-Domínguez et al., 2017). Also, genetic variants in this gene were found nominally associated with an illegal substance and cocaine addiction in two GWAS performed in European-American samples (Drgon et al., 2012). On the other hand, we found that cocaine increased the expression of *PLCB1* both in human dopaminergic neuron-like cells (differentiated SH-SY5Y cells) after acute cocaine exposure and in postmortem samples of the nucleus accumbens (NAc) of cocaine abusers (Cabana-Domínguez et al., 2017). Interestingly, this gene was also found over-expressed in the same brain region in mice after cocaine administration for 7 days and also during withdrawal (Eipper-Mains et al., 2013). All this evidence suggest that *PLCB1* may play a role in cocaine addiction.

The *PLCB1* gene encodes phospholipase C beta 1, and it is highly expressed in the brain, mainly in the frontal cortex, basal ganglia (caudate, putamen, and NAc) and hippocampus. These brain regions are crucial for drug reward and the formation of drug-context associations, both contributing to the development and maintenance of addiction (Bisagno et al., 2016; Castilla-Ortega et al., 2016, 2017; Kutlu and Gould, 2016; Pitts et al., 2016; Cooper et al., 2017). Several neurotransmitters activate this protein, including dopamine through DRD1 and DRD2 (Lee et al., 2004; Rashid et al., 2007), serotonin by 5-HT2A and 2C receptors (Hagberg et al., 1998; Chang et al., 2000) and glutamate by mGluR1 (Conn and Pin, 1997; Hannan et al., 2001). Thus, PLCB1 might be a point of convergence of neurotransmitter systems that play an essential role in the development of addiction (recently reviewed (Howell and Negus, 2014; Moretti et al., 2020)). The activation of PLCB1 produces the cleavage of phosphatidylinositol 4,5-bisphosphate (PIP2) into the second messengers diacylglycerol (DAG) and inositol 1,4,5-trisphosphate (IP3), responsible for intracellular signal transduction. Alterations in PLCB1-mediated signaling in the brain have been associated with other neuropsychiatric disorders such as epilepsy, schizophrenia, and bipolar disorder (Yang et al., 2016).

Here we studied the contribution of the *PLCB1* gene to cocaine addiction and dissected its participation in the different aspects of the addictive process. We used heterozygous knockout (KO) mice (*Plcb1*+/-), as the homozygous KO (*Plcb1*-/-) showed seizure attacks and low viability after birth (Kim et al., 1997). In addition, the use of a constitutive heterozygous KO mouse model allowed us to assess *Plcb1* haploinsufficiency during neurodevelopment in mice similarly to humans, where inherited genetic risk or protective variants can modulate the susceptibility to addiction (Cabana-Domínguez et al., 2017). First, we performed a general phenotypic characterization using a battery of behavioral tests, including memory, anxiety, locomotor activity, coordination, food and water intake, and sucrose preference. Then, we evaluated cocaine operant self-administration, extinction, and cue-induced reinstatement. Finally, we studied transcriptional alterations to further understand the molecular mechanisms involved.

## RESULTS

In the present work, we aimed to assess the contribution of the *PLCB1* gene in the different stages of cocaine addiction using heterozygous knockout mice (*Plcb1*+/-). To do so, we performed a general phenotypic characterization and, then we evaluated cocaine operant self-administration, extinction, and cue-induced reinstatement in those animals. Also, we studied transcriptional changes in mPFC and HPC to understand the mechanisms involved.

### Behavioral tests for phenotype characterization

We first evaluated the effects of the heterozygous deletion of *Plcb1* on general behavioral responses, including locomotor, cognitive and emotional responses, food, and water intake. *Plcb1*+/- mice showed a lower discrimination index in the novel object recognition compared to WT (*t*-test=3.21, P<0.01, Figure 1B), suggesting an impairment in short-term memory. This difference was not influenced by exploration time, as this variable was equal between genotypes (Figure 1C). Furthermore, no differences in locomotor activity were reported between genotypes, discarding an involvement of the *Plcb1* heterozygous deletion in locomotion (Figure 1D). Mutant mice showed increased exploratory activity with a higher number of rearings than the WT mice (*U*-Mann Whitney=67.5, P<0.01, Figure 1E). An anxiogenic profile was revealed in the elevated plus maze in mutants, as shown by the reduced time spent in the open arms (*t*-test=3.02, P<0.001, Figure 1F) and the percentage of time (*t*-test=3.16, P<0.001, Figure 1G). Finally, the rota rod test revealed that the heterozygous deletion of *Plcb1* did not affect motor coordination (Figure 1H-I).

**Figure 1.**
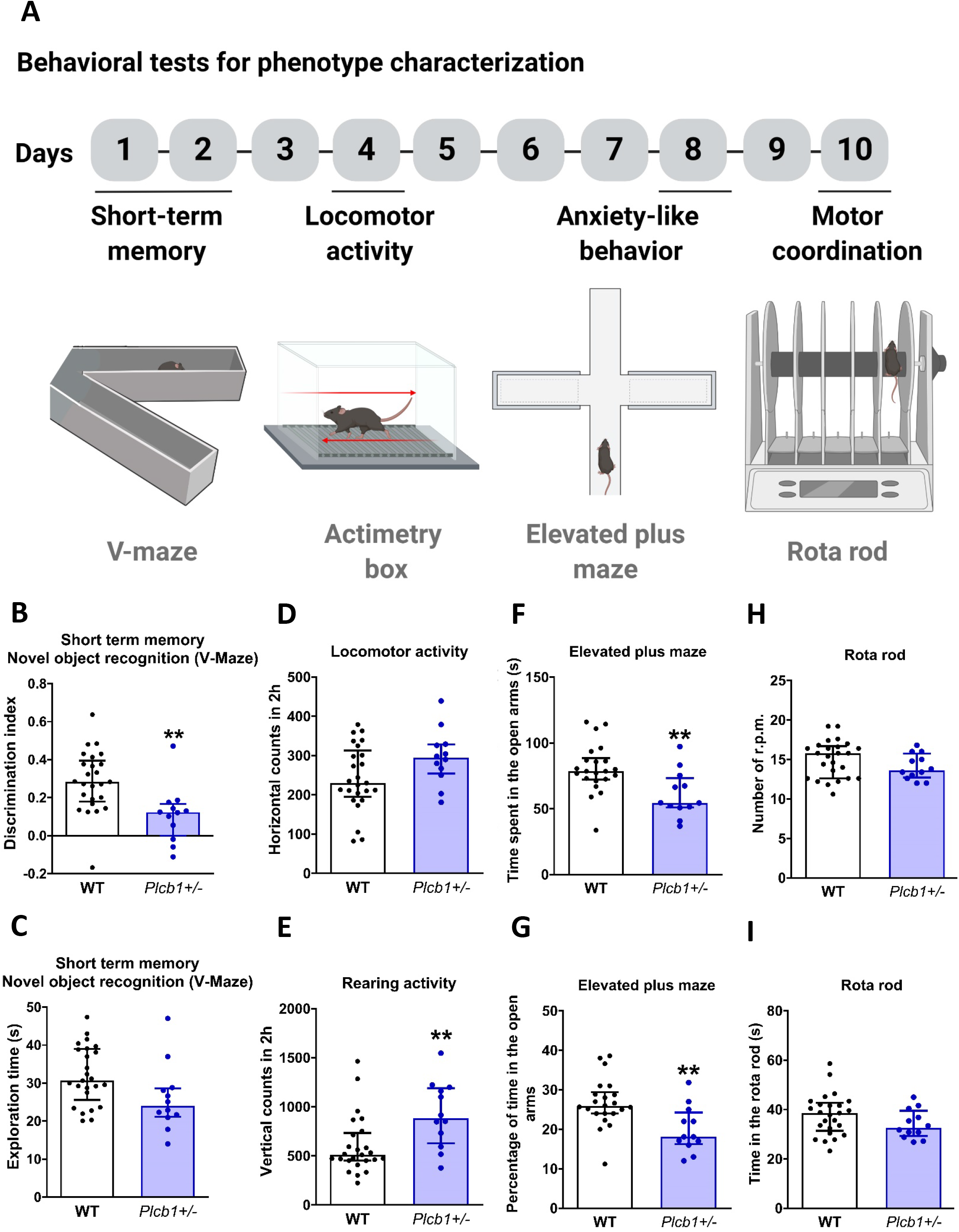
Behavioral tests for phenotype characterization. (A) Timeline of the experimental sequence. (B,C) Short-term memory measured in the V-maze for novel object recognition task, (B) discrimination index and (C) exploration. (D) Locomotor and (E) rearing activity measured in actimetry boxes. (F) Anxiety-like behavior measured by the time spent in the open arms and (G) the percentage of time in the open arms in the elevated plus maze. Motor coordination measured in the rota rod test by the (H) number of r.p.m. that the mice performed in the average trials or the (I) mean time they remain in the apparatus. All data are expressed in median and interquartile range and individual data are shown (WT n=25; Plcb1+/- n=12).

Several consummatory and locomotor parameters were long-term monitored in the PheComp boxes (Supplementary Figure 1A-H, n=6 per genotype). No differences between genotypes were revealed during the whole experimental period in body weight, food, and water intake, levels of sucrose preference, stereotyped movements, horizontal and vertical locomotor activity, indicating no altered behavior in the *Plcb1*+/- mice.

### Operant conditioning maintained by cocaine

Then, we investigated the effects of the heterozygous deletion of *Plcb1* in cocaine behavioral responses related to its addictive properties. For this purpose, *Plcb1*+/- mice and their WT littermates were trained for cocaine operant self-administration (0.5mg/kg/infusion) during FR1 and FR3, progressive ratio, extinction and cue-induced reinstatement (Figure 2A). Control mice trained with saline were included for both genotypes. Results showed that the percentage of mice reaching the criteria of operant conditioning was 100% for both genotypes trained with cocaine and 33% for mice trained with saline: [*Chi-square* test=9.50; P<0.01, *Plcb1*+/- cocaine vs. *Plcb1*+/- saline] and [*Chi-square* test=14.00; P<0.001, WT cocaine vs. WT saline], as expected.

**Figure 2.**
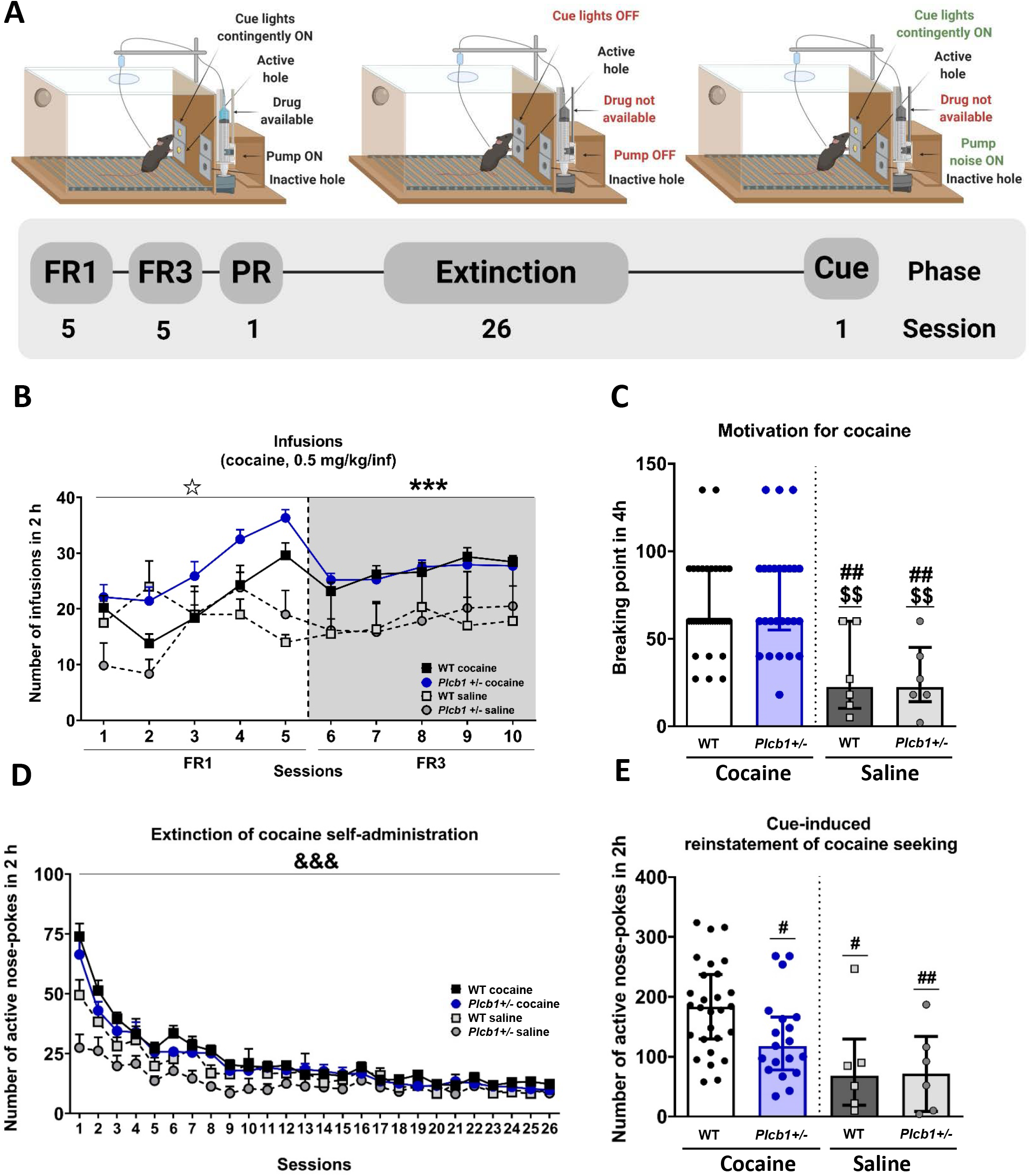
Operant conditioning maintained by cocaine in Plcb1+/-, wild-type (WT) and corresponding control saline mice. (A) Timeline of the experimental sequence. (B) The number of cocaine infusions (0.5 mg/kg/infusion) in both genotypes increased progressively across sessions during FR1 (repeated measures ANOVA, interaction between genotype x drug x sessions, P<0.05) and remained stable during FR3 and higher in mice trained with cocaine (repeated measures ANOVA, main effect of drug, ***P<0.001). The maximum number of infusions was reached on session 5 in both genotypes and was slightly superior in Plcb1+/- (36.35±1.47) than in WT (29.61±2.23) but post hoc analyses demonstrated that this difference was not significant. (C) The motivation for cocaine measured by the breaking point achieved in the progressive ratio schedule of reinforcement (4h) was equivalent in both genotypes trained with cocaine but higher than those trained with saline independently of the genotype (individual data with interquartile range, U Mann-Whitney, ##P<0.01 vs WT cocaine, $$P<0.01 vs Plcb1+/- cocaine). (D) Both genotypes also showed similar levels of extinction that decreased during sessions, but the curve was more pronounced in WT mice than in mutants (repeated measures ANOVA, interaction genotype x sessions, &&&P<0.001). (E) Decreased cue-induced reinstatement of cocaine seeking in Plcb1+/- mice was obtained compared to WT (individual data with interquartile range, U Mann-Whitney, #P<0.05, ##P<0.01 vs WT cocaine). All data are expressed in mean ± SEM when sessions are represented or median and interquartile range when individual data are shown (WT cocaine n=28-36; Plcb1+/- cocaine n=19-26; saline n=6 per genotype). Statistical details are included in Supplementary Table S1.

The primary reinforcing effects and the motivation for cocaine were similar in both genotypes (Figure 2B-C). During FR1, both genotypes similarly increased the number of cocaine infusions across sessions, whereas mice trained with saline remained steady (repeated measures ANOVA, interaction between genotype x drug x sessions, P<0.05, Figure 2B and Supplementary Table S1). Thus, the evolution of operant responding was different in mice trained with cocaine and saline, with an increased responding over nearly all sessions in cocaine-trained mice. This enhancement in cocaine responding was more pronounced in mutants than WT mice. Indeed, WT mice showed a decrease in cocaine intake in the second session. The maximum number of infusions was reached on session 5 in both genotypes and was slightly superior in *Plcb1*+/- mutants (36.35±1.47) than in WT (29.61±2.23) but post hoc analyses demonstrated that this difference in cocaine intake was not significant (Figure 2B). Similar findings were observed for the total number of nose-pokes in which operant responding increased in both genotypes with cocaine but remained stable with saline (Supplementary Figure S2A). When the effort to obtain one dose of cocaine increased to FR3, the number of infusions was stable across sessions in WT and *Plcb1*+/- mice. Similarly, operant responding was higher for all groups than in FR1 with stable higher levels of responding (Supplementary Figure S2A) and a higher number of infusions (repeated measures ANOVA, the main effect of the drug, P<0.001, Figure 2B) in mice trained with cocaine than with saline, independently of the genotype.

Motivation for cocaine was evaluated in a progressive ratio schedule, and no significant differences were obtained between genotypes (Figure 2C). The levels of extinction of the operant behavior were similar between genotypes and decreased progressively across sessions (repeated measures ANOVA, interaction genotype x sessions, P<0.001, Figure 2D). The percentage of mice reaching cocaine-seeking extinction criteria was similar in *Plcb1*+/- (73%) and WT (78%) mice.

Importantly, *Plcb1*+/- mice showed significantly reduced cue-induced reinstatement of cocaine-seeking compared to WT mice (*U* Mann-Whitney, P<0.05, Figure 2E), with 27.59% less active nose-pokes compared to WT mice trained with cocaine. Furthermore, mice trained with saline from both genotypes exhibited 55.84% reduction of active nose-pokes than WT mice trained with cocaine and 39.20% less than mutants trained with cocaine. No significant differences were obtained between genotypes in inactive nose-pokes during operant conditioning maintained by cocaine nor during extinction (Supplementary Figure S2B-C). Both genotypes trained with cocaine acquired the reinstatement criterion (double nose pokes in the active hole than the number of nose pokes during the 3 consecutive days when the mice acquired the extinction criteria) showing their capability to maintain this conditioning learning task. These data showed that *Plcb1*+/- resulted in a phenotype of resistance to cue-induced reinstatement with reduced cocaine-seeking (Figure 2E).

### Brain transcriptomic analysis after the reinstatement of cocaine-seeking behavior

To further understand the role of PLCB1 in the molecular mechanisms involved in this cocaine relapse-related phenotype, we analyzed the transcriptomic profiles of mPFC and HPC immediately after the reinstatement of cocaine-seeking behavior in WT and *Plcb1*+/- mice trained with cocaine. We identified 2,115 protein-coding genes differentially expressed (DEGs) in mPFC (1,231 downregulated and 943 upregulated) and only 12 in HPC, when comparing *Plcb1*+/- and WT animals (Supplementary Tables S2-3). In accordance, PCA analysis and heatmap plots revealed that individuals with the same genotype plotted together only in mPFC, but not in HPC (Supplementary Figure S3). This suggested a more prominent role of mPFC, so further studies were carried out only with DEGs of this brain area. Analysis of functional group over-representation identified several processes previously related to cocaine addiction, including dopaminergic synapse, learning, long-term potentiation (LTP), neurotransmitter secretion, and axon guidance, as well as other relevant signaling pathways such as MAPK, mTOR, and neurotrophin (Figure 3A-B and Supplementary Tables S3-5). We focused on the dopaminergic synapse pathway (enriched in the mPFCs DEGs; *P*_Adj_<0.05), in which phospholipase c (such as Plcb1) is directly participating in signal transmission (Figure 4). Interestingly, many genes coding for proteins in this pathway are differentially expressed in *Plcb1*+/- mice after cue-induced reinstatement (Figure 4 and Supplementary Table S6). Furthermore, we found that several TFBS were over-represented in the DEGs of mPFC, including YY1, MYOD, NRF1, ERR1, FREAC2, NFY and E4F1 (complete list of TFBS in Supplementary Table S7). Then, we filtered the DEGs on mPFC based on fold-change (FC> |1.2|) and obtained a list of 238 genes that we used for gene network construction. This analysis showed a highly scored network (score=62, Figure 3C) that includes 31 DEGs involved in “cellular development, cellular growth and proliferation, nervous system development and function”.

**Figure 3.**
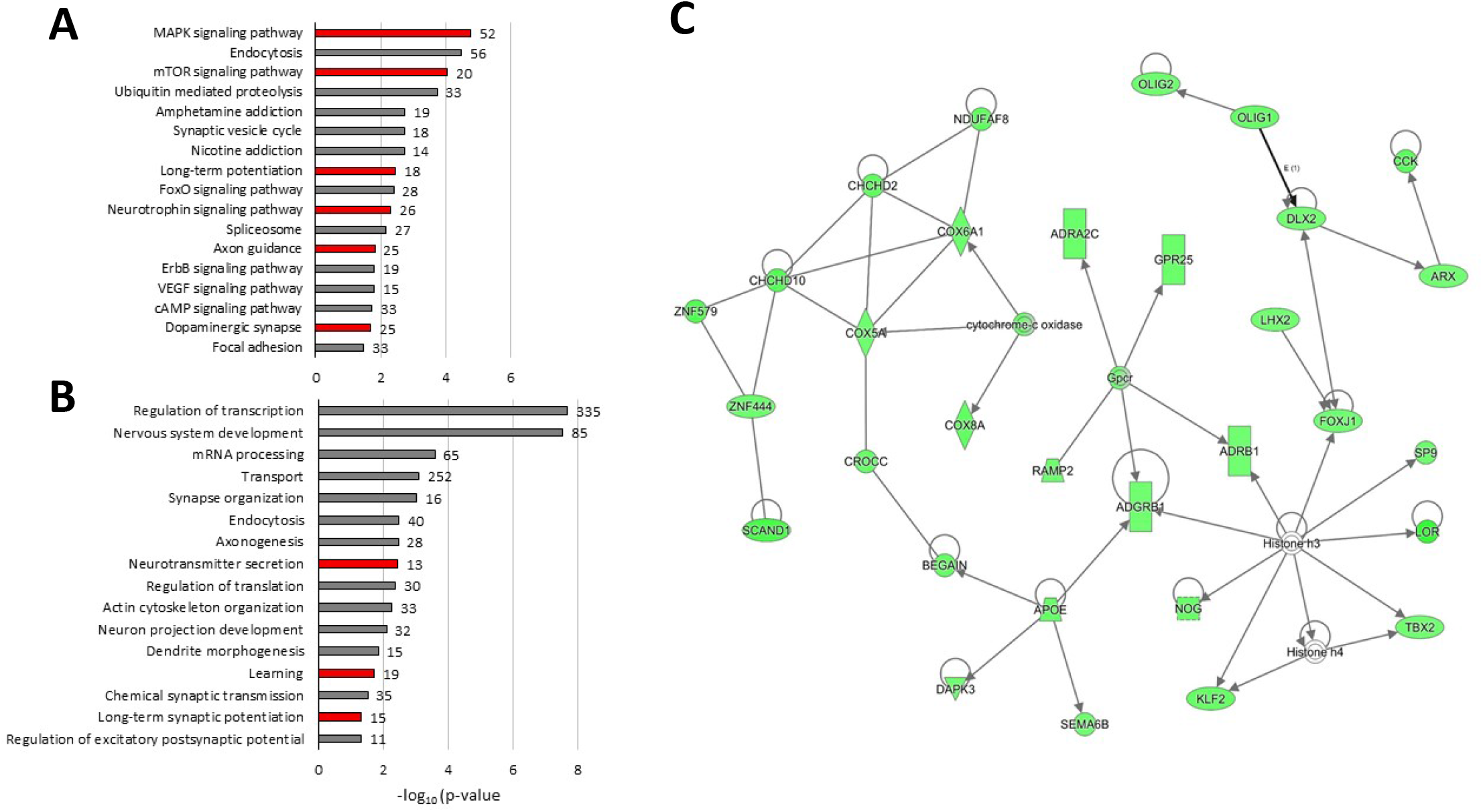
Gene expression changes in mPFC after cue-induced reinstatement of cocaine seeking comparing Plcb1+/- versus wild-type (WT) mice. (A) Selection of over-represented KEGG pathways (Kyoto Encyclopedia of Genes and Genomes) and (B) GO (Gene Ontology) identified by DAVID software among the differentially expressed genes. The number of genes with altered expression included in each category is indicated on the right side of the bar. In red, relevant pathways for the addictive process. (C) Gene network involved in cellular development, cellular growth and proliferation, nervous system development and function (score = 62). The green nodes in the pathway indicate genes with downregulated expression in Plcb1+/- identified in RNAseq.

**Figure 4.**
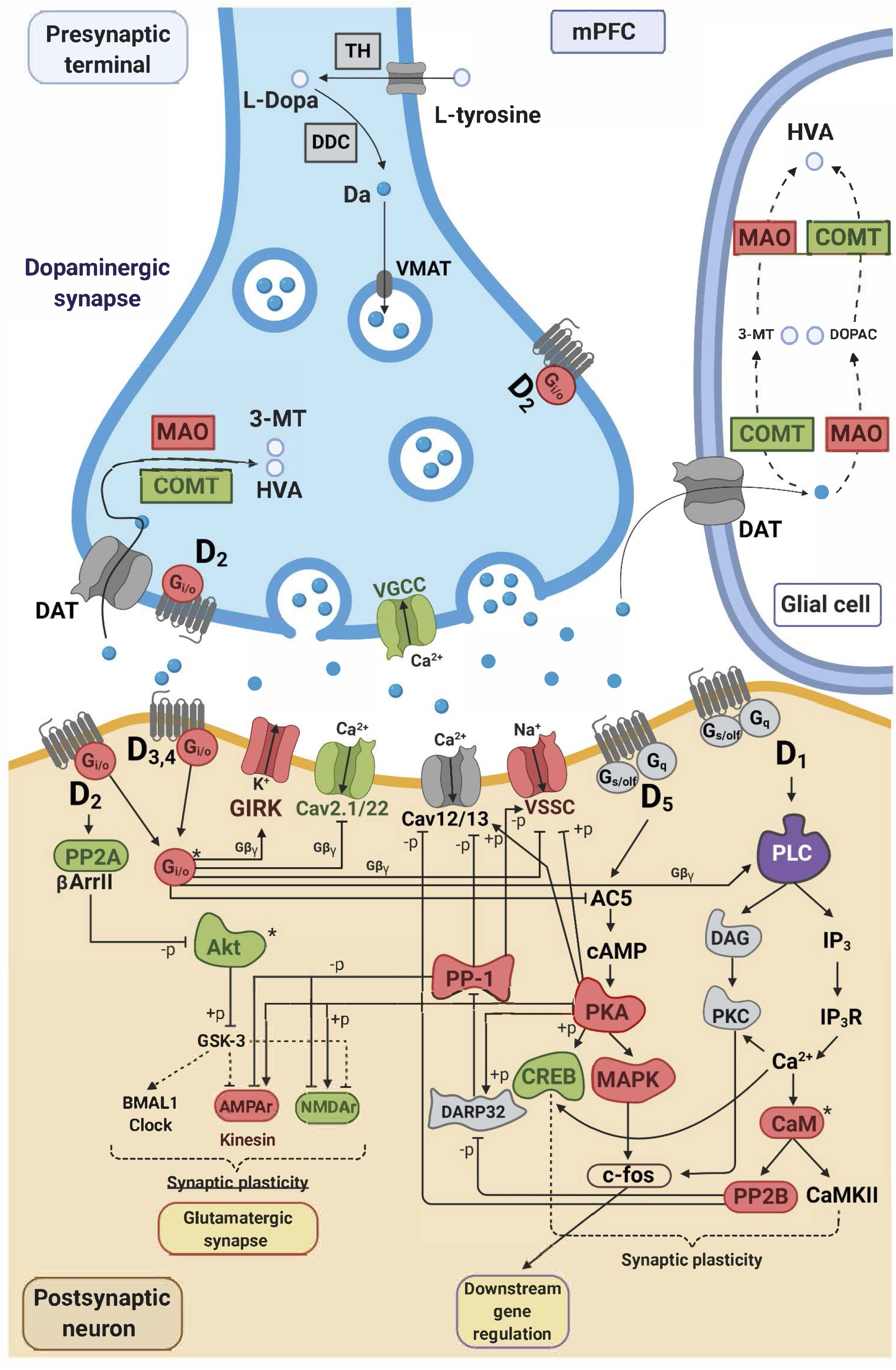
Alterations in expression in the dopaminergic synapse in mPFC of Plcb1+/- mice after cue-induced reinstatement of cocaine seeking. Adapted from KEGG (Kyoto Encyclopedia of Genes and Genomes) pathways (mmu04728). Enriched pathway in DEGs in mPFC, 25 out of 131 genes in the pathway were differently expressed (Praw=2e-03; Pajd=0.02). In red, upregulated genes and in green, downregulated genes in mPFC. *Protein complexes including upregulated and downregulated genes. Correspondence between genes and proteins can be found in Supplementary Table S6.

## DISCUSSION

Here we investigated, for the first time, the contribution of the *PLCB1* gene to cocaine addictive properties using *Plcb1*+/- mice. We found that the heterozygous *Plcb1* genotype resulted in a phenotype of resistance to cue-induced reinstatement of cocaine-seeking behavior. Furthermore, we found relevant transcriptomic differences in *Plcb1*+/- mice compared to WT after cue-induced reinstatement of cocaine seeking in mPFC, a brain area essential in relapse (Jasinska et al., 2015; Moorman et al., 2015). This study supports a role for *PLCB1* in cocaine addiction, confirming previous findings in humans (Cabana-Domínguez et al., 2017), and suggests it may be relevant in relapse to cocaine addiction.

The phenotype of *Plcb1*+/- mice was characterized at different levels to evaluate consummatory and general behavior. In general, mutant mice presented normal levels of body weight, food and water intake, locomotor activity, and motor coordination, demonstrating that this animal model is valid to study the effects of the heterozygous deletion of the *Plcb1* gene in other behavioral responses. *Plcb1*+/- showed increased anxiety in the elevated plus maze, reduced short-term memory in the novel object recognition paradigm, without any sign of depressive-like behavior in the anhedonia sucrose preference test. Thus, the single targeting of the *Plcb1* gene has an impact on selective emotional and cognitive responses.

The short-term memory impairment observed in the mutants did not affect the acquisition of operant associative conditioning nor the instrumental learning that drives the goal-directed action, as shown by similar levels of operant cocaine self-administration and extinction learning than WT. In agreement, both genotypes trained with cocaine accomplish cue-induced reinstatement criterion with high enhancement of responding during this test compared to extinction. Thus, different brain circuits are involved in each kind of learning, with perirhinal-hippocampal structures (Winters et al., 2004) participating in short-term memory in the novel object recognition paradigm and mPFC-dorsal striatum pathway in cue-associated seeking. Also differences in each paradigm such as the acute retrieval or repetitive exposure involved respectively in each task may have a crucial influence in cognition. Hence, cocaine seeking can be multidimensional, involving different types of associative learning that together lead to an extensive repertoire of conditioned and instrumental responding.

The anxiogenic profile of *Plcb1*+/- is in accordance with the results previously reported in the *Plcb4*-/- mice, which were associated with alterations in the cholinergic activity of the medial septum (Shin et al., 2009). However, the selective knock-down of *Plcb1* in the mPFC did not replicate this phenotype in a previous study (Kim et al., 2015), suggesting that other areas may be involved. The anxiogenic profile of *Plcb1*+/- mice had no effect on cocaine self-administration since the acquisition and extinction of this operant behavior was not modified in the mutants. Besides, mutants showed protection against cue-induced reinstatement of cocaine seeking, instead of the expected cocaine seeking promoted by an anxiogenic phenotype (Mantsch et al., 2016).

Therefore, the association of an anxiogenic profile with a phenotype resilient to cocaine seeking in the *Plcb1*+/- mice suggests modifications in specific brain areas involved in cocaine relapse, such as the mPFC. Recently, a phenotype resilient to develop food addiction has been associated with increased strength of pyramidal glutamatergic synaptic transmission in the mPFC related to decreased compulsivity in the face of negative consequences (Domingo-Rodriguez et al., 2020) in mice with an anxiogenic profile (Lutz et al., 2015). Concerning cue-induced cocaine seeking, the mPFC has a crucial role and the network of glutamatergic projections from the prelimbic mPFC to the dorsal striatum participates in this phenotype (Lüscher et al., 2020). In our model, *Plcb1* haploinsufficiency is linked to reduced cue-induced cocaine seeking possibly associated with modifications in this top-down corticolimbic brain network. Furthermore, these glutamatergic projections from mPFC to dorsal striatum receive mesocortical dopaminergic inputs from the VTA that could be crucially involved in the protective effects of *Plcb1*, since this gene plays an essential role in the dopaminergic signal transmission in the mPFC and many of the genes encoding for proteins in this pathway are differentially expressed in *Plcb1*+/- mice after cue-induced reinstatement (Figure 4).

Transcriptomic analyses performed after the cue-induced reinstatement of cocaine-seeking behavior revealed some of the molecular underpinnings that underlie the cocaine relapse-related phenotype observed in the mutant mice. Interestingly, differences in gene expression between *Plcb1*+/- and WT were predominantly found in the mPFC, whereas almost no differences were observed in the HPC. This evidence highlights mPFC as a key region to explain these differences in cocaine-seeking reinstatement observed in *Plcb1*+/- mice. Consistently, in DEGs in mPFC, we found enrichment on pathways that are essential for the development of addiction, including the dopaminergic neurotransmission (Figures 3–4) (Wise and Robble, 2020). *PLCB1* is part of the dopamine-DARPP-32 signaling pathway (Figure 4), which plays a key role in cocaine reward (Borgkvist and Fisone, 2007; Nishi and Shuto, 2017). In a previous study, we also found this pathway enriched in DEGs in the frontal cortex and ventral striatum of mice that showed frustrated expected reward, produced by the cue in the absence of expected reward after a high level of effort in a progressive ratio schedule of reinforcement with palatable food (Martín-García et al., 2015b). Notably, *Plcb1* was upregulated in the frontal cortex of those frustrated mice with increased responses to obtain the reward (palatable food). These data are in line with our findings in which decreased expression of *Plcb1 (Plcb1*+/- mice) results in decreased cue-induced responses to obtain cocaine after extinction. All these results support the participation of *Plcb1* in cue-induced cocaine seeking revealed in the present study and the involvement of mPFC.

Our transcriptomic analyses also found enrichment in genes related to learning and memory processes, such as LTP in the mPFC. The transition from drug use to drug addiction is a maladaptive process that directly affects learning and memory (Torregrossa et al., 2011; Goodman and Packard, 2016). LTP produces long-lasting activity-dependent synaptic modifications that underlie memory, and drugs of abuse alter this mechanism in several brain regions such as mPFC (Huang et al., 2007; Lu et al., 2010), mesocorticolimbic system (Thomas et al., 2008), VTA (Placzek et al., 2016) and hippocampus (Keralapurath et al., 2017; Preston et al., 2019), among others. Importantly, genes related to the glutamatergic system such as *Gria2-4* (glutamate ionotropic receptor AMPA2-4 (alpha 2-4)) and *Grik2* (ionotropic glutamate receptor kainate 2 (beta 2)) were found to be upregulated in *Plcb1*+/- mice. Meanwhile, *Grin1, 2c, and 2d* (glutamate ionotropic receptor NMDA1 (zeta 1, epsilon3 and epsilon4)) were downregulated, suggesting an increase in AMPAR/NMDAR ratio in cocaine-experienced and abstinent *Plcb1* +/- mice as compared to WT, one of the key indicators of LTP induction and increased synaptic strength (Lüscher and Malenka, 2011). Furthermore, we found enrichment in other neuroplasticity (synapse organization, neuron projection development, and axon guidance) (Bahi and Dreyer, 2005; O’Brien, 2009; Cooper et al., 2017; Dong et al., 2017) and signaling pathways (MAPK, mTOR, and neurotrophin) (Li and Wolf, 2015; García-Pardo et al., 2016) related to addiction. The assessment of gene expression in the mPFC of the *Plcb1*+/- mice highlighted alterations in relevant pathways for the addictive process, which could contribute to the results observed in the cue-induced reinstatement.

Nowadays, there are few effective pharmacological treatments available for cocaine use disorders, and frequently, psychosocial interventions in combination with pharmacotherapy are needed (Kampman, 2019). Studies performed in mice have pinpointed multiple potential therapeutic approaches for the reinstatement of cocaine-seeking behavior, targeting the reward circuit (Buchta et al., 2020). In humans, treatment with bupropion, topiramate, or disulfiram has been widely used. Bupropion, a non-tricyclic antidepressant that inhibits dopamine and norepinephrine reuptake, is effective in reducing craving (Frishman, 2007). Topiramate, a GABA/glutamatergic medication, has also been used to treat cocaine use disorder as it reduces the activity of the mesocorticolimbic dopaminergic system (Kampman, 2019). A completely different strategy is the use of disulfiram which potentially inhibits the oxidoreductase dopamine β-hydroxylase (DβH, encoded by the *DBH* gene), which converts dopamine to norepinephrine (Kampangkaew et al., 2019). Significantly, a genetic variation in the *SLC6A3* gene (encoding DAT) has been associated with disulfiram treatment for cocaine addiction, with patients with higher DAT levels having better treatment outcomes than those with lower DAT levels (Kampangkaew et al., 2019). Thus, further studies and new therapeutic targets are needed to obtain effective treatment for cocaine addiction. The results obtained in the present study underscore the relevance of the *Plcb1* gene in the cue-induced reinstatement of cocaine seeking after extinction. Together with previous findings in humans (Drgon et al., 2012; Cabana-Domínguez et al., 2017) and mice (Eipper-Mains et al., 2013), *PLCB1* merits to be further evaluated as a promising novel therapeutic target for preventing relapse and treating cocaine addiction.

The experimental approach used in our study, the *Plcb1*+/- mouse model, allowed us to reproduce better the molecular context observed in humans in comparison to the use of a complete KO mouse (Hall et al., 2013), as these animals preserved, at least, half of the expression of *Plcb1*. However, the haploinsufficiency of *Plcb1* during neurodevelopment in these animals could produce alterations that may contribute to the phenotype observed in the present study. Nevertheless, this may also be similar in humans with genetic risk variants that decrease *PLCB1* expression. Therefore, this approach is appropriate to study a specific genetic alteration that confers susceptibility to drug addiction and to delineate the precise contribution of *PLCB1*.

To sum up, we studied, for the first time, the contribution of the *PLCB1* gene to cocaine addictive properties using *Plcb1*+/- mice. Previous studies have revealed an up-regulation of *Plcb1* in brain areas related to the reward circuit after cocaine exposure in animals (Eipper-Mains et al., 2013) and in human cocaine abusers (Cabana-Domínguez et al., 2017). These changes, together with our results, suggest that cocaine increases the expression of *Plcb1*, and this mechanism plays an essential role in cocaine addiction, as revealed now by the resistance to cue-induced reinstatement of cocaine-seeking behavior exhibited by *Plcb1*+/- mutant mice. These results highlight the importance of the *Plcb1* gene in the development of cocaine addiction and relapse and pinpoint PLCB1 as a promising therapeutic target for cocaine addiction and perhaps other types of addiction.

## METHODS

### Animals

Male mice, 8 weeks old, were housed individually at temperature- and humidity-controlled laboratory conditions (21±1°C, 55±10%) maintained with food and water ad libitum. Mice were tested during the dark phase of a reverse light cycle (lights off at 8.00h and on at 20.00h). *Plcb1*+/- mice in a C57BL/6J background and their wild-type (WT) littermates were used (Kim et al., 1997). All experimental protocols were performed in accordance with the guidelines of the European Communities Council Directive 2010/63/EU and approved by the local ethics committee (Comitè Ètic d’Experimentació Animal-Parc de Recerca Biomèdica de Barcelona, CEEA-PRBB, agreement N°9213). In agreement, maximal efforts were made to reduce the suffering and the number of mice used.

### Genotyping of transgenic mice

*Plcb1*+/- mice in a C57BL/6J background were kindly provided by H.-S. Shin (Kim et al., 1997). Very briefly, the knockout animals were generated by the replacement of exons that encode aminoacid residues 50–82 by a Neomycin cassette that was used both for gene disruption and for positive selection.

To genotype the mice, genomic DNA from ear punches or tail tips was extracted as previously described (Gómez-Grau et al., 2017). Genotyping was carried out by PCR on extracted genomic DNA using three specific PCR primers: a sense primer, 5□-GTTAAGTCCTCAGGCAAACACC-3’, and two antisense primers, 5□-ACCTTGGGAGCTTTGGCGTG3’ and 5□-CTGACTAGGGGAGGAGTAGAAG-3’, that allowed us to amplify a 180bp *Plcb1*+/+ band and a 290bp *Plcb1*-/- band (Filis et al., 2009). Fragments were separated by 2.5% agarose gel electrophoresis. Known WT and *Plcb1*+/- controls were always used to validate genotyping results. The genotype of all animals used was confirmed after the cue-induced reinstatement.

### Behavioral tests for phenotype characterization

A group of 25 WT and 12 *Plcb1*+/- mice underwent different behavioral tests for phenotype characterization for 10 days, as described in Figure 1:

#### Short-term memory

To measure short-term memory, the novel object recognition (NOR) test was used. This test was performed in a V-maze apparatus (40 cm per side, Panlab, Spain) as previously described by us and others (Bura et al., 2010; Planaguma et al., 2015). On day 1 mice were habituated for 9 minutes in the V-maze. On day 2, mice were put back into the V-maze for 9 minutes; two identical objects were presented, and the time the mice spent exploring each object was recorded. After a retention phase of 3 hours, the mice were placed for another 9 minutes into the V-maze; in each paradigm, one of the familiar objects was replaced with a novel object and the total time spent exploring each object (novel and familiar) was registered. Object exploration was defined as the orientation of the nose to the object at a distance of less than 2 cm. A discrimination index was calculated as the difference in the time spent exploring the novel and the time spent exploring the familiar object divided by the total time exploring both objects. A higher discrimination index is considered to reflect more excellent memory retention for the familiar object.

#### Locomotor activity

Locomotor activity was assessed in locomotor activity boxes (9×20×11 cm; Imetronic, Passac, France), equipped with 2 rows of photocell detectors, and placed in a low-luminosity environment (20–25 lux), as previously described (Martin-Garcia et al., 2013). The locomotor activity was recorded for 2 hours, and 2 variables were measured: horizontal activity as horizontal displacement and vertical activity as vertical exploration/elevation.

#### Elevated plus maze test

This test was performed as previously reported (Planaguma et al., 2015). The test measures the conflict between the natural tendency of mice to avoid an illuminated and elevated surface, and the natural tendency to explore new environments. It consisted of a black plastic cross with arms 40 cm long and 6 cm wide placed 50 cm above the floor. Two opposite arms were surrounded by walls (15 cm high, closed arms, 10 lux), while the two other arms were devoid of such walls (open arms, 200 lux). A central platform connected the four arms. At the start of the session, the mouse was placed at the end of a closed arm facing the wall. During the 5-minutes trial, the number of entries and the time spent in each arm were recorded. Anxiety was assessed as both the time spent avoiding the open arms and the number of entries into them.

#### Motor coordination

Rota rod apparatus was used to assess motor coordination. Precisely, this test measures the ability of the mouse to remain on a rotating rod in which the speed of rotation is gradually increased and evaluate general motor coordination. The device is a round drum (5 cm diameter) suspended between 2 plexiglass walls. The drum is suspended at 24 cm from a soft mat covered tabletop. The speed of rotation is controlled by an electric engine with a digital revolution per min (rpm) display. The test consisted in five trial sessions. Each trial started with the mouse being placed on the rotating rod at 4.0 rpm, and then every 3 s the speed increased by 1.0 rpm until 20.0 rpm. The trial terminated when the mouse fell from the rod or after 90 s, whichever occurred first. There was a 10 s interval between trials. The average maximum rota rod speed and time to fall was calculated for each mouse (Martin-Garcia et al., 2013).

#### Experimental procedure for food and drink monitoring

##### Habituation

Six animals per genotype were taken from their home cages and placed in experimental chambers named PHECOMP boxes (Panlab and Harvard Apparatus) equipped with a food, drink and motor activity monitoring system (https://www.panlab.com/en/products/phecomp-system-panlab) for two weeks of habituation. Animals were monitored in these boxes 24 h a day during the whole experimental period.

##### Sucrose preference test

After a period of habituation, a sucrose preference test was performed on the third week of PheComp cage housing during one week, as previously described (Bura et al., 2010; Planaguma et al., 2015). Two bottles of water, one with 2% sucrose and the other without, were placed in the cage. The position of bottles were exchanged everyday, and the consumption was measured after a 24h interval. The preference for sucrose was calculated as the relative amount of water with sucrose versus total liquid (water with and without sucrose) consumed by the mice (Bura et al., 2010).

### Operant conditioning maintained by cocaine

#### Operant self-administration apparatus

The self-administration experiments were conducted in mouse operant chambers (Model ENV-307A-CT; Med Associates Inc., Georgia, VT, USA) equipped with two holes, one randomly selected as the active hole and the other as the inactive hole as previously described (Gutierrez-Cuesta et al., 2014).

#### Operant conditioning maintained by cocaine

Cocaine self-administration experiments were performed as previously described (Gutierrez-Cuesta et al., 2014; Martín-García et al., 2015a). Mice were deeply anesthetized by intraperitoneal injection of a mixture of ketamine (75mg/kg; Imalgène; Merial Laboratorios S.A., Barcelona, Spain) and medetomidine (1mg/kg; Domtor; Esteve, Barcelona, Spain). After surgery, anesthesia was reversed by subcutaneous injection of the synthetic α2 adrenergic receptor antagonist, atipamezole (2.5mg/kg; Revertor; Virbac, Barcelona, Spain) indicated for the reversal of the sedative and analgesic effects of medetomidine (α2 adrenergic receptor agonist). In addition, mice received an intraperitoneal injection of gentamicin (1 mg/kg; Genta-Gobens; Laboratorios Normon, S.A., Madrid, Spain) along with subcutaneous administration of the analgesic meloxicam (2 mg/kg; Metacam; Boehringer Ingelheim, Rhein, Germany). Each 2h daily self-administration session started with a priming injection of the drug. Cocaine (obtained from The Spanish Agency of Drugs, Madrid, Spain) was intravenously infused in 23.5μl over 2s (0.5 mg/kg per injection). Cue light, located above the active hole, was paired with the delivery of the reinforcer. Mice (WT n=36; *Plcb1*+/- n=26) were trained under a fixed ratio 1 schedule of reinforcement (FR1; one nose-poke lead to the delivery of one dose of cocaine) over 5 consecutive daily sessions and under a fixed ratio 3 (FR3) over 5 consecutive daily sessions. Control mice trained with saline were included for both genotypes (WT n=6; *Plcb1*+/- n=6). The timeout period after infusion delivery was 10 s. Responses on the inactive hole and all responses elicited during the 10-s timeout period were also recorded. Responses during the 10-s timeout period were considered a measure of impulsivity reflecting the inability to stop motor behavior once it is initiated. The criteria for self-administration behavior were achieved when all of the following conditions were met: (1) mice maintained stable, responding with 20% deviation from the mean of the total number of reinforcers earned in three consecutive sessions (80% of stability); (2) at least 75% of mice responding on the active hole; and (3) a minimum of 10 reinforcers per session.

After the 10 FR sessions, animals were tested in a progressive ratio schedule of 4h where the response requirement to earn the cocaine escalated according to the following series: 1–2–3–5–12–18–27–40–60–90–135–200–300–450–675–1000. Then, we proceed with the extinction phase, 2h daily sessions following the same experimental conditions than the cocaine self-administration sessions except that cocaine were not available, and cue-light was not presented. Mice were given 2-h daily sessions until they achieved the extinction criterion with a maximum of 26 sessions. The criterion for extinction was achieved when, during 3 consecutive sessions, mice completed a mean number of nose pokes in the active hole consisting of 30% of the mean responses obtained during the 3 days to achieve the acquisition criteria for cocaine self-administration training. On day 27, only mice that accomplished extinction criterion were tested in the cue-induced reinstatement during a 2h session, to evaluate the reinstatement of cocaine-seeking behavior. The test for cue-induced reinstatement was conducted under the same conditions used in the training phase except that cocaine was not available. The reinstatement criterion was achieved when nose pokes in the active hole were double the number of nose pokes in the active hole during the 3 consecutive days when the mice acquired the extinction criteria.

The catheter was flushed daily with heparinized saline (30 USP units/ml). The patency of intravenous catheters was evaluated after the last cocaine self-administration session and whenever the behavior appeared to deviate dramatically from that observed previously by infusion of thiopental through the catheter. If prominent signs of anesthesia were not apparent within 3s of the infusion, the mouse was removed from the experiment. The success rate for maintaining patency of the catheter (mean duration of 11 days) until the end of the cocaine self-administration training was 90%.

After the cue-induced reinstatement, animals were euthanized by decapitation, brains were quickly removed, and the medial prefrontal cortex (mPFC) and hippocampus (HPC) were dissected. Brain tissues were then frozen by immersion in 2-methylbutane surrounded by dry ice and stored at −80°C for later RNA isolation and transcriptomic analyses.

### RNA extraction and RNA sequencing

Total RNA of 12 WT and 11 *Plcb1*+/- mice from mPFC and HPC were isolated using the RNeasy Lipid Tissue Mini Kit (Qiagen Düsseldorf, Germany) according to the manufacturer’s protocol. RNA concentration was determined using the NanoDrop ND-1000 spectrophotometer (NanoDrop Technologies, Wilmington, DE, USA), and integrity was evaluated using the Bioanalyzer2100 platform (Agilent Technologies, Santa Clara, CA, USA). RNA samples were grouped in 4 pools consisting of 3 mice per pool for each experimental group, except for 1 group in the *Plcb1*+/- mice were only 2 animals were pooled. The pools were organized to homogenize the average number of nose pokes in the different pools. The pooled individuals were the same for both mPFC and HPC.

### RNA sequencing

RNA sequencing (RNAseq) was performed by the Centre de Regulació Genòmica (CRG, Barcelona, Spain). Libraries were prepared using the TruSeq Stranded mRNA Sample Prep Kit_v2 (Illumina, San Diego, CA, USA) according to the manufacturer’s protocol and sequenced 2×75 on Illumina’s HiSeq3000 system for both mPFC and HPC. The Bioinformatics service of CRG carried out the analysis of RNAseq. Briefly, FastQC v0.11.5(Andrews S., 2010) was used to inspect the reads quality and CutAdapt 1.7.1 (Martin, 2011) to clean the data of adapters and low-quality reads. Then, reads were mapped to the *Mus musculus* genome of reference (GRCm38/mm10) with STAR 2.5.3a (Dobin et al., 2013), and the differential expression analysis was done by DESeq2 (Love et al., 2014a) to compare WT and *Plcb1+/-o* Corrections for multiple testing were applied by adjusting the p-values with a 5% False Discovery Rate.

RNAseq data of mPFC and HPC were explored on a principal component analysis (PCA) plot using the “plotPCA” method from the DESeq2 package (Love et al., 2014b) and log2 gene expression data. The PCAs were performed with the 500 genes showing the highest variance among the samples to calculate the distance among them. The heatmaps were performed using the “heatmap” function on R, and the hierarchical clustering considered the euclidean distance between the samples considering all the genes or only those with corrected p-value < 1e-05.

### Functional annotation of RNAseq results

We performed a functional group enrichment of differentially expressed genes (DEGs) in mPFC using the DAVID Annotation Tool (http://david.abcc.ncifcrf.gov)(Jiao et al., 2012) considering GO (Gene Ontology) biological processes and KEGG pathways (Kyoto Encyclopedia of Genes and Genomes). Then, we searched for over-represented transcription factor-binding sites (TFBS) using the information of MsigDB (https://www.gsea-msigdb.org/gsea/msigdb) integrated on WebGestalt2Ol9 (http://www.webgestalt.org/)(Liao et al., 2019), and the default parameters, applying the weighted set cover method to reduce redundancy. In both analyses, the Benjamin–Hochberg procedure was performed for multiple testing. Finally, we investigated the existence of gene networks with Ingenuity Pathway Analysis 8.8 software (IPA, http://www.ingenuity.com/products/ipa; Ingenuity Systems, Redwood City, CA, USA) (Krämer et al., 2014) after selecting genes with fold-change > |1.2|.

## Statistical analysis

Three-way ANOVA with repeated measures was used to test the evolution over sessions or days. Sessions or days were used as within-subject factors and genotype (*Plcb1*+/- or WT) and drug (cocaine or saline) were used as between-subjects factors. Post-hoc analyses (Newman-Keuls) were performed when required. Comparisons between two groups were analyzed by Student *t*-test or *U*-Mann-Whitney depending on the distribution defined by the Kolmogorov-Smirnov normality test and the sample size. The chi-square analyses were performed to compare the percentage of mice that acquired operant learning criteria in the different experimental groups. Results are expressed as mean ± SEM or individual values with the median and the interquartile range specified in the figure legend. Differences were considered significant at P<0.05. The sample size was calculated based on power analysis. The statistical analyses were performed using the Statistical Package for Social Science tool SPSS®25.0 (SPSS Inc, Chicago, USA).

See supplementary information for more details of genotyping of transgenic mice, drugs, behavioral tests of phenotype characterization, operant conditioning maintained by cocaine, RNA extraction and sequencing.

## ACKNOWLEDGEMENTS AND FUNDING

We are grateful to H.-S. Shin for providing breeder mice to establish the colony. We thank M. Linares, R. Martín, D. Real, F. Porrón for their technical support, and the Genomics Unit at the CRG for assistance with the mRNAseq.

This work was supported by the Spanish ‘Ministerio de Economía y Competitividad-MINECO’ (#SAF2017-84060-R-AEI/FEDER-UE), the Spanish ‘Instituto de Salud Carlos III, RETICS-RTA’ (#RD12/0028/0023), the ‘Generalitat de Catalunya, AGAUR’ (#2017 SGR-669), ‘ICREA-Acadèmia’ (#2015) and the Spanish ‘Ministerio de Sanidad, Servicios Sociales e Igualdad, ‘Plan Nacional Sobre Drogas of the Spanish Ministry of Health’ (#PNSD-2017I068) to RM, ‘Fundació La Marató-TV3’ (#2016/20-30) and ‘Plan Nacional Sobre Drogas of the Spanish Ministry of Health’ (#PNSD-2019I006) to E.M-G., Spanish ‘Ministerio de Ciencia, Innovación y Universidades^1^ (#RTI2018-100968-B-100), Spanish ‘Ministerio de Economía y Competitividad’ (#SAF2015-68341-R), ‘AGAUR-Generalitat de Catalunya’ (#2017-SGR-738) and Spanish ‘Ministerio de Sanidad, Servicios Sociales e Igualdad, ‘Plan Nacional Sobre Drogas of the Spanish Ministry of Health’ (#PNSD-2017I050) to BC. The research leading to these results has also received funding from the European Union H2020 Program [H2020/2014-2020] under grant agreements n° 667302 (CoCA), 643051 (MiND) and 728018 (Eat2beNICE) to BC. JC-D was supported by the H2020 CoCA and Eat2beNICE projects and NF-C by ‘Centro de Investigación Biomédica en Red de Enfermedades Raras’ (CIBERER).

J.C-D, B.C. and N.F.-C. obtained and maintained the mutant mice (*Plcb1+/-);* E.M.-G. and R.M. conceived and designed the behavioral studies with input from J.C.-D, B.C. and N. F.-C; E.M.-G. and J.C.-D. performed the behavioral phenotype characterization of mutant mice. E.M.-G. performed the surgery for i.v. catheterization and the operant conditioning maintained by cocaine. A.G.-R. collaborated in extinction and cue-induced reinstatement. E.M.-G. and J.C.-D. performed statistical analyses and graphs with the supervision of R.M., B.C. and N.F.-C.; A.G.R. collaborated with the extraction of the samples; J.C.-D. performed the RNA extractions, RNA sequencing, and the bioinformatic analyses supervised by B.C. and N.F.-C.; E.M.-G. and J.C.-D. wrote the manuscript and N.F.C., R.M. and B.C. provided a critical review of the manuscript with inputs from all the other authors.

We have uploaded it to the preprint server bioRxiv (https://doi.org/10.1101/2020.06.18.158964).

## DISCLOURES

The authors have no conflicts of interest.

