## Supplementary Tables for "Reduced cue-induced reinstatement of cocaine-seeking behavior in Plcb1+/- mice"

**Supplementary Table S1.** Operant responding maintained by cocaine during FR1 and FR3 (infusions and active nose-pokes) and extinction (active nose-pokes).

| | Three-way ANOVA | | | |
|---|---|---|---|---|
| | FR1 Infusions | | FR3 Infusions | |
| | *F*-value | *P*-value | *F*-value | *P*-value |
| Genotype | $F_{(1,70)} = 0.40$ | *n.s.* | $F_{(1,70)} = 0.02$ | *n.s.* |
| Drug | $F_{(1,70)} = 5.91$ | $P < 0.05$ | $F_{(1,70)} = 12.78$ | $P < 0.001$ |
| Sessions | $F_{(4,280)} = 9.15$ | $P < 0.001$ | $F_{(4,280)} = 3.34$ | $P < 0.05$ |
| Genotype × Drug | $F_{(1,70)} = 2.41$ | *n.s.* | $F_{(1,70)} = 0.02$ | *n.s.* |
| Genotype × Sessions | $F_{(4,280)} = 3.80$ | $P < 0.01$ | $F_{(4,280)} = 0.30$ | *n.s.* |
| Drug × Sessions | $F_{(4,280)} = 5.25$ | $P < 0.01$ | $F_{(4,280)} = 0.23$ | *n.s.* |
| Genotype × Drug × Sessions | $F_{(4,280)} = 2.81$ | $P < 0.05$ | $F_{(4,280)} = 0.81$ | *n.s.* |

| | FR1 Active nose-pokes | | FR3 Active-nose-pokes | |
|---|---|---|---|---|
| | *F*-value | *P*-value | *F*-value | *P*-value |
| Genotype | $F_{(1,70)} = 0.86$ | *n.s.* | $F_{(1,70)} = 0.10$ | *n.s.* |
| Drug | $F_{(1,70)} = 4.96$ | $P < 0.05$ | $F_{(1,70)} = 9.77$ | $P < 0.01$ |
| Sessions | $F_{(4,280)} = 3.99$ | $P < 0.01$ | $F_{(4,280)} = 3.59$ | $P < 0.01$ |
| Genotype × Drug | $F_{(1,70)} = 1.70$ | *n.s.* | $F_{(1,70)} = 0.08$ | *n.s.* |
| Genotype × Sessions | $F_{(4,280)} = 1.92$ | *n.s.* | $F_{(4,280)} = 0.66$ | *n.s.* |
| Drug × Sessions | $F_{(4,280)} = 3.62$ | $P < 0.01$ | $F_{(4,280)} = 1.36$ | *n.s.* |
| Genotype × Drug × Sessions | $F_{(4,280)} = 1.96$ | *n.s.* | $F_{(4,280)} = 0.85$ | *n.s.* |

| | Extinction | |
|---|---|---|
| | *F*-value | *P*-value |
| Genotype | $F_{(1,55)} = 2.94$ | *n.s.* |
| Drug | $F_{(1,55)} = 8.28$ | $P < 0.01$ |
| Sessions | $F_{(25,1375)} = 41.65$ | $P < 0.001$ |
| Genotype × Drug | $F_{(1,55)} = 0.55$ | *n.s.* |
| Genotype × Sessions | $F_{(25,1375)} = 1.18$ | $P < 0.001$ |
| Drug × Sessions | $F_{(25,1375)} = 4.09$ | *n.s.* |
| Genotype × Drug × Sessions | $F_{(25,1375)} = 0.59$ | *n.s.* |

Three-way ANOVA with genotype and drug as between-subjects factor and repeated measures in the factor sessions. See materials and methods for details. *n.s.:* non significant

**Supplementary Table S2.** Gene expression changes in mPFC (FDR 5%) in Plcb1+/- mice compared to WT after cocaine seeking reinstatement

| **ids** | **baseMean** | **log2FoldChange** | **lfcSE** | **stat** | **pvalue** | **padj** | **Gene Name** | **Gene type** |
|---|---|---|---|---|---|---|---|---|
| ENSMUSG00000095098.1 | 2428.060 | -0.937 | 0.055 | -16.952 | 1.86E-64 | 2.63E-60 | Ccdc85b | protein_coding |
| ENSMUSG00000071076.6 | 2879.048 | -0.949 | 0.060 | -15.696 | 1.60E-55 | 1.13E-51 | Jund | protein_coding |
| ENSMUSG00000020496.9 | 12161.061 | -0.348 | 0.025 | -13.977 | 2.16E-44 | 1.01E-40 | Rnf187 | protein_coding |
| ENSMUSG00000070493.3 | 4689.425 | -0.441 | 0.035 | -12.762 | 2.66E-37 | 9.37E-34 | Chchd2 | protein_coding |
| ENSMUSG00000060032.6 | 950.218 | -0.648 | 0.055 | -11.849 | 2.17E-32 | 6.11E-29 | H2afj | protein_coding |
| ENSMUSG00000044533.15 | 8067.372 | -0.366 | 0.032 | -11.564 | 6.26E-31 | 1.47E-27 | Rps2 | protein_coding |
| ENSMUSG00000054871.5 | 1236.200 | -0.652 | 0.057 | -11.506 | 1.23E-30 | 2.47E-27 | Tmem158 | protein_coding |
| ENSMUSG00000070462.4 | 1279.727 | -0.534 | 0.047 | -11.467 | 1.92E-30 | 3.39E-27 | Mesdc1 | protein_coding |
| ENSMUSG00000040860.16 | 7894.329 | -0.375 | 0.033 | -11.245 | 2.44E-29 | 3.82E-26 | Crocc | protein_coding |
| ENSMUSG00000047085.14 | 6553.856 | -0.469 | 0.042 | -11.236 | 2.72E-29 | 3.84E-26 | Lrrc4b | protein_coding |
| ENSMUSG00000039278.10 | 10220.556 | -0.820 | 0.073 | -11.225 | 3.06E-29 | 3.92E-26 | Pcsk1n | protein_coding |
| ENSMUSG00000078794.4 | 4932.143 | -0.451 | 0.040 | -11.179 | 5.19E-29 | 6.09E-26 | Dact3 | protein_coding |
| ENSMUSG00000029581.14 | 8409.185 | -0.339 | 0.031 | -11.055 | 2.08E-28 | 2.25E-25 | Fscn1 | protein_coding |
| ENSMUSG00000002393.14 | 903.882 | -0.717 | 0.065 | -10.993 | 4.14E-28 | 4.02E-25 | Nr2f6 | protein_coding |
| ENSMUSG00000034685.4 | 3656.144 | -0.459 | 0.042 | -10.990 | 4.28E-28 | 4.02E-25 | Fam171a2 | protein_coding |
| ENSMUSG00000049932.3 | 977.087 | -0.655 | 0.060 | -10.936 | 7.72E-28 | 6.80E-25 | H2afx | protein_coding |
| ENSMUSG00000000740.12 | 4710.356 | -0.556 | 0.051 | -10.823 | 2.69E-27 | 2.23E-24 | Rpl13 | protein_coding |
| ENSMUSG00000043165.7 | 161.243 | -0.817 | 0.077 | -10.627 | 2.22E-26 | 1.74E-23 | Lor | protein_coding |
| ENSMUSG00000034686.7 | 865.971 | -0.716 | 0.068 | -10.476 | 1.11E-25 | 8.26E-23 | Prr7 | protein_coding |
| ENSMUSG00000049422.7 | 3632.917 | -0.545 | 0.052 | -10.421 | 2.00E-25 | 1.41E-22 | Chchd10 | protein_coding |
| ENSMUSG00000054716.4 | 810.956 | -0.763 | 0.074 | -10.243 | 1.27E-24 | 8.55E-22 | Zfp771 | protein_coding |
| ENSMUSG00000039860.19 | 3319.619 | -0.349 | 0.034 | -10.137 | 3.79E-24 | 2.43E-21 | Srrm3 | protein_coding |
| ENSMUSG00000043445.6 | 1473.625 | -0.597 | 0.059 | -10.056 | 8.66E-24 | 5.31E-21 | Pgp | protein_coding |
| ENSMUSG00000022577.16 | 7170.974 | -0.418 | 0.043 | -9.842 | 7.43E-23 | 4.37E-20 | Ly6h | protein_coding |
| ENSMUSG00000025226.11 | 640.336 | -0.617 | 0.063 | -9.833 | 8.08E-23 | 4.56E-20 | Fbxl15 | protein_coding |
| ENSMUSG00000013858.14 | 4459.271 | -0.349 | 0.036 | -9.648 | 5.00E-22 | 2.71E-19 | Tmem259 | protein_coding |
| ENSMUSG00000035885.4 | 9047.900 | -0.409 | 0.042 | -9.622 | 6.48E-22 | 3.38E-19 | Cox8a | protein_coding |
| ENSMUSG00000040794.5 | 2722.233 | -0.701 | 0.074 | -9.506 | 1.97E-21 | 9.62E-19 | C1qtnf4 | protein_coding |
| ENSMUSG00000075227.6 | 616.338 | -0.552 | 0.058 | -9.506 | 1.98E-21 | 9.62E-19 | Znhit2 | protein_coding |
| ENSMUSG00000064264.14 | 891.665 | -0.555 | 0.059 | -9.430 | 4.09E-21 | 1.92E-18 | Zfp428 | protein_coding |

| | | | | | | | | |
|---|---|---|---|---|---|---|---|---|
| ENSMUSG00000040867.12 | 5995.125 | -0.398 | 0.042 | -9.415 | 4.73E-21 | 2.15E-18 | Begain | protein_coding |
| ENSMUSG00000051550.10 | 1024.796 | -0.576 | 0.062 | -9.368 | 7.36E-21 | 3.24E-18 | Zfp579 | protein_coding |
| ENSMUSG00000002985.16 | 41842.922 | -0.360 | 0.038 | -9.362 | 7.80E-21 | 3.33E-18 | Apoe | protein_coding |
| ENSMUSG00000035342.13 | 2569.807 | -0.365 | 0.039 | -9.294 | 1.48E-20 | 5.98E-18 | Lzts2 | protein_coding |
| ENSMUSG00000007944.8 | 2889.222 | -0.418 | 0.045 | -9.294 | 1.49E-20 | 5.98E-18 | Ttc9b | protein_coding |
| ENSMUSG00000070858.12 | 358.331 | -0.665 | 0.073 | -9.095 | 9.45E-20 | 3.66E-17 | Gm1673 | protein_coding |
| ENSMUSG00000030600.15 | 2588.959 | -0.433 | 0.048 | -9.093 | 9.60E-20 | 3.66E-17 | Lrfn1 | protein_coding |
| ENSMUSG00000044030.4 | 1827.063 | -0.360 | 0.040 | -9.035 | 1.65E-19 | 6.11E-17 | Irf2bp1 | protein_coding |
| ENSMUSG00000071341.4 | 1550.943 | -0.693 | 0.077 | -9.002 | 2.22E-19 | 8.03E-17 | Egr4 | protein_coding |
| ENSMUSG00000090071.4 | 8832.339 | -0.379 | 0.042 | -8.989 | 2.49E-19 | 8.77E-17 | Cdk5r2 | protein_coding |
| ENSMUSG00000050896.12 | 1472.949 | -0.506 | 0.057 | -8.896 | 5.77E-19 | 1.98E-16 | Rtn4rl2 | protein_coding |
| ENSMUSG00000046229.9 | 432.648 | -0.664 | 0.075 | -8.832 | 1.03E-18 | 3.45E-16 | Scand1 | protein_coding |
| ENSMUSG00000032174.5 | 12025.867 | -0.339 | 0.038 | -8.813 | 1.22E-18 | 4.01E-16 | Icam5 | protein_coding |
| ENSMUSG00000027447.6 | 26537.549 | -0.248 | 0.028 | -8.792 | 1.46E-18 | 4.69E-16 | Cst3 | protein_coding |
| ENSMUSG00000029419.8 | 6653.533 | -0.321 | 0.037 | -8.762 | 1.92E-18 | 6.02E-16 | Gm996 | protein_coding |
| ENSMUSG00000046215.3 | 1286.568 | -0.536 | 0.061 | -8.717 | 2.86E-18 | 8.75E-16 | Rprml | protein_coding |
| ENSMUSG00000089762.3 | 692.006 | -0.562 | 0.065 | -8.705 | 3.17E-18 | 9.51E-16 | Ier5l | protein_coding |
| ENSMUSG00000070003.7 | 3940.817 | -0.464 | 0.053 | -8.679 | 4.01E-18 | 1.18E-15 | Ssbp4 | protein_coding |
| ENSMUSG00000035011.15 | 6673.966 | -0.360 | 0.042 | -8.566 | 1.07E-17 | 3.07E-15 | Zbtb7a | protein_coding |
| ENSMUSG00000037499.9 | 1132.999 | -0.391 | 0.046 | -8.558 | 1.15E-17 | 3.25E-15 | Nenf | protein_coding |
| ENSMUSG00000020308.6 | 1119.688 | -0.545 | 0.064 | -8.539 | 1.35E-17 | 3.73E-15 | Tpgs1 | protein_coding |
| ENSMUSG00000049036.7 | 489.513 | -0.564 | 0.066 | -8.511 | 1.72E-17 | 4.67E-15 | Tmem121 | protein_coding |
| ENSMUSG00000042388.14 | 14098.397 | -0.368 | 0.043 | -8.506 | 1.79E-17 | 4.77E-15 | Dlgap3 | protein_coding |
| ENSMUSG00000105867.2 | 1502.706 | -0.393 | 0.046 | -8.493 | 2.02E-17 | 5.27E-15 | Gm42517 | protein_coding |
| ENSMUSG00000011751.16 | 14000.264 | -0.373 | 0.044 | -8.434 | 3.33E-17 | 8.53E-15 | Sptbn4 | protein_coding |
| ENSMUSG00000035828.10 | 1592.059 | -0.365 | 0.043 | -8.422 | 3.70E-17 | 9.32E-15 | Pim3 | protein_coding |
| ENSMUSG00000001227.11 | 5284.242 | -0.295 | 0.035 | -8.377 | 5.41E-17 | 1.34E-14 | Sema6b | protein_coding |
| ENSMUSG00000035585.16 | 2037.478 | -0.331 | 0.040 | -8.355 | 6.53E-17 | 1.59E-14 | Tsen34 | protein_coding |
| ENSMUSG00000054364.5 | 7912.063 | -0.257 | 0.031 | -8.336 | 7.71E-17 | 1.84E-14 | Rhob | protein_coding |
| ENSMUSG00000043439.5 | 2609.601 | -0.491 | 0.059 | -8.327 | 8.28E-17 | 1.95E-14 | Epop | protein_coding |
| ENSMUSG00000024873.6 | 6806.672 | -0.278 | 0.033 | -8.317 | 9.02E-17 | 2.08E-14 | Cnih2 | protein_coding |
| ENSMUSG00000044927.6 | 359.776 | -0.527 | 0.064 | -8.183 | 2.77E-16 | 6.29E-14 | H1fx | protein_coding |
| ENSMUSG00000003380.11 | 3581.323 | -0.265 | 0.033 | -8.126 | 4.46E-16 | 9.97E-14 | Rabac1 | protein_coding |

| | | | | | | | | |
|---|---|---|---|---|---|---|---|---|
| ENSMUSG00000028478.18 | 6237.142 | -0.283 | 0.035 | -8.112 | 5.00E-16 | 1.10E-13 | Clta | protein_coding |
| ENSMUSG00000048967.16 | 448.124 | -0.537 | 0.066 | -8.092 | 5.86E-16 | 1.27E-13 | Yjefn3 | protein_coding |
| ENSMUSG00000030685.5 | 3468.793 | -0.267 | 0.033 | -8.073 | 6.84E-16 | 1.46E-13 | Kctd13 | protein_coding |
| ENSMUSG00000042675.15 | 6175.722 | -0.336 | 0.042 | -8.069 | 7.12E-16 | 1.50E-13 | Ypel3 | protein_coding |
| ENSMUSG00000041697.8 | 6569.479 | -0.313 | 0.039 | -8.067 | 7.23E-16 | 1.50E-13 | Cox6a1 | protein_coding |
| ENSMUSG00000033467.12 | 518.566 | -0.465 | 0.058 | -8.011 | 1.13E-15 | 2.32E-13 | Crlf2 | protein_coding |
| ENSMUSG00000052684.4 | 2489.177 | -0.333 | 0.042 | -8.004 | 1.20E-15 | 2.42E-13 | Jun | protein_coding |
| ENSMUSG00000036422.10 | 2306.449 | -0.400 | 0.050 | -7.956 | 1.77E-15 | 3.52E-13 | Pcdh8 | protein_coding |
| ENSMUSG00000078235.3 | 1323.520 | -0.411 | 0.052 | -7.933 | 2.14E-15 | 4.20E-13 | Fam43b | protein_coding |
| ENSMUSG00000011096.17 | 1640.762 | -0.410 | 0.052 | -7.903 | 2.71E-15 | 5.24E-13 | Akt1s1 | protein_coding |
| ENSMUSG00000019261.3 | 3279.321 | -0.348 | 0.044 | -7.885 | 3.15E-15 | 5.93E-13 | Map1s | protein_coding |
| ENSMUSG00000015476.6 | 6427.574 | -0.286 | 0.036 | -7.885 | 3.16E-15 | 5.93E-13 | Prrt1 | protein_coding |
| ENSMUSG00000052981.6 | 9679.894 | -0.240 | 0.030 | -7.878 | 3.32E-15 | 6.17E-13 | Ube2ql1 | protein_coding |
| ENSMUSG00000037236.14 | 14697.288 | 0.240 | 0.031 | 7.849 | 4.21E-15 | 7.70E-13 | Matr3 | protein_coding |
| ENSMUSG00000007950.9 | 10690.024 | -0.328 | 0.042 | -7.809 | 5.76E-15 | 1.04E-12 | Abhd8 | protein_coding |
| ENSMUSG00000028180.18 | 9904.013 | 0.178 | 0.023 | 7.796 | 6.38E-15 | 1.14E-12 | Zranb2 | protein_coding |
| ENSMUSG00000090877.3 | 687.482 | -0.555 | 0.071 | -7.784 | 7.01E-15 | 1.23E-12 | Hspa1b | protein_coding |
| ENSMUSG00000033981.14 | 36232.451 | 0.181 | 0.023 | 7.760 | 8.46E-15 | 1.47E-12 | Gria2 | protein_coding |
| ENSMUSG00000038976.12 | 39823.755 | -0.222 | 0.029 | -7.757 | 8.67E-15 | 1.49E-12 | Ppp1r9b | protein_coding |
| ENSMUSG00000003072.15 | 6475.690 | -0.255 | 0.033 | -7.743 | 9.71E-15 | 1.65E-12 | Atp5d | protein_coding |
| ENSMUSG00000003346.14 | 4043.632 | -0.330 | 0.043 | -7.741 | 9.85E-15 | 1.65E-12 | Abhd17a | protein_coding |
| ENSMUSG00000034201.13 | 3477.939 | -0.333 | 0.043 | -7.740 | 9.97E-15 | 1.65E-12 | Gas2l1 | protein_coding |
| ENSMUSG00000004366.3 | 7344.551 | -0.371 | 0.048 | -7.681 | 1.57E-14 | 2.58E-12 | Sst | protein_coding |
| ENSMUSG00000020087.5 | 711.408 | -0.418 | 0.054 | -7.679 | 1.60E-14 | 2.59E-12 | Tysnd1 | protein_coding |
| ENSMUSG00000041801.5 | 908.207 | -0.382 | 0.050 | -7.670 | 1.72E-14 | 2.76E-12 | Phlda3 | protein_coding |
| ENSMUSG00000000088.7 | 3521.405 | -0.289 | 0.038 | -7.662 | 1.83E-14 | 2.90E-12 | Cox5a | protein_coding |
| ENSMUSG00000002984.17 | 3028.449 | -0.265 | 0.035 | -7.609 | 2.76E-14 | 4.32E-12 | Tomm40 | protein_coding |
| ENSMUSG00000030120.14 | 15767.863 | -0.222 | 0.029 | -7.592 | 3.16E-14 | 4.89E-12 | Mlf2 | protein_coding |
| ENSMUSG00000030811.14 | 4286.799 | -0.243 | 0.032 | -7.567 | 3.81E-14 | 5.83E-12 | Fbxl19 | protein_coding |
| ENSMUSG00000042978.10 | 5699.351 | -0.276 | 0.037 | -7.539 | 4.73E-14 | 7.17E-12 | Sbk1 | protein_coding |
| ENSMUSG00000034854.8 | 1759.471 | -0.296 | 0.039 | -7.511 | 5.86E-14 | 8.78E-12 | Mfsd12 | protein_coding |
| ENSMUSG00000052584.9 | 1884.408 | -0.279 | 0.037 | -7.454 | 9.06E-14 | 1.34E-11 | Serp2 | protein_coding |
| ENSMUSG00000027677.17 | 10489.396 | 0.167 | 0.022 | 7.435 | 1.05E-13 | 1.54E-11 | Ttc14 | protein_coding |

| | | | | | | | | |
|---|---|---|---|---|---|---|---|---|
| ENSMUSG00000048481.15 | 1577.978 | -0.381 | 0.051 | -7.433 | 1.06E-13 | 1.54E-11 | Mypop | protein_coding |
| ENSMUSG00000037979.13 | 3574.295 | -0.243 | 0.033 | -7.422 | 1.15E-13 | 1.65E-11 | Ccdc92 | protein_coding |
| ENSMUSG00000022199.10 | 30829.541 | -0.261 | 0.035 | -7.422 | 1.16E-13 | 1.65E-11 | Slc22a17 | protein_coding |
| ENSMUSG00000028849.17 | 12137.928 | -0.238 | 0.032 | -7.415 | 1.22E-13 | 1.71E-11 | Map7d1 | protein_coding |
| ENSMUSG00000020436.17 | 7711.715 | 0.198 | 0.027 | 7.391 | 1.46E-13 | 2.02E-11 | Gabrg2 | protein_coding |
| ENSMUSG00000059991.7 | 2598.656 | -0.319 | 0.043 | -7.376 | 1.63E-13 | 2.23E-11 | Nptx2 | protein_coding |
| ENSMUSG00000031029.6 | 3853.029 | -0.253 | 0.034 | -7.335 | 2.22E-13 | 3.01E-11 | Eif3f | protein_coding |
| ENSMUSG00000025422.9 | 55624.113 | -0.251 | 0.034 | -7.300 | 2.88E-13 | 3.87E-11 | Agap2 | protein_coding |
| ENSMUSG00000033730.3 | 4336.244 | -0.379 | 0.052 | -7.271 | 3.57E-13 | 4.75E-11 | Egr3 | protein_coding |
| ENSMUSG00000046160.6 | 3270.411 | -0.395 | 0.054 | -7.268 | 3.66E-13 | 4.82E-11 | Olig1 | protein_coding |
| ENSMUSG00000033751.5 | 1164.392 | -0.343 | 0.047 | -7.258 | 3.92E-13 | 5.11E-11 | Gadd45gip1 | protein_coding |
| ENSMUSG00000008140.17 | 7488.264 | -0.245 | 0.034 | -7.242 | 4.41E-13 | 5.71E-11 | Emc10 | protein_coding |
| ENSMUSG00000037428.14 | 10345.284 | -0.378 | 0.052 | -7.227 | 4.93E-13 | 6.32E-11 | Vgf | protein_coding |
| ENSMUSG00000020477.10 | 538.020 | -0.438 | 0.061 | -7.226 | 4.98E-13 | 6.33E-11 | Mrps24 | protein_coding |
| ENSMUSG00000022769.7 | 728.806 | -0.443 | 0.061 | -7.200 | 6.01E-13 | 7.49E-11 | Sdf2l1 | protein_coding |
| ENSMUSG00000030678.7 | 7128.976 | -0.231 | 0.032 | -7.172 | 7.39E-13 | 9.13E-11 | Maz | protein_coding |
| ENSMUSG00000058833.11 | 931.967 | -0.369 | 0.051 | -7.167 | 7.69E-13 | 9.42E-11 | 2810428I15Rik | protein_coding |
| ENSMUSG00000034730.16 | 27122.310 | -0.276 | 0.039 | -7.128 | 1.02E-12 | 1.24E-10 | Adgrb1 | protein_coding |
| ENSMUSG00000061451.13 | 11977.343 | -0.251 | 0.035 | -7.088 | 1.36E-12 | 1.64E-10 | Tmem151a | protein_coding |
| ENSMUSG00000041740.16 | 11504.920 | -0.149 | 0.021 | -7.074 | 1.51E-12 | 1.80E-10 | Rnf10 | protein_coding |
| ENSMUSG00000024614.6 | 3589.782 | 0.224 | 0.032 | 7.069 | 1.56E-12 | 1.85E-10 | Tmx3 | protein_coding |
| ENSMUSG00000020331.9 | 4651.453 | -0.258 | 0.037 | -7.038 | 1.95E-12 | 2.29E-10 | Hcn2 | protein_coding |
| ENSMUSG00000048644.8 | 13242.837 | -0.200 | 0.028 | -7.026 | 2.13E-12 | 2.48E-10 | Ctxn1 | protein_coding |
| ENSMUSG00000055692.18 | 4008.625 | -0.274 | 0.039 | -6.977 | 3.01E-12 | 3.47E-10 | Tmem191c | protein_coding |
| ENSMUSG00000055302.5 | 7188.557 | -0.217 | 0.031 | -6.975 | 3.07E-12 | 3.52E-10 | Mrfap1 | protein_coding |
| ENSMUSG00000029603.15 | 3810.347 | -0.304 | 0.044 | -6.973 | 3.11E-12 | 3.52E-10 | Dtx1 | protein_coding |
| ENSMUSG00000096847.1 | 8989.196 | -0.249 | 0.036 | -6.972 | 3.12E-12 | 3.52E-10 | Tmem151b | protein_coding |
| ENSMUSG00000044876.15 | 1459.512 | -0.308 | 0.044 | -6.971 | 3.15E-12 | 3.52E-10 | Zfp444 | protein_coding |
| ENSMUSG00000004951.10 | 347.637 | -0.536 | 0.077 | -6.968 | 3.21E-12 | 3.56E-10 | Hspb1 | protein_coding |
| ENSMUSG00000034863.9 | 3777.063 | -0.246 | 0.035 | -6.962 | 3.36E-12 | 3.70E-10 | Ano8 | protein_coding |
| ENSMUSG00000040972.8 | 3318.215 | -0.300 | 0.043 | -6.945 | 3.78E-12 | 4.13E-10 | Igsf21 | protein_coding |
| ENSMUSG00000107002.1 | 1016.151 | -0.440 | 0.064 | -6.928 | 4.27E-12 | 4.60E-10 | 0610012G03Rik | protein_coding |
| ENSMUSG00000033735.9 | 929.202 | -0.358 | 0.052 | -6.928 | 4.28E-12 | 4.60E-10 | Spr | protein_coding |

| | | | | | | | | |
|---|---|---|---|---|---|---|---|---|
| ENSMUSG00000015094.16 | 8669.392 | -0.187 | 0.027 | -6.927 | 4.31E-12 | 4.60E-10 | Npdc1 | protein_coding |
| ENSMUSG00000052253.5 | 1244.723 | -0.330 | 0.048 | -6.922 | 4.46E-12 | 4.72E-10 | Zfp622 | protein_coding |
| ENSMUSG00000024790.7 | 351.811 | -0.469 | 0.068 | -6.906 | 4.99E-12 | 5.25E-10 | Sac3d1 | protein_coding |
| ENSMUSG00000021816.10 | 14173.920 | 0.181 | 0.026 | 6.855 | 7.13E-12 | 7.39E-10 | Ppp3cb | protein_coding |
| ENSMUSG00000034168.7 | 3658.536 | -0.354 | 0.052 | -6.843 | 7.73E-12 | 7.95E-10 | Irf2bpl | protein_coding |
| ENSMUSG00000073131.12 | 2191.714 | 0.247 | 0.036 | 6.827 | 8.65E-12 | 8.84E-10 | Vma21 | protein_coding |
| ENSMUSG00000051403.9 | 8650.197 | -0.249 | 0.037 | -6.805 | 1.01E-11 | 1.02E-09 | Ppp1r37 | protein_coding |
| ENSMUSG00000052135.8 | 733.321 | -0.424 | 0.062 | -6.802 | 1.03E-11 | 1.04E-09 | Foxo6 | protein_coding |
| ENSMUSG00000037843.6 | 1653.770 | -0.429 | 0.063 | -6.793 | 1.10E-11 | 1.10E-09 | Vstm2l | protein_coding |
| ENSMUSG00000000838.17 | 3510.761 | 0.217 | 0.032 | 6.791 | 1.11E-11 | 1.11E-09 | Fmr1 | protein_coding |
| ENSMUSG00000007836.6 | 3841.505 | -0.282 | 0.042 | -6.785 | 1.16E-11 | 1.15E-09 | Hnrnpa0 | protein_coding |
| ENSMUSG00000040390.13 | 7779.121 | -0.286 | 0.042 | -6.784 | 1.17E-11 | 1.15E-09 | Map3k10 | protein_coding |
| ENSMUSG00000064115.13 | 9132.273 | 0.203 | 0.030 | 6.761 | 1.37E-11 | 1.33E-09 | Cadm2 | protein_coding |
| ENSMUSG00000050138.7 | 285.223 | -0.517 | 0.077 | -6.754 | 1.44E-11 | 1.39E-09 | Kcnk12 | protein_coding |
| ENSMUSG00000049086.8 | 1317.001 | -0.429 | 0.064 | -6.733 | 1.66E-11 | 1.59E-09 | Bmyc | protein_coding |
| ENSMUSG00000039615.8 | 4325.493 | -0.208 | 0.031 | -6.728 | 1.72E-11 | 1.64E-09 | Stub1 | protein_coding |
| ENSMUSG00000015733.13 | 9149.727 | 0.160 | 0.024 | 6.701 | 2.07E-11 | 1.96E-09 | Capza2 | protein_coding |
| ENSMUSG00000030357.10 | 5983.919 | -0.243 | 0.036 | -6.681 | 2.37E-11 | 2.22E-09 | Fkbp4 | protein_coding |
| ENSMUSG00000041774.14 | 824.274 | -0.345 | 0.052 | -6.661 | 2.73E-11 | 2.55E-09 | Ydjc | protein_coding |
| ENSMUSG00000029725.10 | 261.519 | -0.493 | 0.074 | -6.629 | 3.38E-11 | 3.14E-09 | Ppp1r35 | protein_coding |
| ENSMUSG00000078572.3 | 288.010 | -0.446 | 0.067 | -6.620 | 3.59E-11 | 3.31E-09 | Ndufaf8 | protein_coding |
| ENSMUSG00000022564.6 | 16238.624 | -0.160 | 0.024 | -6.611 | 3.81E-11 | 3.48E-09 | Grina | protein_coding |
| ENSMUSG00000032532.7 | 7293.557 | -0.486 | 0.074 | -6.607 | 3.93E-11 | 3.58E-09 | Cck | protein_coding |
| ENSMUSG00000032328.12 | 18240.021 | 0.163 | 0.025 | 6.600 | 4.12E-11 | 3.72E-09 | Tmem30a | protein_coding |
| ENSMUSG00000061479.15 | 2394.735 | -0.254 | 0.039 | -6.598 | 4.17E-11 | 3.74E-09 | Snrpa | protein_coding |
| ENSMUSG00000039568.6 | 4783.271 | -0.225 | 0.034 | -6.585 | 4.55E-11 | 4.06E-09 | Ubald1 | protein_coding |
| ENSMUSG00000032423.12 | 3578.828 | 0.226 | 0.035 | 6.538 | 6.25E-11 | 5.54E-09 | Syncrip | protein_coding |
| ENSMUSG00000027223.15 | 16938.468 | -0.190 | 0.029 | -6.527 | 6.72E-11 | 5.92E-09 | Mapk8ip1 | protein_coding |
| ENSMUSG00000029823.16 | 9862.856 | 0.170 | 0.026 | 6.509 | 7.57E-11 | 6.63E-09 | Luc7l2 | protein_coding |
| ENSMUSG00000026384.13 | 5677.483 | 0.210 | 0.032 | 6.488 | 8.71E-11 | 7.58E-09 | Ptpn4 | protein_coding |
| ENSMUSG00000031807.10 | 741.956 | -0.372 | 0.057 | -6.484 | 8.93E-11 | 7.72E-09 | Pgls | protein_coding |
| ENSMUSG00000053395.15 | 2532.831 | -0.301 | 0.047 | -6.468 | 9.93E-11 | 8.54E-09 | Cacng8 | protein_coding |
| ENSMUSG00000018199.8 | 3434.332 | 0.210 | 0.033 | 6.440 | 1.19E-10 | 1.02E-08 | Trove2 | protein_coding |

| | | | | | | | | |
|---|---|---|---|---|---|---|---|---|
| ENSMUSG00000024925.10 | 398.444 | -0.458 | 0.071 | -6.439 | 1.20E-10 | 1.02E-08 | Rnaseh2c | protein_coding |
| ENSMUSG00000015837.15 | 19576.624 | -0.139 | 0.022 | -6.435 | 1.23E-10 | 1.04E-08 | Sqstm1 | protein_coding |
| ENSMUSG00000028557.10 | 10518.446 | 0.160 | 0.025 | 6.434 | 1.24E-10 | 1.04E-08 | Rnf11 | protein_coding |
| ENSMUSG00000020108.4 | 1777.174 | -0.341 | 0.053 | -6.429 | 1.29E-10 | 1.07E-08 | Ddit4 | protein_coding |
| ENSMUSG00000014956.15 | 12436.469 | 0.205 | 0.032 | 6.403 | 1.53E-10 | 1.27E-08 | Ppp1cb | protein_coding |
| ENSMUSG00000032997.16 | 3799.357 | -0.309 | 0.048 | -6.388 | 1.68E-10 | 1.39E-08 | Chpf | protein_coding |
| ENSMUSG00000038462.3 | 3926.862 | -0.229 | 0.036 | -6.379 | 1.78E-10 | 1.46E-08 | Uqcrfs1 | protein_coding |
| ENSMUSG00000031229.16 | 11326.121 | 0.179 | 0.028 | 6.366 | 1.94E-10 | 1.58E-08 | Atrx | protein_coding |
| ENSMUSG00000055407.14 | 8844.426 | -0.177 | 0.028 | -6.364 | 1.97E-10 | 1.60E-08 | Map6 | protein_coding |
| ENSMUSG00000021712.14 | 3636.288 | 0.216 | 0.034 | 6.362 | 1.99E-10 | 1.60E-08 | Trim23 | protein_coding |
| ENSMUSG00000034974.13 | 1424.494 | -0.281 | 0.044 | -6.347 | 2.20E-10 | 1.76E-08 | Dapk3 | protein_coding |
| ENSMUSG00000044352.6 | 4046.109 | -0.256 | 0.040 | -6.331 | 2.44E-10 | 1.94E-08 | Sowaha | protein_coding |
| ENSMUSG00000055633.6 | 374.186 | -0.443 | 0.070 | -6.320 | 2.62E-10 | 2.08E-08 | Zfp580 | protein_coding |
| ENSMUSG00000050357.9 | 2527.478 | -0.268 | 0.043 | -6.306 | 2.87E-10 | 2.26E-08 | Carmil2 | protein_coding |
| ENSMUSG00000033768.16 | 16716.471 | -0.266 | 0.042 | -6.305 | 2.89E-10 | 2.26E-08 | Nrxn2 | protein_coding |
| ENSMUSG00000041817.14 | 3530.054 | 0.189 | 0.030 | 6.301 | 2.96E-10 | 2.31E-08 | Fam169a | protein_coding |
| ENSMUSG00000042419.8 | 348.559 | -0.419 | 0.067 | -6.273 | 3.53E-10 | 2.73E-08 | Nfkbil1 | protein_coding |
| ENSMUSG00000043614.13 | 745.874 | -0.363 | 0.058 | -6.269 | 3.63E-10 | 2.80E-08 | Vps37d | protein_coding |
| ENSMUSG00000025856.15 | 1647.820 | -0.263 | 0.042 | -6.244 | 4.25E-10 | 3.26E-08 | Pdgfa | protein_coding |
| ENSMUSG00000075012.4 | 2030.838 | -0.313 | 0.050 | -6.240 | 4.36E-10 | 3.33E-08 | Fjx1 | protein_coding |
| ENSMUSG00000002409.18 | 1582.798 | -0.275 | 0.044 | -6.239 | 4.41E-10 | 3.34E-08 | Dyrk1b | protein_coding |
| ENSMUSG00000024736.14 | 16625.506 | -0.263 | 0.042 | -6.222 | 4.92E-10 | 3.71E-08 | Tmem132a | protein_coding |
| ENSMUSG00000031517.8 | 55977.624 | 0.177 | 0.029 | 6.217 | 5.07E-10 | 3.80E-08 | Gpm6a | protein_coding |
| ENSMUSG00000015599.8 | 7553.553 | -0.209 | 0.034 | -6.207 | 5.39E-10 | 4.01E-08 | Ttbk1 | protein_coding |
| ENSMUSG00000056665.2 | 1334.198 | -0.282 | 0.045 | -6.207 | 5.41E-10 | 4.01E-08 | Them6 | protein_coding |
| ENSMUSG00000042831.13 | 1036.375 | -0.295 | 0.047 | -6.206 | 5.44E-10 | 4.01E-08 | Alkbh6 | protein_coding |
| ENSMUSG00000074576.4 | 308.268 | -0.423 | 0.068 | -6.190 | 6.02E-10 | 4.42E-08 | Mocs3 | protein_coding |
| ENSMUSG00000059326.6 | 1431.179 | -0.359 | 0.058 | -6.177 | 6.53E-10 | 4.77E-08 | Csf2ra | protein_coding |
| ENSMUSG00000035236.17 | 4391.132 | 0.198 | 0.032 | 6.170 | 6.85E-10 | 4.97E-08 | Scai | protein_coding |
| ENSMUSG00000091971.3 | 486.491 | -0.441 | 0.072 | -6.165 | 7.05E-10 | 5.09E-08 | Hspa1a | protein_coding |
| ENSMUSG00000025318.13 | 10013.596 | -0.245 | 0.040 | -6.159 | 7.34E-10 | 5.28E-08 | Jph3 | protein_coding |
| ENSMUSG00000033316.14 | 7251.847 | -0.180 | 0.029 | -6.157 | 7.43E-10 | 5.31E-08 | Galnt9 | protein_coding |
| ENSMUSG00000000948.16 | 469.647 | 0.364 | 0.059 | 6.144 | 8.03E-10 | 5.72E-08 | Gm38393 | protein_coding |

| | | | | | | | | |
|---|---|---|---|---|---|---|---|---|
| ENSMUSG00000036430.8 | 527.997 | -0.398 | 0.065 | -6.132 | 8.68E-10 | 6.15E-08 | Tbcc | protein_coding |
| ENSMUSG00000005483.10 | 4357.706 | -0.223 | 0.036 | -6.125 | 9.08E-10 | 6.40E-08 | Dnajb1 | protein_coding |
| ENSMUSG00000021098.14 | 2293.258 | 0.231 | 0.038 | 6.116 | 9.57E-10 | 6.71E-08 | 4930447C04Rik | protein_coding |
| ENSMUSG00000043323.16 | 3537.032 | -0.296 | 0.048 | -6.098 | 1.07E-09 | 7.50E-08 | Fbrsl1 | protein_coding |
| ENSMUSG00000025326.12 | 6180.965 | 0.185 | 0.030 | 6.083 | 1.18E-09 | 8.18E-08 | Ube3a | protein_coding |
| ENSMUSG00000058301.8 | 4782.017 | -0.166 | 0.027 | -6.071 | 1.27E-09 | 8.81E-08 | Upf1 | protein_coding |
| ENSMUSG00000022436.15 | 8130.343 | -0.182 | 0.030 | -6.061 | 1.36E-09 | 9.32E-08 | Sh3bp1 | protein_coding |
| ENSMUSG00000056596.8 | 3908.967 | -0.192 | 0.032 | -6.060 | 1.36E-09 | 9.32E-08 | Trnp1 | protein_coding |
| ENSMUSG00000078317.6 | 658.459 | -0.361 | 0.060 | -6.027 | 1.67E-09 | 1.14E-07 | F8a | protein_coding |
| ENSMUSG00000063160.12 | 6673.029 | -0.228 | 0.038 | -6.025 | 1.69E-09 | 1.15E-07 | Numbl | protein_coding |
| ENSMUSG00000031290.14 | 1214.162 | 0.279 | 0.046 | 6.015 | 1.79E-09 | 1.20E-07 | Lrch2 | protein_coding |
| ENSMUSG00000062044.15 | 15185.482 | -0.269 | 0.045 | -6.002 | 1.95E-09 | 1.30E-07 | Lmtk3 | protein_coding |
| ENSMUSG00000020300.14 | 5960.585 | 0.171 | 0.029 | 5.992 | 2.08E-09 | 1.38E-07 | Cpeb4 | protein_coding |
| ENSMUSG00000044308.17 | 10143.177 | 0.146 | 0.024 | 5.987 | 2.13E-09 | 1.41E-07 | Ubr3 | protein_coding |
| ENSMUSG00000071337.11 | 6733.098 | 0.183 | 0.031 | 5.980 | 2.23E-09 | 1.47E-07 | Tia1 | protein_coding |
| ENSMUSG00000094724.7 | 84.731 | -0.455 | 0.076 | -5.961 | 2.51E-09 | 1.64E-07 | Rnaset2b | protein_coding |
| ENSMUSG00000051223.14 | 9029.391 | 0.155 | 0.026 | 5.952 | 2.65E-09 | 1.73E-07 | Bzw1 | protein_coding |
| ENSMUSG00000019039.13 | 3290.749 | -0.218 | 0.037 | -5.945 | 2.77E-09 | 1.80E-07 | Dalrd3 | protein_coding |
| ENSMUSG00000010803.13 | 16408.218 | 0.175 | 0.030 | 5.939 | 2.86E-09 | 1.85E-07 | Gabra1 | protein_coding |
| ENSMUSG00000040097.15 | 13012.460 | -0.184 | 0.031 | -5.930 | 3.02E-09 | 1.95E-07 | Flywch1 | protein_coding |
| ENSMUSG00000025034.8 | 2860.118 | -0.266 | 0.045 | -5.923 | 3.17E-09 | 2.03E-07 | Trim8 | protein_coding |
| ENSMUSG00000090125.3 | 821.213 | -0.352 | 0.059 | -5.922 | 3.18E-09 | 2.03E-07 | Pou3f1 | protein_coding |
| ENSMUSG00000038880.13 | 994.468 | -0.328 | 0.055 | -5.921 | 3.21E-09 | 2.04E-07 | Mrps34 | protein_coding |
| ENSMUSG00000035027.18 | 3835.513 | -0.199 | 0.034 | -5.912 | 3.37E-09 | 2.13E-07 | Map2k2 | protein_coding |
| ENSMUSG00000002107.18 | 24736.775 | 0.148 | 0.025 | 5.910 | 3.42E-09 | 2.15E-07 | Celf2 | protein_coding |
| ENSMUSG00000069045.11 | 2655.272 | 0.216 | 0.037 | 5.869 | 4.39E-09 | 2.75E-07 | Ddx3y | protein_coding |
| ENSMUSG00000029267.17 | 2888.283 | 0.226 | 0.039 | 5.867 | 4.44E-09 | 2.77E-07 | Mtf2 | protein_coding |
| ENSMUSG00000051067.8 | 1775.494 | -0.320 | 0.055 | -5.865 | 4.50E-09 | 2.79E-07 | Lingo3 | protein_coding |
| ENSMUSG00000038453.16 | 22631.224 | -0.187 | 0.032 | -5.859 | 4.65E-09 | 2.86E-07 | Srcin1 | protein_coding |
| ENSMUSG00000028134.11 | 4494.322 | 0.174 | 0.030 | 5.859 | 4.65E-09 | 2.86E-07 | Ptbp2 | protein_coding |
| ENSMUSG00000041268.17 | 18258.846 | 0.122 | 0.021 | 5.851 | 4.90E-09 | 3.00E-07 | Dmxl2 | protein_coding |
| ENSMUSG00000003814.8 | 18503.463 | -0.164 | 0.028 | -5.841 | 5.18E-09 | 3.16E-07 | Calr | protein_coding |
| ENSMUSG00000028782.14 | 27737.876 | -0.206 | 0.035 | -5.839 | 5.26E-09 | 3.20E-07 | Adgrb2 | protein_coding |

| | | | | | | | | |
|---|---|---|---|---|---|---|---|---|
| ENSMUSG00000045763.7 | 14642.200 | -0.293 | 0.050 | -5.837 | 5.32E-09 | 3.21E-07 | Basp1 | protein_coding |
| ENSMUSG00000052372.10 | 732.179 | 0.336 | 0.058 | 5.836 | 5.33E-09 | 3.21E-07 | Il1rapl1 | protein_coding |
| ENSMUSG00000095687.1 | 105.105 | -0.451 | 0.077 | -5.831 | 5.52E-09 | 3.31E-07 | Rnaset2a | protein_coding |
| ENSMUSG00000003452.15 | 2465.031 | 0.213 | 0.037 | 5.800 | 6.63E-09 | 3.96E-07 | Bicd1 | protein_coding |
| ENSMUSG00000020441.6 | 598.194 | -0.335 | 0.058 | -5.799 | 6.66E-09 | 3.96E-07 | 2310033P09Rik | protein_coding |
| ENSMUSG00000074405.7 | 1720.240 | -0.275 | 0.047 | -5.798 | 6.71E-09 | 3.97E-07 | Zfp865 | protein_coding |
| ENSMUSG00000020572.8 | 2773.572 | 0.203 | 0.035 | 5.794 | 6.86E-09 | 4.04E-07 | Nampt | protein_coding |
| ENSMUSG00000031198.4 | 1380.431 | 0.254 | 0.044 | 5.782 | 7.38E-09 | 4.31E-07 | Fundc2 | protein_coding |
| ENSMUSG00000008036.11 | 4712.806 | -0.192 | 0.033 | -5.782 | 7.40E-09 | 4.31E-07 | Ap2s1 | protein_coding |
| ENSMUSG00000049295.16 | 1721.449 | -0.263 | 0.045 | -5.775 | 7.70E-09 | 4.45E-07 | Zfp219 | protein_coding |
| ENSMUSG00000034755.18 | 794.634 | 0.320 | 0.055 | 5.775 | 7.71E-09 | 4.45E-07 | Pcdh11x | protein_coding |
| ENSMUSG00000040118.15 | 10049.579 | 0.213 | 0.037 | 5.768 | 8.01E-09 | 4.61E-07 | Cacna2d1 | protein_coding |
| ENSMUSG00000038255.6 | 3058.671 | -0.299 | 0.052 | -5.766 | 8.10E-09 | 4.64E-07 | Neurod2 | protein_coding |
| ENSMUSG00000018846.8 | 3644.024 | 0.198 | 0.034 | 5.764 | 8.23E-09 | 4.70E-07 | Pank3 | protein_coding |
| ENSMUSG00000034949.18 | 4768.263 | -0.230 | 0.040 | -5.739 | 9.53E-09 | 5.40E-07 | Zfr2 | protein_coding |
| ENSMUSG00000026034.17 | 7364.418 | 0.166 | 0.029 | 5.739 | 9.54E-09 | 5.40E-07 | Clk1 | protein_coding |
| ENSMUSG00000035890.9 | 2361.305 | -0.244 | 0.043 | -5.737 | 9.63E-09 | 5.43E-07 | Rnf126 | protein_coding |
| ENSMUSG00000053080.10 | 2091.640 | -0.242 | 0.042 | -5.726 | 1.03E-08 | 5.78E-07 | 2700081O15Rik | protein_coding |
| ENSMUSG00000068267.5 | 6104.550 | -0.187 | 0.033 | -5.725 | 1.03E-08 | 5.78E-07 | Cenpb | protein_coding |
| ENSMUSG00000079658.9 | 4230.038 | 0.196 | 0.034 | 5.722 | 1.05E-08 | 5.86E-07 | Eloc | protein_coding |
| ENSMUSG00000039556.16 | 219.852 | -0.434 | 0.076 | -5.718 | 1.08E-08 | 5.95E-07 | Ppp1r3f | protein_coding |
| ENSMUSG00000048385.8 | 4629.500 | -0.289 | 0.050 | -5.718 | 1.08E-08 | 5.95E-07 | Scrt1 | protein_coding |
| ENSMUSG00000067786.16 | 5994.250 | -0.314 | 0.055 | -5.714 | 1.10E-08 | 6.07E-07 | Nnat | protein_coding |
| ENSMUSG00000038406.17 | 9771.705 | -0.185 | 0.032 | -5.707 | 1.15E-08 | 6.31E-07 | Scaf1 | protein_coding |
| ENSMUSG00000053166.14 | 1615.999 | -0.317 | 0.056 | -5.706 | 1.16E-08 | 6.33E-07 | Cdh22 | protein_coding |
| ENSMUSG00000087403.9 | 1788.113 | 0.212 | 0.037 | 5.700 | 1.20E-08 | 6.51E-07 | Kantr | protein_coding |
| ENSMUSG00000026223.15 | 32457.444 | -0.155 | 0.027 | -5.700 | 1.20E-08 | 6.51E-07 | Itm2c | protein_coding |
| ENSMUSG00000040659.3 | 14846.723 | -0.157 | 0.028 | -5.691 | 1.27E-08 | 6.84E-07 | Efhd2 | protein_coding |
| ENSMUSG00000021288.18 | 31753.493 | -0.161 | 0.028 | -5.687 | 1.30E-08 | 6.97E-07 | Klc1 | protein_coding |
| ENSMUSG00000021687.14 | 10555.336 | 0.133 | 0.023 | 5.683 | 1.33E-08 | 7.11E-07 | Scamp1 | protein_coding |
| ENSMUSG00000021508.10 | 2905.289 | -0.278 | 0.049 | -5.652 | 1.59E-08 | 8.48E-07 | Cxcl14 | protein_coding |
| ENSMUSG00000022208.11 | 20087.777 | -0.195 | 0.035 | -5.651 | 1.59E-08 | 8.48E-07 | Jph4 | protein_coding |
| ENSMUSG00000071637.5 | 99.518 | -0.412 | 0.073 | -5.644 | 1.66E-08 | 8.78E-07 | Cebpd | protein_coding |

| | | | | | | | | |
|---|---|---|---|---|---|---|---|---|
| ENSMUSG00000049807.16 | 7991.453 | -0.216 | 0.038 | -5.641 | 1.69E-08 | 8.92E-07 | Arhgap23 | protein_coding |
| ENSMUSG00000004931.11 | 1109.715 | -0.310 | 0.055 | -5.641 | 1.70E-08 | 8.92E-07 | Apba3 | protein_coding |
| ENSMUSG00000046321.8 | 3321.863 | -0.266 | 0.047 | -5.632 | 1.78E-08 | 9.33E-07 | Hs3st2 | protein_coding |
| ENSMUSG00000037458.14 | 8412.219 | 0.147 | 0.026 | 5.628 | 1.82E-08 | 9.49E-07 | Azin1 | protein_coding |
| ENSMUSG00000034336.3 | 10806.241 | -0.172 | 0.031 | -5.627 | 1.84E-08 | 9.55E-07 | Ina | protein_coding |
| ENSMUSG00000037771.11 | 4816.919 | -0.326 | 0.058 | -5.624 | 1.86E-08 | 9.66E-07 | Slc32a1 | protein_coding |
| ENSMUSG00000045176.3 | 920.352 | -0.325 | 0.058 | -5.622 | 1.89E-08 | 9.71E-07 | Borcs6 | protein_coding |
| ENSMUSG00000024772.9 | 1644.249 | -0.236 | 0.042 | -5.622 | 1.89E-08 | 9.71E-07 | Ehd1 | protein_coding |
| ENSMUSG00000060152.14 | 1332.594 | -0.261 | 0.047 | -5.604 | 2.09E-08 | 1.07E-06 | Pop5 | protein_coding |
| ENSMUSG00000031246.14 | 3153.318 | 0.203 | 0.036 | 5.603 | 2.11E-08 | 1.08E-06 | Sh3bgrl | protein_coding |
| ENSMUSG00000037904.14 | 399.479 | -0.382 | 0.068 | -5.583 | 2.37E-08 | 1.20E-06 | Ankrd9 | protein_coding |
| ENSMUSG00000048920.8 | 1834.603 | -0.229 | 0.041 | -5.568 | 2.58E-08 | 1.31E-06 | Fkrp | protein_coding |
| ENSMUSG00000025531.14 | 2657.289 | 0.199 | 0.036 | 5.564 | 2.64E-08 | 1.33E-06 | Chm | protein_coding |
| ENSMUSG00000046658.16 | 1330.931 | -0.243 | 0.044 | -5.560 | 2.70E-08 | 1.36E-06 | Zfp316 | protein_coding |
| ENSMUSG00000035285.6 | 3131.244 | -0.234 | 0.042 | -5.553 | 2.81E-08 | 1.41E-06 | Nat14 | protein_coding |
| ENSMUSG00000020483.14 | 23272.940 | -0.119 | 0.021 | -5.547 | 2.90E-08 | 1.45E-06 | Dynll2 | protein_coding |
| ENSMUSG00000045348.16 | 5271.072 | -0.174 | 0.031 | -5.546 | 2.92E-08 | 1.46E-06 | Nyap1 | protein_coding |
| ENSMUSG00000037217.15 | 31058.872 | -0.196 | 0.035 | -5.537 | 3.07E-08 | 1.52E-06 | Syn1 | protein_coding |
| ENSMUSG00000033953.10 | 26464.608 | 0.163 | 0.029 | 5.528 | 3.24E-08 | 1.60E-06 | Ppp3r1 | protein_coding |
| ENSMUSG00000030397.10 | 4389.887 | -0.200 | 0.036 | -5.523 | 3.32E-08 | 1.64E-06 | Mark4 | protein_coding |
| ENSMUSG00000025903.14 | 723.674 | 0.313 | 0.057 | 5.519 | 3.41E-08 | 1.68E-06 | Lypla1 | protein_coding |
| ENSMUSG00000002504.14 | 2214.771 | -0.243 | 0.044 | -5.518 | 3.43E-08 | 1.68E-06 | Slc9a3r2 | protein_coding |
| ENSMUSG00000015337.5 | 358.345 | -0.382 | 0.069 | -5.513 | 3.52E-08 | 1.72E-06 | Endog | protein_coding |
| ENSMUSG00000026761.12 | 2719.665 | 0.201 | 0.036 | 5.512 | 3.55E-08 | 1.73E-06 | Orc4 | protein_coding |
| ENSMUSG00000054708.17 | 2697.355 | -0.223 | 0.040 | -5.502 | 3.75E-08 | 1.82E-06 | Ankrd24 | protein_coding |
| ENSMUSG00000060708.4 | 318.361 | -0.382 | 0.069 | -5.500 | 3.80E-08 | 1.83E-06 | Bloc1s4 | protein_coding |
| ENSMUSG00000075467.4 | 1622.585 | -0.244 | 0.044 | -5.498 | 3.83E-08 | 1.84E-06 | Dnlz | protein_coding |
| ENSMUSG00000058709.11 | 3548.943 | -0.197 | 0.036 | -5.495 | 3.91E-08 | 1.88E-06 | Egln2 | protein_coding |
| ENSMUSG00000070705.3 | 990.083 | -0.275 | 0.050 | -5.484 | 4.15E-08 | 1.98E-06 | Eid2b | protein_coding |
| ENSMUSG00000022338.6 | 2616.035 | 0.185 | 0.034 | 5.481 | 4.24E-08 | 2.02E-06 | Eny2 | protein_coding |
| ENSMUSG00000068457.14 | 856.903 | 0.278 | 0.051 | 5.480 | 4.26E-08 | 2.02E-06 | Uty | protein_coding |
| ENSMUSG00000035992.15 | 3098.212 | 0.184 | 0.034 | 5.475 | 4.36E-08 | 2.06E-06 | Fnip1 | protein_coding |
| ENSMUSG00000023272.3 | 1147.467 | -0.253 | 0.046 | -5.468 | 4.55E-08 | 2.13E-06 | Creld2 | protein_coding |

| | | | | | | | | |
|---|---|---|---|---|---|---|---|---|
| ENSMUSG00000022623.15 | 12024.963 | -0.231 | 0.042 | -5.468 | 4.56E-08 | 2.13E-06 | Shank3 | protein_coding |
| ENSMUSG00000024121.13 | 2237.865 | -0.224 | 0.041 | -5.468 | 4.56E-08 | 2.13E-06 | Atp6v0c | protein_coding |
| ENSMUSG00000085795.8 | 1446.007 | -0.297 | 0.054 | -5.466 | 4.61E-08 | 2.15E-06 | Zfp703 | protein_coding |
| ENSMUSG00000041556.8 | 1977.040 | -0.255 | 0.047 | -5.455 | 4.90E-08 | 2.28E-06 | Fbxo2 | protein_coding |
| ENSMUSG00000033701.13 | 1654.348 | -0.210 | 0.038 | -5.454 | 4.93E-08 | 2.28E-06 | Acbd6 | protein_coding |
| ENSMUSG00000042208.15 | 2290.630 | 0.193 | 0.036 | 5.448 | 5.08E-08 | 2.35E-06 | 0610010F05Rik | protein_coding |
| ENSMUSG00000032575.16 | 2227.003 | -0.266 | 0.049 | -5.448 | 5.09E-08 | 2.35E-06 | Manf | protein_coding |
| ENSMUSG00000026977.17 | 3132.445 | 0.178 | 0.033 | 5.439 | 5.35E-08 | 2.46E-06 | March7 | protein_coding |
| ENSMUSG00000094483.2 | 22417.478 | 0.165 | 0.030 | 5.428 | 5.69E-08 | 2.60E-06 | Purb | protein_coding |
| ENSMUSG00000027602.9 | 4853.485 | -0.180 | 0.033 | -5.425 | 5.80E-08 | 2.65E-06 | Map1lc3a | protein_coding |
| ENSMUSG00000036745.15 | 12721.836 | 0.138 | 0.025 | 5.409 | 6.34E-08 | 2.88E-06 | Ttll7 | protein_coding |
| ENSMUSG00000046449.15 | 2597.548 | 0.190 | 0.035 | 5.400 | 6.66E-08 | 3.02E-06 | C77370 | protein_coding |
| ENSMUSG00000040459.11 | 7215.855 | 0.188 | 0.035 | 5.395 | 6.85E-08 | 3.10E-06 | Arglu1 | protein_coding |
| ENSMUSG00000004929.12 | 2702.844 | -0.232 | 0.043 | -5.392 | 6.99E-08 | 3.15E-06 | Thop1 | protein_coding |
| ENSMUSG00000027589.14 | 4786.133 | 0.149 | 0.028 | 5.386 | 7.21E-08 | 3.22E-06 | Pcmtd2 | protein_coding |
| ENSMUSG00000051695.6 | 5522.616 | -0.226 | 0.042 | -5.383 | 7.31E-08 | 3.26E-06 | Pcbp1 | protein_coding |
| ENSMUSG00000036622.15 | 8902.447 | -0.245 | 0.045 | -5.381 | 7.40E-08 | 3.29E-06 | Atp13a2 | protein_coding |
| ENSMUSG00000041309.17 | 320.631 | -0.406 | 0.076 | -5.368 | 7.94E-08 | 3.52E-06 | Nkx6-2 | protein_coding |
| ENSMUSG00000000787.12 | 10537.439 | 0.143 | 0.027 | 5.364 | 8.12E-08 | 3.59E-06 | Ddx3x | protein_coding |
| ENSMUSG00000038916.7 | 2861.898 | -0.210 | 0.039 | -5.363 | 8.18E-08 | 3.60E-06 | Soga3 | protein_coding |
| ENSMUSG00000045318.6 | 685.804 | -0.338 | 0.063 | -5.359 | 8.36E-08 | 3.67E-06 | Adra2c | protein_coding |
| ENSMUSG00000051703.14 | 2997.577 | -0.244 | 0.045 | -5.358 | 8.43E-08 | 3.69E-06 | Tmem198 | protein_coding |
| ENSMUSG00000024962.13 | 1859.051 | -0.253 | 0.047 | -5.347 | 8.95E-08 | 3.90E-06 | Vegfb | protein_coding |
| ENSMUSG00000038174.14 | 6834.272 | 0.178 | 0.033 | 5.342 | 9.18E-08 | 3.99E-06 | Fam126b | protein_coding |
| ENSMUSG00000033186.8 | 3199.245 | 0.179 | 0.034 | 5.341 | 9.22E-08 | 3.99E-06 | Mzt1 | protein_coding |
| ENSMUSG00000052516.17 | 4902.321 | 0.161 | 0.030 | 5.341 | 9.23E-08 | 3.99E-06 | Robo2 | protein_coding |
| ENSMUSG00000021087.18 | 48418.926 | -0.122 | 0.023 | -5.330 | 9.81E-08 | 4.23E-06 | Rtn1 | protein_coding |
| ENSMUSG00000015092.9 | 3057.939 | -0.203 | 0.038 | -5.320 | 1.04E-07 | 4.46E-06 | Edf1 | protein_coding |
| ENSMUSG00000056708.5 | 3385.904 | -0.301 | 0.057 | -5.319 | 1.04E-07 | 4.46E-06 | Ier5 | protein_coding |
| ENSMUSG00000046470.5 | 210.046 | -0.413 | 0.078 | -5.314 | 1.07E-07 | 4.57E-06 | Sox18 | protein_coding |
| ENSMUSG00000002763.16 | 2890.377 | -0.212 | 0.040 | -5.314 | 1.07E-07 | 4.57E-06 | Pex6 | protein_coding |
| ENSMUSG00000039195.4 | 3311.054 | -0.215 | 0.040 | -5.311 | 1.09E-07 | 4.63E-06 | 1110008P14Rik | protein_coding |
| ENSMUSG00000066721.3 | 414.366 | -0.331 | 0.062 | -5.308 | 1.11E-07 | 4.69E-06 | Zfp575 | protein_coding |

| | | | | | | | | |
|---|---|---|---|---|---|---|---|---|
| ENSMUSG00000009207.15 | 2318.693 | 0.194 | 0.037 | 5.304 | 1.13E-07 | 4.77E-06 | Lnpk | protein_coding |
| ENSMUSG00000029050.15 | 8189.933 | -0.147 | 0.028 | -5.304 | 1.13E-07 | 4.77E-06 | Ski | protein_coding |
| ENSMUSG00000026775.9 | 4447.565 | 0.153 | 0.029 | 5.294 | 1.20E-07 | 5.03E-06 | Yme1l1 | protein_coding |
| ENSMUSG00000036353.13 | 1322.658 | 0.254 | 0.048 | 5.293 | 1.21E-07 | 5.04E-06 | P2ry12 | protein_coding |
| ENSMUSG00000037740.8 | 1717.082 | -0.227 | 0.043 | -5.287 | 1.24E-07 | 5.19E-06 | Mrps26 | protein_coding |
| ENSMUSG00000078566.8 | 4013.503 | 0.168 | 0.032 | 5.275 | 1.33E-07 | 5.51E-06 | Bnip3 | protein_coding |
| ENSMUSG00000020167.14 | 1439.219 | -0.229 | 0.044 | -5.263 | 1.42E-07 | 5.88E-06 | Tcf3 | protein_coding |
| ENSMUSG00000022463.7 | 13537.677 | -0.137 | 0.026 | -5.262 | 1.43E-07 | 5.89E-06 | Srebf2 | protein_coding |
| ENSMUSG00000055531.12 | 6485.973 | 0.143 | 0.027 | 5.258 | 1.46E-07 | 6.00E-06 | Cpsf6 | protein_coding |
| ENSMUSG00000034675.17 | 15930.776 | -0.167 | 0.032 | -5.256 | 1.47E-07 | 6.05E-06 | Dbn1 | protein_coding |
| ENSMUSG00000019139.10 | 1684.677 | -0.283 | 0.054 | -5.255 | 1.48E-07 | 6.06E-06 | Isyna1 | protein_coding |
| ENSMUSG00000037126.15 | 20850.649 | -0.206 | 0.039 | -5.251 | 1.51E-07 | 6.17E-06 | Psd | protein_coding |
| ENSMUSG00000030847.8 | 358.786 | -0.339 | 0.065 | -5.248 | 1.54E-07 | 6.26E-06 | Bag3 | protein_coding |
| ENSMUSG00000030846.15 | 5172.613 | 0.147 | 0.028 | 5.245 | 1.56E-07 | 6.35E-06 | Tial1 | protein_coding |
| ENSMUSG00000035640.18 | 14251.594 | -0.216 | 0.041 | -5.240 | 1.61E-07 | 6.51E-06 | Cbarp | protein_coding |
| ENSMUSG00000030759.16 | 6813.646 | 0.150 | 0.029 | 5.230 | 1.70E-07 | 6.84E-06 | Far1 | protein_coding |
| ENSMUSG00000050761.3 | 275.782 | -0.392 | 0.075 | -5.226 | 1.73E-07 | 6.95E-06 | Gp1bb | protein_coding |
| ENSMUSG00000056724.14 | 2154.307 | -0.204 | 0.039 | -5.221 | 1.78E-07 | 7.13E-06 | Nbeal2 | protein_coding |
| ENSMUSG00000056917.12 | 795.472 | -0.309 | 0.059 | -5.218 | 1.80E-07 | 7.20E-06 | Sipa1 | protein_coding |
| ENSMUSG00000067847.13 | 632.458 | -0.326 | 0.063 | -5.212 | 1.87E-07 | 7.44E-06 | Romo1 | protein_coding |
| ENSMUSG00000018293.4 | 3211.829 | -0.247 | 0.047 | -5.210 | 1.89E-07 | 7.51E-06 | Pfn1 | protein_coding |
| ENSMUSG00000025876.15 | 7317.289 | -0.199 | 0.038 | -5.209 | 1.90E-07 | 7.51E-06 | Unc5a | protein_coding |
| ENSMUSG00000066324.2 | 5860.911 | 0.159 | 0.031 | 5.206 | 1.93E-07 | 7.63E-06 | Impad1 | protein_coding |
| ENSMUSG00000020734.13 | 2767.447 | -0.212 | 0.041 | -5.204 | 1.95E-07 | 7.66E-06 | Grin2c | protein_coding |
| ENSMUSG00000039477.16 | 7457.360 | -0.220 | 0.042 | -5.202 | 1.97E-07 | 7.72E-06 | Tnrc18 | protein_coding |
| ENSMUSG00000035967.15 | 5004.849 | 0.156 | 0.030 | 5.202 | 1.97E-07 | 7.72E-06 | Ints6l | protein_coding |
| ENSMUSG00000028851.6 | 3810.918 | -0.168 | 0.032 | -5.200 | 1.99E-07 | 7.79E-06 | Nudc | protein_coding |
| ENSMUSG00000051515.9 | 535.314 | -0.323 | 0.062 | -5.197 | 2.02E-07 | 7.88E-06 | Fam181b | protein_coding |
| ENSMUSG00000056501.3 | 54.047 | -0.389 | 0.075 | -5.193 | 2.07E-07 | NA | Cebpb | protein_coding |
| ENSMUSG00000022378.13 | 6416.637 | 0.159 | 0.031 | 5.193 | 2.07E-07 | 8.04E-06 | Fam49b | protein_coding |
| ENSMUSG00000032115.14 | 8110.712 | -0.147 | 0.028 | -5.190 | 2.11E-07 | 8.15E-06 | Hyou1 | protein_coding |
| ENSMUSG00000002274.12 | 1329.323 | -0.315 | 0.061 | -5.185 | 2.16E-07 | 8.33E-06 | Metrn | protein_coding |
| ENSMUSG00000056342.16 | 11761.357 | 0.126 | 0.024 | 5.181 | 2.21E-07 | 8.51E-06 | Usp34 | protein_coding |

| | | | | | | | | |
|---|---|---|---|---|---|---|---|---|
| ENSMUSG00000001376.17 | 4005.142 | 0.163 | 0.032 | 5.148 | 2.64E-07 | 1.01E-05 | Vps50 | protein_coding |
| ENSMUSG00000027692.16 | 12325.366 | 0.120 | 0.023 | 5.147 | 2.65E-07 | 1.01E-05 | Tnik | protein_coding |
| ENSMUSG00000020232.17 | 579.014 | -0.308 | 0.060 | -5.147 | 2.65E-07 | 1.01E-05 | Hmg20b | protein_coding |
| ENSMUSG00000025375.15 | 20218.030 | -0.175 | 0.034 | -5.144 | 2.68E-07 | 1.02E-05 | Aatk | protein_coding |
| ENSMUSG00000037234.17 | 7525.498 | 0.127 | 0.025 | 5.140 | 2.75E-07 | 1.04E-05 | Hook3 | protein_coding |
| ENSMUSG00000036438.12 | 56102.042 | 0.127 | 0.025 | 5.133 | 2.86E-07 | 1.08E-05 | Calm2 | protein_coding |
| ENSMUSG00000032407.14 | 3961.227 | 0.160 | 0.031 | 5.131 | 2.88E-07 | 1.09E-05 | U2surp | protein_coding |
| ENSMUSG00000055148.7 | 294.128 | -0.388 | 0.076 | -5.124 | 3.00E-07 | 1.13E-05 | Klf2 | protein_coding |
| ENSMUSG00000049556.5 | 14621.710 | -0.207 | 0.040 | -5.123 | 3.01E-07 | 1.13E-05 | Lingo1 | protein_coding |
| ENSMUSG00000006435.15 | 10351.969 | -0.162 | 0.032 | -5.121 | 3.04E-07 | 1.14E-05 | Neurl1a | protein_coding |
| ENSMUSG00000030034.11 | 470.611 | -0.339 | 0.066 | -5.118 | 3.09E-07 | 1.15E-05 | Ino80b | protein_coding |
| ENSMUSG00000037060.3 | 536.011 | -0.327 | 0.064 | -5.113 | 3.17E-07 | 1.18E-05 | Prkcdbp | protein_coding |
| ENSMUSG00000031292.14 | 10932.327 | 0.128 | 0.025 | 5.096 | 3.46E-07 | 1.28E-05 | Cdkl5 | protein_coding |
| ENSMUSG00000042116.3 | 1488.888 | -0.245 | 0.048 | -5.096 | 3.47E-07 | 1.28E-05 | Vwa1 | protein_coding |
| ENSMUSG00000031834.15 | 7480.213 | -0.167 | 0.033 | -5.096 | 3.48E-07 | 1.28E-05 | Pik3r2 | protein_coding |
| ENSMUSG00000030376.8 | 19848.950 | -0.153 | 0.030 | -5.096 | 3.48E-07 | 1.28E-05 | Slc8a2 | protein_coding |
| ENSMUSG00000046822.12 | 4720.074 | -0.163 | 0.032 | -5.095 | 3.48E-07 | 1.28E-05 | Slc39a3 | protein_coding |
| ENSMUSG00000033510.14 | 2291.394 | -0.187 | 0.037 | -5.092 | 3.54E-07 | 1.30E-05 | Otud7a | protein_coding |
| ENSMUSG00000025825.12 | 2008.505 | -0.187 | 0.037 | -5.088 | 3.62E-07 | 1.32E-05 | Iscu | protein_coding |
| ENSMUSG00000041560.12 | 2512.904 | -0.209 | 0.041 | -5.085 | 3.67E-07 | 1.34E-05 | Nop53 | protein_coding |
| ENSMUSG00000046574.8 | 4281.415 | -0.157 | 0.031 | -5.078 | 3.81E-07 | 1.39E-05 | Prr12 | protein_coding |
| ENSMUSG00000063765.11 | 516.745 | -0.335 | 0.066 | -5.076 | 3.86E-07 | 1.40E-05 | Chadl | protein_coding |
| ENSMUSG00000021693.19 | 4302.149 | 0.154 | 0.030 | 5.066 | 4.07E-07 | 1.47E-05 | Kif2a | protein_coding |
| ENSMUSG00000029463.5 | 1956.332 | -0.204 | 0.040 | -5.064 | 4.11E-07 | 1.49E-05 | Fam216a | protein_coding |
| ENSMUSG00000046699.14 | 2150.233 | 0.227 | 0.045 | 5.057 | 4.25E-07 | 1.53E-05 | Slitrk4 | protein_coding |
| ENSMUSG00000032485.14 | 5744.882 | -0.138 | 0.027 | -5.052 | 4.37E-07 | 1.57E-05 | Scap | protein_coding |
| ENSMUSG00000023845.6 | 2653.618 | 0.185 | 0.037 | 5.051 | 4.40E-07 | 1.58E-05 | Lnpep | protein_coding |
| ENSMUSG00000026739.13 | 2076.753 | 0.185 | 0.037 | 5.033 | 4.82E-07 | 1.72E-05 | Bmi1 | protein_coding |
| ENSMUSG00000023353.14 | 12847.615 | -0.169 | 0.034 | -5.032 | 4.85E-07 | 1.73E-05 | Agap3 | protein_coding |
| ENSMUSG00000038550.10 | 753.517 | 0.264 | 0.052 | 5.031 | 4.87E-07 | 1.73E-05 | Ciart | protein_coding |
| ENSMUSG00000020152.7 | 13507.899 | 0.134 | 0.027 | 5.031 | 4.88E-07 | 1.73E-05 | Actr2 | protein_coding |
| ENSMUSG00000022175.8 | 1373.659 | -0.223 | 0.044 | -5.028 | 4.95E-07 | 1.75E-05 | Lrp10 | protein_coding |
| ENSMUSG00000087006.3 | 626.778 | -0.335 | 0.067 | -5.028 | 4.95E-07 | 1.75E-05 | Gm13889 | protein_coding |

| | | | | | | | | |
|---|---|---|---|---|---|---|---|---|
| ENSMUSG00000000202.9 | 458.018 | -0.328 | 0.065 | -5.020 | 5.17E-07 | 1.82E-05 | Btbd17 | protein_coding |
| ENSMUSG00000020362.13 | 3115.284 | 0.171 | 0.034 | 5.016 | 5.27E-07 | 1.85E-05 | Cnot6 | protein_coding |
| ENSMUSG00000096768.7 | 261.480 | -0.321 | 0.064 | -5.014 | 5.32E-07 | 1.86E-05 | Erdr1 | protein_coding |
| ENSMUSG00000029478.16 | 13430.336 | -0.221 | 0.044 | -5.011 | 5.41E-07 | 1.89E-05 | Ncor2 | protein_coding |
| ENSMUSG00000021134.17 | 14604.316 | 0.139 | 0.028 | 5.011 | 5.43E-07 | 1.89E-05 | Srsf5 | protein_coding |
| ENSMUSG00000037608.16 | 7420.576 | 0.163 | 0.033 | 5.010 | 5.43E-07 | 1.89E-05 | Bclaf1 | protein_coding |
| ENSMUSG00000091811.2 | 933.535 | -0.243 | 0.048 | -5.010 | 5.46E-07 | 1.89E-05 | Inafm1 | protein_coding |
| ENSMUSG00000031878.18 | 2484.679 | 0.184 | 0.037 | 5.008 | 5.51E-07 | 1.90E-05 | Nae1 | protein_coding |
| ENSMUSG00000022565.15 | 16094.195 | -0.198 | 0.039 | -5.007 | 5.53E-07 | 1.91E-05 | Plec | protein_coding |
| ENSMUSG00000048616.4 | 242.835 | -0.370 | 0.074 | -5.003 | 5.64E-07 | 1.94E-05 | Nog | protein_coding |
| ENSMUSG00000020687.13 | 3395.348 | 0.157 | 0.031 | 5.001 | 5.71E-07 | 1.96E-05 | Cdc27 | protein_coding |
| ENSMUSG00000038034.15 | 10518.716 | -0.185 | 0.037 | -4.995 | 5.88E-07 | 2.01E-05 | Igsf8 | protein_coding |
| ENSMUSG00000071172.12 | 6515.638 | 0.145 | 0.029 | 4.987 | 6.13E-07 | 2.09E-05 | Srsf3 | protein_coding |
| ENSMUSG00000042570.14 | 2205.728 | -0.228 | 0.046 | -4.985 | 6.21E-07 | 2.11E-05 | Mier2 | protein_coding |
| ENSMUSG00000040855.15 | 13418.012 | 0.150 | 0.030 | 4.982 | 6.31E-07 | 2.14E-05 | Reps2 | protein_coding |
| ENSMUSG00000031565.18 | 4317.262 | -0.197 | 0.040 | -4.980 | 6.37E-07 | 2.16E-05 | Fgfr1 | protein_coding |
| ENSMUSG00000027087.11 | 3732.754 | 0.154 | 0.031 | 4.977 | 6.47E-07 | 2.19E-05 | Itgav | protein_coding |
| ENSMUSG00000058441.7 | 4556.246 | -0.194 | 0.039 | -4.972 | 6.62E-07 | 2.23E-05 | Panx2 | protein_coding |
| ENSMUSG00000025384.15 | 1196.626 | -0.252 | 0.051 | -4.971 | 6.66E-07 | 2.24E-05 | Faap100 | protein_coding |
| ENSMUSG00000042745.9 | 339.512 | -0.359 | 0.072 | -4.970 | 6.69E-07 | 2.24E-05 | Id1 | protein_coding |
| ENSMUSG00000044024.15 | 5485.464 | -0.217 | 0.044 | -4.963 | 6.92E-07 | 2.32E-05 | Rell2 | protein_coding |
| ENSMUSG00000019194.15 | 9276.650 | -0.162 | 0.033 | -4.961 | 7.02E-07 | 2.34E-05 | Scn1b | protein_coding |
| ENSMUSG00000059213.6 | 50720.774 | -0.191 | 0.038 | -4.959 | 7.07E-07 | 2.36E-05 | Ddn | protein_coding |
| ENSMUSG00000025135.12 | 1345.399 | -0.209 | 0.042 | -4.954 | 7.29E-07 | 2.42E-05 | Anapc11 | protein_coding |
| ENSMUSG00000029655.17 | 2534.672 | 0.171 | 0.035 | 4.952 | 7.34E-07 | 2.43E-05 | N4bp2l2 | protein_coding |
| ENSMUSG00000043535.13 | 5199.795 | 0.138 | 0.028 | 4.952 | 7.36E-07 | 2.44E-05 | Setx | protein_coding |
| ENSMUSG00000047067.7 | 734.036 | -0.287 | 0.058 | -4.950 | 7.42E-07 | 2.45E-05 | Dusp28 | protein_coding |
| ENSMUSG00000004151.17 | 3418.371 | 0.187 | 0.038 | 4.945 | 7.61E-07 | 2.51E-05 | Etv1 | protein_coding |
| ENSMUSG00000047428.14 | 2016.425 | -0.227 | 0.046 | -4.943 | 7.71E-07 | 2.53E-05 | Dlk2 | protein_coding |
| ENSMUSG00000021775.10 | 6286.107 | 0.173 | 0.035 | 4.940 | 7.80E-07 | 2.56E-05 | Nr1d2 | protein_coding |
| ENSMUSG00000035133.9 | 6349.622 | 0.142 | 0.029 | 4.939 | 7.84E-07 | 2.56E-05 | Arhgap5 | protein_coding |
| ENSMUSG00000031833.10 | 17478.809 | -0.162 | 0.033 | -4.935 | 8.00E-07 | 2.61E-05 | Mast3 | protein_coding |
| ENSMUSG00000029030.14 | 6057.226 | -0.144 | 0.029 | -4.931 | 8.18E-07 | 2.66E-05 | Tprgl | protein_coding |

| | | | | | | | | |
|---|---|---|---|---|---|---|---|---|
| ENSMUSG00000038777.19 | 1051.875 | -0.252 | 0.051 | -4.930 | 8.21E-07 | 2.66E-05 | Sema6c | protein_coding |
| ENSMUSG00000093803.3 | 830.699 | -0.278 | 0.056 | -4.930 | 8.22E-07 | 2.66E-05 | Ppp2r3d | protein_coding |
| ENSMUSG00000036990.13 | 4374.919 | 0.148 | 0.030 | 4.925 | 8.44E-07 | 2.72E-05 | Otud4 | protein_coding |
| ENSMUSG00000038884.14 | 4858.358 | -0.162 | 0.033 | -4.924 | 8.47E-07 | 2.73E-05 | A230050P20Rik | protein_coding |
| ENSMUSG00000027002.13 | 28005.151 | 0.117 | 0.024 | 4.915 | 8.87E-07 | 2.85E-05 | Nckap1 | protein_coding |
| ENSMUSG00000061028.7 | 4790.275 | -0.152 | 0.031 | -4.913 | 8.97E-07 | 2.87E-05 | Clasrp | protein_coding |
| ENSMUSG00000001065.15 | 1672.734 | -0.219 | 0.045 | -4.912 | 9.00E-07 | 2.88E-05 | Zfp276 | protein_coding |
| ENSMUSG00000037475.15 | 3076.755 | 0.171 | 0.035 | 4.908 | 9.20E-07 | 2.93E-05 | Thoc2 | protein_coding |
| ENSMUSG00000060601.13 | 2186.335 | -0.210 | 0.043 | -4.906 | 9.28E-07 | 2.95E-05 | Nr1h2 | protein_coding |
| ENSMUSG00000029570.5 | 908.815 | -0.263 | 0.054 | -4.906 | 9.29E-07 | 2.95E-05 | Lfng | protein_coding |
| ENSMUSG00000026211.17 | 1272.041 | -0.211 | 0.043 | -4.903 | 9.45E-07 | 2.99E-05 | Obsl1 | protein_coding |
| ENSMUSG00000030590.15 | 1298.243 | -0.210 | 0.043 | -4.899 | 9.63E-07 | 3.04E-05 | Fam98c | protein_coding |
| ENSMUSG00000073062.3 | 575.373 | 0.283 | 0.058 | 4.899 | 9.66E-07 | 3.04E-05 | Zxdb | protein_coding |
| ENSMUSG00000040938.16 | 1202.488 | -0.239 | 0.049 | -4.894 | 9.88E-07 | 3.11E-05 | Slc16a11 | protein_coding |
| ENSMUSG00000019894.14 | 3098.387 | 0.172 | 0.035 | 4.893 | 9.93E-07 | 3.12E-05 | Slc6a15 | protein_coding |
| ENSMUSG00000021486.6 | 4014.659 | -0.153 | 0.031 | -4.891 | 1.00E-06 | 3.14E-05 | Prelid1 | protein_coding |
| ENSMUSG00000001802.16 | 4406.809 | -0.192 | 0.039 | -4.887 | 1.02E-06 | 3.19E-05 | Lrp3 | protein_coding |
| ENSMUSG00000020684.14 | 11478.503 | -0.166 | 0.034 | -4.886 | 1.03E-06 | 3.21E-05 | Rasl10b | protein_coding |
| ENSMUSG00000064368.1 | 1677.830 | 0.318 | 0.065 | 4.882 | 1.05E-06 | 3.26E-05 | mt-Nd6 | protein_coding |
| ENSMUSG00000047044.7 | 1084.476 | -0.246 | 0.050 | -4.876 | 1.08E-06 | 3.36E-05 | D030056L22Rik | protein_coding |
| ENSMUSG00000037416.12 | 4449.057 | 0.136 | 0.028 | 4.870 | 1.11E-06 | 3.45E-05 | Dmxl1 | protein_coding |
| ENSMUSG00000043542.12 | 2796.284 | 0.174 | 0.036 | 4.868 | 1.13E-06 | 3.48E-05 | Zc2hc1a | protein_coding |
| ENSMUSG00000048004.15 | 987.452 | 0.249 | 0.051 | 4.866 | 1.14E-06 | 3.51E-05 | Tmem196 | protein_coding |
| ENSMUSG00000039830.8 | 865.981 | -0.324 | 0.067 | -4.854 | 1.21E-06 | 3.72E-05 | Olig2 | protein_coding |
| ENSMUSG00000042423.9 | 2957.524 | -0.203 | 0.042 | -4.853 | 1.21E-06 | 3.72E-05 | Fbrs | protein_coding |
| ENSMUSG00000036565.7 | 14935.223 | -0.178 | 0.037 | -4.846 | 1.26E-06 | 3.86E-05 | Ttyh3 | protein_coding |
| ENSMUSG00000035283.4 | 934.890 | -0.284 | 0.059 | -4.843 | 1.28E-06 | 3.90E-05 | Adrb1 | protein_coding |
| ENSMUSG00000070436.12 | 1080.476 | -0.319 | 0.066 | -4.842 | 1.28E-06 | 3.90E-05 | Serpinh1 | protein_coding |
| ENSMUSG00000045180.13 | 4364.229 | 0.155 | 0.032 | 4.839 | 1.31E-06 | 3.97E-05 | Shroom2 | protein_coding |
| ENSMUSG00000075700.9 | 7753.872 | 0.142 | 0.029 | 4.834 | 1.34E-06 | 4.06E-05 | Selenot | protein_coding |
| ENSMUSG00000071073.4 | 1390.579 | -0.210 | 0.044 | -4.826 | 1.40E-06 | 4.21E-05 | Lrrc73 | protein_coding |
| ENSMUSG00000041203.9 | 1652.704 | -0.371 | 0.077 | -4.823 | 1.41E-06 | 4.26E-05 | 2310036O22Rik | protein_coding |
| ENSMUSG00000026824.11 | 5376.861 | 0.141 | 0.029 | 4.823 | 1.42E-06 | 4.26E-05 | Kcnj3 | protein_coding |

| | | | | | | | | |
|---|---|---|---|---|---|---|---|---|
| ENSMUSG00000025323.10 | 1241.555 | 0.224 | 0.046 | 4.822 | 1.42E-06 | 4.27E-05 | Sp4 | protein_coding |
| ENSMUSG00000023495.13 | 6842.935 | -0.166 | 0.034 | -4.821 | 1.43E-06 | 4.28E-05 | Pcbp4 | protein_coding |
| ENSMUSG00000021278.7 | 100.978 | -0.375 | 0.078 | -4.816 | 1.46E-06 | 4.37E-05 | Amn | protein_coding |
| ENSMUSG00000053310.11 | 27828.582 | -0.232 | 0.048 | -4.811 | 1.50E-06 | 4.47E-05 | Nrgn | protein_coding |
| ENSMUSG00000031848.15 | 694.391 | -0.314 | 0.065 | -4.809 | 1.52E-06 | 4.51E-05 | Lsm4 | protein_coding |
| ENSMUSG00000025982.13 | 17771.702 | 0.134 | 0.028 | 4.808 | 1.52E-06 | 4.51E-05 | Sf3b1 | protein_coding |
| ENSMUSG00000017291.13 | 8501.026 | 0.125 | 0.026 | 4.807 | 1.54E-06 | 4.55E-05 | Taok1 | protein_coding |
| ENSMUSG00000078578.9 | 10894.424 | 0.109 | 0.023 | 4.805 | 1.55E-06 | 4.57E-05 | Ube2d3 | protein_coding |
| ENSMUSG00000031840.12 | 21304.049 | -0.147 | 0.031 | -4.805 | 1.55E-06 | 4.57E-05 | Rab3a | protein_coding |
| ENSMUSG00000037400.17 | 7409.180 | 0.186 | 0.039 | 4.801 | 1.58E-06 | 4.64E-05 | Atp11b | protein_coding |
| ENSMUSG00000040681.14 | 2148.067 | -0.196 | 0.041 | -4.797 | 1.61E-06 | 4.73E-05 | Hmgn1 | protein_coding |
| ENSMUSG00000047143.3 | 61.797 | -0.356 | 0.074 | -4.795 | 1.63E-06 | NA | Dmrta2 | protein_coding |
| ENSMUSG00000007721.6 | 2184.727 | -0.203 | 0.042 | -4.794 | 1.64E-06 | 4.80E-05 | Ccdc124 | protein_coding |
| ENSMUSG00000022523.9 | 6306.080 | 0.162 | 0.034 | 4.793 | 1.65E-06 | 4.81E-05 | Fgf12 | protein_coding |
| ENSMUSG00000024856.10 | 922.780 | -0.236 | 0.049 | -4.787 | 1.69E-06 | 4.94E-05 | Cdk2ap2 | protein_coding |
| ENSMUSG00000032288.9 | 645.548 | -0.306 | 0.064 | -4.782 | 1.73E-06 | 5.04E-05 | Imp3 | protein_coding |
| ENSMUSG00000034724.17 | 2431.909 | 0.177 | 0.037 | 4.782 | 1.74E-06 | 5.04E-05 | Cnot6l | protein_coding |
| ENSMUSG00000035390.16 | 15931.575 | -0.159 | 0.033 | -4.782 | 1.74E-06 | 5.04E-05 | Brsk1 | protein_coding |
| ENSMUSG00000091337.3 | 8697.179 | 0.124 | 0.026 | 4.777 | 1.78E-06 | 5.14E-05 | Eid1 | protein_coding |
| ENSMUSG00000048249.14 | 3025.014 | 0.172 | 0.036 | 4.769 | 1.85E-06 | 5.34E-05 | Crebrf | protein_coding |
| ENSMUSG00000025860.14 | 5249.636 | 0.145 | 0.030 | 4.768 | 1.86E-06 | 5.37E-05 | Xiap | protein_coding |
| ENSMUSG00000059208.14 | 6162.945 | -0.144 | 0.030 | -4.760 | 1.93E-06 | 5.57E-05 | Hnrnpm | protein_coding |
| ENSMUSG00000000247.11 | 2006.169 | -0.268 | 0.056 | -4.759 | 1.95E-06 | 5.59E-05 | Lhx2 | protein_coding |
| ENSMUSG00000033763.14 | 11198.805 | -0.179 | 0.038 | -4.755 | 1.99E-06 | 5.69E-05 | Mtss1l | protein_coding |
| ENSMUSG00000022791.16 | 15942.079 | -0.168 | 0.035 | -4.752 | 2.01E-06 | 5.75E-05 | Tnk2 | protein_coding |
| ENSMUSG00000025208.8 | 1147.573 | -0.216 | 0.046 | -4.750 | 2.03E-06 | 5.80E-05 | Mrpl43 | protein_coding |
| ENSMUSG00000066607.5 | 6124.985 | -0.188 | 0.039 | -4.750 | 2.04E-06 | 5.80E-05 | 6030419C18Rik | protein_coding |
| ENSMUSG00000026904.17 | 15224.425 | 0.137 | 0.029 | 4.735 | 2.19E-06 | 6.21E-05 | Slc4a10 | protein_coding |
| ENSMUSG00000051185.9 | 1430.540 | -0.200 | 0.042 | -4.731 | 2.23E-06 | 6.33E-05 | Fam174a | protein_coding |
| ENSMUSG00000019188.16 | 3920.503 | -0.151 | 0.032 | -4.730 | 2.25E-06 | 6.37E-05 | H13 | protein_coding |
| ENSMUSG00000024750.10 | 6056.675 | 0.145 | 0.031 | 4.718 | 2.38E-06 | 6.71E-05 | Zfand5 | protein_coding |
| ENSMUSG00000029840.8 | 16026.815 | 0.122 | 0.026 | 4.716 | 2.41E-06 | 6.79E-05 | Mtpn | protein_coding |
| ENSMUSG00000019877.10 | 45123.913 | 0.122 | 0.026 | 4.711 | 2.47E-06 | 6.94E-05 | Serinc1 | protein_coding |

| | | | | | | | | |
|---|---|---|---|---|---|---|---|---|
| ENSMUSG00000019790.17 | 7045.484 | 0.145 | 0.031 | 4.707 | 2.51E-06 | 7.05E-05 | Stxbp5 | protein_coding |
| ENSMUSG00000059742.10 | 5538.595 | 0.186 | 0.040 | 4.706 | 2.53E-06 | 7.09E-05 | Kcnh7 | protein_coding |
| ENSMUSG00000026153.15 | 1816.296 | 0.177 | 0.038 | 4.705 | 2.54E-06 | 7.11E-05 | Fam135a | protein_coding |
| ENSMUSG00000025245.14 | 3365.111 | 0.144 | 0.031 | 4.700 | 2.60E-06 | 7.25E-05 | Lztfl1 | protein_coding |
| ENSMUSG00000033917.15 | 8206.649 | -0.137 | 0.029 | -4.696 | 2.65E-06 | 7.38E-05 | Gde1 | protein_coding |
| ENSMUSG00000020664.10 | 5140.353 | 0.128 | 0.027 | 4.695 | 2.66E-06 | 7.40E-05 | Dld | protein_coding |
| ENSMUSG00000027799.12 | 17367.439 | 0.107 | 0.023 | 4.693 | 2.70E-06 | 7.46E-05 | Nbea | protein_coding |
| ENSMUSG00000074247.10 | 3266.009 | -0.170 | 0.036 | -4.692 | 2.71E-06 | 7.49E-05 | Dda1 | protein_coding |
| ENSMUSG00000019158.9 | 510.387 | -0.331 | 0.071 | -4.684 | 2.81E-06 | 7.76E-05 | Tmem160 | protein_coding |
| ENSMUSG00000025656.17 | 22196.141 | 0.112 | 0.024 | 4.683 | 2.83E-06 | 7.78E-05 | Arhgef9 | protein_coding |
| ENSMUSG00000004364.14 | 6524.337 | 0.124 | 0.027 | 4.681 | 2.85E-06 | 7.83E-05 | Cul3 | protein_coding |
| ENSMUSG00000030869.11 | 1832.502 | -0.196 | 0.042 | -4.671 | 3.00E-06 | 8.23E-05 | Ndufab1 | protein_coding |
| ENSMUSG00000078348.4 | 876.837 | -0.245 | 0.052 | -4.663 | 3.11E-06 | 8.51E-05 | Sf3b5 | protein_coding |
| ENSMUSG00000090100.7 | 4611.823 | 0.143 | 0.031 | 4.663 | 3.11E-06 | 8.51E-05 | Ttbk2 | protein_coding |
| ENSMUSG00000045045.7 | 1931.610 | -0.229 | 0.049 | -4.659 | 3.18E-06 | 8.68E-05 | Lrfn4 | protein_coding |
| ENSMUSG00000001833.17 | 20175.465 | 0.161 | 0.035 | 4.657 | 3.20E-06 | 8.71E-05 | Sept7 | protein_coding |
| ENSMUSG00000016356.17 | 325.596 | -0.323 | 0.069 | -4.652 | 3.29E-06 | 8.93E-05 | Col20a1 | protein_coding |
| ENSMUSG00000045252.11 | 1473.132 | -0.202 | 0.043 | -4.650 | 3.31E-06 | 8.98E-05 | Zfp574 | protein_coding |
| ENSMUSG00000045948.8 | 1289.180 | -0.223 | 0.048 | -4.649 | 3.34E-06 | 9.04E-05 | Mrps12 | protein_coding |
| ENSMUSG00000020385.16 | 4178.366 | 0.144 | 0.031 | 4.648 | 3.36E-06 | 9.07E-05 | Clk4 | protein_coding |
| ENSMUSG00000047264.8 | 1198.834 | -0.257 | 0.055 | -4.644 | 3.41E-06 | 9.19E-05 | Zfp358 | protein_coding |
| ENSMUSG00000001240.13 | 470.613 | -0.321 | 0.069 | -4.643 | 3.43E-06 | 9.22E-05 | Ramp2 | protein_coding |
| ENSMUSG00000024940.10 | 1955.799 | -0.229 | 0.049 | -4.643 | 3.44E-06 | 9.24E-05 | Ltbp3 | protein_coding |
| ENSMUSG00000006262.15 | 1634.485 | 0.204 | 0.044 | 4.642 | 3.45E-06 | 9.25E-05 | Mob1b | protein_coding |
| ENSMUSG00000037703.14 | 14556.195 | -0.122 | 0.026 | -4.641 | 3.47E-06 | 9.29E-05 | Lzts3 | protein_coding |
| ENSMUSG00000036902.11 | 3218.860 | 0.156 | 0.034 | 4.633 | 3.61E-06 | 9.63E-05 | Neto2 | protein_coding |
| ENSMUSG00000020230.15 | 3639.145 | -0.149 | 0.032 | -4.628 | 3.69E-06 | 9.83E-05 | Prmt2 | protein_coding |
| ENSMUSG00000038894.7 | 5506.232 | -0.203 | 0.044 | -4.625 | 3.75E-06 | 9.96E-05 | Irs2 | protein_coding |
| ENSMUSG00000035711.4 | 819.414 | -0.272 | 0.059 | -4.624 | 3.77E-06 | 0.000100076 | Dok3 | protein_coding |
| ENSMUSG00000047260.4 | 1015.646 | -0.262 | 0.057 | -4.620 | 3.83E-06 | 0.000101432 | Emc6 | protein_coding |
| ENSMUSG00000031601.16 | 4603.816 | 0.149 | 0.032 | 4.618 | 3.87E-06 | 0.000102354 | Cnot7 | protein_coding |
| ENSMUSG00000030795.18 | 16057.247 | 0.116 | 0.025 | 4.615 | 3.93E-06 | 0.000103736 | Fus | protein_coding |
| ENSMUSG00000031532.6 | 9864.035 | -0.114 | 0.025 | -4.612 | 3.98E-06 | 0.000104878 | Saraf | protein_coding |

| | | | | | | | | |
|---|---|---|---|---|---|---|---|---|
| ENSMUSG00000047547.14 | 6542.985 | -0.144 | 0.031 | -4.610 | 4.03E-06 | 0.000105843 | Cltb | protein_coding |
| ENSMUSG00000026568.6 | 1975.408 | -0.204 | 0.044 | -4.609 | 4.05E-06 | 0.000106264 | Mpc2 | protein_coding |
| ENSMUSG00000054863.8 | 4024.588 | -0.158 | 0.034 | -4.608 | 4.06E-06 | 0.000106264 | Fam19a5 | protein_coding |
| ENSMUSG00000020516.15 | 2835.399 | 0.188 | 0.041 | 4.605 | 4.13E-06 | 0.000108077 | Rps6kb1 | protein_coding |
| ENSMUSG00000031808.7 | 5653.677 | -0.173 | 0.038 | -4.604 | 4.15E-06 | 0.000108224 | Slc27a1 | protein_coding |
| ENSMUSG00000033545.14 | 7345.622 | -0.159 | 0.034 | -4.603 | 4.17E-06 | 0.000108534 | Znrf1 | protein_coding |
| ENSMUSG00000020519.5 | 636.117 | -0.268 | 0.058 | -4.602 | 4.19E-06 | 0.000108815 | Sap30l | protein_coding |
| ENSMUSG00000006731.10 | 6768.022 | -0.177 | 0.038 | -4.602 | 4.19E-06 | 0.000108815 | B4galnt1 | protein_coding |
| ENSMUSG00000007603.8 | 4004.766 | -0.167 | 0.036 | -4.600 | 4.22E-06 | 0.000109398 | Dus3l | protein_coding |
| ENSMUSG00000027770.5 | 4331.871 | 0.154 | 0.033 | 4.599 | 4.24E-06 | 0.000109727 | Dhx36 | protein_coding |
| ENSMUSG00000029836.15 | 1422.449 | 0.222 | 0.048 | 4.590 | 4.43E-06 | 0.000114375 | Cbx3 | protein_coding |
| ENSMUSG00000044496.6 | 1250.320 | -0.210 | 0.046 | -4.585 | 4.54E-06 | 0.000116849 | 2510039O18Rik | protein_coding |
| ENSMUSG00000084883.2 | 3214.214 | -0.154 | 0.034 | -4.584 | 4.56E-06 | 0.000117123 | Ccdc85c | protein_coding |
| ENSMUSG00000007029.16 | 2620.861 | -0.177 | 0.039 | -4.583 | 4.58E-06 | 0.000117302 | Vars | protein_coding |
| ENSMUSG00000027834.15 | 15446.353 | 0.176 | 0.038 | 4.580 | 4.66E-06 | 0.000119075 | Serpini1 | protein_coding |
| ENSMUSG00000021072.12 | 1862.858 | 0.176 | 0.038 | 4.579 | 4.67E-06 | 0.000119333 | Tmx1 | protein_coding |
| ENSMUSG00000026207.16 | 9317.618 | -0.198 | 0.043 | -4.577 | 4.71E-06 | 0.00012008 | Speg | protein_coding |
| ENSMUSG00000038517.15 | 3557.348 | -0.190 | 0.042 | -4.577 | 4.72E-06 | 0.00012008 | Tbkbp1 | protein_coding |
| ENSMUSG00000050029.7 | 1637.095 | 0.185 | 0.040 | 4.575 | 4.77E-06 | 0.000121114 | Rap2c | protein_coding |
| ENSMUSG00000095407.1 | 600.076 | -0.295 | 0.064 | -4.572 | 4.82E-06 | 0.000122283 | Tmem200c | protein_coding |
| ENSMUSG00000062257.13 | 10301.174 | 0.111 | 0.024 | 4.571 | 4.86E-06 | 0.000123076 | Opcml | protein_coding |
| ENSMUSG00000030265.14 | 5480.667 | 0.134 | 0.030 | 4.556 | 5.21E-06 | 0.000131681 | Kras | protein_coding |
| ENSMUSG00000013833.15 | 2838.964 | -0.201 | 0.044 | -4.552 | 5.30E-06 | 0.000133685 | Med16 | protein_coding |
| ENSMUSG00000036894.3 | 2226.362 | -0.161 | 0.035 | -4.552 | 5.32E-06 | 0.00013394 | Rap2b | protein_coding |
| ENSMUSG00000058239.13 | 2221.399 | -0.230 | 0.050 | -4.551 | 5.33E-06 | 0.00013394 | Usf2 | protein_coding |
| ENSMUSG00000048696.11 | 665.995 | -0.311 | 0.068 | -4.548 | 5.41E-06 | 0.000135744 | Mex3d | protein_coding |
| ENSMUSG00000029455.14 | 3876.341 | -0.181 | 0.040 | -4.547 | 5.43E-06 | 0.000135908 | Aldh2 | protein_coding |
| ENSMUSG00000028020.16 | 8391.639 | 0.125 | 0.027 | 4.546 | 5.47E-06 | 0.000136763 | Glrb | protein_coding |
| ENSMUSG00000022674.14 | 1482.532 | 0.216 | 0.047 | 4.545 | 5.49E-06 | 0.000136928 | Ube2v2 | protein_coding |
| ENSMUSG00000041995.8 | 373.668 | -0.286 | 0.063 | -4.545 | 5.50E-06 | 0.000136938 | Zbed3 | protein_coding |
| ENSMUSG00000027499.12 | 8780.793 | 0.149 | 0.033 | 4.540 | 5.61E-06 | 0.000139523 | Pkia | protein_coding |
| ENSMUSG00000071862.2 | 4644.870 | 0.188 | 0.041 | 4.539 | 5.65E-06 | 0.000140139 | Lrrtm2 | protein_coding |
| ENSMUSG00000045009.9 | 3353.794 | -0.196 | 0.043 | -4.537 | 5.70E-06 | 0.000141146 | Prrt3 | protein_coding |

| | | | | | | | | |
|---|---|---|---|---|---|---|---|---|
| ENSMUSG00000043004.13 | 9114.897 | 0.143 | 0.031 | 4.530 | 5.89E-06 | 0.000145436 | Gng2 | protein_coding |
| ENSMUSG00000037373.12 | 7825.328 | -0.121 | 0.027 | -4.530 | 5.89E-06 | 0.000145436 | Ctbp1 | protein_coding |
| ENSMUSG00000038630.8 | 2580.223 | 0.187 | 0.041 | 4.524 | 6.06E-06 | 0.000149124 | Zkscan16 | protein_coding |
| ENSMUSG00000020484.18 | 5645.125 | -0.138 | 0.030 | -4.523 | 6.10E-06 | 0.000149888 | Xbp1 | protein_coding |
| ENSMUSG00000023020.2 | 1406.775 | -0.209 | 0.046 | -4.520 | 6.17E-06 | 0.000151298 | Cox14 | protein_coding |
| ENSMUSG00000024083.15 | 31156.328 | 0.121 | 0.027 | 4.518 | 6.24E-06 | 0.00015261 | Pja2 | protein_coding |
| ENSMUSG00000048047.3 | 1736.097 | 0.171 | 0.038 | 4.518 | 6.26E-06 | 0.000152787 | Zbtb33 | protein_coding |
| ENSMUSG00000043496.7 | 1953.831 | -0.179 | 0.040 | -4.516 | 6.31E-06 | 0.000153754 | Tril | protein_coding |
| ENSMUSG00000005506.16 | 13353.198 | 0.095 | 0.021 | 4.514 | 6.36E-06 | 0.000154784 | Celf1 | protein_coding |
| ENSMUSG00000028546.17 | 5077.304 | 0.159 | 0.035 | 4.514 | 6.37E-06 | 0.000154784 | Elavl4 | protein_coding |
| ENSMUSG00000020863.15 | 10148.131 | 0.107 | 0.024 | 4.513 | 6.40E-06 | 0.000155242 | Luc7l3 | protein_coding |
| ENSMUSG00000021218.10 | 10278.274 | 0.109 | 0.024 | 4.512 | 6.43E-06 | 0.000155755 | Gdi2 | protein_coding |
| ENSMUSG00000033434.15 | 1421.061 | -0.223 | 0.049 | -4.511 | 6.46E-06 | 0.00015628 | Gtpbp6 | protein_coding |
| ENSMUSG00000034560.6 | 2558.676 | 0.154 | 0.034 | 4.505 | 6.62E-06 | 0.000159685 | Washc4 | protein_coding |
| ENSMUSG00000023909.3 | 2443.085 | -0.173 | 0.038 | -4.505 | 6.63E-06 | 0.000159685 | Paqr4 | protein_coding |
| ENSMUSG00000056895.4 | 321.522 | -0.317 | 0.070 | -4.504 | 6.68E-06 | 0.000160654 | Hist3h2ba | protein_coding |
| ENSMUSG00000022533.13 | 4202.503 | 0.133 | 0.030 | 4.501 | 6.77E-06 | 0.000162515 | Atp13a3 | protein_coding |
| ENSMUSG00000068859.5 | 311.802 | -0.342 | 0.076 | -4.500 | 6.78E-06 | 0.000162622 | Sp9 | protein_coding |
| ENSMUSG00000035277.15 | 895.639 | -0.264 | 0.059 | -4.499 | 6.84E-06 | 0.000163641 | Arx | protein_coding |
| ENSMUSG00000037706.17 | 9050.516 | -0.134 | 0.030 | -4.498 | 6.86E-06 | 0.000163987 | Cd81 | protein_coding |
| ENSMUSG00000035847.15 | 29924.472 | 0.143 | 0.032 | 4.497 | 6.89E-06 | 0.000164307 | Ids | protein_coding |
| ENSMUSG00000033697.15 | 5123.079 | -0.151 | 0.034 | -4.496 | 6.93E-06 | 0.000164938 | Arhgap39 | protein_coding |
| ENSMUSG00000030428.16 | 24779.030 | -0.111 | 0.025 | -4.495 | 6.96E-06 | 0.00016538 | Ttyh1 | protein_coding |
| ENSMUSG00000022016.15 | 15407.076 | 0.116 | 0.026 | 4.491 | 7.08E-06 | 0.000168078 | Akap11 | protein_coding |
| ENSMUSG00000021929.8 | 4476.846 | 0.131 | 0.029 | 4.488 | 7.20E-06 | 0.000170433 | Kpna3 | protein_coding |
| ENSMUSG00000022055.7 | 17096.884 | -0.135 | 0.030 | -4.487 | 7.21E-06 | 0.000170433 | Nefl | protein_coding |
| ENSMUSG00000024480.7 | 1666.321 | 0.175 | 0.039 | 4.486 | 7.27E-06 | 0.000171703 | Ap3s1 | protein_coding |
| ENSMUSG00000020198.8 | 12550.656 | -0.109 | 0.024 | -4.479 | 7.51E-06 | 0.000177063 | Ap3d1 | protein_coding |
| ENSMUSG00000021986.7 | 1210.314 | -0.221 | 0.049 | -4.463 | 8.09E-06 | 0.000190245 | Amer2 | protein_coding |
| ENSMUSG00000060860.8 | 716.132 | -0.237 | 0.053 | -4.460 | 8.19E-06 | 0.000192468 | Ube2s | protein_coding |
| ENSMUSG00000055373.8 | 8076.292 | 0.205 | 0.046 | 4.456 | 8.36E-06 | 0.000196117 | Fut9 | protein_coding |
| ENSMUSG00000026721.15 | 7315.541 | 0.153 | 0.034 | 4.453 | 8.48E-06 | 0.000198542 | Rabgap1l | protein_coding |
| ENSMUSG00000039753.16 | 5065.642 | 0.163 | 0.037 | 4.450 | 8.59E-06 | 0.000200829 | Fbxl5 | protein_coding |

| | | | | | | | | |
|---|---|---|---|---|---|---|---|---|
| ENSMUSG00000003863.18 | 9987.005 | -0.154 | 0.035 | -4.447 | 8.70E-06 | 0.000203089 | Ppfia3 | protein_coding |
| ENSMUSG00000046546.3 | 627.519 | -0.283 | 0.064 | -4.445 | 8.78E-06 | 0.000204521 | Fam43a | protein_coding |
| ENSMUSG00000025964.15 | 6403.633 | 0.116 | 0.026 | 4.444 | 8.81E-06 | 0.000204922 | Adam23 | protein_coding |
| ENSMUSG00000021448.7 | 2913.272 | -0.205 | 0.046 | -4.444 | 8.83E-06 | 0.000204922 | Shc3 | protein_coding |
| ENSMUSG00000013736.16 | 1778.055 | 0.202 | 0.045 | 4.444 | 8.84E-06 | 0.000204922 | Trnt1 | protein_coding |
| ENSMUSG00000050212.4 | 51.204 | -0.331 | 0.075 | -4.438 | 9.09E-06 | NA | Eva1b | protein_coding |
| ENSMUSG00000029571.14 | 8178.343 | 0.140 | 0.032 | 4.437 | 9.11E-06 | 0.000210555 | Tmem106b | protein_coding |
| ENSMUSG00000028248.15 | 15642.569 | 0.120 | 0.027 | 4.437 | 9.11E-06 | 0.000210555 | Pnisr | protein_coding |
| ENSMUSG00000033597.8 | 11960.629 | -0.183 | 0.041 | -4.435 | 9.21E-06 | 0.000212492 | Caskin1 | protein_coding |
| ENSMUSG00000028861.13 | 1115.207 | -0.200 | 0.045 | -4.429 | 9.46E-06 | 0.000217932 | Mrps15 | protein_coding |
| ENSMUSG00000031770.16 | 2880.839 | -0.164 | 0.037 | -4.428 | 9.49E-06 | 0.000218217 | Herpud1 | protein_coding |
| ENSMUSG00000031201.17 | 1463.773 | 0.204 | 0.046 | 4.425 | 9.62E-06 | 0.00022056 | Brcc3 | protein_coding |
| ENSMUSG00000067194.6 | 7792.697 | 0.125 | 0.028 | 4.423 | 9.75E-06 | 0.00022317 | Eif1ax | protein_coding |
| ENSMUSG00000003948.17 | 7125.403 | 0.149 | 0.034 | 4.422 | 9.78E-06 | 0.000223459 | Mmd | protein_coding |
| ENSMUSG00000030019.6 | 1165.017 | -0.222 | 0.050 | -4.421 | 9.82E-06 | 0.000223954 | Fbxl14 | protein_coding |
| ENSMUSG00000034993.7 | 2137.182 | -0.246 | 0.056 | -4.417 | 1.00E-05 | 0.000227796 | Vat1 | protein_coding |
| ENSMUSG00000000581.8 | 1412.740 | 0.187 | 0.042 | 4.415 | 1.01E-05 | 0.000228819 | C1d | protein_coding |
| ENSMUSG00000034209.4 | 1747.091 | -0.216 | 0.049 | -4.415 | 1.01E-05 | 0.000229164 | Rasl10a | protein_coding |
| ENSMUSG00000045427.13 | 5664.287 | 0.113 | 0.026 | 4.413 | 1.02E-05 | 0.000230862 | Hnrnph2 | protein_coding |
| ENSMUSG00000029804.16 | 10451.836 | 0.117 | 0.027 | 4.411 | 1.03E-05 | 0.000232308 | Herc3 | protein_coding |
| ENSMUSG00000034818.16 | 18439.030 | -0.122 | 0.028 | -4.407 | 1.05E-05 | 0.000236272 | Celf5 | protein_coding |
| ENSMUSG00000006476.19 | 26311.380 | -0.148 | 0.034 | -4.405 | 1.06E-05 | 0.000237847 | Nsmf | protein_coding |
| ENSMUSG00000029599.13 | 2489.169 | -0.156 | 0.035 | -4.401 | 1.08E-05 | 0.000242271 | Ddx54 | protein_coding |
| ENSMUSG00000020018.6 | 472.875 | -0.280 | 0.064 | -4.392 | 1.12E-05 | 0.000251563 | Snrpf | protein_coding |
| ENSMUSG00000034799.16 | 28898.141 | -0.137 | 0.031 | -4.392 | 1.12E-05 | 0.000251563 | Unc13a | protein_coding |
| ENSMUSG00000022905.12 | 7678.400 | 0.117 | 0.027 | 4.392 | 1.12E-05 | 0.000251563 | Kpna1 | protein_coding |
| ENSMUSG00000037907.16 | 5736.492 | -0.176 | 0.040 | -4.390 | 1.14E-05 | 0.000253539 | Ankrd13b | protein_coding |
| ENSMUSG00000013662.5 | 4588.092 | 0.130 | 0.030 | 4.384 | 1.17E-05 | 0.000259846 | Atad1 | protein_coding |
| ENSMUSG00000035864.14 | 81951.713 | 0.086 | 0.020 | 4.382 | 1.18E-05 | 0.000262211 | Syt1 | protein_coding |
| ENSMUSG00000031327.10 | 2238.036 | 0.178 | 0.041 | 4.381 | 1.18E-05 | 0.000262211 | Chic1 | protein_coding |
| ENSMUSG00000020153.14 | 2840.089 | -0.166 | 0.038 | -4.380 | 1.19E-05 | 0.00026326 | Ndufs7 | protein_coding |
| ENSMUSG00000034390.16 | 10701.327 | -0.136 | 0.031 | -4.380 | 1.19E-05 | 0.00026326 | Cmip | protein_coding |
| ENSMUSG00000042742.7 | 1676.939 | 0.181 | 0.041 | 4.369 | 1.25E-05 | 0.000276209 | Bmt2 | protein_coding |

| | | | | | | | | |
|---|---|---|---|---|---|---|---|---|
| ENSMUSG00000069565.12 | 1852.233 | -0.192 | 0.044 | -4.368 | 1.25E-05 | 0.000276593 | Dazap1 | protein_coding |
| ENSMUSG00000069662.5 | 5724.284 | 0.148 | 0.034 | 4.368 | 1.25E-05 | 0.000276641 | Marcks | protein_coding |
| ENSMUSG00000036873.13 | 965.281 | -0.222 | 0.051 | -4.367 | 1.26E-05 | 0.000277121 | 2410004B18Rik | protein_coding |
| ENSMUSG00000052837.6 | 2384.262 | -0.331 | 0.076 | -4.366 | 1.27E-05 | 0.000278709 | Junb | protein_coding |
| ENSMUSG00000022788.16 | 1769.344 | 0.183 | 0.042 | 4.365 | 1.27E-05 | 0.000278945 | Fgd4 | protein_coding |
| ENSMUSG00000030122.12 | 13277.412 | -0.141 | 0.032 | -4.364 | 1.28E-05 | 0.000280127 | Ptms | protein_coding |
| ENSMUSG00000028458.12 | 3921.537 | -0.176 | 0.040 | -4.363 | 1.28E-05 | 0.000280127 | Tesk1 | protein_coding |
| ENSMUSG00000062014.12 | 10151.507 | 0.126 | 0.029 | 4.363 | 1.28E-05 | 0.000280731 | Gmfb | protein_coding |
| ENSMUSG00000052759.7 | 102.270 | -0.340 | 0.078 | -4.361 | 1.29E-05 | 0.000282183 | Gpr25 | protein_coding |
| ENSMUSG00000020189.14 | 6913.507 | 0.154 | 0.035 | 4.359 | 1.30E-05 | 0.000283992 | Osbpl8 | protein_coding |
| ENSMUSG00000020311.17 | 2031.130 | 0.172 | 0.039 | 4.358 | 1.31E-05 | 0.000285575 | Erlec1 | protein_coding |
| ENSMUSG00000025854.15 | 1644.819 | -0.209 | 0.048 | -4.357 | 1.32E-05 | 0.000286563 | Fam20c | protein_coding |
| ENSMUSG00000037990.18 | 2964.628 | -0.151 | 0.035 | -4.350 | 1.36E-05 | 0.000295356 | Sh3rf3 | protein_coding |
| ENSMUSG00000020733.3 | 1627.682 | -0.179 | 0.041 | -4.347 | 1.38E-05 | 0.000298669 | Slc9a3r1 | protein_coding |
| ENSMUSG00000027895.9 | 4259.428 | -0.163 | 0.037 | -4.345 | 1.40E-05 | 0.000301629 | Kcnc4 | protein_coding |
| ENSMUSG00000061518.10 | 4540.822 | -0.176 | 0.040 | -4.342 | 1.41E-05 | 0.000304655 | Cox5b | protein_coding |
| ENSMUSG00000003068.16 | 2864.272 | -0.155 | 0.036 | -4.341 | 1.42E-05 | 0.000306021 | Stk11 | protein_coding |
| ENSMUSG00000020644.9 | 2510.015 | 0.168 | 0.039 | 4.340 | 1.43E-05 | 0.000306748 | Id2 | protein_coding |
| ENSMUSG00000019370.10 | 48977.554 | -0.110 | 0.025 | -4.339 | 1.43E-05 | 0.000307524 | Calm3 | protein_coding |
| ENSMUSG00000056076.13 | 4572.935 | -0.126 | 0.029 | -4.333 | 1.47E-05 | 0.000315347 | Eif3b | protein_coding |
| ENSMUSG00000075318.12 | 20326.797 | 0.113 | 0.026 | 4.327 | 1.51E-05 | 0.000323474 | Scn2a | protein_coding |
| ENSMUSG00000032598.8 | 4467.029 | -0.148 | 0.034 | -4.326 | 1.52E-05 | 0.000325315 | Nckipsd | protein_coding |
| ENSMUSG00000022018.7 | 400.258 | -0.314 | 0.073 | -4.325 | 1.53E-05 | 0.000325993 | Rgcc | protein_coding |
| ENSMUSG00000019804.12 | 4526.883 | -0.173 | 0.040 | -4.322 | 1.55E-05 | 0.000330308 | Snx3 | protein_coding |
| ENSMUSG00000037492.16 | 2313.850 | 0.162 | 0.037 | 4.321 | 1.55E-05 | 0.000330308 | Zmat4 | protein_coding |
| ENSMUSG00000039478.15 | 4192.281 | 0.132 | 0.031 | 4.319 | 1.57E-05 | 0.000333283 | Micu3 | protein_coding |
| ENSMUSG00000003308.15 | 3154.709 | -0.158 | 0.036 | -4.318 | 1.57E-05 | 0.000333553 | Keap1 | protein_coding |
| ENSMUSG00000021743.5 | 2664.576 | -0.207 | 0.048 | -4.313 | 1.61E-05 | 0.000341381 | Fezf2 | protein_coding |
| ENSMUSG00000029211.11 | 4683.663 | 0.179 | 0.042 | 4.308 | 1.65E-05 | 0.000349022 | Gabra4 | protein_coding |
| ENSMUSG00000031988.10 | 6278.764 | -0.112 | 0.026 | -4.304 | 1.68E-05 | 0.000353949 | Vps26b | protein_coding |
| ENSMUSG00000003199.16 | 3136.118 | -0.154 | 0.036 | -4.302 | 1.69E-05 | 0.000357183 | Mpnd | protein_coding |
| ENSMUSG00000017548.15 | 2011.574 | 0.160 | 0.037 | 4.300 | 1.70E-05 | 0.000359017 | Suz12 | protein_coding |
| ENSMUSG00000024245.4 | 3806.038 | -0.163 | 0.038 | -4.299 | 1.72E-05 | 0.000361009 | Tmem178 | protein_coding |

| | | | | | | | | |
|---|---|---|---|---|---|---|---|---|
| ENSMUSG00000037014.4 | 1005.966 | -0.231 | 0.054 | -4.296 | 1.74E-05 | 0.000364634 | Sstr4 | protein_coding |
| ENSMUSG00000049097.9 | 3011.872 | -0.191 | 0.044 | -4.294 | 1.75E-05 | 0.000367564 | Ankrd34a | protein_coding |
| ENSMUSG00000038172.14 | 4929.237 | 0.129 | 0.030 | 4.294 | 1.76E-05 | 0.000367704 | Ttc39b | protein_coding |
| ENSMUSG00000029060.17 | 7297.864 | -0.182 | 0.042 | -4.293 | 1.77E-05 | 0.000369177 | Mib2 | protein_coding |
| ENSMUSG00000047036.8 | 12837.570 | 0.121 | 0.028 | 4.291 | 1.78E-05 | 0.000371372 | Zfp445 | protein_coding |
| ENSMUSG00000015937.14 | 3022.496 | -0.149 | 0.035 | -4.288 | 1.80E-05 | 0.000376126 | H2afy | protein_coding |
| ENSMUSG00000011257.19 | 1585.018 | -0.198 | 0.046 | -4.287 | 1.81E-05 | 0.000377661 | Pabpc4 | protein_coding |
| ENSMUSG00000052957.7 | 260.629 | -0.309 | 0.072 | -4.284 | 1.83E-05 | 0.000381259 | Gas1 | protein_coding |
| ENSMUSG00000049265.7 | 1449.372 | -0.206 | 0.048 | -4.284 | 1.84E-05 | 0.000381862 | Kcnk3 | protein_coding |
| ENSMUSG00000039041.15 | 2054.062 | -0.163 | 0.038 | -4.283 | 1.84E-05 | 0.000381933 | Adrm1 | protein_coding |
| ENSMUSG00000037933.16 | 6487.731 | -0.129 | 0.030 | -4.282 | 1.85E-05 | 0.000383869 | Bicd2 | protein_coding |
| ENSMUSG00000024966.8 | 8582.309 | -0.153 | 0.036 | -4.279 | 1.88E-05 | 0.000388025 | Stip1 | protein_coding |
| ENSMUSG00000020388.12 | 336.076 | -0.279 | 0.065 | -4.278 | 1.88E-05 | 0.000388724 | Pdlim4 | protein_coding |
| ENSMUSG00000020620.14 | 1924.488 | 0.159 | 0.037 | 4.277 | 1.89E-05 | 0.000390028 | Abca8b | protein_coding |
| ENSMUSG00000030087.11 | 1344.089 | -0.193 | 0.045 | -4.276 | 1.91E-05 | 0.000392009 | Klf15 | protein_coding |
| ENSMUSG00000031342.17 | 39572.714 | 0.119 | 0.028 | 4.269 | 1.96E-05 | 0.00040268 | Gpm6b | protein_coding |
| ENSMUSG00000013236.16 | 27146.871 | -0.134 | 0.031 | -4.267 | 1.98E-05 | 0.000406718 | Ptprs | protein_coding |
| ENSMUSG00000029413.14 | 1390.187 | -0.185 | 0.043 | -4.265 | 2.00E-05 | 0.000408877 | Naaa | protein_coding |
| ENSMUSG00000038128.6 | 8906.811 | 0.150 | 0.035 | 4.262 | 2.02E-05 | 0.000413875 | Camk4 | protein_coding |
| ENSMUSG00000020235.16 | 3274.937 | -0.141 | 0.033 | -4.257 | 2.07E-05 | 0.000423816 | Fzr1 | protein_coding |
| ENSMUSG00000032589.14 | 40459.199 | -0.155 | 0.037 | -4.252 | 2.12E-05 | 0.00043149 | Bsn | protein_coding |
| ENSMUSG00000055296.14 | 6748.077 | 0.121 | 0.028 | 4.251 | 2.13E-05 | 0.00043329 | Tmem245 | protein_coding |
| ENSMUSG00000002771.12 | 1042.647 | -0.250 | 0.059 | -4.246 | 2.17E-05 | 0.000441663 | Grin2d | protein_coding |
| ENSMUSG00000032239.10 | 892.081 | -0.241 | 0.057 | -4.244 | 2.20E-05 | 0.000446137 | Rp9 | protein_coding |
| ENSMUSG00000026959.13 | 29292.710 | -0.125 | 0.030 | -4.243 | 2.21E-05 | 0.000447187 | Grin1 | protein_coding |
| ENSMUSG00000094870.7 | 2011.578 | 0.175 | 0.041 | 4.242 | 2.21E-05 | 0.000447762 | Zfp131 | protein_coding |
| ENSMUSG00000062563.15 | 988.821 | -0.204 | 0.048 | -4.240 | 2.23E-05 | 0.000451897 | Cys1 | protein_coding |
| ENSMUSG00000032293.8 | 3859.027 | 0.137 | 0.032 | 4.236 | 2.28E-05 | 0.000459809 | Ireb2 | protein_coding |
| ENSMUSG00000059824.12 | 2799.446 | -0.183 | 0.043 | -4.235 | 2.28E-05 | 0.000459809 | Dbp | protein_coding |
| ENSMUSG00000000078.7 | 2735.456 | 0.147 | 0.035 | 4.232 | 2.32E-05 | 0.00046661 | Klf6 | protein_coding |
| ENSMUSG00000047126.17 | 34456.705 | 0.106 | 0.025 | 4.231 | 2.32E-05 | 0.000466772 | Cltc | protein_coding |
| ENSMUSG00000024261.5 | 12393.616 | 0.119 | 0.028 | 4.226 | 2.38E-05 | 0.000477297 | Syt4 | protein_coding |
| ENSMUSG00000004934.14 | 1057.440 | -0.221 | 0.052 | -4.224 | 2.40E-05 | 0.000480258 | Pias4 | protein_coding |

| | | | | | | | | |
|---|---|---|---|---|---|---|---|---|
| ENSMUSG00000017221.14 | 5706.538 | -0.114 | 0.027 | -4.218 | 2.46E-05 | 0.00049307 | Psmd3 | protein_coding |
| ENSMUSG00000060275.11 | 388.874 | -0.273 | 0.065 | -4.211 | 2.54E-05 | 0.000507677 | Nrg2 | protein_coding |
| ENSMUSG00000030409.15 | 1512.995 | -0.217 | 0.052 | -4.209 | 2.57E-05 | 0.000512652 | Dmpk | protein_coding |
| ENSMUSG00000024286.7 | 5882.664 | 0.131 | 0.031 | 4.200 | 2.67E-05 | 0.000532722 | Ccny | protein_coding |
| ENSMUSG00000074358.9 | 392.893 | -0.277 | 0.066 | -4.199 | 2.68E-05 | 0.000533091 | Ccdc61 | protein_coding |
| ENSMUSG00000045103.17 | 4424.498 | 0.119 | 0.028 | 4.199 | 2.68E-05 | 0.000533091 | Dmd | protein_coding |
| ENSMUSG00000036138.16 | 3507.773 | -0.158 | 0.038 | -4.198 | 2.69E-05 | 0.000533884 | Acaa1a | protein_coding |
| ENSMUSG00000031239.5 | 1736.406 | 0.177 | 0.042 | 4.192 | 2.76E-05 | 0.000547255 | Itm2a | protein_coding |
| ENSMUSG00000003226.7 | 9429.758 | 0.112 | 0.027 | 4.190 | 2.79E-05 | 0.000551715 | Ranbp2 | protein_coding |
| ENSMUSG00000053716.9 | 3136.885 | -0.175 | 0.042 | -4.187 | 2.83E-05 | 0.00055887 | Dusp7 | protein_coding |
| ENSMUSG00000032297.11 | 1513.449 | -0.225 | 0.054 | -4.186 | 2.84E-05 | 0.000559749 | Celf6 | protein_coding |
| ENSMUSG00000040957.14 | 1077.296 | -0.225 | 0.054 | -4.186 | 2.84E-05 | 0.000559749 | Cables1 | protein_coding |
| ENSMUSG00000020135.13 | 11839.130 | -0.160 | 0.038 | -4.185 | 2.86E-05 | 0.00056206 | Apc2 | protein_coding |
| ENSMUSG00000034880.7 | 466.094 | -0.280 | 0.067 | -4.177 | 2.95E-05 | 0.000580158 | Mrpl34 | protein_coding |
| ENSMUSG00000018800.14 | 3119.502 | 0.142 | 0.034 | 4.177 | 2.96E-05 | 0.000580158 | Abca5 | protein_coding |
| ENSMUSG00000051495.7 | 1462.992 | -0.191 | 0.046 | -4.176 | 2.96E-05 | 0.000580562 | Irf2bp2 | protein_coding |
| ENSMUSG00000024212.8 | 4763.208 | -0.133 | 0.032 | -4.175 | 2.98E-05 | 0.000581535 | Mllt1 | protein_coding |
| ENSMUSG00000026484.13 | 1334.780 | 0.196 | 0.047 | 4.175 | 2.98E-05 | 0.000581535 | Rnf2 | protein_coding |
| ENSMUSG00000003528.14 | 1484.074 | -0.192 | 0.046 | -4.175 | 2.98E-05 | 0.000581535 | Slc25a1 | protein_coding |
| ENSMUSG00000031242.7 | 3444.417 | 0.155 | 0.037 | 4.173 | 3.00E-05 | 0.000584929 | 2610002M06Rik | protein_coding |
| ENSMUSG00000017412.15 | 12016.001 | 0.146 | 0.035 | 4.169 | 3.06E-05 | 0.000595306 | Cacnb4 | protein_coding |
| ENSMUSG00000028524.21 | 9249.259 | 0.096 | 0.023 | 4.169 | 3.06E-05 | 0.000595306 | Sgip1 | protein_coding |
| ENSMUSG00000078570.10 | 939.177 | -0.225 | 0.054 | -4.168 | 3.07E-05 | 0.000596795 | 1110065P20Rik | protein_coding |
| ENSMUSG00000003345.16 | 6717.407 | -0.150 | 0.036 | -4.167 | 3.09E-05 | 0.000598887 | Csnk1g2 | protein_coding |
| ENSMUSG00000060279.14 | 15957.828 | -0.124 | 0.030 | -4.163 | 3.14E-05 | 0.000607212 | Ap2a1 | protein_coding |
| ENSMUSG00000000326.12 | 3676.874 | -0.126 | 0.030 | -4.159 | 3.20E-05 | 0.00061871 | Comt | protein_coding |
| ENSMUSG00000014782.16 | 457.830 | -0.276 | 0.066 | -4.154 | 3.26E-05 | 0.0006297 | Plekhg4 | protein_coding |
| ENSMUSG00000028572.13 | 5780.891 | 0.122 | 0.029 | 4.154 | 3.27E-05 | 0.000630571 | Hook1 | protein_coding |
| ENSMUSG00000022789.14 | 12472.224 | 0.096 | 0.023 | 4.153 | 3.27E-05 | 0.000630571 | Dnm1l | protein_coding |
| ENSMUSG00000028212.16 | 227.726 | 0.303 | 0.073 | 4.152 | 3.29E-05 | 0.000632572 | Ccne2 | protein_coding |
| ENSMUSG00000036948.17 | 3214.988 | -0.181 | 0.044 | -4.152 | 3.29E-05 | 0.000632572 | BC037034 | protein_coding |
| ENSMUSG00000041351.16 | 16365.580 | -0.133 | 0.032 | -4.150 | 3.33E-05 | 0.000638546 | Rap1gap | protein_coding |
| ENSMUSG00000027109.16 | 2620.821 | 0.175 | 0.042 | 4.149 | 3.34E-05 | 0.000639651 | Sp3 | protein_coding |

| | | | | | | | | |
|---|---|---|---|---|---|---|---|---|
| ENSMUSG00000047181.12 | 2922.636 | -0.213 | 0.051 | -4.147 | 3.36E-05 | 0.00064256 | Samd14 | protein_coding |
| ENSMUSG00000012422.14 | 2625.639 | 0.145 | 0.035 | 4.147 | 3.36E-05 | 0.00064256 | Tmem167 | protein_coding |
| ENSMUSG00000007812.16 | 1771.830 | 0.157 | 0.038 | 4.144 | 3.41E-05 | 0.000649331 | Zfp655 | protein_coding |
| ENSMUSG00000006676.16 | 11511.042 | -0.123 | 0.030 | -4.141 | 3.46E-05 | 0.00065818 | Usp19 | protein_coding |
| ENSMUSG00000019428.16 | 14967.886 | -0.153 | 0.037 | -4.135 | 3.56E-05 | 0.000675418 | Fkbp8 | protein_coding |
| ENSMUSG00000052917.14 | 2071.501 | 0.173 | 0.042 | 4.131 | 3.61E-05 | 0.000683845 | Senp7 | protein_coding |
| ENSMUSG00000046791.8 | 664.713 | -0.219 | 0.053 | -4.126 | 3.70E-05 | 0.000700254 | Riox1 | protein_coding |
| ENSMUSG00000035203.16 | 18960.868 | -0.193 | 0.047 | -4.125 | 3.71E-05 | 0.000701119 | Epn1 | protein_coding |
| ENSMUSG00000052133.16 | 2091.271 | -0.165 | 0.040 | -4.124 | 3.72E-05 | 0.000702311 | Sema5b | protein_coding |
| ENSMUSG00000036815.16 | 8615.556 | 0.108 | 0.026 | 4.124 | 3.73E-05 | 0.000704015 | Dpp10 | protein_coding |
| ENSMUSG00000020859.16 | 21485.161 | 0.085 | 0.021 | 4.120 | 3.79E-05 | 0.000713206 | Spag9 | protein_coding |
| ENSMUSG00000040640.10 | 8036.130 | 0.137 | 0.033 | 4.118 | 3.82E-05 | 0.000718337 | Erc2 | protein_coding |
| ENSMUSG00000031167.16 | 3869.963 | 0.142 | 0.035 | 4.118 | 3.82E-05 | 0.000718337 | Rbm3 | protein_coding |
| ENSMUSG00000002968.8 | 4758.469 | -0.166 | 0.040 | -4.116 | 3.85E-05 | 0.000721817 | Med25 | protein_coding |
| ENSMUSG00000062078.14 | 13028.032 | 0.169 | 0.041 | 4.116 | 3.85E-05 | 0.000721817 | Qk | protein_coding |
| ENSMUSG00000008489.18 | 4231.942 | 0.152 | 0.037 | 4.116 | 3.86E-05 | 0.000721817 | Elavl2 | protein_coding |
| ENSMUSG00000001986.16 | 15880.195 | 0.125 | 0.030 | 4.109 | 3.98E-05 | 0.000743749 | Gria3 | protein_coding |
| ENSMUSG00000000531.4 | 2049.137 | -0.214 | 0.052 | -4.108 | 3.99E-05 | 0.000743749 | Grasp | protein_coding |
| ENSMUSG00000022800.13 | 4196.382 | 0.122 | 0.030 | 4.108 | 3.99E-05 | 0.000743749 | Fyttd1 | protein_coding |
| ENSMUSG00000031360.14 | 2461.223 | 0.152 | 0.037 | 4.108 | 3.99E-05 | 0.000743749 | Ctps2 | protein_coding |
| ENSMUSG00000040760.10 | 6638.965 | 0.107 | 0.026 | 4.107 | 4.02E-05 | 0.000744945 | Appl1 | protein_coding |
| ENSMUSG00000059674.6 | 837.640 | -0.247 | 0.060 | -4.107 | 4.02E-05 | 0.000744945 | Cdh24 | protein_coding |
| ENSMUSG00000040563.13 | 9375.418 | -0.178 | 0.043 | -4.106 | 4.02E-05 | 0.000744945 | Plppr2 | protein_coding |
| ENSMUSG00000021266.16 | 4587.164 | -0.120 | 0.029 | -4.102 | 4.09E-05 | 0.000757728 | Wars | protein_coding |
| ENSMUSG00000039770.5 | 3249.533 | 0.143 | 0.035 | 4.098 | 4.17E-05 | 0.000770811 | Ypel5 | protein_coding |
| ENSMUSG00000033943.15 | 5184.026 | 0.119 | 0.029 | 4.097 | 4.19E-05 | 0.000774115 | Mga | protein_coding |
| ENSMUSG00000031644.19 | 3130.640 | 0.137 | 0.033 | 4.094 | 4.24E-05 | 0.000781971 | Nek1 | protein_coding |
| ENSMUSG00000021484.7 | 4572.064 | -0.127 | 0.031 | -4.093 | 4.26E-05 | 0.000785302 | Lman2 | protein_coding |
| ENSMUSG00000059142.15 | 975.835 | 0.215 | 0.052 | 4.092 | 4.29E-05 | 0.000788438 | Zfp945 | protein_coding |
| ENSMUSG00000032336.17 | 24772.810 | 0.121 | 0.030 | 4.089 | 4.33E-05 | 0.000795748 | Nptn | protein_coding |
| ENSMUSG00000005148.7 | 383.648 | -0.269 | 0.066 | -4.086 | 4.38E-05 | 0.000804206 | Klf5 | protein_coding |
| ENSMUSG00000042133.16 | 3739.601 | 0.134 | 0.033 | 4.086 | 4.40E-05 | 0.000805013 | Ppig | protein_coding |
| ENSMUSG00000027365.14 | 3378.412 | 0.154 | 0.038 | 4.086 | 4.40E-05 | 0.000805013 | Trpm7 | protein_coding |

| | | | | | | | | |
|---|---|---|---|---|---|---|---|---|
| ENSMUSG00000082229.1 | 5269.514 | 0.152 | 0.037 | 4.082 | 4.47E-05 | 0.000817958 | Nap1l2 | protein_coding |
| ENSMUSG00000029253.12 | 1289.513 | 0.187 | 0.046 | 4.075 | 4.61E-05 | 0.000841475 | Cenpc1 | protein_coding |
| ENSMUSG00000032253.13 | 3692.887 | 0.138 | 0.034 | 4.074 | 4.62E-05 | 0.00084326 | Phip | protein_coding |
| ENSMUSG00000072969.2 | 983.647 | 0.206 | 0.050 | 4.073 | 4.64E-05 | 0.00084425 | Armcx5 | protein_coding |
| ENSMUSG00000071074.9 | 1986.061 | -0.154 | 0.038 | -4.071 | 4.68E-05 | 0.000850285 | Yipf3 | protein_coding |
| ENSMUSG00000092083.4 | 1386.745 | 0.232 | 0.057 | 4.070 | 4.69E-05 | 0.000852111 | Kcnb2 | protein_coding |
| ENSMUSG00000026520.8 | 1090.736 | -0.197 | 0.049 | -4.069 | 4.72E-05 | 0.000855596 | Pycr2 | protein_coding |
| ENSMUSG00000024097.10 | 5477.947 | 0.118 | 0.029 | 4.067 | 4.76E-05 | 0.000861954 | Srsf7 | protein_coding |
| ENSMUSG00000028743.7 | 805.055 | -0.237 | 0.058 | -4.066 | 4.78E-05 | 0.000865547 | Akr7a5 | protein_coding |
| ENSMUSG00000062373.7 | 3804.516 | 0.139 | 0.034 | 4.065 | 4.81E-05 | 0.00086844 | Tmem65 | protein_coding |
| ENSMUSG00000034889.8 | 1231.390 | -0.203 | 0.050 | -4.063 | 4.84E-05 | 0.000873683 | Cactin | protein_coding |
| ENSMUSG00000074649.6 | 2536.414 | -0.168 | 0.041 | -4.061 | 4.88E-05 | 0.000879005 | BC029722 | protein_coding |
| ENSMUSG00000030465.19 | 35699.300 | 0.112 | 0.028 | 4.058 | 4.94E-05 | 0.000888505 | Psd3 | protein_coding |
| ENSMUSG00000031788.14 | 2966.609 | -0.173 | 0.043 | -4.058 | 4.94E-05 | 0.000888505 | Kifc3 | protein_coding |
| ENSMUSG00000058503.11 | 1510.184 | 0.175 | 0.043 | 4.057 | 4.96E-05 | 0.000889668 | Fam133b | protein_coding |
| ENSMUSG00000070639.5 | 5562.006 | 0.125 | 0.031 | 4.057 | 4.97E-05 | 0.000889668 | Lrrc8b | protein_coding |
| ENSMUSG00000060257.2 | 875.266 | -0.258 | 0.064 | -4.057 | 4.97E-05 | 0.000889668 | Scrt2 | protein_coding |
| ENSMUSG00000033707.8 | 1706.935 | -0.182 | 0.045 | -4.051 | 5.09E-05 | 0.000910658 | Lrrc24 | protein_coding |
| ENSMUSG00000025372.16 | 20267.635 | -0.109 | 0.027 | -4.045 | 5.23E-05 | 0.000933689 | Baiap2 | protein_coding |
| ENSMUSG00000062209.15 | 4757.202 | 0.137 | 0.034 | 4.045 | 5.24E-05 | 0.000935336 | Erbb4 | protein_coding |
| ENSMUSG00000051518.7 | 583.086 | -0.226 | 0.056 | -4.043 | 5.28E-05 | 0.000940818 | Rps19bp1 | protein_coding |
| ENSMUSG00000038539.15 | 553.353 | -0.242 | 0.060 | -4.042 | 5.29E-05 | 0.000942044 | Atf5 | protein_coding |
| ENSMUSG00000028833.13 | 74877.484 | -0.171 | 0.042 | -4.042 | 5.31E-05 | 0.0009429 | Ncdn | protein_coding |
| ENSMUSG00000034254.17 | 5873.582 | -0.134 | 0.033 | -4.041 | 5.33E-05 | 0.000946517 | Agpat1 | protein_coding |
| ENSMUSG00000029208.16 | 2455.072 | 0.161 | 0.040 | 4.039 | 5.36E-05 | 0.000949981 | Guf1 | protein_coding |
| ENSMUSG00000021936.13 | 6893.509 | 0.121 | 0.030 | 4.037 | 5.41E-05 | 0.000957892 | Mapk8 | protein_coding |
| ENSMUSG00000036442.8 | 1224.673 | -0.195 | 0.048 | -4.032 | 5.53E-05 | 0.000978312 | Thap11 | protein_coding |
| ENSMUSG00000027660.16 | 4301.093 | 0.136 | 0.034 | 4.030 | 5.58E-05 | 0.000984922 | Skil | protein_coding |
| ENSMUSG00000020238.14 | 3382.963 | -0.159 | 0.040 | -4.030 | 5.58E-05 | 0.000984922 | Ncln | protein_coding |
| ENSMUSG00000060002.14 | 2289.315 | 0.147 | 0.036 | 4.029 | 5.61E-05 | 0.000988402 | Chpt1 | protein_coding |
| ENSMUSG00000030660.9 | 1700.586 | 0.166 | 0.041 | 4.022 | 5.77E-05 | 0.001014592 | Pik3c2a | protein_coding |
| ENSMUSG00000022452.14 | 1843.344 | -0.167 | 0.042 | -4.021 | 5.79E-05 | 0.00101667 | Smdt1 | protein_coding |
| ENSMUSG00000030739.18 | 1801.879 | -0.182 | 0.045 | -4.019 | 5.83E-05 | 0.001023942 | Myh14 | protein_coding |

| | | | | | | | | |
|---|---|---|---|---|---|---|---|---|
| ENSMUSG00000061136.14 | 2131.757 | 0.150 | 0.037 | 4.018 | 5.87E-05 | 0.001028287 | Prpf40a | protein_coding |
| ENSMUSG00000055675.6 | 7191.365 | -0.113 | 0.028 | -4.014 | 5.98E-05 | 0.001046171 | Kbtbd11 | protein_coding |
| ENSMUSG00000034636.9 | 13777.982 | 0.096 | 0.024 | 4.011 | 6.04E-05 | 0.001056245 | Zyg11b | protein_coding |
| ENSMUSG00000038552.14 | 3920.748 | -0.128 | 0.032 | -4.008 | 6.11E-05 | 0.001067463 | Fndc4 | protein_coding |
| ENSMUSG00000001016.12 | 3901.820 | 0.124 | 0.031 | 4.007 | 6.15E-05 | 0.001072221 | Ilf2 | protein_coding |
| ENSMUSG00000031007.2 | 12846.466 | 0.118 | 0.029 | 4.005 | 6.19E-05 | 0.001078896 | Atp6ap2 | protein_coding |
| ENSMUSG00000010608.15 | 8299.323 | 0.107 | 0.027 | 4.003 | 6.24E-05 | 0.001086461 | Rbm25 | protein_coding |
| ENSMUSG00000029328.15 | 7926.574 | 0.114 | 0.028 | 4.002 | 6.28E-05 | 0.001091336 | Hnrnpdl | protein_coding |
| ENSMUSG00000059852.7 | 299.643 | -0.299 | 0.075 | -4.001 | 6.30E-05 | 0.001093023 | Kcng2 | protein_coding |
| ENSMUSG00000055022.14 | 19856.627 | 0.099 | 0.025 | 4.001 | 6.32E-05 | 0.001095371 | Cntn1 | protein_coding |
| ENSMUSG00000029071.16 | 6646.963 | -0.130 | 0.032 | -3.999 | 6.36E-05 | 0.001101371 | Dvl1 | protein_coding |
| ENSMUSG00000066392.11 | 14638.938 | 0.106 | 0.027 | 3.995 | 6.47E-05 | 0.001118634 | Nrxn3 | protein_coding |
| ENSMUSG00000025104.13 | 5721.694 | 0.126 | 0.032 | 3.991 | 6.58E-05 | 0.001137222 | Hdgfrp3 | protein_coding |
| ENSMUSG00000036550.16 | 7045.852 | 0.096 | 0.024 | 3.989 | 6.63E-05 | 0.001143655 | Cnot1 | protein_coding |
| ENSMUSG00000032740.16 | 4407.494 | 0.135 | 0.034 | 3.989 | 6.64E-05 | 0.001144045 | Ccdc88a | protein_coding |
| ENSMUSG00000021587.5 | 3960.658 | 0.142 | 0.036 | 3.988 | 6.67E-05 | 0.001147926 | Pcsk1 | protein_coding |
| ENSMUSG00000017418.13 | 1451.123 | 0.186 | 0.047 | 3.985 | 6.74E-05 | 0.001157767 | Arl5b | protein_coding |
| ENSMUSG00000013663.7 | 7340.641 | 0.126 | 0.032 | 3.984 | 6.77E-05 | 0.001161715 | Pten | protein_coding |
| ENSMUSG00000049577.15 | 1265.901 | -0.231 | 0.058 | -3.978 | 6.96E-05 | 0.001193063 | Zfpm1 | protein_coding |
| ENSMUSG00000052144.8 | 3270.947 | 0.146 | 0.037 | 3.977 | 6.97E-05 | 0.001193063 | Ppp4r2 | protein_coding |
| ENSMUSG00000053550.13 | 15473.744 | -0.137 | 0.035 | -3.977 | 6.98E-05 | 0.001193063 | Shisa7 | protein_coding |
| ENSMUSG00000052137.11 | 750.519 | 0.219 | 0.055 | 3.974 | 7.06E-05 | 0.001206155 | Rbm12b2 | protein_coding |
| ENSMUSG00000035621.13 | 3133.392 | -0.184 | 0.046 | -3.972 | 7.12E-05 | 0.001215266 | Midn | protein_coding |
| ENSMUSG00000001270.9 | 45300.289 | -0.125 | 0.031 | -3.971 | 7.15E-05 | 0.001217825 | Ckb | protein_coding |
| ENSMUSG00000070576.4 | 2122.775 | -0.173 | 0.044 | -3.971 | 7.15E-05 | 0.001217825 | Mn1 | protein_coding |
| ENSMUSG00000020948.9 | 1562.364 | 0.167 | 0.042 | 3.969 | 7.22E-05 | 0.001227643 | Klhl28 | protein_coding |
| ENSMUSG00000037243.17 | 2306.567 | -0.164 | 0.041 | -3.968 | 7.25E-05 | 0.001230676 | Zfp692 | protein_coding |
| ENSMUSG00000042104.18 | 2126.291 | 0.154 | 0.039 | 3.967 | 7.28E-05 | 0.001234713 | Uggt2 | protein_coding |
| ENSMUSG00000028403.15 | 3207.858 | 0.146 | 0.037 | 3.967 | 7.29E-05 | 0.001234713 | Zdhhc21 | protein_coding |
| ENSMUSG00000030605.15 | 8758.575 | -0.122 | 0.031 | -3.965 | 7.35E-05 | 0.001243446 | Mfge8 | protein_coding |
| ENSMUSG00000047141.5 | 1396.310 | 0.162 | 0.041 | 3.964 | 7.36E-05 | 0.001243446 | Zfp654 | protein_coding |
| ENSMUSG00000043099.4 | 167.706 | -0.308 | 0.078 | -3.963 | 7.40E-05 | 0.001248498 | Hic1 | protein_coding |
| ENSMUSG00000025907.14 | 5217.351 | 0.146 | 0.037 | 3.962 | 7.43E-05 | 0.001252859 | Rb1cc1 | protein_coding |

| | | | | | | | | |
|---|---|---|---|---|---|---|---|---|
| ENSMUSG00000004319.15 | 9346.959 | 0.097 | 0.025 | 3.962 | 7.44E-05 | 0.001253524 | Clcn3 | protein_coding |
| ENSMUSG00000110185.1 | 2185.219 | 0.180 | 0.046 | 3.959 | 7.54E-05 | 0.00126812 | Igip | protein_coding |
| ENSMUSG00000074622.4 | 896.402 | -0.224 | 0.057 | -3.957 | 7.60E-05 | 0.001276473 | Mafb | protein_coding |
| ENSMUSG00000018102.4 | 546.925 | -0.236 | 0.060 | -3.955 | 7.67E-05 | 0.001285083 | Hist1h2bc | protein_coding |
| ENSMUSG00000041378.1 | 1295.476 | -0.235 | 0.059 | -3.955 | 7.67E-05 | 0.001285083 | Cldn5 | protein_coding |
| ENSMUSG00000004980.16 | 23414.877 | 0.085 | 0.021 | 3.951 | 7.77E-05 | 0.001298932 | Hnrnpa2b1 | protein_coding |
| ENSMUSG00000035437.15 | 4855.177 | 0.121 | 0.031 | 3.951 | 7.77E-05 | 0.001298932 | Rabgap1 | protein_coding |
| ENSMUSG00000022139.16 | 10395.162 | 0.123 | 0.031 | 3.950 | 7.83E-05 | 0.001306781 | Mbnl2 | protein_coding |
| ENSMUSG00000021537.8 | 1582.711 | 0.179 | 0.045 | 3.947 | 7.91E-05 | 0.001319931 | Cetn3 | protein_coding |
| ENSMUSG00000074968.11 | 5203.695 | 0.187 | 0.047 | 3.946 | 7.93E-05 | 0.001319931 | Ano3 | protein_coding |
| ENSMUSG00000021453.2 | 567.834 | -0.242 | 0.061 | -3.946 | 7.93E-05 | 0.001319931 | Gadd45g | protein_coding |
| ENSMUSG00000021102.4 | 879.454 | -0.207 | 0.052 | -3.944 | 8.00E-05 | 0.001329569 | Glrx5 | protein_coding |
| ENSMUSG00000047213.14 | 3844.728 | 0.143 | 0.036 | 3.941 | 8.10E-05 | 0.001340933 | Ythdf3 | protein_coding |
| ENSMUSG00000058835.14 | 5010.307 | 0.124 | 0.032 | 3.941 | 8.10E-05 | 0.001340933 | Abi1 | protein_coding |
| ENSMUSG00000036890.13 | 1895.182 | 0.167 | 0.042 | 3.941 | 8.11E-05 | 0.001340933 | Gtdc1 | protein_coding |
| ENSMUSG00000042156.15 | 5300.384 | -0.124 | 0.031 | -3.940 | 8.14E-05 | 0.001344518 | Dzip1 | protein_coding |
| ENSMUSG00000019966.18 | 1957.055 | 0.186 | 0.047 | 3.939 | 8.20E-05 | 0.001352747 | Kitl | protein_coding |
| ENSMUSG00000067851.11 | 6551.314 | 0.120 | 0.030 | 3.938 | 8.21E-05 | 0.001353206 | Arfgef1 | protein_coding |
| ENSMUSG00000051285.17 | 4803.962 | 0.132 | 0.033 | 3.937 | 8.26E-05 | 0.001360409 | Pcmtd1 | protein_coding |
| ENSMUSG00000066613.14 | 1989.437 | 0.167 | 0.042 | 3.935 | 8.32E-05 | 0.001368095 | Zfp932 | protein_coding |
| ENSMUSG00000021910.15 | 30861.392 | -0.103 | 0.026 | -3.931 | 8.47E-05 | 0.001389478 | Nisch | protein_coding |
| ENSMUSG00000024797.12 | 1974.195 | -0.165 | 0.042 | -3.930 | 8.48E-05 | 0.001390183 | Vps51 | protein_coding |
| ENSMUSG00000037270.18 | 11067.487 | 0.102 | 0.026 | 3.926 | 8.65E-05 | 0.001415958 | 4932438A13Rik | protein_coding |
| ENSMUSG00000029403.14 | 2254.115 | 0.148 | 0.038 | 3.925 | 8.68E-05 | 0.001417354 | Cdkl2 | protein_coding |
| ENSMUSG00000017707.9 | 12835.124 | 0.104 | 0.027 | 3.925 | 8.69E-05 | 0.001417354 | Serinc3 | protein_coding |
| ENSMUSG00000009013.5 | 8345.902 | -0.121 | 0.031 | -3.925 | 8.69E-05 | 0.001417354 | Dynll1 | protein_coding |
| ENSMUSG00000020063.16 | 997.776 | 0.188 | 0.048 | 3.922 | 8.78E-05 | 0.001430229 | Sirt1 | protein_coding |
| ENSMUSG00000025037.6 | 1626.704 | 0.158 | 0.040 | 3.918 | 8.91E-05 | 0.00145044 | Maoa | protein_coding |
| ENSMUSG00000022568.16 | 3684.525 | -0.155 | 0.039 | -3.918 | 8.95E-05 | 0.001453865 | Scrib | protein_coding |
| ENSMUSG00000040268.17 | 5026.734 | 0.130 | 0.033 | 3.917 | 8.95E-05 | 0.001453865 | Plekha1 | protein_coding |
| ENSMUSG00000022799.4 | 2750.728 | -0.143 | 0.037 | -3.909 | 9.25E-05 | 0.001500457 | Arhgap31 | protein_coding |
| ENSMUSG00000037098.17 | 7912.251 | -0.114 | 0.029 | -3.909 | 9.27E-05 | 0.001502288 | Rab11fip3 | protein_coding |
| ENSMUSG00000034647.14 | 4780.631 | 0.121 | 0.031 | 3.907 | 9.34E-05 | 0.001510971 | Ankrd12 | protein_coding |

| | | | | | | | | |
|---|---|---|---|---|---|---|---|---|
| ENSMUSG00000027965.15 | 1801.733 | 0.186 | 0.048 | 3.907 | 9.35E-05 | 0.001510971 | Olfm3 | protein_coding |
| ENSMUSG00000061650.6 | 1645.251 | -0.169 | 0.043 | -3.906 | 9.40E-05 | 0.001517923 | Med9 | protein_coding |
| ENSMUSG00000029838.11 | 9071.774 | 0.120 | 0.031 | 3.905 | 9.42E-05 | 0.001518331 | Ptn | protein_coding |
| ENSMUSG00000029088.16 | 2677.045 | 0.154 | 0.039 | 3.902 | 9.54E-05 | 0.001535975 | Kcnip4 | protein_coding |
| ENSMUSG00000038271.17 | 3124.942 | -0.143 | 0.037 | -3.900 | 9.63E-05 | 0.001549383 | Iffo1 | protein_coding |
| ENSMUSG00000054452.10 | 13443.879 | -0.178 | 0.046 | -3.897 | 9.74E-05 | 0.001565556 | Aes | protein_coding |
| ENSMUSG00000028879.4 | 6683.620 | 0.106 | 0.027 | 3.896 | 9.80E-05 | 0.001572844 | Stx12 | protein_coding |
| ENSMUSG00000015468.14 | 489.075 | -0.227 | 0.058 | -3.893 | 9.90E-05 | 0.001586131 | Notch4 | protein_coding |
| ENSMUSG00000031955.10 | 2641.582 | -0.151 | 0.039 | -3.891 | 9.98E-05 | 0.001596701 | Bcar1 | protein_coding |
| ENSMUSG00000037221.13 | 524.173 | -0.253 | 0.065 | -3.889 | 0.00010049 | 0.001605786 | Mospd3 | protein_coding |
| ENSMUSG00000021147.17 | 3716.970 | 0.129 | 0.033 | 3.888 | 0.000101159 | 0.001614649 | Wdr37 | protein_coding |
| ENSMUSG00000028252.20 | 1594.252 | 0.159 | 0.041 | 3.887 | 0.000101348 | 0.00161583 | Ccnc | protein_coding |
| ENSMUSG00000038502.16 | 6790.840 | -0.172 | 0.044 | -3.887 | 0.000101578 | 0.001617674 | Ptov1 | protein_coding |
| ENSMUSG00000072949.6 | 326.881 | -0.289 | 0.074 | -3.885 | 0.000102364 | 0.001628349 | Acot1 | protein_coding |
| ENSMUSG00000035478.14 | 4084.279 | -0.122 | 0.031 | -3.884 | 0.000102832 | 0.001633947 | Mbd3 | protein_coding |
| ENSMUSG00000033014.12 | 3997.628 | 0.123 | 0.032 | 3.881 | 0.000104103 | 0.001652283 | Trim33 | protein_coding |
| ENSMUSG00000037826.5 | 4023.394 | 0.179 | 0.046 | 3.876 | 0.000105987 | 0.001679534 | Ppm1k | protein_coding |
| ENSMUSG00000060988.12 | 1689.871 | 0.196 | 0.051 | 3.876 | 0.000106058 | 0.001679534 | Galnt13 | protein_coding |
| ENSMUSG00000064329.13 | 6720.124 | 0.133 | 0.034 | 3.876 | 0.000106273 | 0.001679584 | Scn1a | protein_coding |
| ENSMUSG00000020571.12 | 6811.223 | -0.129 | 0.033 | -3.876 | 0.000106366 | 0.001679584 | Pdia6 | protein_coding |
| ENSMUSG00000026361.9 | 2575.398 | 0.146 | 0.038 | 3.875 | 0.000106419 | 0.001679584 | Cdc73 | protein_coding |
| ENSMUSG00000055707.13 | 2668.208 | -0.133 | 0.034 | -3.875 | 0.000106584 | 0.001680313 | Klhl26 | protein_coding |
| ENSMUSG00000002058.13 | 945.810 | -0.192 | 0.049 | -3.875 | 0.000106797 | 0.001681779 | Unc119 | protein_coding |
| ENSMUSG00000022483.16 | 131.226 | -0.300 | 0.077 | -3.874 | 0.00010715 | 0.001685463 | Col2a1 | protein_coding |
| ENSMUSG00000038679.16 | 1150.411 | 0.181 | 0.047 | 3.873 | 0.000107459 | 0.001688429 | Trps1 | protein_coding |
| ENSMUSG00000020594.14 | 10067.640 | 0.110 | 0.028 | 3.872 | 0.000107785 | 0.001691669 | Pum2 | protein_coding |
| ENSMUSG00000027162.7 | 3312.108 | 0.148 | 0.038 | 3.871 | 0.000108372 | 0.0016971 | Lin7c | protein_coding |
| ENSMUSG00000030016.14 | 6253.297 | 0.127 | 0.033 | 3.868 | 0.000109528 | 0.001713304 | Zfp638 | protein_coding |
| ENSMUSG00000020827.18 | 19435.072 | -0.095 | 0.024 | -3.867 | 0.000110159 | 0.001721259 | Mink1 | protein_coding |
| ENSMUSG00000084845.9 | 1727.111 | -0.182 | 0.047 | -3.867 | 0.000110322 | 0.001721905 | Tmem240 | protein_coding |
| ENSMUSG00000023991.16 | 1764.699 | -0.174 | 0.045 | -3.866 | 0.00011073 | 0.001724608 | Foxp4 | protein_coding |
| ENSMUSG00000044134.9 | 727.135 | -0.214 | 0.055 | -3.866 | 0.00011074 | 0.001724608 | Fam109a | protein_coding |
| ENSMUSG00000059518.14 | 1311.902 | -0.162 | 0.042 | -3.865 | 0.000111207 | 0.001728093 | Znhit1 | protein_coding |

| | | | | | | | | |
|---|---|---|---|---|---|---|---|---|
| ENSMUSG00000023067.13 | 1378.022 | -0.208 | 0.054 | -3.865 | 0.000111209 | 0.001728093 | Cdkn1a | protein_coding |
| ENSMUSG00000066551.12 | 3091.449 | 0.129 | 0.033 | 3.863 | 0.000112057 | 0.001739349 | Hmgb1 | protein_coding |
| ENSMUSG00000041459.15 | 11620.754 | 0.104 | 0.027 | 3.859 | 0.000113968 | 0.001767069 | Tardbp | protein_coding |
| ENSMUSG00000079442.12 | 1858.365 | -0.145 | 0.038 | -3.857 | 0.000114793 | 0.001777903 | St6galnac4 | protein_coding |
| ENSMUSG00000025862.14 | 3380.772 | 0.154 | 0.040 | 3.853 | 0.000116727 | 0.00180588 | Stag2 | protein_coding |
| ENSMUSG00000062931.15 | 909.360 | 0.181 | 0.047 | 3.848 | 0.000119181 | 0.001841815 | Zfp938 | protein_coding |
| ENSMUSG00000034165.16 | 1161.221 | -0.209 | 0.054 | -3.847 | 0.000119623 | 0.001846618 | Ccnd3 | protein_coding |
| ENSMUSG00000061024.8 | 787.701 | -0.201 | 0.052 | -3.846 | 0.000120278 | 0.001854705 | Rrs1 | protein_coding |
| ENSMUSG00000041638.18 | 5761.890 | -0.116 | 0.030 | -3.843 | 0.000121428 | 0.001870387 | Gcn1l1 | protein_coding |
| ENSMUSG00000039137.18 | 2368.206 | -0.219 | 0.057 | -3.841 | 0.000122754 | 0.001888749 | Whrn | protein_coding |
| ENSMUSG00000019796.12 | 8042.054 | -0.104 | 0.027 | -3.840 | 0.000123087 | 0.001891813 | Lrp11 | protein_coding |
| ENSMUSG00000051146.2 | 1924.225 | -0.192 | 0.050 | -3.840 | 0.000123239 | 0.001892089 | Camk2n2 | protein_coding |
| ENSMUSG00000035632.16 | 2933.993 | -0.132 | 0.034 | -3.838 | 0.000123822 | 0.001898956 | Cnot3 | protein_coding |
| ENSMUSG00000040229.11 | 419.140 | 0.253 | 0.066 | 3.836 | 0.000124951 | 0.001912898 | Gpr34 | protein_coding |
| ENSMUSG00000028919.11 | 1137.518 | -0.204 | 0.053 | -3.836 | 0.000125002 | 0.001912898 | Arhgef19 | protein_coding |
| ENSMUSG00000046836.13 | 3236.576 | 0.125 | 0.033 | 3.836 | 0.000125207 | 0.001913962 | Brox | protein_coding |
| ENSMUSG00000005871.14 | 23902.687 | 0.117 | 0.031 | 3.831 | 0.000127486 | 0.001946679 | Apc | protein_coding |
| ENSMUSG00000007850.16 | 14203.591 | 0.105 | 0.027 | 3.829 | 0.000128691 | 0.001962953 | Hnrnph1 | protein_coding |
| ENSMUSG00000020627.10 | 3732.272 | -0.145 | 0.038 | -3.825 | 0.000130783 | 0.001992705 | Klhl29 | protein_coding |
| ENSMUSG00000047879.16 | 6869.032 | 0.139 | 0.036 | 3.821 | 0.000132897 | 0.002022735 | Usp14 | protein_coding |
| ENSMUSG00000070520.4 | 908.389 | -0.189 | 0.050 | -3.820 | 0.000133648 | 0.002031972 | Nsmce3 | protein_coding |
| ENSMUSG00000080316.10 | 2956.360 | 0.126 | 0.033 | 3.819 | 0.000133831 | 0.002032557 | Spaca6 | protein_coding |
| ENSMUSG00000039661.14 | 4133.984 | -0.149 | 0.039 | -3.817 | 0.000134835 | 0.002045603 | Dusp26 | protein_coding |
| ENSMUSG00000020130.9 | 1633.491 | 0.151 | 0.039 | 3.816 | 0.000135618 | 0.002053205 | Tbc1d15 | protein_coding |
| ENSMUSG00000030495.12 | 1675.080 | -0.179 | 0.047 | -3.816 | 0.000135627 | 0.002053205 | Slc7a10 | protein_coding |
| ENSMUSG00000025626.16 | 1581.392 | 0.151 | 0.040 | 3.812 | 0.000137886 | 0.002085161 | Phf6 | protein_coding |
| ENSMUSG00000035596.14 | 10480.264 | -0.131 | 0.034 | -3.811 | 0.000138614 | 0.002093923 | Mboat7 | protein_coding |
| ENSMUSG00000017740.17 | 35519.639 | -0.128 | 0.034 | -3.810 | 0.000139153 | 0.002099809 | Slc12a5 | protein_coding |
| ENSMUSG00000030729.17 | 23544.131 | 0.112 | 0.029 | 3.809 | 0.000139501 | 0.00210261 | Pgm2l1 | protein_coding |
| ENSMUSG00000040124.5 | 439.603 | 0.232 | 0.061 | 3.809 | 0.000139637 | 0.00210261 | Gorab | protein_coding |
| ENSMUSG00000025020.11 | 7466.519 | -0.150 | 0.039 | -3.808 | 0.000140089 | 0.00210717 | Slit1 | protein_coding |
| ENSMUSG00000074922.1 | 419.314 | -0.264 | 0.069 | -3.807 | 0.000140464 | 0.002110549 | Fam122a | protein_coding |
| ENSMUSG00000034227.7 | 473.187 | -0.287 | 0.075 | -3.805 | 0.000141739 | 0.00212745 | Foxj1 | protein_coding |

| | | | | | | | | |
|---|---|---|---|---|---|---|---|---|
| ENSMUSG00000034610.14 | 2173.374 | 0.145 | 0.038 | 3.801 | 0.000143915 | 0.002155897 | Zcchc11 | protein_coding |
| ENSMUSG00000021054.6 | 1725.853 | 0.165 | 0.043 | 3.801 | 0.000143941 | 0.002155897 | Sgpp1 | protein_coding |
| ENSMUSG00000046785.8 | 10590.732 | 0.106 | 0.028 | 3.800 | 0.000144417 | 0.002160742 | Epm2aip1 | protein_coding |
| ENSMUSG00000069049.11 | 2658.010 | 0.196 | 0.052 | 3.799 | 0.0001451 | 0.00216865 | Eif2s3y | protein_coding |
| ENSMUSG00000023175.15 | 13864.858 | -0.190 | 0.050 | -3.797 | 0.000146454 | 0.002186568 | Bsg | protein_coding |
| ENSMUSG00000027339.15 | 3532.234 | -0.168 | 0.044 | -3.794 | 0.000148241 | 0.002210907 | Rassf2 | protein_coding |
| ENSMUSG00000037808.13 | 1045.376 | 0.200 | 0.053 | 3.788 | 0.000152089 | 0.002265902 | Fam76b | protein_coding |
| ENSMUSG00000052726.15 | 1728.732 | 0.161 | 0.043 | 3.786 | 0.000152928 | 0.002275993 | Kcnt2 | protein_coding |
| ENSMUSG00000025894.11 | 1329.652 | 0.157 | 0.041 | 3.783 | 0.00015488 | 0.002302619 | Aasdhppt | protein_coding |
| ENSMUSG00000040797.16 | 12574.165 | -0.136 | 0.036 | -3.781 | 0.000156071 | 0.00231787 | Iqsec3 | protein_coding |
| ENSMUSG00000024661.6 | 47673.458 | -0.259 | 0.069 | -3.781 | 0.000156483 | 0.002321543 | Fth1 | protein_coding |
| ENSMUSG00000070803.6 | 159.831 | -0.294 | 0.078 | -3.780 | 0.000156995 | 0.002326691 | Cited4 | protein_coding |
| ENSMUSG00000058355.8 | 4905.180 | 0.110 | 0.029 | 3.779 | 0.0001572 | 0.002327294 | Abce1 | protein_coding |
| ENSMUSG00000051579.10 | 1296.991 | 0.170 | 0.045 | 3.776 | 0.000159489 | 0.002358699 | Tceal8 | protein_coding |
| ENSMUSG00000028719.8 | 3093.061 | 0.126 | 0.033 | 3.773 | 0.00016139 | 0.002384233 | Cmpk1 | protein_coding |
| ENSMUSG00000032637.15 | 11366.564 | -0.120 | 0.032 | -3.772 | 0.000161712 | 0.002384233 | Atxn2l | protein_coding |
| ENSMUSG00000050222.10 | 298.849 | -0.272 | 0.072 | -3.772 | 0.000161813 | 0.002384233 | Il17d | protein_coding |
| ENSMUSG00000033307.7 | 3077.700 | -0.159 | 0.042 | -3.772 | 0.000162062 | 0.002384233 | Mif | protein_coding |
| ENSMUSG00000059474.13 | 1622.313 | 0.162 | 0.043 | 3.770 | 0.000163051 | 0.002396291 | Mbtd1 | protein_coding |
| ENSMUSG00000036529.17 | 13115.271 | -0.095 | 0.025 | -3.769 | 0.000164016 | 0.00240796 | Sbf1 | protein_coding |
| ENSMUSG00000011831.16 | 4898.911 | 0.135 | 0.036 | 3.768 | 0.000164787 | 0.002416769 | Evi5 | protein_coding |
| ENSMUSG00000006699.17 | 12147.616 | 0.116 | 0.031 | 3.766 | 0.000166057 | 0.002432849 | Cdc42 | protein_coding |
| ENSMUSG00000040010.10 | 3577.071 | -0.130 | 0.034 | -3.762 | 0.000168233 | 0.002462174 | Slc7a5 | protein_coding |
| ENSMUSG00000063704.12 | 199.479 | -0.291 | 0.077 | -3.762 | 0.000168586 | 0.002464777 | Mapk15 | protein_coding |
| ENSMUSG00000027466.15 | 5162.882 | -0.126 | 0.033 | -3.760 | 0.000170022 | 0.002482566 | Rbck1 | protein_coding |
| ENSMUSG00000033964.12 | 1408.394 | 0.160 | 0.042 | 3.759 | 0.000170299 | 0.002482566 | Zbtb41 | protein_coding |
| ENSMUSG00000054477.15 | 2111.492 | -0.161 | 0.043 | -3.759 | 0.000170425 | 0.002482566 | Kcnn2 | protein_coding |
| ENSMUSG00000064367.1 | 151372.231 | 0.084 | 0.022 | 3.759 | 0.000170507 | 0.002482566 | mt-Nd5 | protein_coding |
| ENSMUSG00000030276.19 | 1201.185 | -0.203 | 0.054 | -3.759 | 0.000170793 | 0.002484172 | Ttll3 | protein_coding |
| ENSMUSG00000005625.15 | 3947.932 | -0.115 | 0.031 | -3.757 | 0.000172059 | 0.002499994 | Psmd4 | protein_coding |
| ENSMUSG00000012296.15 | 1517.257 | -0.155 | 0.041 | -3.756 | 0.000172798 | 0.002508144 | Tjap1 | protein_coding |
| ENSMUSG00000030237.14 | 2531.906 | 0.158 | 0.042 | 3.754 | 0.000173927 | 0.002521945 | Slco1a4 | protein_coding |
| ENSMUSG00000034994.10 | 54876.426 | -0.100 | 0.027 | -3.754 | 0.000174207 | 0.002523412 | Eef2 | protein_coding |

| | | | | | | | | |
|---|---|---|---|---|---|---|---|---|
| ENSMUSG00000026753.5 | 2592.882 | 0.145 | 0.039 | 3.753 | 0.000174886 | 0.002530633 | Ppp6c | protein_coding |
| ENSMUSG00000062604.11 | 12371.138 | 0.091 | 0.024 | 3.752 | 0.000175635 | 0.002538877 | Srpk2 | protein_coding |
| ENSMUSG00000028937.14 | 12117.157 | -0.112 | 0.030 | -3.749 | 0.000177496 | 0.002563139 | Acot7 | protein_coding |
| ENSMUSG00000047013.15 | 8527.247 | -0.132 | 0.035 | -3.748 | 0.000178128 | 0.00256964 | Fbxo41 | protein_coding |
| ENSMUSG00000025738.7 | 50611.909 | -0.124 | 0.033 | -3.746 | 0.00017983 | 0.002591535 | Fbxl16 | protein_coding |
| ENSMUSG00000021959.15 | 1773.887 | -0.176 | 0.047 | -3.742 | 0.00018226 | 0.002621544 | Lats2 | protein_coding |
| ENSMUSG00000037622.14 | 4442.374 | -0.143 | 0.038 | -3.742 | 0.000182375 | 0.002621544 | Wdtc1 | protein_coding |
| ENSMUSG00000063887.13 | 3216.069 | 0.132 | 0.035 | 3.742 | 0.00018247 | 0.002621544 | Nlgn1 | protein_coding |
| ENSMUSG00000045967.11 | 7787.592 | 0.096 | 0.026 | 3.741 | 0.000183111 | 0.002628072 | Gpr158 | protein_coding |
| ENSMUSG00000004460.16 | 2562.283 | -0.165 | 0.044 | -3.741 | 0.000183643 | 0.002633023 | Dnajb11 | protein_coding |
| ENSMUSG00000000804.14 | 9146.575 | 0.090 | 0.024 | 3.739 | 0.00018509 | 0.002649443 | Usp32 | protein_coding |
| ENSMUSG00000019726.11 | 5389.856 | 0.115 | 0.031 | 3.738 | 0.000185164 | 0.002649443 | Lyst | protein_coding |
| ENSMUSG00000034259.7 | 725.291 | -0.202 | 0.054 | -3.736 | 0.000186866 | 0.002671083 | Exosc4 | protein_coding |
| ENSMUSG00000041935.9 | 3876.310 | 0.130 | 0.035 | 3.735 | 0.000187907 | 0.002683245 | AW549877 | protein_coding |
| ENSMUSG00000022037.14 | 29068.723 | -0.103 | 0.028 | -3.732 | 0.000190023 | 0.002708213 | Clu | protein_coding |
| ENSMUSG00000028063.15 | 2615.100 | -0.164 | 0.044 | -3.732 | 0.000190223 | 0.002708213 | Lmna | protein_coding |
| ENSMUSG00000038664.16 | 21283.798 | 0.095 | 0.025 | 3.732 | 0.000190232 | 0.002708213 | Herc1 | protein_coding |
| ENSMUSG00000109336.1 | 5085.260 | -0.125 | 0.033 | -3.731 | 0.000190811 | 0.002711178 | Samd4b | protein_coding |
| ENSMUSG00000022203.6 | 728.461 | -0.239 | 0.064 | -3.731 | 0.000190825 | 0.002711178 | Efs | protein_coding |
| ENSMUSG00000034312.14 | 16351.984 | -0.106 | 0.029 | -3.730 | 0.000191242 | 0.002711628 | Iqsec1 | protein_coding |
| ENSMUSG00000031202.5 | 2265.154 | 0.149 | 0.040 | 3.729 | 0.000192131 | 0.002721508 | Rab39b | protein_coding |
| ENSMUSG00000045589.7 | 17877.210 | 0.138 | 0.037 | 3.729 | 0.000192569 | 0.002724453 | Frrs1l | protein_coding |
| ENSMUSG00000074212.7 | 6138.466 | 0.103 | 0.028 | 3.728 | 0.000192726 | 0.002724453 | Dnajb14 | protein_coding |
| ENSMUSG00000035597.19 | 3227.089 | 0.134 | 0.036 | 3.727 | 0.00019384 | 0.002734712 | Prpf39 | protein_coding |
| ENSMUSG00000025575.14 | 3365.043 | -0.126 | 0.034 | -3.725 | 0.000195403 | 0.002753781 | Cant1 | protein_coding |
| ENSMUSG00000031827.13 | 3256.800 | -0.177 | 0.048 | -3.725 | 0.000195582 | 0.002753781 | Cotl1 | protein_coding |
| ENSMUSG00000063358.15 | 29527.718 | 0.100 | 0.027 | 3.721 | 0.000198664 | 0.002794387 | Mapk1 | protein_coding |
| ENSMUSG00000033793.12 | 7455.181 | 0.114 | 0.031 | 3.716 | 0.00020244 | 0.002844653 | Atp6v1h | protein_coding |
| ENSMUSG00000036896.5 | 1476.221 | -0.163 | 0.044 | -3.715 | 0.000203081 | 0.002850819 | C1qc | protein_coding |
| ENSMUSG00000057715.13 | 4004.061 | 0.146 | 0.039 | 3.713 | 0.000204922 | 0.002873808 | A830018L16Rik | protein_coding |
| ENSMUSG00000068328.9 | 2853.439 | -0.151 | 0.041 | -3.710 | 0.000206946 | 0.002899304 | Aup1 | protein_coding |
| ENSMUSG00000025451.15 | 2447.654 | 0.127 | 0.034 | 3.710 | 0.000207303 | 0.002901039 | Paip1 | protein_coding |
| ENSMUSG00000021733.9 | 1627.277 | 0.149 | 0.040 | 3.710 | 0.000207482 | 0.002901039 | Slc4a7 | protein_coding |

| | | | | | | | | |
|---|---|---|---|---|---|---|---|---|
| ENSMUSG00000079523.8 | 2574.010 | -0.217 | 0.058 | -3.709 | 0.000207909 | 0.002904135 | Tmsb10 | protein_coding |
| ENSMUSG00000051790.15 | 13196.225 | -0.130 | 0.035 | -3.708 | 0.000208546 | 0.002910151 | Nlgn2 | protein_coding |
| ENSMUSG00000005893.14 | 7353.490 | 0.096 | 0.026 | 3.708 | 0.000208819 | 0.002910586 | Nr2c2 | protein_coding |
| ENSMUSG00000002718.14 | 3337.236 | 0.122 | 0.033 | 3.708 | 0.000209155 | 0.002910586 | Cse1l | protein_coding |
| ENSMUSG00000009470.16 | 4049.508 | 0.121 | 0.033 | 3.708 | 0.000209197 | 0.002910586 | Tnpo1 | protein_coding |
| ENSMUSG00000025793.15 | 4599.888 | -0.120 | 0.032 | -3.704 | 0.000212033 | 0.002947137 | Hgs | protein_coding |
| ENSMUSG00000048578.11 | 8476.310 | -0.131 | 0.035 | -3.704 | 0.000212248 | 0.002947209 | Mlec | protein_coding |
| ENSMUSG00000039936.18 | 2298.493 | -0.145 | 0.039 | -3.703 | 0.000212768 | 0.002951531 | Pik3cd | protein_coding |
| ENSMUSG00000035226.5 | 6408.633 | -0.128 | 0.035 | -3.700 | 0.000215733 | 0.002989721 | Rims4 | protein_coding |
| ENSMUSG00000026103.14 | 23210.226 | 0.091 | 0.025 | 3.699 | 0.000216606 | 0.00299886 | Gls | protein_coding |
| ENSMUSG00000031885.14 | 1076.848 | 0.173 | 0.047 | 3.698 | 0.000217192 | 0.003004031 | Cbfb | protein_coding |
| ENSMUSG00000028525.16 | 6049.386 | 0.111 | 0.030 | 3.698 | 0.000217548 | 0.003006001 | Pde4b | protein_coding |
| ENSMUSG00000050812.18 | 7687.097 | 0.093 | 0.025 | 3.697 | 0.000218238 | 0.003009628 | AI314180 | protein_coding |
| ENSMUSG00000027797.15 | 46345.698 | 0.083 | 0.022 | 3.695 | 0.00022003 | 0.003031383 | Dclk1 | protein_coding |
| ENSMUSG00000002799.6 | 3010.339 | -0.152 | 0.041 | -3.694 | 0.00022077 | 0.003036273 | Jag2 | protein_coding |
| ENSMUSG00000030583.16 | 3978.172 | -0.120 | 0.032 | -3.694 | 0.000220816 | 0.003036273 | Sipa1l3 | protein_coding |
| ENSMUSG00000032827.15 | 17732.730 | 0.108 | 0.029 | 3.690 | 0.000224321 | 0.003081468 | Ppp1r9a | protein_coding |
| ENSMUSG00000070808.4 | 1522.987 | -0.180 | 0.049 | -3.685 | 0.000229088 | 0.003143883 | Gltscr1 | protein_coding |
| ENSMUSG00000045665.4 | 1427.447 | -0.155 | 0.042 | -3.677 | 0.000235861 | 0.003232862 | Mfsd5 | protein_coding |
| ENSMUSG00000022797.15 | 4932.903 | 0.169 | 0.046 | 3.677 | 0.000236031 | 0.003232862 | Tfrc | protein_coding |
| ENSMUSG00000030431.8 | 40.243 | -0.262 | 0.071 | -3.673 | 0.000239456 | NA | Tmem238 | protein_coding |
| ENSMUSG00000027273.13 | 101399.955 | 0.074 | 0.020 | 3.671 | 0.000241331 | 0.003300402 | Snap25 | protein_coding |
| ENSMUSG00000020458.16 | 28796.564 | 0.098 | 0.027 | 3.671 | 0.00024143 | 0.003300402 | Rtn4 | protein_coding |
| ENSMUSG00000031557.16 | 1518.222 | 0.158 | 0.043 | 3.669 | 0.000243038 | 0.003319168 | Plekha2 | protein_coding |
| ENSMUSG00000001761.7 | 616.586 | -0.208 | 0.057 | -3.667 | 0.000245563 | 0.003350401 | Smo | protein_coding |
| ENSMUSG00000067199.4 | 361.897 | -0.243 | 0.066 | -3.664 | 0.000248458 | 0.003385093 | Frat1 | protein_coding |
| ENSMUSG00000023391.8 | 328.657 | -0.274 | 0.075 | -3.664 | 0.000248586 | 0.003385093 | Dlx2 | protein_coding |
| ENSMUSG00000028132.15 | 3460.879 | 0.149 | 0.041 | 3.663 | 0.000249115 | 0.003386342 | Tmem56 | protein_coding |
| ENSMUSG00000007610.15 | 1257.687 | -0.188 | 0.051 | -3.663 | 0.000249158 | 0.003386342 | Gtpbp3 | protein_coding |
| ENSMUSG00000022551.7 | 5871.997 | -0.120 | 0.033 | -3.663 | 0.000249689 | 0.003390292 | Cyc1 | protein_coding |
| ENSMUSG00000026986.14 | 901.230 | 0.193 | 0.053 | 3.662 | 0.000250369 | 0.003396246 | Hnmt | protein_coding |
| ENSMUSG00000052581.13 | 1444.430 | 0.193 | 0.053 | 3.661 | 0.00025103 | 0.003401935 | Lrrtm4 | protein_coding |
| ENSMUSG00000024234.6 | 1127.433 | 0.178 | 0.049 | 3.660 | 0.000251778 | 0.003408794 | Mtpap | protein_coding |

| | | | | | | | | |
|---|---|---|---|---|---|---|---|---|
| ENSMUSG00000005610.17 | 31177.861 | 0.083 | 0.023 | 3.660 | 0.000252341 | 0.003410107 | Eif4g2 | protein_coding |
| ENSMUSG00000046432.12 | 4685.756 | -0.110 | 0.030 | -3.660 | 0.000252359 | 0.003410107 | Bex3 | protein_coding |
| ENSMUSG00000027006.13 | 3971.255 | 0.119 | 0.032 | 3.659 | 0.000252711 | 0.003411603 | Dnajc10 | protein_coding |
| ENSMUSG00000022811.16 | 5052.471 | 0.098 | 0.027 | 3.656 | 0.000255786 | 0.003449812 | Zfp148 | protein_coding |
| ENSMUSG00000021750.15 | 20236.274 | -0.144 | 0.039 | -3.655 | 0.000256796 | 0.003460114 | Fam107a | protein_coding |
| ENSMUSG00000058756.13 | 38093.905 | -0.135 | 0.037 | -3.655 | 0.000257641 | 0.003468188 | Thra | protein_coding |
| ENSMUSG00000063273.11 | 3315.081 | 0.112 | 0.031 | 3.654 | 0.000258196 | 0.003472345 | Naa15 | protein_coding |
| ENSMUSG00000024325.8 | 1734.000 | -0.165 | 0.045 | -3.654 | 0.000258665 | 0.003472915 | Ring1 | protein_coding |
| ENSMUSG00000024425.3 | 15580.524 | -0.126 | 0.034 | -3.653 | 0.000258731 | 0.003472915 | Ndfip1 | protein_coding |
| ENSMUSG00000035773.6 | 432.181 | -0.241 | 0.066 | -3.653 | 0.000259441 | 0.003476721 | Kiss1r | protein_coding |
| ENSMUSG00000033350.7 | 6240.698 | -0.131 | 0.036 | -3.653 | 0.000259508 | 0.003476721 | Chst2 | protein_coding |
| ENSMUSG00000039585.15 | 8822.190 | 0.092 | 0.025 | 3.652 | 0.000260578 | 0.003487734 | Myo9a | protein_coding |
| ENSMUSG00000034488.14 | 5436.135 | 0.157 | 0.043 | 3.650 | 0.000261928 | 0.003501568 | Edil3 | protein_coding |
| ENSMUSG00000020015.10 | 9014.222 | 0.120 | 0.033 | 3.650 | 0.000262108 | 0.003501568 | Cdk17 | protein_coding |
| ENSMUSG00000041685.15 | 2509.995 | 0.144 | 0.039 | 3.648 | 0.000264138 | 0.003525015 | Fcho2 | protein_coding |
| ENSMUSG00000051451.6 | 2638.249 | -0.164 | 0.045 | -3.648 | 0.000264364 | 0.003525015 | Crebzf | protein_coding |
| ENSMUSG00000028552.13 | 12314.819 | 0.109 | 0.030 | 3.645 | 0.000267664 | 0.003565651 | Eps15 | protein_coding |
| ENSMUSG00000004056.15 | 4656.402 | -0.121 | 0.033 | -3.643 | 0.000269644 | 0.003588635 | Akt2 | protein_coding |
| ENSMUSG00000063646.16 | 4331.434 | -0.139 | 0.038 | -3.639 | 0.000273857 | 0.003641265 | Jakmip1 | protein_coding |
| ENSMUSG00000038738.15 | 32130.870 | -0.181 | 0.050 | -3.637 | 0.000275408 | 0.00365844 | Shank1 | protein_coding |
| ENSMUSG00000044628.4 | 3894.366 | -0.146 | 0.040 | -3.634 | 0.000278834 | 0.003698916 | Rnf208 | protein_coding |
| ENSMUSG00000040447.15 | 3755.046 | -0.113 | 0.031 | -3.634 | 0.00027898 | 0.003698916 | Spns2 | protein_coding |
| ENSMUSG00000030788.16 | 3309.213 | 0.134 | 0.037 | 3.633 | 0.000280648 | 0.003716309 | Rnf141 | protein_coding |
| ENSMUSG00000089862.8 | 1466.257 | 0.154 | 0.042 | 3.632 | 0.000280819 | 0.003716309 | Umad1 | protein_coding |
| ENSMUSG00000051674.14 | 5330.973 | 0.117 | 0.032 | 3.629 | 0.00028438 | 0.003759898 | Dcun1d4 | protein_coding |
| ENSMUSG00000000093.6 | 101.014 | -0.282 | 0.078 | -3.625 | 0.00028932 | 0.003821631 | Tbx2 | protein_coding |
| ENSMUSG00000050373.13 | 1787.290 | -0.161 | 0.044 | -3.624 | 0.0002906 | 0.003831569 | Snx21 | protein_coding |
| ENSMUSG00000046096.7 | 3451.773 | 0.131 | 0.036 | 3.623 | 0.000290866 | 0.003831569 | BC030336 | protein_coding |
| ENSMUSG00000021189.11 | 1630.956 | 0.153 | 0.042 | 3.623 | 0.000290888 | 0.003831569 | Atxn3 | protein_coding |
| ENSMUSG00000021326.12 | 1074.768 | -0.196 | 0.054 | -3.623 | 0.000291679 | 0.003838392 | Trim27 | protein_coding |
| ENSMUSG00000038301.15 | 6332.728 | 0.134 | 0.037 | 3.622 | 0.000292006 | 0.003839113 | Snx10 | protein_coding |
| ENSMUSG00000027206.13 | 3966.601 | 0.141 | 0.039 | 3.621 | 0.000293067 | 0.003849478 | Cops2 | protein_coding |
| ENSMUSG00000060882.5 | 3915.755 | 0.115 | 0.032 | 3.621 | 0.00029386 | 0.003856302 | Kcnd2 | protein_coding |

| | | | | | | | | |
|---|---|---|---|---|---|---|---|---|
| ENSMUSG00000057335.11 | 5456.041 | 0.104 | 0.029 | 3.614 | 0.000301868 | 0.0039577 | Cep170 | protein_coding |
| ENSMUSG00000025240.9 | 4441.391 | 0.120 | 0.033 | 3.613 | 0.000302432 | 0.003961411 | Sacm1l | protein_coding |
| ENSMUSG00000045608.6 | 532.576 | -0.208 | 0.058 | -3.613 | 0.000302919 | 0.00396411 | Dbx2 | protein_coding |
| ENSMUSG00000025609.15 | 5707.663 | 0.100 | 0.028 | 3.612 | 0.000303599 | 0.003969318 | Mkln1 | protein_coding |
| ENSMUSG00000021400.7 | 2840.929 | -0.162 | 0.045 | -3.608 | 0.000308153 | 0.004021399 | Wrnip1 | protein_coding |
| ENSMUSG00000029028.14 | 2985.254 | -0.115 | 0.032 | -3.608 | 0.000308812 | 0.004026263 | Lrrc47 | protein_coding |
| ENSMUSG00000014776.7 | 882.424 | -0.204 | 0.057 | -3.606 | 0.000310608 | 0.004045946 | Nol3 | protein_coding |
| ENSMUSG00000024870.5 | 5671.922 | -0.113 | 0.031 | -3.601 | 0.000317196 | 0.00412104 | Rab1b | protein_coding |
| ENSMUSG00000062300.14 | 304.768 | -0.255 | 0.071 | -3.601 | 0.000317231 | 0.00412104 | Nectin2 | protein_coding |
| ENSMUSG00000058153.15 | 16697.456 | -0.104 | 0.029 | -3.601 | 0.00031725 | 0.00412104 | Sez6l | protein_coding |
| ENSMUSG00000044098.14 | 1839.150 | 0.148 | 0.041 | 3.600 | 0.000318823 | 0.004137652 | Rsbn1 | protein_coding |
| ENSMUSG00000028780.13 | 1510.248 | 0.195 | 0.054 | 3.598 | 0.000320834 | 0.00415992 | Sema3c | protein_coding |
| ENSMUSG00000020396.8 | 2499.672 | -0.178 | 0.049 | -3.596 | 0.000323053 | 0.004181344 | Nefh | protein_coding |
| ENSMUSG00000038319.14 | 1901.623 | -0.200 | 0.055 | -3.596 | 0.00032308 | 0.004181344 | Kcnh2 | protein_coding |
| ENSMUSG00000067336.6 | 8649.416 | 0.126 | 0.035 | 3.596 | 0.000323542 | 0.004183493 | Bmpr2 | protein_coding |
| ENSMUSG00000058975.7 | 10061.760 | -0.134 | 0.037 | -3.592 | 0.000328016 | 0.004237152 | Kcnc1 | protein_coding |
| ENSMUSG00000074733.14 | 4031.357 | 0.139 | 0.039 | 3.592 | 0.000328294 | 0.004237152 | Zfp950 | protein_coding |
| ENSMUSG00000034006.16 | 856.322 | -0.176 | 0.049 | -3.591 | 0.000329594 | 0.004243894 | Pqlc1 | protein_coding |
| ENSMUSG00000029436.9 | 11660.321 | -0.124 | 0.035 | -3.591 | 0.000329679 | 0.004243894 | Mmp17 | protein_coding |
| ENSMUSG00000035863.14 | 12457.747 | -0.122 | 0.034 | -3.591 | 0.000329719 | 0.004243894 | Palm | protein_coding |
| ENSMUSG00000022314.9 | 5778.816 | 0.124 | 0.035 | 3.588 | 0.000332637 | 0.004277541 | Rad21 | protein_coding |
| ENSMUSG00000063253.11 | 4454.781 | 0.116 | 0.032 | 3.588 | 0.000333621 | 0.004286287 | Scoc | protein_coding |
| ENSMUSG00000036095.11 | 7669.177 | 0.141 | 0.039 | 3.586 | 0.000336149 | 0.004314825 | Dgkb | protein_coding |
| ENSMUSG00000029203.16 | 4698.903 | 0.111 | 0.031 | 3.585 | 0.000336463 | 0.004314937 | Ube2k | protein_coding |
| ENSMUSG00000028528.16 | 19738.325 | 0.094 | 0.026 | 3.585 | 0.000336827 | 0.004315678 | Dnajc6 | protein_coding |
| ENSMUSG00000031278.12 | 4641.231 | 0.118 | 0.033 | 3.585 | 0.000337503 | 0.004320405 | Acsl4 | protein_coding |
| ENSMUSG00000019254.16 | 6588.564 | -0.126 | 0.035 | -3.583 | 0.000340157 | 0.004350432 | Ppp1r12c | protein_coding |
| ENSMUSG00000025393.12 | 47150.991 | -0.074 | 0.021 | -3.582 | 0.000340688 | 0.004353265 | Atp5b | protein_coding |
| ENSMUSG00000026241.5 | 142.068 | -0.277 | 0.077 | -3.582 | 0.000341395 | 0.004358355 | Nppc | protein_coding |
| ENSMUSG00000044562.12 | 339.366 | -0.260 | 0.073 | -3.577 | 0.000347001 | 0.004425907 | Rasip1 | protein_coding |
| ENSMUSG00000058793.14 | 36320.828 | -0.074 | 0.021 | -3.576 | 0.000349139 | 0.004449158 | Cds2 | protein_coding |
| ENSMUSG00000027634.14 | 23822.698 | 0.068 | 0.019 | 3.575 | 0.000350648 | 0.004464351 | Ndrg3 | protein_coding |
| ENSMUSG00000022151.16 | 1726.599 | 0.146 | 0.041 | 3.574 | 0.00035154 | 0.004471664 | Ttc33 | protein_coding |

| | | | | | | | | |
|---|---|---|---|---|---|---|---|---|
| ENSMUSG00000027582.16 | 3736.896 | -0.127 | 0.036 | -3.573 | 0.000353062 | 0.004486525 | Zgpat | protein_coding |
| ENSMUSG00000024164.15 | 77.285 | -0.240 | 0.067 | -3.573 | 0.000353345 | 0.004486525 | C3 | protein_coding |
| ENSMUSG00000041483.14 | 2332.896 | 0.123 | 0.034 | 3.570 | 0.000357408 | 0.004530714 | Zfp281 | protein_coding |
| ENSMUSG00000059895.12 | 2602.634 | -0.143 | 0.040 | -3.570 | 0.000357468 | 0.004530714 | Ptp4a3 | protein_coding |
| ENSMUSG00000023990.18 | 383.046 | -0.239 | 0.067 | -3.569 | 0.000358127 | 0.004531529 | Tfeb | protein_coding |
| ENSMUSG00000044501.17 | 524.901 | 0.221 | 0.062 | 3.569 | 0.000358207 | 0.004531529 | Zfp758 | protein_coding |
| ENSMUSG00000035969.15 | 8931.934 | -0.109 | 0.031 | -3.567 | 0.000361347 | 0.004560332 | Rusc2 | protein_coding |
| ENSMUSG00000031748.16 | 41778.564 | -0.072 | 0.020 | -3.567 | 0.000361423 | 0.004560332 | Gnao1 | protein_coding |
| ENSMUSG00000027438.14 | 29828.806 | 0.096 | 0.027 | 3.566 | 0.000361939 | 0.004562768 | Napb | protein_coding |
| ENSMUSG00000028804.20 | 7522.495 | -0.128 | 0.036 | -3.565 | 0.000363633 | 0.004575926 | Csmd2 | protein_coding |
| ENSMUSG00000022553.15 | 2649.414 | -0.122 | 0.034 | -3.564 | 0.000364974 | 0.004588711 | Maf1 | protein_coding |
| ENSMUSG00000030824.17 | 3914.507 | -0.114 | 0.032 | -3.564 | 0.000365921 | 0.004592676 | Nucb1 | protein_coding |
| ENSMUSG00000007594.10 | 4333.406 | -0.170 | 0.048 | -3.564 | 0.000365941 | 0.004592676 | Hapln4 | protein_coding |
| ENSMUSG00000019943.10 | 34531.921 | 0.096 | 0.027 | 3.562 | 0.000367555 | 0.004608826 | Atp2b1 | protein_coding |
| ENSMUSG00000030180.15 | 2883.416 | 0.121 | 0.034 | 3.561 | 0.000369247 | 0.004625922 | Kdm5a | protein_coding |
| ENSMUSG00000031214.13 | 3052.052 | 0.111 | 0.031 | 3.560 | 0.000370293 | 0.004634912 | Ophn1 | protein_coding |
| ENSMUSG00000032745.17 | 4681.432 | 0.112 | 0.031 | 3.556 | 0.000376967 | 0.004714265 | Gpbp1 | protein_coding |
| ENSMUSG00000057342.15 | 1642.843 | -0.178 | 0.050 | -3.554 | 0.000379001 | 0.004735502 | Sphk2 | protein_coding |
| ENSMUSG00000021209.12 | 1926.483 | 0.150 | 0.042 | 3.552 | 0.000381708 | 0.004765092 | Ppp4r4 | protein_coding |
| ENSMUSG00000061904.12 | 16465.016 | -0.095 | 0.027 | -3.552 | 0.000382882 | 0.004775515 | Slc25a3 | protein_coding |
| ENSMUSG00000020745.15 | 28358.370 | 0.101 | 0.028 | 3.550 | 0.000385488 | 0.004803771 | Pafah1b1 | protein_coding |
| ENSMUSG00000035798.14 | 7142.290 | 0.098 | 0.028 | 3.548 | 0.000388145 | 0.004832611 | Zdhhc17 | protein_coding |
| ENSMUSG00000031592.10 | 5050.383 | 0.130 | 0.037 | 3.547 | 0.000389063 | 0.004839766 | Pcm1 | protein_coding |
| ENSMUSG00000048379.8 | 910.172 | 0.173 | 0.049 | 3.546 | 0.000391185 | 0.004861865 | Socs4 | protein_coding |
| ENSMUSG00000029212.11 | 9798.783 | 0.083 | 0.024 | 3.541 | 0.000397895 | 0.004940395 | Gabrb1 | protein_coding |
| ENSMUSG00000021113.5 | 2954.815 | 0.127 | 0.036 | 3.541 | 0.000398204 | 0.004940395 | Snapc1 | protein_coding |
| ENSMUSG00000043460.6 | 7992.118 | -0.144 | 0.041 | -3.540 | 0.000399449 | 0.004951478 | Elfn2 | protein_coding |
| ENSMUSG00000063802.5 | 3194.451 | -0.147 | 0.042 | -3.540 | 0.000400096 | 0.004955138 | Hspbp1 | protein_coding |
| ENSMUSG00000073664.11 | 1934.775 | 0.140 | 0.039 | 3.540 | 0.00040049 | 0.004955663 | Nbeal1 | protein_coding |
| ENSMUSG00000004054.8 | 1271.192 | -0.156 | 0.044 | -3.536 | 0.000406173 | 0.005014959 | Map3k11 | protein_coding |
| ENSMUSG00000040488.17 | 4583.160 | -0.156 | 0.044 | -3.536 | 0.000406349 | 0.005014959 | Ltbp4 | protein_coding |
| ENSMUSG00000034839.5 | 1478.734 | -0.142 | 0.040 | -3.532 | 0.000412808 | 0.005090217 | Larp6 | protein_coding |
| ENSMUSG00000033423.16 | 5161.290 | -0.094 | 0.027 | -3.531 | 0.000413301 | 0.005091843 | Eri3 | protein_coding |

| | | | | | | | | |
|---|---|---|---|---|---|---|---|---|
| ENSMUSG00000041058.15 | 2095.836 | 0.140 | 0.040 | 3.531 | 0.000414531 | 0.005102527 | Wwp1 | protein_coding |
| ENSMUSG00000075232.5 | 3271.674 | 0.124 | 0.035 | 3.528 | 0.00041833 | 0.005144796 | Amd1 | protein_coding |
| ENSMUSG00000026864.13 | 19795.951 | -0.133 | 0.038 | -3.527 | 0.00042058 | 0.005167966 | Hspa5 | protein_coding |
| ENSMUSG00000054000.5 | 94.976 | -0.273 | 0.078 | -3.525 | 0.000423192 | 0.005195526 | Tusc1 | protein_coding |
| ENSMUSG00000074211.4 | 297.930 | -0.236 | 0.067 | -3.523 | 0.000427122 | 0.005237018 | Sdhaf1 | protein_coding |
| ENSMUSG00000050288.6 | 411.682 | -0.255 | 0.072 | -3.523 | 0.000427315 | 0.005237018 | Fzd2 | protein_coding |
| ENSMUSG00000037710.8 | 2405.474 | -0.128 | 0.036 | -3.522 | 0.000428769 | 0.005248463 | Cisd1 | protein_coding |
| ENSMUSG00000019878.8 | 1464.577 | 0.151 | 0.043 | 3.521 | 0.000429366 | 0.005248463 | Hsf2 | protein_coding |
| ENSMUSG00000016995.17 | 702.461 | -0.192 | 0.054 | -3.520 | 0.000430842 | 0.00526016 | Matn4 | protein_coding |
| ENSMUSG00000068748.7 | 12731.922 | 0.118 | 0.033 | 3.520 | 0.000431069 | 0.00526016 | Ptprz1 | protein_coding |
| ENSMUSG00000030086.16 | 1933.121 | -0.136 | 0.039 | -3.518 | 0.000434824 | 0.005301389 | Chchd6 | protein_coding |
| ENSMUSG00000007050.17 | 216.569 | -0.257 | 0.073 | -3.517 | 0.000435919 | 0.00531015 | Lsm2 | protein_coding |
| ENSMUSG00000028971.4 | 255.446 | -0.260 | 0.074 | -3.516 | 0.000437781 | 0.005327042 | Cort | protein_coding |
| ENSMUSG00000026820.5 | 1659.747 | -0.149 | 0.042 | -3.516 | 0.000438062 | 0.005327042 | Ptges2 | protein_coding |
| ENSMUSG00000017485.10 | 5475.424 | 0.124 | 0.035 | 3.515 | 0.00043918 | 0.005334737 | Top2b | protein_coding |
| ENSMUSG00000026791.14 | 1225.822 | -0.166 | 0.047 | -3.515 | 0.000439796 | 0.005334737 | Slc2a8 | protein_coding |
| ENSMUSG00000024392.17 | 11694.491 | -0.106 | 0.030 | -3.515 | 0.00043983 | 0.005334737 | Bag6 | protein_coding |
| ENSMUSG00000004264.17 | 2681.058 | -0.127 | 0.036 | -3.514 | 0.000442164 | 0.005354826 | Phb2 | protein_coding |
| ENSMUSG00000020444.19 | 2960.142 | -0.127 | 0.036 | -3.513 | 0.000442246 | 0.005354826 | Guk1 | protein_coding |
| ENSMUSG00000070802.5 | 22025.654 | -0.118 | 0.033 | -3.513 | 0.000442679 | 0.005355469 | Pnmal2 | protein_coding |
| ENSMUSG00000024949.17 | 8737.071 | -0.085 | 0.024 | -3.509 | 0.000450024 | 0.005439652 | Sf1 | protein_coding |
| ENSMUSG00000037326.10 | 4290.107 | -0.131 | 0.037 | -3.508 | 0.000450669 | 0.005442781 | Capn15 | protein_coding |
| ENSMUSG00000033900.13 | 6789.646 | 0.113 | 0.032 | 3.507 | 0.000453308 | 0.005469965 | Map9 | protein_coding |
| ENSMUSG00000046593.3 | 97.146 | 0.268 | 0.076 | 3.507 | 0.000453735 | 0.005470433 | Tmem215 | protein_coding |
| ENSMUSG00000022837.13 | 1432.661 | 0.179 | 0.051 | 3.506 | 0.000454607 | 0.00547626 | Iqcb1 | protein_coding |
| ENSMUSG00000037999.13 | 5931.370 | 0.127 | 0.036 | 3.503 | 0.000459898 | 0.005535273 | Arap2 | protein_coding |
| ENSMUSG00000031860.17 | 479.662 | -0.210 | 0.060 | -3.502 | 0.000462129 | 0.005557383 | Pbx4 | protein_coding |
| ENSMUSG00000024294.13 | 9211.754 | 0.106 | 0.030 | 3.501 | 0.0004642 | 0.005573328 | Mib1 | protein_coding |
| ENSMUSG00000040446.3 | 3672.494 | 0.107 | 0.030 | 3.501 | 0.000464246 | 0.005573328 | Rprd1a | protein_coding |
| ENSMUSG00000025940.6 | 1464.413 | -0.150 | 0.043 | -3.500 | 0.000465929 | 0.005588766 | Tmem70 | protein_coding |
| ENSMUSG00000021259.5 | 10067.359 | -0.137 | 0.039 | -3.498 | 0.000468441 | 0.00561412 | Cyp46a1 | protein_coding |
| ENSMUSG00000032187.16 | 9331.978 | -0.099 | 0.028 | -3.496 | 0.000472183 | 0.00565253 | Smarca4 | protein_coding |
| ENSMUSG00000036499.9 | 3153.080 | 0.143 | 0.041 | 3.496 | 0.000472448 | 0.00565253 | Eea1 | protein_coding |

| | | | | | | | | |
|---|---|---|---|---|---|---|---|---|
| ENSMUSG00000008348.9 | 1368.538 | -0.158 | 0.045 | -3.495 | 0.000473279 | 0.005653172 | Ubc | protein_coding |
| ENSMUSG00000058240.13 | 3214.449 | 0.115 | 0.033 | 3.495 | 0.000473304 | 0.005653172 | Cryzl1 | protein_coding |
| ENSMUSG00000026781.15 | 3085.593 | 0.135 | 0.039 | 3.495 | 0.000473706 | 0.005653185 | Acbd5 | protein_coding |
| ENSMUSG00000024335.19 | 8629.357 | -0.084 | 0.024 | -3.495 | 0.000474233 | 0.005654691 | Brd2 | protein_coding |
| ENSMUSG00000026434.12 | 5108.303 | 0.153 | 0.044 | 3.493 | 0.00047703 | 0.005678428 | Nucks1 | protein_coding |
| ENSMUSG00000042216.13 | 10116.678 | -0.133 | 0.038 | -3.491 | 0.000482099 | 0.005730376 | Sgsm1 | protein_coding |
| ENSMUSG00000021096.11 | 11674.416 | 0.087 | 0.025 | 3.490 | 0.000482207 | 0.005730376 | Ppm1a | protein_coding |
| ENSMUSG00000000223.13 | 5815.648 | 0.110 | 0.032 | 3.490 | 0.000482811 | 0.005732726 | Drp2 | protein_coding |
| ENSMUSG00000021773.10 | 432.534 | -0.210 | 0.060 | -3.488 | 0.000485758 | 0.005762861 | Comtd1 | protein_coding |
| ENSMUSG00000026889.12 | 2731.056 | 0.123 | 0.035 | 3.488 | 0.000487533 | 0.005779053 | Rbm18 | protein_coding |
| ENSMUSG00000022091.5 | 1953.552 | -0.163 | 0.047 | -3.484 | 0.000493064 | 0.005839699 | Sorbs3 | protein_coding |
| ENSMUSG00000042807.15 | 2774.596 | 0.130 | 0.037 | 3.481 | 0.000499202 | 0.005907435 | Hecw2 | protein_coding |
| ENSMUSG00000018974.7 | 2701.335 | -0.116 | 0.033 | -3.481 | 0.000500257 | 0.005914952 | Sart3 | protein_coding |
| ENSMUSG00000022228.13 | 7106.188 | 0.088 | 0.025 | 3.480 | 0.000502213 | 0.005933106 | Zscan26 | protein_coding |
| ENSMUSG00000048707.9 | 814.301 | -0.217 | 0.062 | -3.479 | 0.000502953 | 0.005936865 | Tprn | protein_coding |
| ENSMUSG00000004473.10 | 191.582 | -0.263 | 0.076 | -3.479 | 0.000504198 | 0.005946578 | Clec11a | protein_coding |
| ENSMUSG00000048834.16 | 5211.346 | 0.145 | 0.042 | 3.477 | 0.000507906 | 0.005985307 | Vstm2a | protein_coding |
| ENSMUSG00000027433.5 | 2363.510 | 0.128 | 0.037 | 3.476 | 0.00050854 | 0.005987772 | Xrn2 | protein_coding |
| ENSMUSG00000034566.10 | 5223.329 | 0.108 | 0.031 | 3.476 | 0.000509333 | 0.0059921 | Atp5h | protein_coding |
| ENSMUSG00000024579.7 | 1094.501 | -0.171 | 0.049 | -3.476 | 0.000509785 | 0.005992424 | Pcyox1l | protein_coding |
| ENSMUSG00000005057.13 | 266.087 | -0.248 | 0.071 | -3.471 | 0.000517946 | 0.006083275 | Sh2b2 | protein_coding |
| ENSMUSG00000073639.5 | 4814.906 | 0.117 | 0.034 | 3.471 | 0.000519286 | 0.006093935 | Rab18 | protein_coding |
| ENSMUSG00000001911.16 | 10874.911 | -0.170 | 0.049 | -3.470 | 0.000520772 | 0.006106292 | Nfix | protein_coding |
| ENSMUSG00000034891.13 | 12073.629 | -0.146 | 0.042 | -3.468 | 0.000523451 | 0.006132605 | Sncb | protein_coding |
| ENSMUSG00000007656.13 | 12599.868 | 0.124 | 0.036 | 3.468 | 0.000524293 | 0.00613736 | Arpp19 | protein_coding |
| ENSMUSG00000031703.8 | 11346.985 | 0.085 | 0.025 | 3.467 | 0.000527212 | 0.006161351 | Itfg1 | protein_coding |
| ENSMUSG00000038628.8 | 1999.807 | 0.132 | 0.038 | 3.467 | 0.000527217 | 0.006161351 | Polr3k | protein_coding |
| ENSMUSG00000021945.8 | 5020.241 | 0.099 | 0.028 | 3.466 | 0.000527953 | 0.006164847 | Zmym2 | protein_coding |
| ENSMUSG00000019841.15 | 2901.951 | 0.139 | 0.040 | 3.463 | 0.000534624 | 0.006237578 | Rev3l | protein_coding |
| ENSMUSG00000028124.15 | 2394.901 | 0.142 | 0.041 | 3.462 | 0.000535935 | 0.006242606 | Gclm | protein_coding |
| ENSMUSG00000039126.9 | 4225.706 | 0.130 | 0.037 | 3.462 | 0.000535941 | 0.006242606 | Prune2 | protein_coding |
| ENSMUSG00000042155.3 | 3062.745 | 0.145 | 0.042 | 3.461 | 0.000538447 | 0.006266612 | Klhl23 | protein_coding |
| ENSMUSG00000049600.8 | 1115.392 | -0.191 | 0.055 | -3.460 | 0.000539749 | 0.00627659 | Zbtb45 | protein_coding |

| | | | | | | | | |
|---|---|---|---|---|---|---|---|---|
| ENSMUSG00000041763.14 | 2918.798 | 0.122 | 0.035 | 3.460 | 0.000540632 | 0.00628167 | Tpp2 | protein_coding |
| ENSMUSG00000046798.14 | 2342.337 | 0.117 | 0.034 | 3.458 | 0.000544373 | 0.006318227 | Cldn12 | protein_coding |
| ENSMUSG00000039697.16 | 4676.619 | 0.105 | 0.031 | 3.458 | 0.000544675 | 0.006318227 | Ncoa7 | protein_coding |
| ENSMUSG00000042699.11 | 7470.811 | 0.108 | 0.031 | 3.457 | 0.000545494 | 0.006318299 | Dhx9 | protein_coding |
| ENSMUSG00000024383.8 | 2222.111 | 0.135 | 0.039 | 3.457 | 0.000545578 | 0.006318299 | Map3k2 | protein_coding |
| ENSMUSG00000038127.14 | 3459.048 | 0.123 | 0.036 | 3.457 | 0.000546442 | 0.006323116 | Ccdc50 | protein_coding |
| ENSMUSG00000022619.5 | 18252.596 | -0.103 | 0.030 | -3.455 | 0.000549525 | 0.006353575 | Mapk8ip2 | protein_coding |
| ENSMUSG00000003378.9 | 12661.282 | -0.147 | 0.043 | -3.452 | 0.000555731 | 0.006413728 | Grik5 | protein_coding |
| ENSMUSG00000033184.14 | 4298.847 | 0.118 | 0.034 | 3.452 | 0.000555848 | 0.006413728 | Tmed7 | protein_coding |
| ENSMUSG00000062961.7 | 803.326 | -0.201 | 0.058 | -3.452 | 0.000556093 | 0.006413728 | Ccdc177 | protein_coding |
| ENSMUSG00000010154.7 | 881.424 | -0.175 | 0.051 | -3.451 | 0.000557525 | 0.006424984 | Spire2 | protein_coding |
| ENSMUSG00000031696.9 | 8271.302 | 0.109 | 0.031 | 3.449 | 0.000562603 | 0.006478204 | Vps35 | protein_coding |
| ENSMUSG00000039809.10 | 17433.164 | -0.121 | 0.035 | -3.447 | 0.000565829 | 0.006510033 | Gabbr2 | protein_coding |
| ENSMUSG00000073775.4 | 448.991 | -0.223 | 0.065 | -3.446 | 0.000569408 | 0.006545871 | Kti12 | protein_coding |
| ENSMUSG00000022971.18 | 466.869 | -0.220 | 0.064 | -3.445 | 0.000570219 | 0.006549854 | Ifnar2 | protein_coding |
| ENSMUSG00000002413.15 | 7727.602 | 0.107 | 0.031 | 3.445 | 0.000571053 | 0.006554084 | Braf | protein_coding |
| ENSMUSG00000028278.14 | 6029.259 | 0.095 | 0.028 | 3.445 | 0.000571742 | 0.006556655 | Rragd | protein_coding |
| ENSMUSG00000041180.13 | 1558.120 | 0.146 | 0.042 | 3.444 | 0.000572642 | 0.006561645 | Hectd2 | protein_coding |
| ENSMUSG00000020794.13 | 2123.614 | 0.126 | 0.037 | 3.444 | 0.000573884 | 0.00657053 | Ube2g1 | protein_coding |
| ENSMUSG00000075376.10 | 3462.216 | 0.147 | 0.043 | 3.442 | 0.000577777 | 0.006607137 | Rc3h2 | protein_coding |
| ENSMUSG00000025409.14 | 3334.404 | -0.157 | 0.046 | -3.442 | 0.000578215 | 0.006607137 | Mbd6 | protein_coding |
| ENSMUSG00000079184.10 | 2723.726 | 0.122 | 0.035 | 3.442 | 0.000578488 | 0.006607137 | Mphosph8 | protein_coding |
| ENSMUSG00000035278.9 | 809.064 | -0.186 | 0.054 | -3.440 | 0.0005817 | 0.006638451 | Plekhj1 | protein_coding |
| ENSMUSG00000025586.17 | 5259.590 | 0.106 | 0.031 | 3.439 | 0.000584287 | 0.006662577 | Cpeb1 | protein_coding |
| ENSMUSG00000051166.10 | 7507.555 | 0.114 | 0.033 | 3.436 | 0.000590515 | 0.006728144 | Eml5 | protein_coding |
| ENSMUSG00000026831.16 | 166.521 | -0.264 | 0.077 | -3.436 | 0.000591138 | 0.006729804 | 1700007K13Rik | protein_coding |
| ENSMUSG00000024811.11 | 8264.088 | 0.093 | 0.027 | 3.434 | 0.000594847 | 0.00676656 | Tnks2 | protein_coding |
| ENSMUSG00000079598.3 | 1700.123 | -0.137 | 0.040 | -3.430 | 0.000604521 | 0.006869955 | Clec2l | protein_coding |
| ENSMUSG00000087408.10 | 2163.305 | -0.145 | 0.042 | -3.429 | 0.000604911 | 0.006869955 | Cers1 | protein_coding |
| ENSMUSG00000024287.7 | 1646.626 | 0.162 | 0.047 | 3.428 | 0.00060865 | 0.006906849 | Thoc1 | protein_coding |
| ENSMUSG00000027342.14 | 1253.220 | -0.156 | 0.045 | -3.427 | 0.000609931 | 0.00691582 | Pcna | protein_coding |
| ENSMUSG00000028127.10 | 3518.106 | 0.124 | 0.036 | 3.426 | 0.000611973 | 0.006933399 | Abcd3 | protein_coding |
| ENSMUSG00000025870.10 | 658.683 | -0.205 | 0.060 | -3.424 | 0.000616447 | 0.006978476 | Arl10 | protein_coding |

| | | | | | | | | |
|---|---|---|---|---|---|---|---|---|
| ENSMUSG00000029815.12 | 405.744 | -0.236 | 0.069 | -3.424 | 0.000617888 | 0.006989177 | Malsu1 | protein_coding |
| ENSMUSG00000024096.16 | 4807.583 | -0.100 | 0.029 | -3.423 | 0.000618642 | 0.006992088 | Ralbp1 | protein_coding |
| ENSMUSG00000071711.11 | 886.680 | -0.165 | 0.048 | -3.423 | 0.00061973 | 0.006998773 | Mpst | protein_coding |
| ENSMUSG00000016344.14 | 1230.770 | -0.175 | 0.051 | -3.422 | 0.000622082 | 0.007019712 | Ppdpf | protein_coding |
| ENSMUSG00000058624.12 | 18026.028 | 0.120 | 0.035 | 3.421 | 0.000623741 | 0.007032802 | Gda | protein_coding |
| ENSMUSG00000038145.17 | 4788.358 | -0.113 | 0.033 | -3.420 | 0.000627349 | 0.007067828 | Snrk | protein_coding |
| ENSMUSG00000062115.15 | 6353.833 | -0.143 | 0.042 | -3.419 | 0.000629491 | 0.007086304 | Rai1 | protein_coding |
| ENSMUSG00000024347.16 | 3388.725 | -0.138 | 0.040 | -3.416 | 0.000635592 | 0.007149267 | Psd2 | protein_coding |
| ENSMUSG00000028776.14 | 367.695 | -0.255 | 0.075 | -3.414 | 0.000639852 | 0.00719116 | Tinagl1 | protein_coding |
| ENSMUSG00000043059.16 | 1024.499 | -0.165 | 0.048 | -3.414 | 0.000640337 | 0.00719116 | Zfp513 | protein_coding |
| ENSMUSG00000072941.5 | 383.878 | -0.258 | 0.076 | -3.413 | 0.00064165 | 0.007200176 | Sod3 | protein_coding |
| ENSMUSG00000025508.13 | 2725.956 | -0.139 | 0.041 | -3.412 | 0.000645058 | 0.007232173 | Rplp2 | protein_coding |
| ENSMUSG00000020366.18 | 11766.295 | 0.091 | 0.027 | 3.412 | 0.000645566 | 0.007232173 | Mapk9 | protein_coding |
| ENSMUSG00000057963.9 | 3037.746 | -0.118 | 0.035 | -3.412 | 0.000646041 | 0.007232173 | Itpk1 | protein_coding |
| ENSMUSG00000031765.8 | 3412.140 | -0.166 | 0.049 | -3.410 | 0.000649665 | 0.007258634 | Mt1 | protein_coding |
| ENSMUSG00000046556.7 | 1690.039 | -0.166 | 0.049 | -3.410 | 0.000650533 | 0.007258634 | Zfp319 | protein_coding |
| ENSMUSG00000039483.10 | 1846.790 | -0.150 | 0.044 | -3.410 | 0.000650715 | 0.007258634 | Asb6 | protein_coding |
| ENSMUSG00000033054.7 | 1514.406 | 0.164 | 0.048 | 3.409 | 0.000650854 | 0.007258634 | Npat | protein_coding |
| ENSMUSG00000046994.9 | 1499.290 | 0.145 | 0.043 | 3.408 | 0.000653986 | 0.007286383 | Mars2 | protein_coding |
| ENSMUSG00000029629.17 | 3178.344 | 0.125 | 0.037 | 3.408 | 0.000654733 | 0.007288945 | Phf14 | protein_coding |
| ENSMUSG00000038888.8 | 404.871 | -0.220 | 0.065 | -3.408 | 0.000655288 | 0.007289362 | Ctu1 | protein_coding |
| ENSMUSG00000026796.16 | 2345.033 | -0.144 | 0.042 | -3.407 | 0.000656009 | 0.007291629 | Fam129b | protein_coding |
| ENSMUSG00000040721.9 | 5939.219 | -0.125 | 0.037 | -3.407 | 0.00065706 | 0.007297565 | Zfhx2 | protein_coding |
| ENSMUSG00000040537.17 | 13537.484 | 0.076 | 0.022 | 3.406 | 0.000659065 | 0.007314064 | Adam22 | protein_coding |
| ENSMUSG00000074738.2 | 888.257 | -0.181 | 0.053 | -3.406 | 0.000660047 | 0.007319198 | Fndc10 | protein_coding |
| ENSMUSG00000028883.17 | 945.381 | 0.170 | 0.050 | 3.405 | 0.000662172 | 0.007336996 | Sema3a | protein_coding |
| ENSMUSG00000049482.16 | 854.706 | -0.167 | 0.049 | -3.402 | 0.000668217 | 0.007398152 | Ctu2 | protein_coding |
| ENSMUSG00000026516.8 | 5259.000 | 0.099 | 0.029 | 3.401 | 0.000671249 | 0.0074161 | Nvl | protein_coding |
| ENSMUSG00000014959.12 | 4505.750 | -0.102 | 0.030 | -3.401 | 0.000671355 | 0.0074161 | Gorasp2 | protein_coding |
| ENSMUSG00000004677.17 | 6520.185 | -0.107 | 0.031 | -3.401 | 0.000671416 | 0.0074161 | Myo9b | protein_coding |
| ENSMUSG00000034265.7 | 1955.903 | -0.174 | 0.051 | -3.400 | 0.000674577 | 0.007443278 | Zdhhc14 | protein_coding |
| ENSMUSG00000002661.14 | 327.537 | -0.226 | 0.067 | -3.399 | 0.000675329 | 0.007443278 | Alkbh7 | protein_coding |
| ENSMUSG00000022051.15 | 5039.275 | 0.113 | 0.033 | 3.399 | 0.000675939 | 0.007443278 | Bnip3l | protein_coding |

| | | | | | | | | |
|---|---|---|---|---|---|---|---|---|
| ENSMUSG00000012017.3 | 129.578 | -0.263 | 0.077 | -3.399 | 0.00067599 | 0.007443278 | Scarf2 | protein_coding |
| ENSMUSG00000029291.11 | 12512.966 | 0.088 | 0.026 | 3.392 | 0.000692936 | 0.007617967 | Rufy3 | protein_coding |
| ENSMUSG00000028986.12 | 5032.600 | 0.104 | 0.031 | 3.391 | 0.00069525 | 0.007637455 | Klhl7 | protein_coding |
| ENSMUSG00000045817.8 | 817.580 | -0.224 | 0.066 | -3.390 | 0.000698369 | 0.007665745 | Zfp36l2 | protein_coding |
| ENSMUSG00000042032.13 | 4853.874 | 0.111 | 0.033 | 3.390 | 0.000699168 | 0.007668535 | Mat2b | protein_coding |
| ENSMUSG00000028221.3 | 2005.598 | 0.127 | 0.038 | 3.388 | 0.00070418 | 0.007717507 | Tmem55a | protein_coding |
| ENSMUSG00000053293.9 | 4969.677 | -0.113 | 0.034 | -3.384 | 0.00071566 | 0.007834232 | Pom121 | protein_coding |
| ENSMUSG00000038393.14 | 1533.545 | -0.212 | 0.063 | -3.383 | 0.000715942 | 0.007834232 | Txnip | protein_coding |
| ENSMUSG00000026234.12 | 12042.221 | 0.100 | 0.030 | 3.381 | 0.000721785 | 0.007892034 | Ncl | protein_coding |
| ENSMUSG00000011589.8 | 2335.155 | -0.121 | 0.036 | -3.381 | 0.00072248 | 0.007893516 | Fsd1 | protein_coding |
| ENSMUSG00000036111.8 | 359.009 | -0.227 | 0.067 | -3.380 | 0.000725051 | 0.007915466 | Lmo1 | protein_coding |
| ENSMUSG00000027346.15 | 4381.756 | 0.116 | 0.034 | 3.378 | 0.000730013 | 0.007957637 | Gpcpd1 | protein_coding |
| ENSMUSG00000040009.6 | 3968.154 | -0.105 | 0.031 | -3.378 | 0.000730043 | 0.007957637 | Gnaz | protein_coding |
| ENSMUSG00000001229.7 | 4690.090 | -0.104 | 0.031 | -3.377 | 0.0007339 | 0.007993503 | Dpp9 | protein_coding |
| ENSMUSG00000034220.7 | 5400.391 | -0.141 | 0.042 | -3.376 | 0.000734838 | 0.007997531 | Gpc1 | protein_coding |
| ENSMUSG00000038467.15 | 6234.715 | -0.099 | 0.029 | -3.375 | 0.000736821 | 0.008012925 | Chmp4b | protein_coding |
| ENSMUSG00000058966.13 | 2889.995 | -0.143 | 0.042 | -3.373 | 0.000744141 | 0.00808629 | Fam57b | protein_coding |
| ENSMUSG00000047658.6 | 2097.567 | -0.155 | 0.046 | -3.370 | 0.000750658 | 0.008150822 | Gal3st3 | protein_coding |
| ENSMUSG00000027282.17 | 3741.159 | 0.102 | 0.030 | 3.370 | 0.000752059 | 0.008159754 | Mtch2 | protein_coding |
| ENSMUSG00000030275.6 | 28808.730 | 0.079 | 0.023 | 3.367 | 0.000760288 | 0.008242693 | Etnk1 | protein_coding |
| ENSMUSG00000045515.3 | 4033.301 | -0.160 | 0.048 | -3.366 | 0.000763676 | 0.008273063 | Pou3f3 | protein_coding |
| ENSMUSG00000059273.9 | 3620.776 | -0.119 | 0.035 | -3.365 | 0.00076503 | 0.008281363 | Zc3h4 | protein_coding |
| ENSMUSG00000043881.8 | 2301.050 | 0.135 | 0.040 | 3.364 | 0.000769375 | 0.008322008 | Kbtbd7 | protein_coding |
| ENSMUSG00000033128.8 | 3938.673 | -0.138 | 0.041 | -3.363 | 0.000770802 | 0.008331048 | Gga1 | protein_coding |
| ENSMUSG00000060510.14 | 2120.750 | 0.122 | 0.036 | 3.363 | 0.000772028 | 0.008337901 | Zfp266 | protein_coding |
| ENSMUSG00000040734.14 | 278.776 | -0.236 | 0.070 | -3.361 | 0.000775955 | 0.008373902 | Ppp1r13l | protein_coding |
| ENSMUSG00000024360.7 | 4078.360 | 0.105 | 0.031 | 3.359 | 0.000782515 | 0.00843823 | Etf1 | protein_coding |
| ENSMUSG00000027763.13 | 4287.676 | 0.118 | 0.035 | 3.358 | 0.000786309 | 0.008472659 | Mbnl1 | protein_coding |
| ENSMUSG00000037062.13 | 4405.240 | 0.096 | 0.028 | 3.356 | 0.000789487 | 0.008500403 | Sh3glb1 | protein_coding |
| ENSMUSG00000045039.9 | 14917.441 | -0.132 | 0.039 | -3.354 | 0.000795496 | 0.008558564 | Megf8 | protein_coding |
| ENSMUSG00000046792.8 | 514.953 | -0.207 | 0.062 | -3.352 | 0.000801075 | 0.008612018 | Zfp787 | protein_coding |
| ENSMUSG00000006019.15 | 653.241 | -0.185 | 0.055 | -3.352 | 0.000802751 | 0.008623459 | Dhx34 | protein_coding |
| ENSMUSG00000027993.16 | 15899.492 | 0.089 | 0.026 | 3.351 | 0.000805729 | 0.008648856 | Trim2 | protein_coding |

| | | | | | | | | |
|---|---|---|---|---|---|---|---|---|
| ENSMUSG00000042272.17 | 4019.280 | 0.101 | 0.030 | 3.346 | 0.000819547 | 0.008790481 | Sestd1 | protein_coding |
| ENSMUSG00000036492.12 | 219.643 | -0.259 | 0.077 | -3.346 | 0.000820759 | 0.008796794 | Rnf39 | protein_coding |
| ENSMUSG00000021025.8 | 794.081 | -0.236 | 0.071 | -3.345 | 0.000821468 | 0.0087977 | Nfkbia | protein_coding |
| ENSMUSG00000022337.6 | 2355.328 | 0.130 | 0.039 | 3.344 | 0.000825264 | 0.008818547 | Emc2 | protein_coding |
| ENSMUSG00000027401.9 | 158.423 | 0.261 | 0.078 | 3.344 | 0.000825292 | 0.008818547 | Tgm3 | protein_coding |
| ENSMUSG00000015671.11 | 3369.978 | 0.104 | 0.031 | 3.344 | 0.00082679 | 0.008824467 | Psma2 | protein_coding |
| ENSMUSG00000047824.13 | 2998.569 | -0.116 | 0.035 | -3.344 | 0.000827098 | 0.008824467 | Pygo2 | protein_coding |
| ENSMUSG00000057182.14 | 3437.671 | 0.111 | 0.033 | 3.343 | 0.000829186 | 0.008840056 | Scn3a | protein_coding |
| ENSMUSG00000046351.10 | 1998.731 | 0.143 | 0.043 | 3.338 | 0.000843855 | 0.008989637 | Zfp322a | protein_coding |
| ENSMUSG00000063406.11 | 1350.574 | 0.147 | 0.044 | 3.338 | 0.000844777 | 0.008992661 | Tmed5 | protein_coding |
| ENSMUSG00000000915.15 | 4917.733 | -0.100 | 0.030 | -3.337 | 0.000847578 | 0.009015668 | Hip1r | protein_coding |
| ENSMUSG00000025732.5 | 274.915 | -0.243 | 0.073 | -3.335 | 0.000853073 | 0.009067275 | Fam195a | protein_coding |
| ENSMUSG00000018849.6 | 6010.810 | -0.123 | 0.037 | -3.334 | 0.000856764 | 0.009099646 | Wwc1 | protein_coding |
| ENSMUSG00000024570.6 | 1011.667 | -0.157 | 0.047 | -3.334 | 0.000857422 | 0.009099778 | Rbfa | protein_coding |
| ENSMUSG00000039470.15 | 2248.516 | 0.144 | 0.043 | 3.329 | 0.000870753 | 0.009234308 | Zdhhc2 | protein_coding |
| ENSMUSG00000038437.11 | 12141.616 | -0.109 | 0.033 | -3.329 | 0.000873018 | 0.009251362 | Mllt6 | protein_coding |
| ENSMUSG00000037296.7 | 646.029 | -0.193 | 0.058 | -3.325 | 0.000883533 | 0.009355753 | Lsm1 | protein_coding |
| ENSMUSG00000068882.13 | 4739.950 | 0.111 | 0.033 | 3.325 | 0.00088479 | 0.009362033 | Ssb | protein_coding |
| ENSMUSG00000039656.16 | 3247.027 | -0.127 | 0.038 | -3.324 | 0.000885938 | 0.009367149 | Rxrb | protein_coding |
| ENSMUSG00000028575.11 | 72.690 | 0.256 | 0.077 | 3.323 | 0.000889627 | NA | Eqtn | protein_coding |
| ENSMUSG00000035131.14 | 879.241 | 0.192 | 0.058 | 3.321 | 0.000896052 | 0.009466984 | Brinp3 | protein_coding |
| ENSMUSG00000031865.16 | 23289.070 | -0.099 | 0.030 | -3.319 | 0.000902132 | 0.009524085 | Dctn1 | protein_coding |
| ENSMUSG00000033411.16 | 1163.940 | 0.163 | 0.049 | 3.319 | 0.000903301 | 0.009529281 | Ctdspl2 | protein_coding |
| ENSMUSG00000064293.14 | 2883.290 | 0.144 | 0.043 | 3.315 | 0.000915658 | 0.009652417 | Cntn4 | protein_coding |
| ENSMUSG00000035762.10 | 969.610 | 0.167 | 0.050 | 3.310 | 0.000931592 | 0.00980874 | Tmem161b | protein_coding |
| ENSMUSG00000033569.17 | 5208.784 | 0.106 | 0.032 | 3.310 | 0.000931879 | 0.00980874 | Adgrb3 | protein_coding |
| ENSMUSG00000043929.16 | 784.702 | 0.196 | 0.059 | 3.308 | 0.000938475 | 0.009870137 | Klhl15 | protein_coding |
| ENSMUSG00000090015.8 | 1686.958 | 0.137 | 0.041 | 3.308 | 0.000939113 | 0.009870137 | Gm15446 | protein_coding |
| ENSMUSG00000028433.14 | 2677.349 | -0.121 | 0.036 | -3.308 | 0.00094048 | 0.009877145 | Ubap2 | protein_coding |
| ENSMUSG00000028282.12 | 856.403 | 0.169 | 0.051 | 3.307 | 0.000944394 | 0.009910863 | Casp8ap2 | protein_coding |
| ENSMUSG00000031786.7 | 148.729 | -0.236 | 0.071 | -3.305 | 0.000949804 | 0.009960221 | Drc7 | protein_coding |
| ENSMUSG00000036644.15 | 12650.004 | -0.084 | 0.025 | -3.304 | 0.000952321 | 0.009978723 | Tbc1d9b | protein_coding |
| ENSMUSG00000052681.8 | 2943.987 | 0.110 | 0.033 | 3.304 | 0.000952984 | 0.009978723 | Rap1b | protein_coding |

| | | | | | | | | |
|---|---|---|---|---|---|---|---|---|
| ENSMUSG00000059325.14 | 1671.228 | -0.180 | 0.054 | -3.303 | 0.00095657 | 0.010008828 | Hopx | protein_coding |
| ENSMUSG00000035559.9 | 444.680 | -0.204 | 0.062 | -3.302 | 0.000959781 | 0.010034977 | Mpv17l2 | protein_coding |
| ENSMUSG00000073411.11 | 1572.079 | -0.220 | 0.067 | -3.302 | 0.000960493 | 0.010034978 | H2-D1 | protein_coding |
| ENSMUSG00000039989.7 | 1945.186 | -0.131 | 0.040 | -3.301 | 0.000962519 | 0.010048696 | Cbx4 | protein_coding |
| ENSMUSG00000024958.12 | 3619.926 | -0.118 | 0.036 | -3.300 | 0.000968146 | 0.010099966 | Gpr137 | protein_coding |
| ENSMUSG00000036620.14 | 2486.763 | -0.120 | 0.036 | -3.299 | 0.00096983 | 0.010110045 | Mgat4b | protein_coding |
| ENSMUSG00000026239.14 | 1137.089 | -0.162 | 0.049 | -3.299 | 0.00097162 | 0.010121221 | Pde6d | protein_coding |
| ENSMUSG00000010080.15 | 52.725 | -0.194 | 0.059 | -3.297 | 0.000976777 | NA | Epn3 | protein_coding |
| ENSMUSG00000030967.15 | 2489.935 | 0.114 | 0.035 | 3.296 | 0.000979561 | 0.0101964 | Zranb1 | protein_coding |
| ENSMUSG00000028005.13 | 9567.633 | 0.120 | 0.036 | 3.295 | 0.00098298 | 0.010220499 | Gucy1b3 | protein_coding |
| ENSMUSG00000040123.16 | 2456.995 | 0.127 | 0.038 | 3.295 | 0.000983326 | 0.010220499 | Zmym5 | protein_coding |
| ENSMUSG00000028249.15 | 7637.736 | 0.093 | 0.028 | 3.294 | 0.000988168 | 0.010263258 | Sdcbp | protein_coding |
| ENSMUSG00000023938.7 | 1890.005 | -0.166 | 0.050 | -3.292 | 0.000993009 | 0.010305239 | Aars2 | protein_coding |
| ENSMUSG00000029246.14 | 826.366 | 0.165 | 0.050 | 3.292 | 0.000993672 | 0.010305239 | Ppat | protein_coding |
| ENSMUSG00000020608.7 | 2057.113 | 0.123 | 0.037 | 3.291 | 0.00099684 | 0.010323634 | Smc6 | protein_coding |
| ENSMUSG00000044576.6 | 709.716 | -0.175 | 0.053 | -3.291 | 0.000996911 | 0.010323634 | Garem2 | protein_coding |
| ENSMUSG00000021770.10 | 4247.179 | 0.098 | 0.030 | 3.291 | 0.000998698 | 0.010330755 | Samd8 | protein_coding |
| ENSMUSG00000023232.17 | 711.636 | -0.187 | 0.057 | -3.291 | 0.000999065 | 0.010330755 | Serinc2 | protein_coding |
| ENSMUSG00000060780.2 | 2235.204 | -0.127 | 0.039 | -3.289 | 0.001005085 | 0.010385384 | Lrrtm1 | protein_coding |
| ENSMUSG00000090000.1 | 906.811 | 0.167 | 0.051 | 3.289 | 0.001006772 | 0.010395199 | Ier3ip1 | protein_coding |
| ENSMUSG00000022292.15 | 1995.888 | 0.127 | 0.039 | 3.286 | 0.001016444 | 0.010478072 | Rrm2b | protein_coding |
| ENSMUSG00000038822.15 | 3430.263 | 0.123 | 0.037 | 3.286 | 0.00101702 | 0.010478072 | Hace1 | protein_coding |
| ENSMUSG00000052751.16 | 1837.640 | -0.159 | 0.048 | -3.286 | 0.001017029 | 0.010478072 | Repin1 | protein_coding |
| ENSMUSG00000055421.8 | 5189.959 | 0.135 | 0.041 | 3.285 | 0.00101858 | 0.01048367 | Pcdh9 | protein_coding |
| ENSMUSG00000004892.13 | 12932.699 | -0.130 | 0.039 | -3.285 | 0.00101906 | 0.01048367 | Bcan | protein_coding |
| ENSMUSG00000030062.7 | 6516.429 | -0.091 | 0.028 | -3.285 | 0.001019978 | 0.010485467 | Rpn1 | protein_coding |
| ENSMUSG00000074781.5 | 6408.179 | 0.091 | 0.028 | 3.285 | 0.001020917 | 0.010487471 | Ube2n | protein_coding |
| ENSMUSG00000039219.18 | 3177.541 | 0.139 | 0.042 | 3.284 | 0.001024189 | 0.010513413 | Arid4b | protein_coding |
| ENSMUSG00000040423.10 | 2495.050 | 0.150 | 0.046 | 3.284 | 0.001024954 | 0.010513609 | Rc3h1 | protein_coding |
| ENSMUSG00000041028.15 | 16381.306 | 0.081 | 0.025 | 3.282 | 0.001029288 | 0.010550394 | Ghitm | protein_coding |
| ENSMUSG00000032932.14 | 2452.164 | 0.130 | 0.040 | 3.282 | 0.001031 | 0.010560255 | Hspa13 | protein_coding |
| ENSMUSG00000079509.10 | 1216.756 | 0.149 | 0.045 | 3.281 | 0.001032595 | 0.010568918 | Zfx | protein_coding |
| ENSMUSG00000026374.14 | 4138.734 | 0.114 | 0.035 | 3.281 | 0.001036196 | 0.010598078 | Tsn | protein_coding |

| | | | | | | | | |
|---|---|---|---|---|---|---|---|---|
| ENSMUSG00000042834.14 | 1520.390 | 0.141 | 0.043 | 3.280 | 0.001037175 | 0.010600398 | Nrep | protein_coding |
| ENSMUSG00000030030.5 | 1374.621 | -0.135 | 0.041 | -3.280 | 0.001039751 | 0.010611329 | 1700003E16Rik | protein_coding |
| ENSMUSG00000015461.15 | 3114.680 | -0.116 | 0.036 | -3.279 | 0.001043408 | 0.010634412 | Atf6b | protein_coding |
| ENSMUSG00000042625.16 | 6523.260 | -0.117 | 0.036 | -3.279 | 0.001043522 | 0.010634412 | Safb2 | protein_coding |
| ENSMUSG00000048078.16 | 11210.618 | -0.097 | 0.030 | -3.276 | 0.001051896 | 0.010707286 | Tenm4 | protein_coding |
| ENSMUSG00000067608.4 | 82.546 | 0.121 | 0.037 | 3.276 | 0.001052192 | 0.010707286 | Pcna-ps2 | protein_coding |
| ENSMUSG00000060681.15 | 6340.197 | 0.098 | 0.030 | 3.275 | 0.001054849 | 0.01071885 | Slc9a6 | protein_coding |
| ENSMUSG00000025040.13 | 1562.073 | 0.152 | 0.047 | 3.273 | 0.001064185 | 0.010805922 | Fundc1 | protein_coding |
| ENSMUSG00000039220.16 | 2901.628 | -0.135 | 0.041 | -3.272 | 0.001066937 | 0.010826066 | Ppp1r10 | protein_coding |
| ENSMUSG00000022450.5 | 1721.600 | -0.150 | 0.046 | -3.272 | 0.001067949 | 0.010828541 | Ndufa6 | protein_coding |
| ENSMUSG00000039157.12 | 5637.059 | -0.117 | 0.036 | -3.271 | 0.001069794 | 0.010839454 | Fam102a | protein_coding |
| ENSMUSG00000030020.13 | 12163.407 | 0.079 | 0.024 | 3.270 | 0.001074363 | 0.010877923 | Prickle2 | protein_coding |
| ENSMUSG00000023806.9 | 274.647 | -0.233 | 0.071 | -3.270 | 0.001076353 | 0.010890251 | Rsph3b | protein_coding |
| ENSMUSG00000024479.2 | 7699.821 | 0.120 | 0.037 | 3.270 | 0.001077186 | 0.010890863 | Mal2 | protein_coding |
| ENSMUSG00000028691.12 | 4838.204 | 0.118 | 0.036 | 3.268 | 0.001081194 | 0.01092355 | Prdx1 | protein_coding |
| ENSMUSG00000078862.10 | 853.625 | 0.162 | 0.049 | 3.268 | 0.001082399 | 0.010927883 | Gm14326 | protein_coding |
| ENSMUSG00000074406.5 | 562.418 | -0.192 | 0.059 | -3.266 | 0.001091782 | 0.01101473 | Zfp628 | protein_coding |
| ENSMUSG00000025199.16 | 3538.759 | 0.110 | 0.034 | 3.264 | 0.001097039 | 0.011059844 | Chuk | protein_coding |
| ENSMUSG00000060166.5 | 5776.769 | -0.152 | 0.047 | -3.263 | 0.001102649 | 0.011108455 | Zdhhc8 | protein_coding |
| ENSMUSG00000040710.10 | 854.910 | 0.166 | 0.051 | 3.263 | 0.001104082 | 0.011114948 | St8sia4 | protein_coding |
| ENSMUSG00000023025.15 | 2263.371 | 0.135 | 0.041 | 3.260 | 0.001112166 | 0.011188344 | Larp4 | protein_coding |
| ENSMUSG00000040549.16 | 9504.899 | 0.090 | 0.028 | 3.259 | 0.001119567 | 0.011254762 | Ckap5 | protein_coding |
| ENSMUSG00000042460.5 | 1130.914 | 0.171 | 0.053 | 3.258 | 0.001120667 | 0.011257794 | C1galt1 | protein_coding |
| ENSMUSG00000037351.9 | 15502.008 | -0.074 | 0.023 | -3.257 | 0.001123991 | 0.011283145 | Actr1b | protein_coding |
| ENSMUSG00000034912.17 | 2397.017 | 0.123 | 0.038 | 3.257 | 0.001125093 | 0.011286164 | Mdga2 | protein_coding |
| ENSMUSG00000031661.12 | 735.868 | -0.183 | 0.056 | -3.256 | 0.001129441 | 0.011320972 | Nkd1 | protein_coding |
| ENSMUSG00000026037.14 | 1496.651 | 0.148 | 0.046 | 3.256 | 0.001130708 | 0.011320972 | Orc2 | protein_coding |
| ENSMUSG00000028975.16 | 2085.046 | -0.130 | 0.040 | -3.256 | 0.001130973 | 0.011320972 | Pex14 | protein_coding |
| ENSMUSG00000021967.3 | 654.203 | -0.199 | 0.061 | -3.255 | 0.00113421 | 0.011345316 | Mrpl57 | protein_coding |
| ENSMUSG00000041120.6 | 2304.267 | -0.174 | 0.054 | -3.255 | 0.001135604 | 0.011348028 | Nbl1 | protein_coding |
| ENSMUSG00000001729.14 | 7839.082 | -0.095 | 0.029 | -3.254 | 0.001136091 | 0.011348028 | Akt1 | protein_coding |
| ENSMUSG00000073433.10 | 1949.666 | -0.148 | 0.046 | -3.253 | 0.001143275 | 0.011410745 | Arhgdig | protein_coding |
| ENSMUSG00000031431.13 | 3665.975 | -0.154 | 0.047 | -3.252 | 0.001144699 | 0.011410745 | Tsc22d3 | protein_coding |

| | | | | | | | | |
|---|---|---|---|---|---|---|---|---|
| ENSMUSG00000027011.14 | 2595.216 | 0.120 | 0.037 | 3.252 | 0.001144799 | 0.011410745 | Ube2e3 | protein_coding |
| ENSMUSG00000020307.15 | 1248.299 | -0.159 | 0.049 | -3.251 | 0.001149793 | 0.011449701 | Cdc34 | protein_coding |
| ENSMUSG00000032580.12 | 9284.368 | 0.088 | 0.027 | 3.249 | 0.001156264 | 0.011500625 | Rbm5 | protein_coding |
| ENSMUSG00000023017.9 | 4202.407 | -0.103 | 0.032 | -3.249 | 0.001158385 | 0.011513595 | Asic1 | protein_coding |
| ENSMUSG00000062647.16 | 6556.315 | -0.095 | 0.029 | -3.249 | 0.001160023 | 0.011521747 | Rpl7a | protein_coding |
| ENSMUSG00000022342.5 | 3981.111 | 0.127 | 0.039 | 3.248 | 0.00116211 | 0.01153435 | Kcnv1 | protein_coding |
| ENSMUSG00000025227.14 | 1287.587 | -0.137 | 0.042 | -3.248 | 0.001162954 | 0.011534607 | Mfsd13a | protein_coding |
| ENSMUSG00000036466.17 | 832.379 | -0.184 | 0.057 | -3.246 | 0.001172227 | 0.011618398 | Megf11 | protein_coding |
| ENSMUSG00000031622.16 | 5257.382 | -0.113 | 0.035 | -3.244 | 0.00117991 | 0.011679163 | Sin3b | protein_coding |
| ENSMUSG00000019802.13 | 4650.027 | 0.101 | 0.031 | 3.244 | 0.001180015 | 0.011679163 | Sec63 | protein_coding |
| ENSMUSG00000048807.2 | 1114.240 | -0.183 | 0.056 | -3.243 | 0.001180971 | 0.011680421 | Slc35e4 | protein_coding |
| ENSMUSG00000021294.7 | 891.365 | -0.203 | 0.063 | -3.242 | 0.001187944 | 0.011741155 | Kif26a | protein_coding |
| ENSMUSG00000046364.14 | 3816.168 | -0.139 | 0.043 | -3.240 | 0.001194232 | 0.01179503 | Rpl27a | protein_coding |
| ENSMUSG00000040455.17 | 2611.528 | 0.114 | 0.035 | 3.239 | 0.001197441 | 0.011818441 | Usp45 | protein_coding |
| ENSMUSG00000028522.16 | 1614.640 | 0.151 | 0.047 | 3.239 | 0.001198504 | 0.011820659 | Mier1 | protein_coding |
| ENSMUSG00000028312.19 | 625.633 | 0.178 | 0.055 | 3.237 | 0.001209496 | 0.011920727 | Smc2 | protein_coding |
| ENSMUSG00000040044.11 | 1500.497 | 0.149 | 0.046 | 3.236 | 0.001210654 | 0.011923803 | Orc3 | protein_coding |
| ENSMUSG00000058690.14 | 6122.043 | 0.086 | 0.027 | 3.233 | 0.001226965 | 0.012076012 | Ccser2 | protein_coding |
| ENSMUSG00000057561.9 | 2079.058 | 0.130 | 0.040 | 3.230 | 0.001237426 | 0.012170466 | Eif1a | protein_coding |
| ENSMUSG00000044349.15 | 146191.718 | 0.070 | 0.022 | 3.229 | 0.001242187 | 0.012208778 | Snhg11 | protein_coding |
| ENSMUSG00000030061.16 | 3302.816 | 0.110 | 0.034 | 3.229 | 0.00124313 | 0.012209526 | Uba3 | protein_coding |
| ENSMUSG00000024073.14 | 11043.469 | 0.078 | 0.024 | 3.228 | 0.001245903 | 0.012228245 | Birc6 | protein_coding |
| ENSMUSG00000031134.16 | 3420.313 | 0.100 | 0.031 | 3.228 | 0.001247013 | 0.012230623 | Rbmx | protein_coding |
| ENSMUSG00000000276.11 | 4939.458 | 0.107 | 0.033 | 3.226 | 0.001254276 | 0.012293297 | Dgke | protein_coding |
| ENSMUSG00000005732.14 | 1458.257 | -0.155 | 0.048 | -3.226 | 0.001256647 | 0.012307975 | Ranbp1 | protein_coding |
| ENSMUSG00000032046.15 | 11123.209 | -0.114 | 0.035 | -3.225 | 0.001257803 | 0.012310745 | Abhd12 | protein_coding |
| ENSMUSG00000044700.15 | 5147.725 | -0.116 | 0.036 | -3.225 | 0.001259262 | 0.012316471 | Tmem201 | protein_coding |
| ENSMUSG00000070000.13 | 5323.079 | -0.133 | 0.041 | -3.224 | 0.001262348 | 0.012336226 | Fcho1 | protein_coding |
| ENSMUSG00000024109.18 | 17820.089 | 0.067 | 0.021 | 3.224 | 0.001263032 | 0.012336226 | Nrxn1 | protein_coding |
| ENSMUSG00000021596.16 | 3988.672 | 0.154 | 0.048 | 3.222 | 0.001271759 | 0.012412862 | Mctp1 | protein_coding |
| ENSMUSG00000022311.15 | 2725.836 | 0.124 | 0.038 | 3.222 | 0.001273945 | 0.012425594 | Csmd3 | protein_coding |
| ENSMUSG00000037486.18 | 2066.521 | 0.116 | 0.036 | 3.221 | 0.001279157 | 0.012467798 | Asxl2 | protein_coding |
| ENSMUSG00000021109.13 | 2897.629 | 0.128 | 0.040 | 3.220 | 0.001282863 | 0.012494086 | Hif1a | protein_coding |

| | | | | | | | | |
|---|---|---|---|---|---|---|---|---|
| ENSMUSG00000054499.9 | 1480.281 | -0.174 | 0.054 | -3.220 | 0.001283627 | 0.012494086 | Dedd2 | protein_coding |
| ENSMUSG00000021550.5 | 333.007 | -0.217 | 0.068 | -3.217 | 0.001295406 | 0.012600036 | 2210016F16Rik | protein_coding |
| ENSMUSG00000059851.15 | 1090.301 | -0.146 | 0.045 | -3.212 | 0.001316059 | 0.01278192 | Kmt5c | protein_coding |
| ENSMUSG00000028603.15 | 2744.473 | 0.115 | 0.036 | 3.212 | 0.001316755 | 0.01278192 | Scp2 | protein_coding |
| ENSMUSG00000109901.1 | 1107.645 | 0.158 | 0.049 | 3.212 | 0.001316826 | 0.01278192 | Chmp1b | protein_coding |
| ENSMUSG00000060073.9 | 3337.244 | 0.108 | 0.034 | 3.211 | 0.001323825 | 0.012841014 | Psma3 | protein_coding |
| ENSMUSG00000049796.4 | 130.381 | -0.250 | 0.078 | -3.207 | 0.001339931 | 0.012988296 | Crh | protein_coding |
| ENSMUSG00000022119.15 | 2986.386 | 0.107 | 0.033 | 3.206 | 0.001344242 | 0.013013251 | Rbm26 | protein_coding |
| ENSMUSG00000025059.16 | 1440.527 | 0.129 | 0.040 | 3.206 | 0.001344352 | 0.013013251 | Gk | protein_coding |
| ENSMUSG00000002342.17 | 1460.558 | -0.158 | 0.049 | -3.204 | 0.001354686 | 0.0130953 | Tmem161a | protein_coding |
| ENSMUSG00000049659.15 | 5001.066 | 0.111 | 0.035 | 3.204 | 0.001356068 | 0.013099675 | Aftph | protein_coding |
| ENSMUSG00000045210.8 | 4305.115 | 0.095 | 0.030 | 3.202 | 0.001364134 | 0.013168566 | Vcpip1 | protein_coding |
| ENSMUSG00000054737.11 | 755.876 | 0.166 | 0.052 | 3.202 | 0.001365578 | 0.0131705 | Zfp182 | protein_coding |
| ENSMUSG00000022240.9 | 10594.049 | -0.101 | 0.032 | -3.202 | 0.001366203 | 0.0131705 | Ctnnd2 | protein_coding |
| ENSMUSG00000061689.15 | 15796.965 | -0.117 | 0.037 | -3.201 | 0.001370406 | 0.013201984 | Dlgap4 | protein_coding |
| ENSMUSG00000028676.17 | 3582.685 | 0.115 | 0.036 | 3.199 | 0.001379168 | 0.013277314 | Srsf10 | protein_coding |
| ENSMUSG00000034653.11 | 2352.634 | 0.127 | 0.040 | 3.198 | 0.001384598 | 0.013320492 | Ythdc2 | protein_coding |
| ENSMUSG00000028034.15 | 7704.143 | 0.090 | 0.028 | 3.197 | 0.001386512 | 0.013329809 | Fubp1 | protein_coding |
| ENSMUSG00000022490.6 | 2260.824 | -0.140 | 0.044 | -3.196 | 0.001394811 | 0.01340045 | Ppp1r1a | protein_coding |
| ENSMUSG00000034297.14 | 4542.876 | 0.116 | 0.036 | 3.195 | 0.001398469 | 0.01341179 | Med13 | protein_coding |
| ENSMUSG00000066798.3 | 814.080 | 0.173 | 0.054 | 3.195 | 0.001398689 | 0.01341179 | Zbtb6 | protein_coding |
| ENSMUSG00000092486.1 | 288.292 | 0.221 | 0.069 | 3.195 | 0.001399077 | 0.01341179 | 2610524H06Rik | protein_coding |
| ENSMUSG00000021704.7 | 2964.747 | 0.103 | 0.032 | 3.195 | 0.001399797 | 0.01341179 | Mtx3 | protein_coding |
| ENSMUSG00000050822.11 | 358.190 | -0.243 | 0.076 | -3.192 | 0.001411448 | 0.013509196 | Slc29a4 | protein_coding |
| ENSMUSG00000002980.14 | 726.290 | -0.210 | 0.066 | -3.192 | 0.001411881 | 0.013509196 | Bcam | protein_coding |
| ENSMUSG00000033342.13 | 2345.788 | 0.114 | 0.036 | 3.188 | 0.001432329 | 0.013689355 | Plppr5 | protein_coding |
| ENSMUSG00000020949.9 | 4557.417 | 0.105 | 0.033 | 3.188 | 0.001432652 | 0.013689355 | Fkbp3 | protein_coding |
| ENSMUSG00000022307.15 | 15880.246 | 0.091 | 0.028 | 3.187 | 0.001435099 | 0.013703448 | Oxr1 | protein_coding |
| ENSMUSG00000021068.16 | 5199.583 | 0.118 | 0.037 | 3.187 | 0.001437177 | 0.013713998 | Nin | protein_coding |
| ENSMUSG00000036202.15 | 1965.542 | 0.115 | 0.036 | 3.186 | 0.001441559 | 0.013737866 | Rif1 | protein_coding |
| ENSMUSG00000034154.15 | 2088.869 | 0.111 | 0.035 | 3.186 | 0.001441628 | 0.013737866 | Ino80 | protein_coding |
| ENSMUSG00000025993.10 | 543.093 | 0.216 | 0.068 | 3.185 | 0.001447043 | 0.013780148 | Slc40a1 | protein_coding |
| ENSMUSG00000054942.13 | 5935.442 | 0.092 | 0.029 | 3.184 | 0.001450629 | 0.013800101 | Miga1 | protein_coding |

| | | | | | | | | |
|---|---|---|---|---|---|---|---|---|
| ENSMUSG00000023010.14 | 13319.872 | -0.077 | 0.024 | -3.184 | 0.001451096 | 0.013800101 | Tmbim6 | protein_coding |
| ENSMUSG00000064065.15 | 6992.609 | 0.102 | 0.032 | 3.183 | 0.001458973 | 0.013856307 | Ipcef1 | protein_coding |
| ENSMUSG00000027459.16 | 262.946 | -0.237 | 0.075 | -3.182 | 0.001462978 | 0.013884996 | Fam110a | protein_coding |
| ENSMUSG00000045538.6 | 524.597 | -0.189 | 0.059 | -3.182 | 0.00146458 | 0.013890839 | Ddx28 | protein_coding |
| ENSMUSG00000036887.5 | 1424.052 | -0.147 | 0.046 | -3.180 | 0.001471136 | 0.013943033 | C1qa | protein_coding |
| ENSMUSG00000021226.7 | 240.802 | -0.237 | 0.074 | -3.180 | 0.001472061 | 0.013943033 | Acot2 | protein_coding |
| ENSMUSG00000027900.15 | 974.332 | 0.158 | 0.050 | 3.180 | 0.001473356 | 0.013945926 | Dram2 | protein_coding |
| ENSMUSG00000054162.15 | 3467.321 | 0.163 | 0.051 | 3.179 | 0.001477668 | 0.013977351 | Spock3 | protein_coding |
| ENSMUSG00000031333.7 | 2188.897 | 0.110 | 0.035 | 3.177 | 0.001487104 | 0.014057174 | Abcb7 | protein_coding |
| ENSMUSG00000087370.3 | 3783.515 | 0.144 | 0.045 | 3.177 | 0.001488924 | 0.014064944 | Tmem170b | protein_coding |
| ENSMUSG00000037138.17 | 3870.592 | -0.102 | 0.032 | -3.176 | 0.001491173 | 0.014076754 | Aff3 | protein_coding |
| ENSMUSG00000025949.16 | 3455.633 | 0.092 | 0.029 | 3.176 | 0.001495106 | 0.014104433 | Pikfyve | protein_coding |
| ENSMUSG00000024112.16 | 6634.118 | -0.161 | 0.051 | -3.175 | 0.001500171 | 0.014142748 | Cacna1h | protein_coding |
| ENSMUSG00000072501.4 | 3698.171 | 0.119 | 0.037 | 3.174 | 0.001502066 | 0.014151145 | Phf20l1 | protein_coding |
| ENSMUSG00000053929.16 | 6660.372 | -0.078 | 0.025 | -3.173 | 0.001510194 | 0.014208725 | Cyhr1 | protein_coding |
| ENSMUSG00000044365.15 | 2164.088 | 0.114 | 0.036 | 3.171 | 0.001516542 | 0.014258933 | Cxxc4 | protein_coding |
| ENSMUSG00000039704.6 | 2477.362 | 0.107 | 0.034 | 3.169 | 0.001527699 | 0.014348442 | Lmbrd2 | protein_coding |
| ENSMUSG00000048388.3 | 21205.677 | 0.102 | 0.032 | 3.169 | 0.001528098 | 0.014348442 | Fam171b | protein_coding |
| ENSMUSG00000059890.16 | 4402.156 | 0.096 | 0.030 | 3.169 | 0.001530878 | 0.014364973 | Ube4a | protein_coding |
| ENSMUSG00000028657.14 | 5154.742 | 0.094 | 0.030 | 3.168 | 0.00153702 | 0.014413014 | Ppt1 | protein_coding |
| ENSMUSG00000001750.15 | 1037.739 | -0.155 | 0.049 | -3.167 | 0.001541289 | 0.014443436 | Tcirg1 | protein_coding |
| ENSMUSG00000026159.13 | 5669.656 | 0.093 | 0.029 | 3.166 | 0.001545988 | 0.01447784 | Agfg1 | protein_coding |
| ENSMUSG00000019856.14 | 1660.072 | 0.163 | 0.052 | 3.165 | 0.001550176 | 0.014507423 | Fam184a | protein_coding |
| ENSMUSG00000035007.5 | 2906.173 | -0.107 | 0.034 | -3.165 | 0.001552752 | 0.014521885 | Rundc1 | protein_coding |
| ENSMUSG00000021494.8 | 2258.826 | -0.110 | 0.035 | -3.164 | 0.001553898 | 0.014522965 | Ddx41 | protein_coding |
| ENSMUSG00000030878.11 | 637.753 | -0.184 | 0.058 | -3.164 | 0.001554981 | 0.014523465 | Cdr2 | protein_coding |
| ENSMUSG00000029516.19 | 9066.984 | -0.117 | 0.037 | -3.162 | 0.001566851 | 0.014624634 | Cit | protein_coding |
| ENSMUSG00000006575.14 | 14526.732 | -0.116 | 0.037 | -3.161 | 0.00156979 | 0.014642373 | Rundc3a | protein_coding |
| ENSMUSG00000020836.15 | 3065.815 | -0.128 | 0.041 | -3.159 | 0.00158417 | 0.014766726 | Coro6 | protein_coding |
| ENSMUSG00000035513.19 | 1056.523 | -0.189 | 0.060 | -3.158 | 0.001586156 | 0.014775467 | Ntng2 | protein_coding |
| ENSMUSG00000060301.6 | 1702.396 | 0.124 | 0.039 | 3.156 | 0.001597592 | 0.014872165 | 2610008E11Rik | protein_coding |
| ENSMUSG00000025158.7 | 3045.088 | -0.134 | 0.042 | -3.156 | 0.001600936 | 0.014893457 | Rfng | protein_coding |
| ENSMUSG00000029359.13 | 1229.718 | -0.185 | 0.059 | -3.154 | 0.001611436 | 0.014981256 | Tesc | protein_coding |

| | | | | | | | | |
|---|---|---|---|---|---|---|---|---|
| ENSMUSG00000050587.14 | 5701.233 | 0.101 | 0.032 | 3.152 | 0.001622717 | 0.015076186 | Lrrc4c | protein_coding |
| ENSMUSG00000025731.15 | 799.947 | -0.193 | 0.061 | -3.151 | 0.001625744 | 0.015093818 | Mettl26 | protein_coding |
| ENSMUSG00000028465.16 | 3626.520 | -0.132 | 0.042 | -3.151 | 0.001627405 | 0.015093818 | Tln1 | protein_coding |
| ENSMUSG00000031343.13 | 8450.417 | 0.093 | 0.030 | 3.151 | 0.001628328 | 0.015093818 | Gabra3 | protein_coding |
| ENSMUSG00000073563.2 | 2982.742 | 0.098 | 0.031 | 3.151 | 0.001629513 | 0.015093818 | Csnk1g3 | protein_coding |
| ENSMUSG00000028382.15 | 1249.187 | 0.158 | 0.050 | 3.150 | 0.00162997 | 0.015093818 | Ptbp3 | protein_coding |
| ENSMUSG00000035206.10 | 2559.734 | -0.137 | 0.044 | -3.150 | 0.001632945 | 0.015111444 | Sppl2b | protein_coding |
| ENSMUSG00000026728.9 | 2018.692 | -0.191 | 0.061 | -3.150 | 0.001634475 | 0.015115676 | Vim | protein_coding |
| ENSMUSG00000060126.14 | 8546.325 | -0.150 | 0.048 | -3.149 | 0.001636845 | 0.015127666 | Tpt1 | protein_coding |
| ENSMUSG00000008683.16 | 6385.038 | 0.081 | 0.026 | 3.149 | 0.001639494 | 0.015142216 | Rps15a | protein_coding |
| ENSMUSG00000020205.8 | 862.068 | -0.176 | 0.056 | -3.147 | 0.00164857 | 0.015216076 | Phlda1 | protein_coding |
| ENSMUSG00000031729.6 | 5618.037 | 0.095 | 0.030 | 3.146 | 0.001652701 | 0.015244218 | Ist1 | protein_coding |
| ENSMUSG00000054976.14 | 2215.629 | 0.145 | 0.046 | 3.146 | 0.001655313 | 0.015258328 | Nyap2 | protein_coding |
| ENSMUSG00000049252.17 | 4318.109 | 0.155 | 0.049 | 3.145 | 0.001663276 | 0.015321711 | Lrp1b | protein_coding |
| ENSMUSG00000022884.14 | 26994.710 | 0.083 | 0.027 | 3.144 | 0.001664929 | 0.015326917 | Eif4a2 | protein_coding |
| ENSMUSG00000022760.15 | 830.444 | -0.187 | 0.059 | -3.143 | 0.001672566 | 0.015387169 | Thap7 | protein_coding |
| ENSMUSG00000032737.13 | 2278.939 | -0.130 | 0.041 | -3.140 | 0.001691813 | 0.015554086 | Inppl1 | protein_coding |
| ENSMUSG00000051062.6 | 1090.086 | -0.165 | 0.053 | -3.134 | 0.001724812 | 0.01584713 | Fbll1 | protein_coding |
| ENSMUSG00000008575.17 | 4789.824 | 0.105 | 0.034 | 3.133 | 0.001728109 | 0.015867081 | Nfib | protein_coding |
| ENSMUSG00000052040.10 | 9479.757 | -0.080 | 0.026 | -3.133 | 0.001729726 | 0.01587159 | Klf13 | protein_coding |
| ENSMUSG00000031591.14 | 3161.398 | 0.102 | 0.033 | 3.133 | 0.001732631 | 0.015887901 | Asah1 | protein_coding |
| ENSMUSG00000044147.8 | 898.342 | -0.190 | 0.061 | -3.132 | 0.001735842 | 0.015906994 | Arf6 | protein_coding |
| ENSMUSG00000043670.4 | 5540.276 | -0.113 | 0.036 | -3.131 | 0.001743691 | 0.015968536 | Diras1 | protein_coding |
| ENSMUSG00000002910.11 | 636.054 | -0.210 | 0.067 | -3.129 | 0.001751567 | 0.016030254 | Arrdc2 | protein_coding |
| ENSMUSG00000037685.15 | 16500.266 | 0.084 | 0.027 | 3.129 | 0.001756823 | 0.016067921 | Atp8a1 | protein_coding |
| ENSMUSG00000024924.14 | 4599.902 | 0.094 | 0.030 | 3.126 | 0.001773924 | 0.016213801 | Vldlr | protein_coding |
| ENSMUSG00000025171.1 | 342.850 | -0.208 | 0.066 | -3.125 | 0.001775184 | 0.016214804 | Ubtd1 | protein_coding |
| ENSMUSG00000000420.15 | 1901.030 | 0.115 | 0.037 | 3.125 | 0.001777889 | 0.016228999 | Galnt1 | protein_coding |
| ENSMUSG00000022391.15 | 13369.638 | -0.068 | 0.022 | -3.122 | 0.001793432 | 0.016360276 | Rangap1 | protein_coding |
| ENSMUSG00000004591.16 | 1625.558 | 0.125 | 0.040 | 3.122 | 0.001798365 | 0.016390308 | Pkn2 | protein_coding |
| ENSMUSG00000041930.7 | 460.537 | -0.203 | 0.065 | -3.122 | 0.00179905 | 0.016390308 | Fam222a | protein_coding |
| ENSMUSG00000024908.14 | 4768.656 | 0.092 | 0.029 | 3.120 | 0.001805472 | 0.016438195 | Ppp6r3 | protein_coding |
| ENSMUSG00000006651.8 | 57322.095 | -0.112 | 0.036 | -3.120 | 0.00181073 | 0.016475424 | Aplp1 | protein_coding |

| | | | | | | | | |
|---|---|---|---|---|---|---|---|---|
| ENSMUSG00000042323.16 | 3024.193 | 0.113 | 0.036 | 3.118 | 0.001817993 | 0.016530835 | Pbrm1 | protein_coding |
| ENSMUSG00000066037.14 | 5069.949 | 0.108 | 0.035 | 3.117 | 0.001824343 | 0.016577878 | Hnrnpr | protein_coding |
| ENSMUSG00000031451.6 | 3746.138 | -0.115 | 0.037 | -3.116 | 0.001831739 | 0.016634363 | Gas6 | protein_coding |
| ENSMUSG00000045136.6 | 4269.495 | -0.136 | 0.044 | -3.116 | 0.001834055 | 0.016635021 | Tubb2b | protein_coding |
| ENSMUSG00000005442.13 | 12064.247 | -0.119 | 0.038 | -3.116 | 0.001834172 | 0.016635021 | Cic | protein_coding |
| ENSMUSG00000039275.9 | 5285.538 | -0.090 | 0.029 | -3.114 | 0.00184317 | 0.016705878 | Foxk2 | protein_coding |
| ENSMUSG00000052296.7 | 10285.219 | -0.104 | 0.033 | -3.113 | 0.001850117 | 0.016753009 | Ppp6r1 | protein_coding |
| ENSMUSG00000028538.12 | 2014.598 | -0.124 | 0.040 | -3.113 | 0.001850747 | 0.016753009 | St3gal3 | protein_coding |
| ENSMUSG00000032030.16 | 3010.361 | 0.118 | 0.038 | 3.113 | 0.001852007 | 0.016753646 | Cul5 | protein_coding |
| ENSMUSG00000028747.10 | 216.931 | -0.227 | 0.073 | -3.112 | 0.001860288 | 0.01681777 | Htr6 | protein_coding |
| ENSMUSG00000034480.18 | 1439.452 | 0.134 | 0.043 | 3.111 | 0.001862143 | 0.016823749 | Diaph2 | protein_coding |
| ENSMUSG00000061306.15 | 3574.673 | -0.105 | 0.034 | -3.111 | 0.001864407 | 0.01683341 | Slc38a10 | protein_coding |
| ENSMUSG00000029344.14 | 823.111 | -0.168 | 0.054 | -3.110 | 0.00186772 | 0.016848737 | Tpst2 | protein_coding |
| ENSMUSG00000031486.15 | 1036.728 | -0.165 | 0.053 | -3.110 | 0.001868495 | 0.016848737 | Adgra2 | protein_coding |
| ENSMUSG00000046711.15 | 1186.150 | -0.145 | 0.047 | -3.109 | 0.001877607 | 0.016920071 | Hmga1 | protein_coding |
| ENSMUSG00000000902.13 | 2814.029 | -0.111 | 0.036 | -3.108 | 0.001881497 | 0.016935965 | Smarcb1 | protein_coding |
| ENSMUSG00000019852.7 | 10251.713 | 0.074 | 0.024 | 3.108 | 0.001882601 | 0.016935965 | Arfgef3 | protein_coding |
| ENSMUSG00000028670.14 | 3152.069 | -0.150 | 0.048 | -3.108 | 0.001882976 | 0.016935965 | Lypla2 | protein_coding |
| ENSMUSG00000017677.11 | 5274.898 | 0.121 | 0.039 | 3.108 | 0.001885523 | 0.016948058 | Wsb1 | protein_coding |
| ENSMUSG00000020463.15 | 2433.737 | 0.133 | 0.043 | 3.107 | 0.001886945 | 0.016950032 | Ppp4r3b | protein_coding |
| ENSMUSG00000034593.16 | 34855.551 | 0.079 | 0.026 | 3.107 | 0.001892328 | 0.016984954 | Myo5a | protein_coding |
| ENSMUSG00000028911.16 | 3089.587 | -0.103 | 0.033 | -3.106 | 0.001894007 | 0.016984954 | Srsf4 | protein_coding |
| ENSMUSG00000032913.13 | 2355.151 | 0.111 | 0.036 | 3.106 | 0.001894448 | 0.016984954 | Lrig2 | protein_coding |
| ENSMUSG00000020866.17 | 9702.261 | -0.140 | 0.045 | -3.106 | 0.00189846 | 0.017010103 | Cacna1g | protein_coding |
| ENSMUSG00000021302.10 | 2796.741 | 0.132 | 0.043 | 3.105 | 0.001904887 | 0.017056844 | Ggps1 | protein_coding |
| ENSMUSG00000025823.9 | 3072.387 | -0.138 | 0.044 | -3.104 | 0.001910209 | 0.017090395 | Pdia4 | protein_coding |
| ENSMUSG00000093930.1 | 13245.928 | 0.103 | 0.033 | 3.104 | 0.001911211 | 0.017090395 | Hmgcs1 | protein_coding |
| ENSMUSG00000046985.11 | 2546.388 | 0.103 | 0.033 | 3.104 | 0.001912271 | 0.017090395 | Tapt1 | protein_coding |
| ENSMUSG00000000355.13 | 1609.907 | 0.132 | 0.042 | 3.103 | 0.001914319 | 0.017097853 | Mcts1 | protein_coding |
| ENSMUSG00000029608.10 | 28222.538 | -0.098 | 0.032 | -3.103 | 0.001916738 | 0.017108615 | Rph3a | protein_coding |
| ENSMUSG00000026883.17 | 7739.318 | -0.124 | 0.040 | -3.100 | 0.001935263 | 0.017252119 | Dab2ip | protein_coding |
| ENSMUSG00000026028.12 | 11084.594 | 0.089 | 0.029 | 3.099 | 0.001940477 | 0.017287667 | Trak2 | protein_coding |
| ENSMUSG00000022054.11 | 7336.520 | -0.160 | 0.051 | -3.098 | 0.001947636 | 0.017340479 | Nefm | protein_coding |

| | | | | | | | | |
|---|---|---|---|---|---|---|---|---|
| ENSMUSG00000042063.11 | 785.545 | 0.153 | 0.050 | 3.098 | 0.001948929 | 0.017341037 | Zfp386 | protein_coding |
| ENSMUSG00000033249.10 | 826.873 | -0.179 | 0.058 | -3.098 | 0.001951264 | 0.017350858 | Hsf4 | protein_coding |
| ENSMUSG00000026594.14 | 2487.217 | 0.110 | 0.035 | 3.095 | 0.001970326 | 0.017496816 | Ralgps2 | protein_coding |
| ENSMUSG00000040414.7 | 1387.129 | -0.126 | 0.041 | -3.095 | 0.001971265 | 0.017496816 | Slc25a28 | protein_coding |
| ENSMUSG00000029406.15 | 18972.160 | -0.120 | 0.039 | -3.094 | 0.001972426 | 0.017496816 | Pitpnm2 | protein_coding |
| ENSMUSG00000032966.14 | 18962.860 | -0.096 | 0.031 | -3.094 | 0.001972644 | 0.017496816 | Fkbp1a | protein_coding |
| ENSMUSG00000070709.12 | 490.840 | 0.196 | 0.063 | 3.093 | 0.001984738 | 0.017591033 | Zfp974 | protein_coding |
| ENSMUSG00000035172.15 | 520.123 | -0.182 | 0.059 | -3.092 | 0.001985762 | 0.017591033 | Plekhh3 | protein_coding |
| ENSMUSG00000034160.13 | 23360.071 | 0.068 | 0.022 | 3.092 | 0.001987704 | 0.017597174 | Ogt | protein_coding |
| ENSMUSG00000019146.2 | 3840.417 | -0.104 | 0.034 | -3.092 | 0.001989165 | 0.017599052 | Cacng2 | protein_coding |
| ENSMUSG00000040811.15 | 8066.826 | -0.112 | 0.036 | -3.091 | 0.001997348 | 0.017660366 | Eml2 | protein_coding |
| ENSMUSG00000018068.13 | 1445.517 | 0.129 | 0.042 | 3.088 | 0.002013871 | 0.017795297 | Ints2 | protein_coding |
| ENSMUSG00000025892.16 | 5004.834 | 0.097 | 0.031 | 3.087 | 0.002020053 | 0.01783874 | Gria4 | protein_coding |
| ENSMUSG00000030861.15 | 2932.393 | 0.102 | 0.033 | 3.087 | 0.002024775 | 0.017869245 | Acadsb | protein_coding |
| ENSMUSG00000020650.15 | 2199.452 | 0.119 | 0.039 | 3.086 | 0.002026117 | 0.017869894 | Bcap29 | protein_coding |
| ENSMUSG00000070509.15 | 2119.425 | -0.136 | 0.044 | -3.086 | 0.002030679 | 0.017898926 | Rgma | protein_coding |
| ENSMUSG00000020305.13 | 809.235 | 0.170 | 0.055 | 3.085 | 0.002033927 | 0.017903052 | Asb3 | protein_coding |
| ENSMUSG00000030595.15 | 889.640 | -0.182 | 0.059 | -3.085 | 0.002034614 | 0.017903052 | Nfkbib | protein_coding |
| ENSMUSG00000034037.14 | 771.798 | -0.208 | 0.068 | -3.085 | 0.002034957 | 0.017903052 | Fgd5 | protein_coding |
| ENSMUSG00000030774.13 | 20460.957 | 0.083 | 0.027 | 3.085 | 0.002036561 | 0.017905982 | Pak1 | protein_coding |
| ENSMUSG00000010054.4 | 3373.730 | -0.104 | 0.034 | -3.082 | 0.002058254 | 0.018080479 | Tusc2 | protein_coding |
| ENSMUSG00000027203.15 | 724.529 | -0.173 | 0.056 | -3.082 | 0.002058973 | 0.018080479 | Dut | protein_coding |
| ENSMUSG00000022176.11 | 406.252 | -0.220 | 0.071 | -3.080 | 0.002066791 | 0.018137824 | Rem2 | protein_coding |
| ENSMUSG00000036295.5 | 3100.128 | 0.117 | 0.038 | 3.080 | 0.00207103 | 0.018163718 | Lrrn3 | protein_coding |
| ENSMUSG00000027544.16 | 927.360 | -0.153 | 0.050 | -3.080 | 0.002072687 | 0.018166944 | Nfatc2 | protein_coding |
| ENSMUSG00000041609.16 | 3776.343 | -0.114 | 0.037 | -3.079 | 0.002076616 | 0.018190075 | Bicdl1 | protein_coding |
| ENSMUSG00000037643.14 | 3082.726 | 0.103 | 0.033 | 3.078 | 0.002080861 | 0.018215931 | Prkci | protein_coding |
| ENSMUSG00000023915.4 | 4185.831 | -0.111 | 0.036 | -3.078 | 0.002087118 | 0.018258393 | Tnfrsf21 | protein_coding |
| ENSMUSG00000025060.14 | 5397.162 | 0.092 | 0.030 | 3.077 | 0.002089472 | 0.018258393 | Slk | protein_coding |
| ENSMUSG00000025016.10 | 8262.641 | 0.091 | 0.030 | 3.077 | 0.002089598 | 0.018258393 | Tm9sf3 | protein_coding |
| ENSMUSG00000020811.16 | 4170.770 | -0.117 | 0.038 | -3.076 | 0.002095539 | 0.01829896 | Wscd1 | protein_coding |
| ENSMUSG00000028445.7 | 1571.805 | -0.180 | 0.058 | -3.075 | 0.00210303 | 0.018353009 | Enho | protein_coding |
| ENSMUSG00000027637.3 | 1202.630 | -0.135 | 0.044 | -3.075 | 0.002107344 | 0.018376893 | 1110008F13Rik | protein_coding |

| | | | | | | | | |
|---|---|---|---|---|---|---|---|---|
| ENSMUSG00000001525.10 | 25815.111 | -0.095 | 0.031 | -3.075 | 0.002108375 | 0.018376893 | Tubb5 | protein_coding |
| ENSMUSG00000045333.15 | 947.121 | -0.197 | 0.064 | -3.074 | 0.002110179 | 0.018381251 | Zfp423 | protein_coding |
| ENSMUSG00000040536.15 | 6630.909 | 0.109 | 0.035 | 3.074 | 0.00211253 | 0.01839036 | Necab1 | protein_coding |
| ENSMUSG00000050705.15 | 2191.961 | -0.117 | 0.038 | -3.073 | 0.002117915 | 0.018425862 | 2310061I04Rik | protein_coding |
| ENSMUSG00000020868.15 | 1337.446 | -0.136 | 0.044 | -3.072 | 0.002122833 | 0.018457257 | Xylt2 | protein_coding |
| ENSMUSG00000034656.17 | 9972.353 | -0.128 | 0.042 | -3.072 | 0.002126837 | 0.018471794 | Cacna1a | protein_coding |
| ENSMUSG00000052459.13 | 27929.577 | 0.082 | 0.027 | 3.072 | 0.002128273 | 0.018471794 | Atp6v1a | protein_coding |
| ENSMUSG00000006362.16 | 2195.179 | -0.159 | 0.052 | -3.071 | 0.00213109 | 0.018471794 | Cbfa2t3 | protein_coding |
| ENSMUSG00000036898.16 | 1348.430 | 0.136 | 0.044 | 3.071 | 0.002135882 | 0.018471794 | Zfp157 | protein_coding |
| ENSMUSG00000032875.8 | 11778.126 | -0.116 | 0.038 | -3.071 | 0.002136296 | 0.018471794 | Arhgef17 | protein_coding |
| ENSMUSG00000013878.18 | 1250.670 | 0.144 | 0.047 | 3.071 | 0.002136531 | 0.018471794 | Rnf170 | protein_coding |
| ENSMUSG00000022099.16 | 16684.674 | -0.095 | 0.031 | -3.071 | 0.002136572 | 0.018471794 | Dmtn | protein_coding |
| ENSMUSG00000036957.8 | 1249.380 | -0.171 | 0.056 | -3.070 | 0.002137205 | 0.018471794 | Lrfn3 | protein_coding |
| ENSMUSG00000034684.12 | 574.760 | -0.193 | 0.063 | -3.070 | 0.002138155 | 0.018471794 | Sema3f | protein_coding |
| ENSMUSG00000041629.7 | 1294.135 | -0.151 | 0.049 | -3.070 | 0.002138855 | 0.018471794 | Fam104a | protein_coding |
| ENSMUSG00000022305.12 | 2067.776 | 0.133 | 0.043 | 3.070 | 0.002138922 | 0.018471794 | Lrp12 | protein_coding |
| ENSMUSG00000030410.16 | 6307.371 | -0.106 | 0.035 | -3.070 | 0.002142002 | 0.018487065 | Dmwd | protein_coding |
| ENSMUSG00000034789.7 | 2171.990 | -0.118 | 0.039 | -3.069 | 0.00214441 | 0.018496524 | Rab24 | protein_coding |
| ENSMUSG00000046111.17 | 2048.140 | 0.110 | 0.036 | 3.069 | 0.002147294 | 0.018510067 | Cep295 | protein_coding |
| ENSMUSG00000026275.13 | 5571.158 | 0.089 | 0.029 | 3.067 | 0.002162705 | 0.018631523 | Ppp1r7 | protein_coding |
| ENSMUSG00000035835.14 | 3654.829 | -0.150 | 0.049 | -3.066 | 0.002167509 | 0.018661498 | Plppr3 | protein_coding |
| ENSMUSG00000031176.8 | 5495.761 | 0.100 | 0.033 | 3.065 | 0.00217446 | 0.018707934 | Dynlt3 | protein_coding |
| ENSMUSG00000020440.13 | 3362.578 | -0.142 | 0.046 | -3.065 | 0.002175557 | 0.018707934 | Arf5 | protein_coding |
| ENSMUSG00000003929.10 | 1299.587 | 0.149 | 0.049 | 3.065 | 0.002179312 | 0.018728794 | Zfp81 | protein_coding |
| ENSMUSG00000028444.17 | 3482.437 | -0.119 | 0.039 | -3.064 | 0.002183534 | 0.018753645 | Cntfr | protein_coding |
| ENSMUSG00000050830.17 | 399.358 | -0.201 | 0.066 | -3.063 | 0.002193106 | 0.018824381 | Vwc2 | protein_coding |
| ENSMUSG00000072825.11 | 18244.681 | -0.120 | 0.039 | -3.063 | 0.002194664 | 0.018826292 | Cep170b | protein_coding |
| ENSMUSG00000039789.7 | 1238.902 | 0.134 | 0.044 | 3.061 | 0.002205487 | 0.018907624 | Zfp597 | protein_coding |
| ENSMUSG00000025875.14 | 1751.518 | -0.142 | 0.047 | -3.059 | 0.002221177 | 0.01903056 | Tspan17 | protein_coding |
| ENSMUSG00000003184.15 | 1381.303 | -0.144 | 0.047 | -3.058 | 0.002226506 | 0.019064629 | Irf3 | protein_coding |
| ENSMUSG00000015222.17 | 69761.750 | 0.082 | 0.027 | 3.058 | 0.002230129 | 0.019084056 | Map2 | protein_coding |
| ENSMUSG00000052214.9 | 3306.462 | -0.102 | 0.033 | -3.057 | 0.002233723 | 0.01909272 | Opa3 | protein_coding |
| ENSMUSG00000027030.15 | 3311.872 | 0.100 | 0.033 | 3.057 | 0.002233851 | 0.01909272 | Stk39 | protein_coding |

| | | | | | | | | |
|---|---|---|---|---|---|---|---|---|
| ENSMUSG00000002108.10 | 159.195 | -0.233 | 0.076 | -3.056 | 0.002240778 | 0.019131402 | Nr1h3 | protein_coding |
| ENSMUSG00000021071.16 | 19254.231 | -0.078 | 0.026 | -3.056 | 0.002241092 | 0.019131402 | Trim9 | protein_coding |
| ENSMUSG00000017774.19 | 1355.492 | -0.127 | 0.042 | -3.055 | 0.002248733 | 0.019176227 | Myo1c | protein_coding |
| ENSMUSG00000019471.10 | 5407.596 | -0.095 | 0.031 | -3.055 | 0.002249064 | 0.019176227 | Cdc37 | protein_coding |
| ENSMUSG00000009075.2 | 1417.401 | -0.145 | 0.048 | -3.055 | 0.002252391 | 0.019192984 | Cabp7 | protein_coding |
| ENSMUSG00000072946.12 | 1647.531 | 0.125 | 0.041 | 3.054 | 0.002256223 | 0.019214022 | Ptgr2 | protein_coding |
| ENSMUSG00000030557.17 | 7978.722 | 0.102 | 0.033 | 3.054 | 0.002259022 | 0.019226241 | Mef2a | protein_coding |
| ENSMUSG00000028289.12 | 3497.949 | 0.152 | 0.050 | 3.052 | 0.002270438 | 0.019311744 | Epha7 | protein_coding |
| ENSMUSG00000043964.14 | 645.103 | -0.167 | 0.055 | -3.051 | 0.00227951 | 0.019365527 | Orai3 | protein_coding |
| ENSMUSG00000024163.17 | 32535.428 | -0.084 | 0.028 | -3.051 | 0.002281852 | 0.019371937 | Mapk8ip3 | protein_coding |
| ENSMUSG00000024217.9 | 769.455 | -0.160 | 0.052 | -3.051 | 0.002283013 | 0.019371937 | Snrpc | protein_coding |
| ENSMUSG00000026259.14 | 15333.253 | -0.089 | 0.029 | -3.050 | 0.002284704 | 0.019374616 | Ngef | protein_coding |
| ENSMUSG00000005583.16 | 27198.322 | 0.126 | 0.041 | 3.048 | 0.002304909 | 0.019524923 | Mef2c | protein_coding |
| ENSMUSG00000054693.14 | 4663.605 | 0.102 | 0.034 | 3.048 | 0.002305199 | 0.019524923 | Adam10 | protein_coding |
| ENSMUSG00000027366.12 | 2261.430 | 0.112 | 0.037 | 3.048 | 0.002307203 | 0.019530164 | Sppl2a | protein_coding |
| ENSMUSG00000030604.9 | 675.464 | 0.173 | 0.057 | 3.047 | 0.002310036 | 0.019537894 | Zfp626 | protein_coding |
| ENSMUSG00000039089.15 | 1723.868 | -0.129 | 0.042 | -3.047 | 0.002310889 | 0.019537894 | L3mbtl3 | protein_coding |
| ENSMUSG00000049321.17 | 1326.397 | 0.157 | 0.052 | 3.045 | 0.002330115 | 0.019688633 | Zfp2 | protein_coding |
| ENSMUSG00000045087.8 | 441.256 | -0.225 | 0.074 | -3.041 | 0.002355771 | 0.019892317 | S1pr5 | protein_coding |
| ENSMUSG00000054793.8 | 10867.068 | -0.093 | 0.030 | -3.041 | 0.002357043 | 0.019892317 | Cadm4 | protein_coding |
| ENSMUSG00000032184.5 | 1520.814 | -0.150 | 0.049 | -3.041 | 0.002360453 | 0.019909173 | Lysmd2 | protein_coding |
| ENSMUSG00000022556.9 | 2655.307 | -0.139 | 0.046 | -3.040 | 0.002362894 | 0.019909766 | Hsf1 | protein_coding |
| ENSMUSG00000020590.16 | 4527.788 | 0.089 | 0.029 | 3.040 | 0.002363349 | 0.019909766 | Snx13 | protein_coding |
| ENSMUSG00000035623.14 | 3477.318 | 0.100 | 0.033 | 3.039 | 0.002375135 | 0.019985165 | Rsf1 | protein_coding |
| ENSMUSG00000005802.12 | 3646.402 | 0.128 | 0.042 | 3.039 | 0.002376731 | 0.019986661 | Slc30a4 | protein_coding |
| ENSMUSG00000036093.7 | 3868.096 | 0.090 | 0.030 | 3.038 | 0.002383328 | 0.020030184 | Arl5a | protein_coding |
| ENSMUSG00000029475.17 | 1275.278 | -0.134 | 0.044 | -3.037 | 0.002385877 | 0.020039664 | Kdm2b | protein_coding |
| ENSMUSG00000041168.9 | 4287.412 | -0.093 | 0.031 | -3.037 | 0.002393344 | 0.020090408 | Lonp1 | protein_coding |
| ENSMUSG00000031217.8 | 822.786 | -0.187 | 0.062 | -3.036 | 0.002396866 | 0.020107991 | Efnb1 | protein_coding |
| ENSMUSG00000021476.9 | 10185.776 | -0.072 | 0.024 | -3.036 | 0.002400147 | 0.020123538 | Habp4 | protein_coding |
| ENSMUSG00000028568.16 | 1398.101 | 0.135 | 0.044 | 3.035 | 0.002402325 | 0.020129823 | Btf3l4 | protein_coding |
| ENSMUSG00000001829.17 | 3283.329 | -0.093 | 0.031 | -3.034 | 0.002413163 | 0.020208628 | Clpb | protein_coding |
| ENSMUSG00000028211.11 | 953.452 | 0.147 | 0.048 | 3.033 | 0.002418527 | 0.020241517 | Trp53inp1 | protein_coding |

| | | | | | | | | |
|---|---|---|---|---|---|---|---|---|
| ENSMUSG00000024953.16 | 5501.191 | -0.089 | 0.029 | -3.031 | 0.002433792 | 0.020357193 | Prdx5 | protein_coding |
| ENSMUSG00000022329.14 | 602.845 | 0.165 | 0.055 | 3.031 | 0.002435803 | 0.02036193 | Stk3 | protein_coding |
| ENSMUSG00000019888.16 | 484.961 | 0.205 | 0.068 | 3.031 | 0.002441078 | 0.020393928 | Mgat4c | protein_coding |
| ENSMUSG00000029265.4 | 1839.851 | 0.118 | 0.039 | 3.029 | 0.002452264 | 0.020475239 | Dr1 | protein_coding |
| ENSMUSG00000025579.14 | 23494.737 | -0.140 | 0.046 | -3.028 | 0.002459884 | 0.020520028 | Gaa | protein_coding |
| ENSMUSG00000024947.16 | 3190.785 | -0.110 | 0.036 | -3.028 | 0.00246054 | 0.020520028 | Men1 | protein_coding |
| ENSMUSG00000059005.13 | 7946.507 | 0.083 | 0.027 | 3.028 | 0.002462326 | 0.020522783 | Hnrnpa3 | protein_coding |
| ENSMUSG00000027434.11 | 157.344 | -0.233 | 0.077 | -3.026 | 0.002475159 | 0.02061755 | Nkx2-2 | protein_coding |
| ENSMUSG00000028917.14 | 5411.116 | -0.092 | 0.030 | -3.026 | 0.002477821 | 0.020627532 | Plekhm2 | protein_coding |
| ENSMUSG00000000197.7 | 6805.936 | 0.101 | 0.034 | 3.025 | 0.002482582 | 0.020654968 | Nalcn | protein_coding |
| ENSMUSG00000034998.18 | 612.856 | 0.174 | 0.057 | 3.022 | 0.002507581 | 0.020850317 | Foxn2 | protein_coding |
| ENSMUSG00000050310.8 | 4076.484 | 0.100 | 0.033 | 3.022 | 0.002509601 | 0.020850317 | Rictor | protein_coding |
| ENSMUSG00000029223.12 | 23107.798 | -0.083 | 0.027 | -3.022 | 0.0025105 | 0.020850317 | Uchl1 | protein_coding |
| ENSMUSG00000025511.14 | 379.054 | -0.205 | 0.068 | -3.022 | 0.002512353 | 0.020853416 | Tspan4 | protein_coding |
| ENSMUSG00000018417.14 | 1410.640 | 0.150 | 0.050 | 3.021 | 0.002518354 | 0.020890921 | Myo1b | protein_coding |
| ENSMUSG00000052534.15 | 10634.226 | 0.111 | 0.037 | 3.019 | 0.002538114 | 0.021042462 | Pbx1 | protein_coding |
| ENSMUSG00000034040.16 | 2885.020 | -0.102 | 0.034 | -3.018 | 0.002542304 | 0.021064803 | Wbscr17 | protein_coding |
| ENSMUSG00000021413.9 | 7240.185 | 0.098 | 0.032 | 3.018 | 0.002547481 | 0.021095295 | Prpf4b | protein_coding |
| ENSMUSG00000034936.2 | 861.964 | -0.168 | 0.056 | -3.017 | 0.00254913 | 0.021096554 | Arl4d | protein_coding |
| ENSMUSG00000033152.13 | 6793.555 | -0.116 | 0.039 | -3.014 | 0.002580683 | 0.021345154 | Podxl2 | protein_coding |
| ENSMUSG00000000552.9 | 2916.151 | -0.152 | 0.050 | -3.013 | 0.002583867 | 0.021358957 | Zfp385a | protein_coding |
| ENSMUSG00000052934.14 | 6915.708 | -0.097 | 0.032 | -3.009 | 0.002623033 | 0.02165467 | Fbxo31 | protein_coding |
| ENSMUSG00000068154.5 | 216.802 | -0.226 | 0.075 | -3.009 | 0.00262425 | 0.02165467 | Insm1 | protein_coding |
| ENSMUSG00000078137.1 | 2646.828 | -0.162 | 0.054 | -3.008 | 0.002626718 | 0.02166235 | Ankrd63 | protein_coding |
| ENSMUSG00000017747.13 | 406.008 | -0.199 | 0.066 | -3.008 | 0.002629571 | 0.021671781 | Ghdc | protein_coding |
| ENSMUSG00000015749.12 | 4443.365 | 0.093 | 0.031 | 3.008 | 0.002630936 | 0.021671781 | Anp32e | protein_coding |
| ENSMUSG00000041073.10 | 2830.326 | -0.125 | 0.041 | -3.008 | 0.002633845 | 0.021683069 | Nacad | protein_coding |
| ENSMUSG00000002820.6 | 1386.867 | -0.128 | 0.043 | -3.007 | 0.002636034 | 0.021688416 | Atg4d | protein_coding |
| ENSMUSG00000036097.7 | 4639.637 | 0.092 | 0.031 | 3.007 | 0.002638062 | 0.021692441 | Fam178a | protein_coding |
| ENSMUSG00000003970.8 | 9386.655 | -0.070 | 0.023 | -3.006 | 0.002651398 | 0.021773239 | Rpl8 | protein_coding |
| ENSMUSG00000014426.8 | 2882.504 | -0.097 | 0.032 | -3.005 | 0.002652253 | 0.021773239 | Map3k4 | protein_coding |
| ENSMUSG00000027893.14 | 16834.084 | 0.068 | 0.023 | 3.005 | 0.002652522 | 0.021773239 | Ahcyl1 | protein_coding |
| ENSMUSG00000020589.17 | 12626.732 | 0.100 | 0.033 | 3.005 | 0.002654674 | 0.021778213 | Fam49a | protein_coding |

| | | | | | | | | |
|---|---|---|---|---|---|---|---|---|
| ENSMUSG00000021282.17 | 9602.115 | 0.080 | 0.027 | 3.005 | 0.002659678 | 0.021806578 | Eif5 | protein_coding |
| ENSMUSG00000031872.14 | 785.673 | -0.176 | 0.059 | -3.004 | 0.002666136 | 0.021846814 | Bean1 | protein_coding |
| ENSMUSG00000022865.13 | 2050.309 | 0.120 | 0.040 | 3.003 | 0.002676554 | 0.021908723 | Cxadr | protein_coding |
| ENSMUSG00000004937.6 | 6029.400 | -0.096 | 0.032 | -3.003 | 0.002677614 | 0.021908723 | Sgta | protein_coding |
| ENSMUSG00000028677.19 | 7524.714 | -0.087 | 0.029 | -3.002 | 0.002678355 | 0.021908723 | Rnf220 | protein_coding |
| ENSMUSG00000033096.7 | 2092.395 | 0.106 | 0.035 | 3.002 | 0.002684184 | 0.021930954 | Apmap | protein_coding |
| ENSMUSG00000025423.15 | 3550.861 | 0.107 | 0.036 | 3.000 | 0.002696309 | 0.02201725 | Pias2 | protein_coding |
| ENSMUSG00000027220.2 | 11689.569 | -0.078 | 0.026 | -3.000 | 0.002699792 | 0.022032927 | Syt13 | protein_coding |
| ENSMUSG00000022389.14 | 24485.777 | 0.087 | 0.029 | 3.000 | 0.00270236 | 0.022041122 | Tef | protein_coding |
| ENSMUSG00000037286.15 | 2612.804 | 0.108 | 0.036 | 3.000 | 0.002704011 | 0.022041836 | Stag1 | protein_coding |
| ENSMUSG00000063297.7 | 7726.271 | 0.147 | 0.049 | 2.999 | 0.002707601 | 0.022053298 | Luzp2 | protein_coding |
| ENSMUSG00000024937.14 | 1673.536 | -0.124 | 0.041 | -2.999 | 0.002709736 | 0.022053298 | Ehbp1l1 | protein_coding |
| ENSMUSG00000067942.5 | 955.772 | 0.170 | 0.057 | 2.999 | 0.002710112 | 0.022053298 | Zfp160 | protein_coding |
| ENSMUSG00000034902.17 | 24405.657 | -0.124 | 0.041 | -2.998 | 0.002713668 | 0.022069497 | Pip5k1c | protein_coding |
| ENSMUSG00000035246.16 | 2845.405 | 0.096 | 0.032 | 2.998 | 0.002717922 | 0.022091344 | Pcyt1b | protein_coding |
| ENSMUSG00000026585.13 | 14338.445 | 0.077 | 0.026 | 2.996 | 0.002731199 | 0.022183194 | Kifap3 | protein_coding |
| ENSMUSG00000029392.12 | 2482.236 | -0.127 | 0.042 | -2.996 | 0.00273237 | 0.022183194 | Rilpl1 | protein_coding |
| ENSMUSG00000026782.15 | 10478.011 | 0.092 | 0.031 | 2.995 | 0.002741619 | 0.022245472 | Abi2 | protein_coding |
| ENSMUSG00000026443.3 | 5850.729 | -0.118 | 0.039 | -2.993 | 0.002760328 | 0.022384389 | Lrrn2 | protein_coding |
| ENSMUSG00000030035.14 | 1146.721 | -0.140 | 0.047 | -2.992 | 0.002775717 | 0.02249624 | Wbp1 | protein_coding |
| ENSMUSG00000063052.9 | 1085.639 | 0.142 | 0.047 | 2.990 | 0.002786152 | 0.022563601 | Lrrc40 | protein_coding |
| ENSMUSG00000063605.5 | 54.709 | -0.219 | 0.073 | -2.990 | 0.002786456 | NA | Ccdc102a | protein_coding |
| ENSMUSG00000029404.15 | 1114.516 | -0.144 | 0.048 | -2.990 | 0.002787231 | 0.022563601 | Arl6ip4 | protein_coding |
| ENSMUSG00000029416.17 | 1174.106 | -0.154 | 0.051 | -2.989 | 0.002796548 | 0.022626028 | Slc15a4 | protein_coding |
| ENSMUSG00000064061.13 | 5365.475 | 0.095 | 0.032 | 2.987 | 0.002818654 | 0.0227918 | Dzip3 | protein_coding |
| ENSMUSG00000051391.9 | 54193.219 | -0.074 | 0.025 | -2.985 | 0.002831893 | 0.022873863 | Ywhag | protein_coding |
| ENSMUSG00000034412.13 | 1305.701 | -0.137 | 0.046 | -2.985 | 0.002832048 | 0.022873863 | Tbc1d10a | protein_coding |
| ENSMUSG00000009628.14 | 59.235 | 0.222 | 0.074 | 2.985 | 0.002832051 | NA | Tex15 | protein_coding |
| ENSMUSG00000028541.14 | 3394.699 | -0.119 | 0.040 | -2.984 | 0.002840989 | 0.022932929 | B4galt2 | protein_coding |
| ENSMUSG00000028484.16 | 9914.699 | 0.082 | 0.028 | 2.983 | 0.002854188 | 0.023026285 | Psip1 | protein_coding |
| ENSMUSG00000031647.10 | 2827.961 | 0.126 | 0.042 | 2.983 | 0.002857236 | 0.023037691 | Mfap3l | protein_coding |
| ENSMUSG00000038594.9 | 1293.424 | 0.122 | 0.041 | 2.982 | 0.002867 | 0.023085463 | Cep85l | protein_coding |
| ENSMUSG00000015087.14 | 8396.827 | -0.077 | 0.026 | -2.982 | 0.002867864 | 0.023085463 | Rabl6 | protein_coding |

| | | | | | | | | |
|---|---|---|---|---|---|---|---|---|
| ENSMUSG00000033256.14 | 1188.943 | -0.132 | 0.044 | -2.982 | 0.002868075 | 0.023085463 | Shf | protein_coding |
| ENSMUSG00000051243.14 | 4117.812 | -0.136 | 0.045 | -2.981 | 0.002869952 | 0.023087387 | Islr2 | protein_coding |
| ENSMUSG00000037972.6 | 8061.827 | 0.084 | 0.028 | 2.981 | 0.002875158 | 0.023116071 | Snn | protein_coding |
| ENSMUSG00000026020.9 | 2015.834 | 0.117 | 0.039 | 2.977 | 0.002914499 | 0.023400905 | Nop58 | protein_coding |
| ENSMUSG00000062901.2 | 4053.071 | 0.109 | 0.037 | 2.977 | 0.002914576 | 0.023400905 | Klhl24 | protein_coding |
| ENSMUSG00000014850.15 | 1085.959 | 0.145 | 0.049 | 2.977 | 0.002915566 | 0.023400905 | Msh3 | protein_coding |
| ENSMUSG00000037166.5 | 330.219 | -0.232 | 0.078 | -2.976 | 0.002924018 | 0.023455381 | Ppp1r14a | protein_coding |
| ENSMUSG00000003581.14 | 1540.026 | -0.142 | 0.048 | -2.974 | 0.002940465 | 0.023560982 | Rnf215 | protein_coding |
| ENSMUSG00000002222.14 | 7426.127 | 0.092 | 0.031 | 2.974 | 0.002940526 | 0.023560982 | Rmnd5a | protein_coding |
| ENSMUSG00000063146.11 | 9410.584 | -0.114 | 0.038 | -2.974 | 0.002943478 | 0.023571238 | Clip2 | protein_coding |
| ENSMUSG00000026678.10 | 4103.300 | 0.121 | 0.041 | 2.972 | 0.002956914 | 0.023664692 | Rgs5 | protein_coding |
| ENSMUSG00000021076.6 | 5244.536 | 0.087 | 0.029 | 2.972 | 0.002958506 | 0.023664692 | Actr10 | protein_coding |
| ENSMUSG00000025968.16 | 6733.695 | 0.087 | 0.029 | 2.972 | 0.002960331 | 0.023665854 | Ndufs1 | protein_coding |
| ENSMUSG00000048988.8 | 2006.964 | -0.158 | 0.053 | -2.971 | 0.002967148 | 0.023706908 | Elfn1 | protein_coding |
| ENSMUSG00000008855.17 | 10957.776 | -0.139 | 0.047 | -2.971 | 0.002972506 | 0.023736259 | Hdac5 | protein_coding |
| ENSMUSG00000024268.15 | 26685.449 | -0.089 | 0.030 | -2.968 | 0.002993118 | 0.023873797 | Celf4 | protein_coding |
| ENSMUSG00000031805.17 | 439.281 | -0.222 | 0.075 | -2.967 | 0.003010441 | 0.023998394 | Jak3 | protein_coding |
| ENSMUSG00000038538.17 | 5138.697 | 0.098 | 0.033 | 2.966 | 0.003012865 | 0.024004137 | Ubn2 | protein_coding |
| ENSMUSG00000044807.13 | 2524.240 | 0.117 | 0.040 | 2.966 | 0.003019837 | 0.024046089 | Zfp354c | protein_coding |
| ENSMUSG00000026203.16 | 4424.098 | -0.104 | 0.035 | -2.965 | 0.003025097 | 0.024063972 | Dnajb2 | protein_coding |
| ENSMUSG00000037533.19 | 2966.900 | 0.098 | 0.033 | 2.965 | 0.003025497 | 0.024063972 | Rapgef6 | protein_coding |
| ENSMUSG00000044667.12 | 11409.375 | 0.085 | 0.029 | 2.965 | 0.003027374 | 0.024065322 | Plppr4 | protein_coding |
| ENSMUSG00000013419.7 | 4639.140 | -0.114 | 0.039 | -2.965 | 0.00303076 | 0.024078657 | Zfp651 | protein_coding |
| ENSMUSG00000024759.13 | 2211.379 | 0.137 | 0.046 | 2.964 | 0.003035529 | 0.024102952 | Atl3 | protein_coding |
| ENSMUSG00000006456.10 | 1367.546 | -0.140 | 0.047 | -2.963 | 0.003048495 | 0.02419228 | Rbm14 | protein_coding |
| ENSMUSG00000043259.15 | 4062.159 | 0.097 | 0.033 | 2.961 | 0.003062487 | 0.024289641 | Fam13c | protein_coding |
| ENSMUSG00000039233.12 | 3388.504 | 0.105 | 0.036 | 2.961 | 0.003066807 | 0.024307159 | Tbce | protein_coding |
| ENSMUSG00000038805.10 | 272.478 | -0.231 | 0.078 | -2.961 | 0.003069018 | 0.024307159 | Six3 | protein_coding |
| ENSMUSG00000030751.18 | 3189.382 | 0.092 | 0.031 | 2.961 | 0.00306987 | 0.024307159 | Psma1 | protein_coding |
| ENSMUSG00000039000.8 | 4877.043 | 0.088 | 0.030 | 2.959 | 0.003081999 | 0.024378678 | Ube3c | protein_coding |
| ENSMUSG00000071302.10 | 939.612 | 0.167 | 0.056 | 2.959 | 0.003082362 | 0.024378678 | 2610044O15Rik8 | protein_coding |
| ENSMUSG00000030757.13 | 1787.015 | 0.130 | 0.044 | 2.959 | 0.003085593 | 0.024390545 | Zkscan2 | protein_coding |
| ENSMUSG00000062627.9 | 2165.648 | 0.121 | 0.041 | 2.958 | 0.003092187 | 0.024428972 | Mysm1 | protein_coding |

| | | | | | | | | |
|---|---|---|---|---|---|---|---|---|
| ENSMUSG00000031706.7 | 2777.073 | -0.145 | 0.049 | -2.958 | 0.003096244 | 0.024447314 | Rfx1 | protein_coding |
| ENSMUSG00000020923.17 | 6791.025 | -0.073 | 0.025 | -2.957 | 0.003106394 | 0.024513727 | Ubtf | protein_coding |
| ENSMUSG00000096606.2 | 177.647 | -0.227 | 0.077 | -2.956 | 0.003114249 | 0.024561963 | Tpbgl | protein_coding |
| ENSMUSG00000020719.14 | 44186.496 | 0.079 | 0.027 | 2.956 | 0.003118123 | 0.024578763 | Ddx5 | protein_coding |
| ENSMUSG00000028207.18 | 3862.112 | 0.102 | 0.035 | 2.956 | 0.003121483 | 0.024591495 | Asph | protein_coding |
| ENSMUSG00000031095.15 | 2138.040 | 0.143 | 0.048 | 2.954 | 0.003133669 | 0.024673703 | Cul4b | protein_coding |
| ENSMUSG00000079157.4 | 3834.770 | -0.132 | 0.045 | -2.954 | 0.003139976 | 0.024709562 | Fam155a | protein_coding |
| ENSMUSG00000002803.14 | 2315.856 | -0.118 | 0.040 | -2.954 | 0.00314173 | 0.024709566 | Btbd6 | protein_coding |
| ENSMUSG00000027580.17 | 627.529 | -0.179 | 0.061 | -2.953 | 0.003151999 | 0.024776509 | Helz2 | protein_coding |
| ENSMUSG00000018661.17 | 3447.199 | -0.087 | 0.029 | -2.952 | 0.003155231 | 0.024788086 | Cog1 | protein_coding |
| ENSMUSG00000001260.10 | 1321.826 | 0.155 | 0.052 | 2.952 | 0.003161423 | 0.024818092 | Gabrg1 | protein_coding |
| ENSMUSG00000044709.6 | 731.726 | -0.158 | 0.053 | -2.951 | 0.003162572 | 0.024818092 | Gemin7 | protein_coding |
| ENSMUSG00000015363.13 | 1101.142 | -0.137 | 0.047 | -2.951 | 0.003170458 | 0.024853316 | Trabd | protein_coding |
| ENSMUSG00000074227.12 | 596.567 | -0.218 | 0.074 | -2.951 | 0.003170588 | 0.024853316 | Spint2 | protein_coding |
| ENSMUSG00000040021.13 | 2316.840 | 0.100 | 0.034 | 2.949 | 0.003182893 | 0.024935906 | Lats1 | protein_coding |
| ENSMUSG00000032977.9 | 1999.707 | -0.132 | 0.045 | -2.949 | 0.003188428 | 0.024965388 | Fam207a | protein_coding |
| ENSMUSG00000027457.15 | 22056.889 | -0.100 | 0.034 | -2.948 | 0.00319418 | 0.024996544 | Snph | protein_coding |
| ENSMUSG00000019699.16 | 8256.218 | 0.073 | 0.025 | 2.947 | 0.003206156 | 0.02507634 | Akt3 | protein_coding |
| ENSMUSG00000010529.6 | 102.059 | -0.230 | 0.078 | -2.947 | 0.003208969 | 0.025084418 | Gm266 | protein_coding |
| ENSMUSG00000044894.14 | 2396.720 | -0.135 | 0.046 | -2.945 | 0.003229642 | 0.025232023 | Uqcrq | protein_coding |
| ENSMUSG00000021831.8 | 2363.176 | 0.101 | 0.034 | 2.945 | 0.003231738 | 0.025234414 | Ero1l | protein_coding |
| ENSMUSG00000026885.13 | 1480.602 | -0.119 | 0.040 | -2.944 | 0.003243182 | 0.025302111 | Ttll11 | protein_coding |
| ENSMUSG00000037674.15 | 2769.055 | 0.106 | 0.036 | 2.944 | 0.003243998 | 0.025302111 | Rfx7 | protein_coding |
| ENSMUSG00000070934.6 | 3967.576 | -0.097 | 0.033 | -2.943 | 0.003254193 | 0.025367587 | Rraga | protein_coding |
| ENSMUSG00000030854.17 | 14938.124 | -0.114 | 0.039 | -2.940 | 0.00328247 | 0.02557387 | Ptpn5 | protein_coding |
| ENSMUSG00000033726.8 | 353.572 | -0.203 | 0.069 | -2.940 | 0.003285065 | 0.02557995 | Emx1 | protein_coding |
| ENSMUSG00000035696.15 | 2944.225 | 0.121 | 0.041 | 2.938 | 0.003304227 | 0.02571495 | Rnf38 | protein_coding |
| ENSMUSG00000004895.9 | 1761.275 | -0.118 | 0.040 | -2.934 | 0.0033416 | 0.02599145 | Prcc | protein_coding |
| ENSMUSG00000048490.13 | 2233.891 | 0.136 | 0.047 | 2.934 | 0.003346541 | 0.026013897 | Nrip1 | protein_coding |
| ENSMUSG00000029547.10 | 6976.670 | -0.114 | 0.039 | -2.934 | 0.003348177 | 0.026013897 | Ints1 | protein_coding |
| ENSMUSG00000061111.8 | 2330.486 | -0.123 | 0.042 | -2.933 | 0.003354452 | 0.026048289 | Fam195b | protein_coding |
| ENSMUSG00000042256.4 | 203.995 | 0.218 | 0.074 | 2.932 | 0.003368861 | 0.026145773 | Ptchd4 | protein_coding |
| ENSMUSG00000029676.15 | 1022.906 | 0.148 | 0.050 | 2.931 | 0.003374558 | 0.026175576 | Pot1a | protein_coding |

| | | | | | | | | |
|---|---|---|---|---|---|---|---|---|
| ENSMUSG00000070823.6 | 2389.175 | -0.111 | 0.038 | -2.930 | 0.003387976 | 0.026258778 | Gm1043 | protein_coding |
| ENSMUSG00000009092.5 | 103.062 | -0.227 | 0.078 | -2.930 | 0.003389011 | 0.026258778 | Derl3 | protein_coding |
| ENSMUSG00000039715.8 | 1641.974 | -0.141 | 0.048 | -2.929 | 0.003399997 | 0.026329427 | Wdr34 | protein_coding |
| ENSMUSG00000022434.7 | 926.459 | 0.143 | 0.049 | 2.928 | 0.003411668 | 0.026405297 | Fam118a | protein_coding |
| ENSMUSG00000022108.7 | 20290.390 | -0.077 | 0.026 | -2.928 | 0.003414487 | 0.026412612 | Itm2b | protein_coding |
| ENSMUSG00000005779.12 | 4176.240 | -0.092 | 0.032 | -2.927 | 0.003417663 | 0.026422679 | Psmb4 | protein_coding |
| ENSMUSG00000017754.13 | 2463.737 | -0.110 | 0.038 | -2.927 | 0.003422757 | 0.026447555 | Pltp | protein_coding |
| ENSMUSG00000040675.17 | 800.901 | 0.158 | 0.054 | 2.925 | 0.003441569 | 0.026568021 | Mthfd1l | protein_coding |
| ENSMUSG00000009681.10 | 9052.838 | -0.095 | 0.032 | -2.925 | 0.003442118 | 0.026568021 | Bcr | protein_coding |
| ENSMUSG00000002111.8 | 114.203 | -0.227 | 0.077 | -2.925 | 0.003448423 | 0.026596511 | Spi1 | protein_coding |
| ENSMUSG00000029726.16 | 1730.369 | -0.122 | 0.042 | -2.925 | 0.003449583 | 0.026596511 | Mepce | protein_coding |
| ENSMUSG00000027981.14 | 1764.300 | 0.137 | 0.047 | 2.924 | 0.003453679 | 0.026613532 | Rnpc3 | protein_coding |
| ENSMUSG00000027312.14 | 13771.411 | 0.064 | 0.022 | 2.924 | 0.003458092 | 0.02663298 | Atrn | protein_coding |
| ENSMUSG00000029998.14 | 5836.564 | 0.081 | 0.028 | 2.923 | 0.003461463 | 0.026644382 | Pcyox1 | protein_coding |
| ENSMUSG00000036377.18 | 4071.097 | -0.121 | 0.041 | -2.923 | 0.003463684 | 0.026646924 | C530008M17Rik | protein_coding |
| ENSMUSG00000029152.13 | 12479.510 | 0.067 | 0.023 | 2.921 | 0.003484314 | 0.026791011 | Ociad1 | protein_coding |
| ENSMUSG00000029095.17 | 8259.541 | -0.081 | 0.028 | -2.921 | 0.003493682 | 0.026848393 | Ablim2 | protein_coding |
| ENSMUSG00000007613.15 | 1273.818 | 0.148 | 0.051 | 2.920 | 0.003503177 | 0.026906689 | Tgfbr1 | protein_coding |
| ENSMUSG00000021730.8 | 4343.334 | 0.085 | 0.029 | 2.919 | 0.003513564 | 0.026971772 | Hcn1 | protein_coding |
| ENSMUSG00000002812.4 | 4380.215 | -0.084 | 0.029 | -2.918 | 0.003521682 | 0.027019371 | Flii | protein_coding |
| ENSMUSG00000046058.7 | 920.375 | -0.182 | 0.063 | -2.917 | 0.003531381 | 0.027061196 | Eid2 | protein_coding |
| ENSMUSG00000022124.14 | 4356.387 | 0.094 | 0.032 | 2.917 | 0.003532238 | 0.027061196 | Fbxl3 | protein_coding |
| ENSMUSG00000050628.6 | 292.103 | -0.199 | 0.068 | -2.917 | 0.003532893 | 0.027061196 | Ubald2 | protein_coding |
| ENSMUSG00000048787.13 | 1092.244 | 0.129 | 0.044 | 2.917 | 0.003537703 | 0.027083315 | Dcun1d3 | protein_coding |
| ENSMUSG00000040502.5 | 755.224 | -0.159 | 0.054 | -2.916 | 0.003544328 | 0.027119303 | March9 | protein_coding |
| ENSMUSG00000028645.11 | 4542.588 | -0.121 | 0.042 | -2.916 | 0.00354739 | 0.027128005 | Slc2a1 | protein_coding |
| ENSMUSG00000027546.15 | 19938.446 | -0.084 | 0.029 | -2.914 | 0.003573217 | 0.027310697 | Atp9a | protein_coding |
| ENSMUSG00000024293.15 | 1036.715 | 0.131 | 0.045 | 2.913 | 0.003583334 | 0.027373174 | Esco1 | protein_coding |
| ENSMUSG00000022295.7 | 11430.140 | 0.073 | 0.025 | 2.912 | 0.003587046 | 0.027386687 | Atp6v1c1 | protein_coding |
| ENSMUSG00000039100.9 | 14363.913 | 0.080 | 0.027 | 2.911 | 0.003597267 | 0.027449851 | March6 | protein_coding |
| ENSMUSG00000017943.15 | 3486.246 | -0.096 | 0.033 | -2.911 | 0.003602073 | 0.027471655 | Gdap1l1 | protein_coding |
| ENSMUSG00000022562.14 | 1567.172 | -0.135 | 0.046 | -2.911 | 0.003606678 | 0.02748331 | Oplah | protein_coding |
| ENSMUSG00000047669.8 | 429.224 | -0.196 | 0.067 | -2.911 | 0.003607501 | 0.02748331 | Msl3l2 | protein_coding |

| | | | | | | | | |
|---|---|---|---|---|---|---|---|---|
| ENSMUSG00000061458.9 | 1047.613 | 0.143 | 0.049 | 2.909 | 0.003623654 | 0.027591456 | Nol10 | protein_coding |
| ENSMUSG00000024862.16 | 11500.407 | -0.099 | 0.034 | -2.909 | 0.003625651 | 0.027591752 | Klc2 | protein_coding |
| ENSMUSG00000005069.12 | 3618.804 | -0.098 | 0.034 | -2.909 | 0.003628131 | 0.027595722 | Pex5 | protein_coding |
| ENSMUSG00000037003.15 | 1247.349 | -0.156 | 0.054 | -2.906 | 0.003656008 | 0.027792759 | Tns2 | protein_coding |
| ENSMUSG00000021754.17 | 586.429 | 0.209 | 0.072 | 2.905 | 0.003673792 | 0.027912895 | Map3k1 | protein_coding |
| ENSMUSG00000030541.16 | 2148.486 | -0.121 | 0.042 | -2.902 | 0.003712711 | 0.02818808 | Idh2 | protein_coding |
| ENSMUSG00000059173.19 | 12566.060 | 0.108 | 0.037 | 2.901 | 0.003716475 | 0.02818808 | Pde1a | protein_coding |
| ENSMUSG00000022961.17 | 27423.576 | 0.057 | 0.020 | 2.901 | 0.003717555 | 0.02818808 | Son | protein_coding |
| ENSMUSG00000038812.16 | 517.784 | -0.166 | 0.057 | -2.901 | 0.003718011 | 0.02818808 | Trmt112 | protein_coding |
| ENSMUSG00000015882.17 | 1012.881 | 0.138 | 0.047 | 2.900 | 0.003733103 | 0.028287285 | Lcorl | protein_coding |
| ENSMUSG00000055493.4 | 1579.358 | 0.136 | 0.047 | 2.899 | 0.003740647 | 0.028329217 | Epm2a | protein_coding |
| ENSMUSG00000026004.15 | 692.207 | 0.178 | 0.061 | 2.899 | 0.003748208 | 0.02837124 | Kansl1l | protein_coding |
| ENSMUSG00000007021.4 | 8637.358 | -0.092 | 0.032 | -2.898 | 0.003750662 | 0.028374571 | Syngr3 | protein_coding |
| ENSMUSG00000036641.8 | 798.561 | 0.148 | 0.051 | 2.897 | 0.003771061 | 0.028513593 | Ccdc148 | protein_coding |
| ENSMUSG00000002083.13 | 249.253 | -0.209 | 0.072 | -2.896 | 0.003779588 | 0.028562738 | Bbc3 | protein_coding |
| ENSMUSG00000021024.14 | 3100.554 | 0.100 | 0.035 | 2.893 | 0.003813635 | 0.028804596 | Psma6 | protein_coding |
| ENSMUSG00000074829.10 | 833.306 | 0.159 | 0.055 | 2.893 | 0.003816824 | 0.02881324 | 2010315B03Rik | protein_coding |
| ENSMUSG00000029283.17 | 1393.085 | 0.117 | 0.040 | 2.892 | 0.003825653 | 0.028863807 | Cdc7 | protein_coding |
| ENSMUSG00000040940.18 | 4651.912 | -0.115 | 0.040 | -2.892 | 0.003830462 | 0.028863807 | Arhgef1 | protein_coding |
| ENSMUSG00000000561.14 | 1490.969 | 0.126 | 0.043 | 2.892 | 0.003831652 | 0.028863807 | Wdr77 | protein_coding |
| ENSMUSG00000020538.15 | 3082.030 | -0.138 | 0.048 | -2.892 | 0.003831714 | 0.028863807 | Srebf1 | protein_coding |
| ENSMUSG00000029723.16 | 2949.047 | -0.109 | 0.038 | -2.890 | 0.003846367 | 0.028958708 | Tsc22d4 | protein_coding |
| ENSMUSG00000029462.18 | 4216.778 | 0.097 | 0.034 | 2.889 | 0.003869613 | 0.029118169 | Vps29 | protein_coding |
| ENSMUSG00000091956.2 | 32.221 | -0.179 | 0.062 | -2.889 | 0.003869925 | NA | C2cd4b | protein_coding |
| ENSMUSG00000039163.10 | 310.748 | 0.194 | 0.067 | 2.888 | 0.003873526 | 0.029132059 | Cmc1 | protein_coding |
| ENSMUSG00000020986.13 | 5605.758 | 0.090 | 0.031 | 2.887 | 0.003892832 | 0.029261638 | Sec23a | protein_coding |
| ENSMUSG00000024220.13 | 5710.187 | -0.107 | 0.037 | -2.886 | 0.003898454 | 0.029288282 | Zfp523 | protein_coding |
| ENSMUSG00000025266.11 | 11716.046 | 0.080 | 0.028 | 2.884 | 0.003920773 | 0.029440261 | Gnl3l | protein_coding |
| ENSMUSG00000030960.16 | 664.343 | 0.155 | 0.054 | 2.882 | 0.003947563 | 0.029625642 | Mettl10 | protein_coding |
| ENSMUSG00000031813.8 | 775.359 | -0.161 | 0.056 | -2.882 | 0.003949961 | 0.029627858 | Mvb12a | protein_coding |
| ENSMUSG00000018677.9 | 2573.863 | -0.142 | 0.049 | -2.881 | 0.003961619 | 0.029699499 | Slc25a39 | protein_coding |
| ENSMUSG00000061524.8 | 854.621 | -0.209 | 0.072 | -2.879 | 0.003988925 | 0.029866952 | Zic2 | protein_coding |
| ENSMUSG00000038039.13 | 4280.819 | 0.087 | 0.030 | 2.879 | 0.003989146 | 0.029866952 | Gcc2 | protein_coding |

| | | | | | | | | |
|---|---|---|---|---|---|---|---|---|
| ENSMUSG00000024807.17 | 3449.735 | -0.114 | 0.040 | -2.879 | 0.003990313 | 0.029866952 | Syvn1 | protein_coding |
| ENSMUSG00000020358.17 | 4222.207 | -0.111 | 0.038 | -2.879 | 0.003994706 | 0.029883964 | Hnrnpab | protein_coding |
| ENSMUSG00000041570.14 | 9699.306 | 0.071 | 0.025 | 2.878 | 0.003998301 | 0.02989499 | Camsap2 | protein_coding |
| ENSMUSG00000029381.15 | 273.473 | -0.219 | 0.076 | -2.878 | 0.004001258 | 0.029901232 | Shroom3 | protein_coding |
| ENSMUSG00000022559.7 | 1359.811 | -0.131 | 0.045 | -2.877 | 0.004017849 | 0.030009306 | Fbxl6 | protein_coding |
| ENSMUSG00000055471.6 | 437.764 | -0.213 | 0.074 | -2.875 | 0.004039517 | 0.030155167 | Alk | protein_coding |
| ENSMUSG00000027274.16 | 1701.579 | 0.133 | 0.046 | 2.873 | 0.004060939 | 0.030299032 | Mkks | protein_coding |
| ENSMUSG00000027122.15 | 1751.407 | 0.109 | 0.038 | 2.873 | 0.004066632 | 0.030325454 | Arl14ep | protein_coding |
| ENSMUSG00000020090.7 | 228.748 | -0.224 | 0.078 | -2.872 | 0.004079794 | 0.03039211 | Npffr1 | protein_coding |
| ENSMUSG00000050796.4 | 483.713 | -0.178 | 0.062 | -2.872 | 0.004082039 | 0.03039211 | B3galt6 | protein_coding |
| ENSMUSG00000026219.14 | 12298.936 | 0.071 | 0.025 | 2.871 | 0.004093044 | 0.030457953 | Trip12 | protein_coding |
| ENSMUSG00000095224.1 | 33.520 | -0.187 | 0.065 | -2.870 | 0.004099328 | NA | F930015N05Rik | protein_coding |
| ENSMUSG00000045896.14 | 4282.507 | 0.101 | 0.035 | 2.870 | 0.004101729 | 0.030506471 | Paip2b | protein_coding |
| ENSMUSG00000073295.4 | 1365.631 | 0.121 | 0.042 | 2.870 | 0.004108414 | 0.030540078 | Nudt11 | protein_coding |
| ENSMUSG00000051951.5 | 2172.903 | 0.105 | 0.036 | 2.868 | 0.004126198 | 0.030656103 | Xkr4 | protein_coding |
| ENSMUSG00000030284.11 | 4354.857 | -0.099 | 0.034 | -2.866 | 0.004156924 | 0.030868116 | Creld1 | protein_coding |
| ENSMUSG00000055733.6 | 2486.578 | 0.106 | 0.037 | 2.865 | 0.00417592 | 0.03099215 | Nap1l3 | protein_coding |
| ENSMUSG00000041528.15 | 4775.163 | -0.086 | 0.030 | -2.864 | 0.004178273 | 0.03099215 | Rnf123 | protein_coding |
| ENSMUSG00000018809.2 | 683.226 | 0.159 | 0.056 | 2.864 | 0.004180726 | 0.03099215 | Smyd4 | protein_coding |
| ENSMUSG00000074102.4 | 3178.611 | -0.097 | 0.034 | -2.864 | 0.004182423 | 0.03099215 | Rbm15b | protein_coding |
| ENSMUSG00000007659.18 | 5113.133 | -0.080 | 0.028 | -2.864 | 0.004186887 | 0.030993083 | Bcl2l1 | protein_coding |
| ENSMUSG00000047412.16 | 1323.609 | 0.148 | 0.052 | 2.864 | 0.004188874 | 0.030993083 | Zbtb44 | protein_coding |
| ENSMUSG00000053799.6 | 1417.664 | 0.135 | 0.047 | 2.864 | 0.004189146 | 0.030993083 | Exoc6 | protein_coding |
| ENSMUSG00000039375.16 | 1969.057 | 0.141 | 0.049 | 2.861 | 0.004220222 | 0.031198206 | Wdr17 | protein_coding |
| ENSMUSG00000027550.14 | 1081.901 | 0.130 | 0.045 | 2.861 | 0.004222124 | 0.031198206 | Lrrcc1 | protein_coding |
| ENSMUSG00000036908.16 | 699.321 | -0.175 | 0.061 | -2.861 | 0.004225796 | 0.031198206 | Unc93b1 | protein_coding |
| ENSMUSG00000061589.14 | 7852.959 | -0.120 | 0.042 | -2.861 | 0.004225844 | 0.031198206 | Dot1l | protein_coding |
| ENSMUSG00000056073.16 | 4267.930 | 0.112 | 0.039 | 2.861 | 0.004227939 | 0.031198206 | Grik2 | protein_coding |
| ENSMUSG00000059981.12 | 9912.784 | -0.099 | 0.035 | -2.859 | 0.004256135 | 0.031387988 | Taok2 | protein_coding |
| ENSMUSG00000020102.15 | 2436.958 | 0.126 | 0.044 | 2.858 | 0.004258112 | 0.031387988 | Slc16a7 | protein_coding |
| ENSMUSG00000024943.8 | 1749.443 | 0.122 | 0.043 | 2.858 | 0.004262882 | 0.03140672 | Smc5 | protein_coding |
| ENSMUSG00000027335.9 | 1020.145 | -0.223 | 0.078 | -2.857 | 0.004282678 | 0.031536084 | Adra1d | protein_coding |
| ENSMUSG00000039081.9 | 149.357 | -0.221 | 0.077 | -2.856 | 0.004288181 | 0.031560116 | Zfp503 | protein_coding |

| | | | | | | | | |
|---|---|---|---|---|---|---|---|---|
| ENSMUSG00000029213.11 | 1603.446 | 0.119 | 0.042 | 2.855 | 0.004309594 | 0.031686385 | Commd8 | protein_coding |
| ENSMUSG00000038733.13 | 7341.153 | 0.068 | 0.024 | 2.855 | 0.004309834 | 0.031686385 | Wdr26 | protein_coding |
| ENSMUSG00000031917.12 | 990.362 | 0.140 | 0.049 | 2.854 | 0.00431307 | 0.031693643 | Nip7 | protein_coding |
| ENSMUSG00000067212.8 | 872.700 | -0.188 | 0.066 | -2.854 | 0.004318576 | 0.031717568 | H2-T23 | protein_coding |
| ENSMUSG00000031776.17 | 5533.946 | -0.084 | 0.029 | -2.854 | 0.004323844 | 0.031739714 | Arl2bp | protein_coding |
| ENSMUSG00000081058.5 | 47.844 | -0.207 | 0.073 | -2.852 | 0.004342478 | NA | Hist2h3c2 | protein_coding |
| ENSMUSG00000042203.7 | 2451.620 | -0.107 | 0.037 | -2.852 | 0.004343863 | 0.03187007 | Tbc1d22b | protein_coding |
| ENSMUSG00000025958.14 | 3011.460 | 0.096 | 0.034 | 2.851 | 0.00435359 | 0.031924816 | Creb1 | protein_coding |
| ENSMUSG00000039110.14 | 195.719 | -0.220 | 0.077 | -2.850 | 0.004365995 | 0.031999137 | Mycbpap | protein_coding |
| ENSMUSG00000012640.16 | 1783.452 | 0.109 | 0.038 | 2.850 | 0.004370688 | 0.032016877 | Zfp715 | protein_coding |
| ENSMUSG00000024122.16 | 8309.132 | 0.074 | 0.026 | 2.849 | 0.004383893 | 0.032096926 | Pdpk1 | protein_coding |
| ENSMUSG00000038415.10 | 133.771 | -0.221 | 0.078 | -2.849 | 0.004392286 | 0.032141681 | Foxq1 | protein_coding |
| ENSMUSG00000000058.6 | 1457.141 | 0.134 | 0.047 | 2.847 | 0.004411064 | 0.032262345 | Cav2 | protein_coding |
| ENSMUSG00000062190.12 | 4117.026 | 0.100 | 0.035 | 2.847 | 0.00441492 | 0.032273801 | Lancl2 | protein_coding |
| ENSMUSG00000045980.13 | 1095.528 | -0.154 | 0.054 | -2.846 | 0.004420507 | 0.03229789 | Tmem104 | protein_coding |
| ENSMUSG00000026623.16 | 16016.804 | 0.074 | 0.026 | 2.845 | 0.004444631 | 0.032457323 | Lpgat1 | protein_coding |
| ENSMUSG00000003438.17 | 1207.996 | -0.121 | 0.043 | -2.843 | 0.004464717 | 0.03258712 | Timm50 | protein_coding |
| ENSMUSG00000034152.6 | 5909.334 | 0.090 | 0.031 | 2.843 | 0.004474989 | 0.032638421 | Exoc3 | protein_coding |
| ENSMUSG00000037818.16 | 1205.024 | 0.137 | 0.048 | 2.842 | 0.004476378 | 0.032638421 | Abhd18 | protein_coding |
| ENSMUSG00000033216.9 | 783.972 | -0.150 | 0.053 | -2.841 | 0.004492265 | 0.03273732 | Eefsec | protein_coding |
| ENSMUSG00000025130.11 | 5505.445 | -0.080 | 0.028 | -2.840 | 0.004507173 | 0.032828994 | P4hb | protein_coding |
| ENSMUSG00000024851.13 | 12086.263 | -0.117 | 0.041 | -2.837 | 0.004554349 | 0.033138355 | Pitpnm1 | protein_coding |
| ENSMUSG00000019539.11 | 167.063 | -0.218 | 0.077 | -2.837 | 0.004559046 | 0.033155412 | Rcn3 | protein_coding |
| ENSMUSG00000029408.13 | 2561.511 | -0.128 | 0.045 | -2.836 | 0.00456438 | 0.033177085 | Abcb9 | protein_coding |
| ENSMUSG00000016757.10 | 930.409 | -0.136 | 0.048 | -2.835 | 0.004576449 | 0.033247667 | Ttll12 | protein_coding |
| ENSMUSG00000037364.12 | 7777.017 | -0.076 | 0.027 | -2.835 | 0.004580435 | 0.033259483 | Srrt | protein_coding |
| ENSMUSG00000027581.12 | 14149.775 | -0.080 | 0.028 | -2.834 | 0.004594615 | 0.03332125 | Stmn3 | protein_coding |
| ENSMUSG00000038366.15 | 9096.112 | -0.087 | 0.031 | -2.834 | 0.004595448 | 0.03332125 | Lasp1 | protein_coding |
| ENSMUSG00000029245.16 | 6123.933 | 0.112 | 0.040 | 2.834 | 0.004596034 | 0.03332125 | Epha5 | protein_coding |
| ENSMUSG00000014846.12 | 2026.408 | -0.143 | 0.050 | -2.833 | 0.00460418 | 0.033333759 | Tppp3 | protein_coding |
| ENSMUSG00000056258.8 | 9482.170 | 0.103 | 0.036 | 2.833 | 0.004604787 | 0.033333759 | Kcnq3 | protein_coding |
| ENSMUSG00000017286.15 | 2324.834 | 0.104 | 0.037 | 2.833 | 0.004604855 | 0.033333759 | Glod4 | protein_coding |
| ENSMUSG00000027248.13 | 9807.582 | -0.108 | 0.038 | -2.832 | 0.004619637 | 0.033423596 | Pdia3 | protein_coding |

| | | | | | | | | |
|---|---|---|---|---|---|---|---|---|
| ENSMUSG00000034471.12 | 2006.459 | -0.123 | 0.043 | -2.832 | 0.004627907 | 0.033466247 | Caskin2 | protein_coding |
| ENSMUSG00000025425.17 | 2589.963 | -0.103 | 0.036 | -2.829 | 0.004669854 | 0.033752269 | St8sia5 | protein_coding |
| ENSMUSG00000030226.12 | 4956.855 | 0.155 | 0.055 | 2.828 | 0.004677427 | 0.033789673 | Lmo3 | protein_coding |
| ENSMUSG00000007670.9 | 5115.982 | -0.097 | 0.034 | -2.828 | 0.004682195 | 0.03380679 | Khsrp | protein_coding |
| ENSMUSG00000071369.11 | 2636.207 | -0.099 | 0.035 | -2.828 | 0.004685263 | 0.033811622 | Map3k5 | protein_coding |
| ENSMUSG00000020570.15 | 2038.760 | 0.129 | 0.046 | 2.827 | 0.00469803 | 0.033886403 | Sypl | protein_coding |
| ENSMUSG00000042662.16 | 1088.737 | -0.130 | 0.046 | -2.826 | 0.004715355 | 0.03399397 | Dusp15 | protein_coding |
| ENSMUSG00000031634.13 | 1845.088 | 0.120 | 0.042 | 2.825 | 0.004730072 | 0.03407211 | Ufsp2 | protein_coding |
| ENSMUSG00000035403.10 | 464.103 | -0.186 | 0.066 | -2.825 | 0.004731029 | 0.03407211 | Crb2 | protein_coding |
| ENSMUSG00000020591.11 | 4510.472 | -0.084 | 0.030 | -2.825 | 0.004734878 | 0.034082418 | Ntsr2 | protein_coding |
| ENSMUSG00000028033.16 | 4456.268 | 0.103 | 0.036 | 2.824 | 0.004741451 | 0.034099745 | Kcnq5 | protein_coding |
| ENSMUSG00000033671.18 | 4661.247 | 0.079 | 0.028 | 2.824 | 0.004742124 | 0.034099745 | Cep350 | protein_coding |
| ENSMUSG00000055884.8 | 736.958 | 0.146 | 0.052 | 2.822 | 0.004773188 | 0.034305613 | Fancm | protein_coding |
| ENSMUSG00000037376.14 | 1310.073 | -0.130 | 0.046 | -2.822 | 0.004779664 | 0.034332589 | Trmt6 | protein_coding |
| ENSMUSG00000037509.19 | 11931.775 | -0.080 | 0.028 | -2.821 | 0.004783695 | 0.034332589 | Arhgef4 | protein_coding |
| ENSMUSG00000028518.8 | 2769.534 | 0.101 | 0.036 | 2.821 | 0.004784512 | 0.034332589 | Prkaa2 | protein_coding |
| ENSMUSG00000019970.15 | 4033.305 | -0.219 | 0.078 | -2.821 | 0.004786685 | 0.034332589 | Sgk1 | protein_coding |
| ENSMUSG00000091264.2 | 11522.341 | 0.082 | 0.029 | 2.821 | 0.004793677 | 0.034365248 | Smim13 | protein_coding |
| ENSMUSG00000024201.12 | 4733.544 | -0.084 | 0.030 | -2.820 | 0.004801317 | 0.034393205 | Kdm4b | protein_coding |
| ENSMUSG00000042446.16 | 3214.860 | 0.089 | 0.032 | 2.820 | 0.004802457 | 0.034393205 | Zmym4 | protein_coding |
| ENSMUSG00000033728.14 | 926.267 | -0.151 | 0.054 | -2.819 | 0.004814755 | 0.034463769 | Lrrc14 | protein_coding |
| ENSMUSG00000039361.11 | 7728.129 | 0.090 | 0.032 | 2.818 | 0.004834198 | 0.034585377 | Picalm | protein_coding |
| ENSMUSG00000036698.10 | 9195.413 | -0.062 | 0.022 | -2.816 | 0.004861759 | 0.034764904 | ago-02 | protein_coding |
| ENSMUSG00000028397.13 | 1301.415 | 0.138 | 0.049 | 2.815 | 0.004884951 | 0.034900442 | Kdm4c | protein_coding |
| ENSMUSG00000054455.4 | 7009.041 | -0.081 | 0.029 | -2.814 | 0.004885666 | 0.034900442 | Vapb | protein_coding |
| ENSMUSG00000033287.15 | 5947.733 | -0.096 | 0.034 | -2.814 | 0.004894437 | 0.034945388 | Kctd17 | protein_coding |
| ENSMUSG00000000776.12 | 803.949 | -0.154 | 0.055 | -2.814 | 0.004899679 | 0.034965105 | Polr3d | protein_coding |
| ENSMUSG00000021290.6 | 1707.258 | 0.119 | 0.042 | 2.813 | 0.004905902 | 0.03499179 | 2010107E04Rik | protein_coding |
| ENSMUSG00000033430.10 | 2967.685 | -0.095 | 0.034 | -2.812 | 0.004916008 | 0.035046138 | Terf2ip | protein_coding |
| ENSMUSG00000027613.15 | 1607.451 | -0.126 | 0.045 | -2.812 | 0.00492641 | 0.035102537 | Eif6 | protein_coding |
| ENSMUSG00000040813.16 | 2429.520 | -0.105 | 0.037 | -2.811 | 0.00494198 | 0.035178427 | Tex264 | protein_coding |
| ENSMUSG00000025658.16 | 12177.632 | 0.094 | 0.034 | 2.811 | 0.004942052 | 0.035178427 | Cnksr2 | protein_coding |
| ENSMUSG00000042286.13 | 1297.622 | -0.160 | 0.057 | -2.809 | 0.004961971 | 0.035302381 | Stab1 | protein_coding |

| | | | | | | | | |
|---|---|---|---|---|---|---|---|---|
| ENSMUSG00000030528.12 | 229.010 | 0.199 | 0.071 | 2.809 | 0.004968199 | 0.035328857 | Blm | protein_coding |
| ENSMUSG00000048279.17 | 5423.330 | 0.086 | 0.031 | 2.807 | 0.00500054 | 0.035540906 | Sacs | protein_coding |
| ENSMUSG00000020954.16 | 3645.356 | 0.099 | 0.035 | 2.807 | 0.005006569 | 0.035558457 | Strn3 | protein_coding |
| ENSMUSG00000029061.2 | 149.090 | -0.217 | 0.077 | -2.807 | 0.005008056 | 0.035558457 | Mmp23 | protein_coding |
| ENSMUSG00000079084.10 | 1635.208 | 0.141 | 0.050 | 2.806 | 0.005015205 | 0.035591286 | Ccdc82 | protein_coding |
| ENSMUSG00000047446.18 | 1627.763 | 0.120 | 0.043 | 2.804 | 0.005046804 | 0.035790326 | Arl4a | protein_coding |
| ENSMUSG00000018593.13 | 7566.220 | -0.134 | 0.048 | -2.804 | 0.00504907 | 0.035790326 | Sparc | protein_coding |
| ENSMUSG00000002808.7 | 5220.543 | -0.086 | 0.031 | -2.804 | 0.00505087 | 0.035790326 | Epdr1 | protein_coding |
| ENSMUSG00000018474.17 | 40392.474 | -0.081 | 0.029 | -2.803 | 0.005057989 | 0.035813577 | Chd3 | protein_coding |
| ENSMUSG00000031715.14 | 2584.249 | 0.102 | 0.036 | 2.803 | 0.005059983 | 0.035813577 | Smarca5 | protein_coding |
| ENSMUSG00000079317.10 | 1364.598 | 0.126 | 0.045 | 2.803 | 0.005061774 | 0.035813577 | Trappc2 | protein_coding |
| ENSMUSG00000059669.8 | 905.192 | 0.143 | 0.051 | 2.803 | 0.005068909 | 0.03583094 | Taf1b | protein_coding |
| ENSMUSG00000090112.8 | 3283.617 | 0.094 | 0.034 | 2.803 | 0.005069313 | 0.03583094 | Shprh | protein_coding |
| ENSMUSG00000038084.16 | 8417.630 | 0.076 | 0.027 | 2.802 | 0.005076634 | 0.035861985 | Opa1 | protein_coding |
| ENSMUSG00000019817.18 | 1365.174 | 0.130 | 0.047 | 2.802 | 0.005078794 | 0.035861985 | Plagl1 | protein_coding |
| ENSMUSG00000024423.5 | 14629.599 | 0.087 | 0.031 | 2.801 | 0.005096157 | 0.035966571 | Impact | protein_coding |
| ENSMUSG00000001143.13 | 3114.395 | -0.105 | 0.038 | -2.801 | 0.005100714 | 0.035978228 | Lman2l | protein_coding |
| ENSMUSG00000049744.15 | 393.411 | 0.213 | 0.076 | 2.800 | 0.005102915 | 0.035978228 | Arhgap15 | protein_coding |
| ENSMUSG00000040875.12 | 2174.705 | -0.115 | 0.041 | -2.799 | 0.005120472 | 0.036083966 | Osbpl10 | protein_coding |
| ENSMUSG00000036315.13 | 428.333 | 0.170 | 0.061 | 2.799 | 0.005128896 | 0.036125266 | Znrd1 | protein_coding |
| ENSMUSG00000018848.4 | 1883.804 | 0.107 | 0.038 | 2.797 | 0.005151132 | 0.036263764 | Rars | protein_coding |
| ENSMUSG00000040785.18 | 57443.698 | 0.049 | 0.017 | 2.797 | 0.005156703 | 0.036283546 | Ttc3 | protein_coding |
| ENSMUSG00000041891.15 | 1859.862 | 0.107 | 0.038 | 2.797 | 0.005160279 | 0.036283546 | Lman1 | protein_coding |
| ENSMUSG00000079657.10 | 3066.927 | -0.121 | 0.043 | -2.797 | 0.005161665 | 0.036283546 | Rab26 | protein_coding |
| ENSMUSG00000078234.6 | 705.399 | -0.182 | 0.065 | -2.796 | 0.0051724 | 0.036331245 | Klhdc7a | protein_coding |
| ENSMUSG00000036242.14 | 1256.763 | 0.125 | 0.045 | 2.796 | 0.005173606 | 0.036331245 | 3632451O06Rik | protein_coding |
| ENSMUSG00000043843.16 | 1477.358 | -0.160 | 0.057 | -2.796 | 0.005178287 | 0.036346006 | Tmem145 | protein_coding |
| ENSMUSG00000027977.15 | 2952.218 | 0.112 | 0.040 | 2.795 | 0.005191014 | 0.036412328 | Ndst3 | protein_coding |
| ENSMUSG00000001383.8 | 5937.772 | 0.094 | 0.034 | 2.795 | 0.005195378 | 0.036412328 | Zmat2 | protein_coding |
| ENSMUSG00000032097.10 | 8456.316 | 0.081 | 0.029 | 2.794 | 0.005198277 | 0.036412328 | Ddx6 | protein_coding |
| ENSMUSG00000030682.4 | 6057.526 | -0.079 | 0.028 | -2.794 | 0.00520168 | 0.036412328 | Cdipt | protein_coding |
| ENSMUSG00000020814.13 | 293.498 | -0.207 | 0.074 | -2.794 | 0.005203141 | 0.036412328 | Mxra7 | protein_coding |
| ENSMUSG00000028232.13 | 1629.116 | 0.121 | 0.043 | 2.794 | 0.005203237 | 0.036412328 | Tmem68 | protein_coding |

| | | | | | | | | |
|---|---|---|---|---|---|---|---|---|
| ENSMUSG00000006423.15 | 1915.349 | 0.107 | 0.038 | 2.794 | 0.005210308 | 0.036443715 | C330007P06Rik | protein_coding |
| ENSMUSG00000022443.16 | 7353.311 | -0.118 | 0.042 | -2.792 | 0.005230901 | 0.036569602 | Myh9 | protein_coding |
| ENSMUSG00000040396.12 | 1884.509 | 0.102 | 0.037 | 2.791 | 0.005249978 | 0.036684775 | Abhd13 | protein_coding |
| ENSMUSG00000022382.14 | 2139.972 | -0.128 | 0.046 | -2.791 | 0.005257933 | 0.036722157 | Wnt7b | protein_coding |
| ENSMUSG00000015605.5 | 3418.427 | -0.122 | 0.044 | -2.790 | 0.00526776 | 0.036759408 | Srf | protein_coding |
| ENSMUSG00000020078.16 | 2662.592 | 0.124 | 0.044 | 2.790 | 0.005268483 | 0.036759408 | Vps26a | protein_coding |
| ENSMUSG00000033389.16 | 7933.180 | -0.089 | 0.032 | -2.788 | 0.00529651 | 0.03692171 | Arhgap44 | protein_coding |
| ENSMUSG00000016256.10 | 1407.544 | -0.121 | 0.043 | -2.788 | 0.005296984 | 0.03692171 | Ctsz | protein_coding |
| ENSMUSG00000031837.14 | 2895.485 | -0.151 | 0.054 | -2.787 | 0.005316324 | 0.037038196 | Necab2 | protein_coding |
| ENSMUSG00000009633.3 | 144.464 | -0.217 | 0.078 | -2.787 | 0.005322191 | 0.037060144 | G0s2 | protein_coding |
| ENSMUSG00000019055.15 | 1263.282 | -0.163 | 0.059 | -2.787 | 0.005324733 | 0.037060144 | Plod1 | protein_coding |
| ENSMUSG00000032087.10 | 4527.800 | -0.114 | 0.041 | -2.786 | 0.00532979 | 0.03707703 | Dscaml1 | protein_coding |
| ENSMUSG00000024193.7 | 2505.588 | -0.115 | 0.041 | -2.786 | 0.005333565 | 0.037084987 | Phf1 | protein_coding |
| ENSMUSG00000039716.12 | 18783.837 | 0.068 | 0.024 | 2.786 | 0.005338257 | 0.037099307 | Dock3 | protein_coding |
| ENSMUSG00000052915.14 | 9299.328 | -0.066 | 0.024 | -2.785 | 0.005352187 | 0.037177783 | Msl1 | protein_coding |
| ENSMUSG00000020042.15 | 1637.831 | -0.117 | 0.042 | -2.785 | 0.005357703 | 0.037197769 | Btbd11 | protein_coding |
| ENSMUSG00000074698.10 | 3945.323 | 0.082 | 0.029 | 2.781 | 0.005419775 | 0.037610194 | Csnk2a1 | protein_coding |
| ENSMUSG00000040007.8 | 2720.039 | -0.127 | 0.046 | -2.779 | 0.005452442 | 0.037818268 | Bahd1 | protein_coding |
| ENSMUSG00000071252.5 | 483.675 | 0.166 | 0.060 | 2.779 | 0.005456501 | 0.037827805 | 2210408I21Rik | protein_coding |
| ENSMUSG00000042599.8 | 4016.723 | 0.112 | 0.040 | 2.777 | 0.005486038 | 0.038013876 | Kdm7a | protein_coding |
| ENSMUSG00000029860.16 | 2184.588 | -0.105 | 0.038 | -2.776 | 0.005509872 | 0.038160264 | Zyx | protein_coding |
| ENSMUSG00000040520.7 | 1643.982 | 0.125 | 0.045 | 2.775 | 0.005517989 | 0.03819771 | Manea | protein_coding |
| ENSMUSG00000041445.8 | 245.744 | -0.215 | 0.077 | -2.775 | 0.005522707 | 0.0382116 | Mmrn2 | protein_coding |
| ENSMUSG00000020053.18 | 365.508 | 0.191 | 0.069 | 2.775 | 0.00552578 | 0.038214106 | Igf1 | protein_coding |
| ENSMUSG00000052397.8 | 3327.604 | -0.115 | 0.041 | -2.774 | 0.005541827 | 0.038306284 | Ezr | protein_coding |
| ENSMUSG00000041729.15 | 8383.494 | -0.092 | 0.033 | -2.774 | 0.00554493 | 0.03830894 | Coro2b | protein_coding |
| ENSMUSG00000025743.14 | 12097.867 | -0.112 | 0.040 | -2.772 | 0.005576636 | 0.03850912 | Sdc3 | protein_coding |
| ENSMUSG00000029401.8 | 496.165 | -0.171 | 0.062 | -2.770 | 0.00559875 | 0.038634996 | Rilpl2 | protein_coding |
| ENSMUSG00000042404.16 | 3734.127 | -0.093 | 0.033 | -2.770 | 0.005600347 | 0.038634996 | Dennd4b | protein_coding |
| ENSMUSG00000047368.3 | 1283.838 | 0.130 | 0.047 | 2.770 | 0.005609163 | 0.038676883 | Abhd17b | protein_coding |
| ENSMUSG00000073725.8 | 9290.077 | 0.072 | 0.026 | 2.769 | 0.005617691 | 0.038716744 | Lmbrd1 | protein_coding |
| ENSMUSG00000021774.12 | 2554.221 | 0.100 | 0.036 | 2.769 | 0.005628913 | 0.03877512 | Ube2e1 | protein_coding |
| ENSMUSG00000063260.2 | 302.870 | 0.197 | 0.071 | 2.768 | 0.005633208 | 0.038785754 | Syt10 | protein_coding |

| | | | | | | | | |
|---|---|---|---|---|---|---|---|---|
| ENSMUSG00000050910.5 | 1115.463 | -0.134 | 0.048 | -2.768 | 0.005638877 | 0.038794395 | Cdr2l | protein_coding |
| ENSMUSG00000004771.12 | 3830.992 | 0.084 | 0.030 | 2.768 | 0.005639968 | 0.038794395 | Rab11a | protein_coding |
| ENSMUSG00000055567.18 | 24243.610 | 0.075 | 0.027 | 2.767 | 0.005654084 | 0.038872518 | Unc80 | protein_coding |
| ENSMUSG00000035293.13 | 1037.761 | 0.157 | 0.057 | 2.767 | 0.005661781 | 0.038906456 | G2e3 | protein_coding |
| ENSMUSG00000035840.6 | 399.358 | 0.173 | 0.063 | 2.766 | 0.005669875 | 0.038943089 | Lysmd3 | protein_coding |
| ENSMUSG00000019173.11 | 5180.158 | -0.076 | 0.027 | -2.765 | 0.005685611 | 0.039025495 | Rab5c | protein_coding |
| ENSMUSG00000054934.10 | 3130.202 | -0.122 | 0.044 | -2.765 | 0.005687686 | 0.039025495 | Kcnmb4 | protein_coding |
| ENSMUSG00000021039.9 | 3336.649 | 0.092 | 0.033 | 2.765 | 0.00569018 | 0.039025495 | Snw1 | protein_coding |
| ENSMUSG00000026825.17 | 65705.309 | -0.072 | 0.026 | -2.764 | 0.005715101 | 0.039177351 | Dnm1 | protein_coding |
| ENSMUSG00000029674.13 | 3929.279 | -0.110 | 0.040 | -2.763 | 0.005728089 | 0.039243635 | Limk1 | protein_coding |
| ENSMUSG00000042178.6 | 675.698 | -0.160 | 0.058 | -2.763 | 0.005730339 | 0.039243635 | Armc5 | protein_coding |
| ENSMUSG00000040249.15 | 40651.053 | -0.097 | 0.035 | -2.763 | 0.005735388 | 0.039254812 | Lrp1 | protein_coding |
| ENSMUSG00000023143.10 | 514.390 | -0.184 | 0.066 | -2.762 | 0.005737542 | 0.039254812 | Nagpa | protein_coding |
| ENSMUSG00000024045.5 | 6238.589 | 0.079 | 0.029 | 2.761 | 0.005755902 | 0.039361321 | Akap8 | protein_coding |
| ENSMUSG00000016526.8 | 420.795 | -0.179 | 0.065 | -2.760 | 0.005779913 | 0.0394872 | Dyrk3 | protein_coding |
| ENSMUSG00000006095.12 | 2408.184 | -0.097 | 0.035 | -2.760 | 0.005788023 | 0.039523449 | Tbcb | protein_coding |
| ENSMUSG00000024381.15 | 12840.022 | -0.083 | 0.030 | -2.759 | 0.005806029 | 0.03961539 | Bin1 | protein_coding |
| ENSMUSG00000035314.10 | 2094.494 | -0.121 | 0.044 | -2.758 | 0.005807109 | 0.03961539 | Gdpd5 | protein_coding |
| ENSMUSG00000033813.15 | 1668.487 | 0.108 | 0.039 | 2.758 | 0.00581374 | 0.039641436 | Tcea1 | protein_coding |
| ENSMUSG00000066647.6 | 1585.416 | 0.114 | 0.041 | 2.757 | 0.005827577 | 0.039716574 | Gm5113 | protein_coding |
| ENSMUSG00000026834.13 | 1244.311 | 0.137 | 0.050 | 2.757 | 0.005833485 | 0.039737623 | Acvr1c | protein_coding |
| ENSMUSG00000045036.14 | 124.341 | 0.215 | 0.078 | 2.756 | 0.005845567 | 0.039800685 | Tmem232 | protein_coding |
| ENSMUSG00000035284.10 | 8268.370 | 0.080 | 0.029 | 2.756 | 0.005854228 | 0.039840407 | Vps13c | protein_coding |
| ENSMUSG00000024137.8 | 2338.670 | -0.115 | 0.042 | -2.755 | 0.005863247 | 0.039882529 | E4f1 | protein_coding |
| ENSMUSG00000013155.10 | 387.411 | -0.182 | 0.066 | -2.755 | 0.005874729 | 0.039941353 | Enkd1 | protein_coding |
| ENSMUSG00000027708.14 | 1468.397 | 0.122 | 0.044 | 2.754 | 0.005887303 | 0.039988263 | Dcun1d1 | protein_coding |
| ENSMUSG00000001552.14 | 3210.399 | -0.116 | 0.042 | -2.753 | 0.005906036 | 0.040093642 | Jup | protein_coding |
| ENSMUSG00000037989.15 | 10860.162 | -0.102 | 0.037 | -2.753 | 0.005911169 | 0.040093642 | Wnk2 | protein_coding |
| ENSMUSG00000008305.18 | 1931.676 | -0.119 | 0.043 | -2.753 | 0.005911352 | 0.040093642 | Tle1 | protein_coding |
| ENSMUSG00000050697.8 | 1417.420 | 0.125 | 0.046 | 2.751 | 0.005948761 | 0.040314754 | Prkaa1 | protein_coding |
| ENSMUSG00000024172.10 | 2834.719 | 0.108 | 0.039 | 2.751 | 0.005949673 | 0.040314754 | St6gal2 | protein_coding |
| ENSMUSG00000033615.9 | 21615.743 | -0.083 | 0.030 | -2.750 | 0.005952727 | 0.040316064 | Cplx1 | protein_coding |
| ENSMUSG00000025008.15 | 1934.860 | 0.102 | 0.037 | 2.750 | 0.005956814 | 0.040324372 | Tctn3 | protein_coding |

| | | | | | | | | |
|---|---|---|---|---|---|---|---|---|
| ENSMUSG00000068551.12 | 1088.754 | -0.138 | 0.050 | -2.750 | 0.005961202 | 0.040334697 | Zfp467 | protein_coding |
| ENSMUSG00000069631.14 | 1626.706 | 0.123 | 0.045 | 2.749 | 0.00597548 | 0.040411909 | Strada | protein_coding |
| ENSMUSG00000025790.14 | 2271.448 | -0.118 | 0.043 | -2.748 | 0.005987484 | 0.040473672 | Slco3a1 | protein_coding |
| ENSMUSG00000021832.7 | 3314.268 | 0.092 | 0.033 | 2.748 | 0.005998556 | 0.040529071 | Psmc6 | protein_coding |
| ENSMUSG00000049285.4 | 99.117 | -0.214 | 0.078 | -2.748 | 0.006004683 | 0.04055103 | Mblac1 | protein_coding |
| ENSMUSG00000047604.3 | 212.635 | -0.199 | 0.072 | -2.747 | 0.006007727 | 0.040552159 | Frat2 | protein_coding |
| ENSMUSG00000039782.14 | 3982.258 | 0.110 | 0.040 | 2.745 | 0.006048589 | 0.04080843 | Cpeb2 | protein_coding |
| ENSMUSG00000032397.7 | 333.840 | 0.188 | 0.069 | 2.744 | 0.00606369 | 0.040881917 | Tipin | protein_coding |
| ENSMUSG00000039578.16 | 477.317 | 0.178 | 0.065 | 2.744 | 0.006065282 | 0.040881917 | Ccser1 | protein_coding |
| ENSMUSG00000054752.16 | 2312.713 | 0.101 | 0.037 | 2.742 | 0.006103073 | 0.041116976 | Fsd1l | protein_coding |
| ENSMUSG00000001496.15 | 62.217 | -0.190 | 0.069 | -2.742 | 0.006109188 | NA | Nkx2-1 | protein_coding |
| ENSMUSG00000038530.11 | 15289.276 | 0.084 | 0.031 | 2.742 | 0.006112611 | 0.041161555 | Rgs4 | protein_coding |
| ENSMUSG00000027488.12 | 1497.250 | -0.132 | 0.048 | -2.741 | 0.006124783 | 0.041223825 | Snta1 | protein_coding |
| ENSMUSG00000027955.16 | 208.751 | 0.208 | 0.076 | 2.739 | 0.006159335 | 0.041436593 | Fam198b | protein_coding |
| ENSMUSG00000018481.9 | 5367.124 | 0.084 | 0.031 | 2.739 | 0.00616238 | 0.041437298 | Appbp2 | protein_coding |
| ENSMUSG00000037820.15 | 877.950 | -0.198 | 0.072 | -2.737 | 0.006199061 | 0.041664073 | Tgm2 | protein_coding |
| ENSMUSG00000049612.10 | 4633.062 | 0.085 | 0.031 | 2.735 | 0.006234373 | 0.041869309 | Omg | protein_coding |
| ENSMUSG00000021848.15 | 82.083 | -0.167 | 0.061 | -2.735 | 0.006235539 | 0.041869309 | Otx2 | protein_coding |
| ENSMUSG00000094114.1 | 413.993 | 0.204 | 0.075 | 2.734 | 0.006248919 | 0.041939176 | Gm21967 | protein_coding |
| ENSMUSG00000059839.8 | 826.854 | 0.152 | 0.056 | 2.734 | 0.006263913 | 0.042016847 | Zfp874b | protein_coding |
| ENSMUSG00000073792.11 | 624.500 | 0.158 | 0.058 | 2.734 | 0.006266455 | 0.042016847 | Alg6 | protein_coding |
| ENSMUSG00000074165.10 | 1768.465 | 0.128 | 0.047 | 2.733 | 0.006275422 | 0.042056968 | Zfp788 | protein_coding |
| ENSMUSG00000047648.13 | 412.804 | 0.177 | 0.065 | 2.732 | 0.006286652 | 0.042112204 | Fbxo30 | protein_coding |
| ENSMUSG00000022983.15 | 2360.445 | -0.110 | 0.040 | -2.732 | 0.006291571 | 0.042125132 | Scaf4 | protein_coding |
| ENSMUSG00000066735.8 | 3528.781 | 0.090 | 0.033 | 2.732 | 0.006302111 | 0.042175665 | Vkorc1l1 | protein_coding |
| ENSMUSG00000060639.5 | 52.900 | -0.201 | 0.074 | -2.731 | 0.006314248 | NA | Hist1h4i | protein_coding |
| ENSMUSG00000027810.14 | 1679.940 | 0.132 | 0.048 | 2.730 | 0.006334064 | 0.042369384 | Eif2a | protein_coding |
| ENSMUSG00000033849.3 | 3236.461 | 0.125 | 0.046 | 2.730 | 0.006341409 | 0.042398394 | B3galt2 | protein_coding |
| ENSMUSG00000005034.15 | 28854.178 | 0.061 | 0.022 | 2.729 | 0.006355942 | 0.042475414 | Prkacb | protein_coding |
| ENSMUSG00000042851.17 | 553.603 | 0.152 | 0.056 | 2.728 | 0.006369746 | 0.04254749 | Zc3h6 | protein_coding |
| ENSMUSG00000033055.10 | 815.844 | -0.136 | 0.050 | -2.728 | 0.006374 | 0.042555737 | Ankrd54 | protein_coding |
| ENSMUSG00000026269.14 | 2295.990 | -0.114 | 0.042 | -2.727 | 0.006398805 | 0.042701115 | Rnpepl1 | protein_coding |
| ENSMUSG00000044477.10 | 3343.999 | -0.118 | 0.043 | -2.726 | 0.0064047 | 0.042706017 | Zfand3 | protein_coding |

| | | | | | | | | |
|---|---|---|---|---|---|---|---|---|
| ENSMUSG00000024420.8 | 717.480 | -0.196 | 0.072 | -2.726 | 0.0064056 | 0.042706017 | Zfp521 | protein_coding |
| ENSMUSG00000021171.8 | 3538.054 | 0.091 | 0.033 | 2.726 | 0.006413177 | 0.042725841 | Esyt2 | protein_coding |
| ENSMUSG00000074129.13 | 235.171 | -0.195 | 0.072 | -2.726 | 0.006414636 | 0.042725841 | Rpl13a | protein_coding |
| ENSMUSG00000041014.17 | 1717.614 | 0.123 | 0.045 | 2.726 | 0.006419917 | 0.042740816 | Nrg3 | protein_coding |
| ENSMUSG00000024560.6 | 2503.614 | -0.103 | 0.038 | -2.725 | 0.006427504 | 0.042771122 | Cxxc1 | protein_coding |
| ENSMUSG00000014444.17 | 663.923 | -0.180 | 0.066 | -2.725 | 0.006433626 | 0.042791657 | Piezo1 | protein_coding |
| ENSMUSG00000030401.16 | 3844.811 | -0.124 | 0.046 | -2.724 | 0.006456423 | 0.042922783 | Rtn2 | protein_coding |
| ENSMUSG00000015656.17 | 65683.816 | -0.083 | 0.030 | -2.723 | 0.006459431 | 0.042922783 | Hspa8 | protein_coding |
| ENSMUSG00000031916.17 | 1122.218 | -0.126 | 0.046 | -2.723 | 0.006465627 | 0.04294371 | Cog8 | protein_coding |
| ENSMUSG00000021037.7 | 4742.147 | -0.080 | 0.029 | -2.722 | 0.006487034 | 0.043065593 | Ahsa1 | protein_coding |
| ENSMUSG00000043448.13 | 547.355 | -0.208 | 0.076 | -2.717 | 0.006582017 | 0.043674573 | Gjc2 | protein_coding |
| ENSMUSG00000089911.4 | 2239.967 | 0.113 | 0.041 | 2.717 | 0.006584963 | 0.043674573 | Mfsd14a | protein_coding |
| ENSMUSG00000031150.12 | 792.878 | -0.136 | 0.050 | -2.717 | 0.006591749 | 0.043699014 | Ccdc120 | protein_coding |
| ENSMUSG00000032788.15 | 12161.495 | -0.102 | 0.038 | -2.716 | 0.006615343 | 0.043834808 | Pdxk | protein_coding |
| ENSMUSG00000040648.14 | 1916.818 | 0.125 | 0.046 | 2.715 | 0.006626085 | 0.043885356 | Ppip5k2 | protein_coding |
| ENSMUSG00000006154.13 | 408.391 | -0.167 | 0.062 | -2.715 | 0.006635473 | 0.043926896 | Eps8l1 | protein_coding |
| ENSMUSG00000027009.18 | 851.224 | 0.190 | 0.070 | 2.714 | 0.006641903 | 0.043948819 | Itga4 | protein_coding |
| ENSMUSG00000043683.3 | 2939.697 | -0.105 | 0.039 | -2.714 | 0.006651113 | 0.043989107 | Fem1a | protein_coding |
| ENSMUSG00000020614.13 | 831.777 | -0.149 | 0.055 | -2.713 | 0.006672281 | 0.044108409 | Fam20a | protein_coding |
| ENSMUSG00000024777.9 | 6707.032 | -0.097 | 0.036 | -2.711 | 0.006698487 | 0.04426089 | Ppp2r5b | protein_coding |
| ENSMUSG00000032171.7 | 2803.766 | -0.109 | 0.040 | -2.711 | 0.006706056 | 0.044290135 | Pin1 | protein_coding |
| ENSMUSG00000026767.12 | 1645.453 | 0.110 | 0.041 | 2.708 | 0.006771409 | 0.044700812 | Fam188a | protein_coding |
| ENSMUSG00000036907.9 | 43.696 | -0.191 | 0.071 | -2.706 | 0.006803759 | NA | C1ql2 | protein_coding |
| ENSMUSG00000026648.18 | 872.248 | 0.142 | 0.052 | 2.705 | 0.006821518 | 0.045010522 | Dclre1c | protein_coding |
| ENSMUSG00000025321.14 | 2513.808 | 0.110 | 0.041 | 2.705 | 0.006832186 | 0.045059816 | Itgb8 | protein_coding |
| ENSMUSG00000007216.6 | 292.151 | -0.190 | 0.070 | -2.704 | 0.006841802 | 0.045074542 | Zfp775 | protein_coding |
| ENSMUSG00000014859.9 | 1489.180 | -0.115 | 0.042 | -2.704 | 0.006844013 | 0.045074542 | E2f4 | protein_coding |
| ENSMUSG00000045107.4 | 278.928 | -0.196 | 0.072 | -2.703 | 0.006878541 | 0.045280786 | Saysd1 | protein_coding |
| ENSMUSG00000068039.12 | 8061.654 | 0.082 | 0.031 | 2.702 | 0.006882277 | 0.045284225 | Tcp1 | protein_coding |
| ENSMUSG00000008226.14 | 1345.839 | 0.125 | 0.046 | 2.702 | 0.006889846 | 0.045312871 | Scrn3 | protein_coding |
| ENSMUSG00000026610.13 | 1123.516 | 0.149 | 0.055 | 2.702 | 0.006899454 | 0.045349659 | Esrrg | protein_coding |
| ENSMUSG00000049878.13 | 2161.002 | 0.105 | 0.039 | 2.702 | 0.006901874 | 0.045349659 | Rlf | protein_coding |
| ENSMUSG00000052364.4 | 1371.834 | -0.131 | 0.049 | -2.701 | 0.006911133 | 0.045389334 | B630019K06Rik | protein_coding |

| | | | | | | | | |
|---|---|---|---|---|---|---|---|---|
| ENSMUSG00000042410.15 | 2344.567 | 0.107 | 0.040 | 2.701 | 0.006922701 | 0.04544413 | Agps | protein_coding |
| ENSMUSG00000030513.14 | 1121.150 | -0.136 | 0.050 | -2.700 | 0.006936232 | 0.04548413 | Pcsk6 | protein_coding |
| ENSMUSG00000045509.6 | 193.486 | -0.198 | 0.073 | -2.700 | 0.006938222 | 0.04548413 | Gpr150 | protein_coding |
| ENSMUSG00000091735.2 | 290.588 | -0.204 | 0.076 | -2.700 | 0.006938476 | 0.04548413 | Gpr62 | protein_coding |
| ENSMUSG00000038967.13 | 5227.473 | -0.088 | 0.033 | -2.699 | 0.006947404 | 0.045521485 | Pdk2 | protein_coding |
| ENSMUSG00000059878.12 | 967.968 | 0.125 | 0.046 | 2.699 | 0.006952317 | 0.045532504 | Zfp422 | protein_coding |
| ENSMUSG00000005262.12 | 3552.824 | 0.088 | 0.033 | 2.699 | 0.006964035 | 0.04558807 | Ufd1 | protein_coding |
| ENSMUSG00000038615.17 | 18271.517 | -0.058 | 0.022 | -2.698 | 0.006976156 | 0.045646216 | Nfe2l1 | protein_coding |
| ENSMUSG00000028496.17 | 1575.401 | 0.120 | 0.045 | 2.697 | 0.006990241 | 0.045674758 | Mllt3 | protein_coding |
| ENSMUSG00000027111.15 | 1265.562 | 0.126 | 0.047 | 2.696 | 0.007008464 | 0.045772609 | Itga6 | protein_coding |
| ENSMUSG00000060904.14 | 5263.984 | 0.079 | 0.029 | 2.693 | 0.007072686 | 0.046170653 | Arl1 | protein_coding |
| ENSMUSG00000032570.17 | 8878.145 | 0.073 | 0.027 | 2.693 | 0.007090744 | 0.046267107 | Atp2c1 | protein_coding |
| ENSMUSG00000016382.15 | 10739.546 | 0.104 | 0.039 | 2.692 | 0.007103226 | 0.046327102 | Pls3 | protein_coding |
| ENSMUSG00000036916.13 | 1515.935 | 0.142 | 0.053 | 2.692 | 0.007109242 | 0.046344895 | Zfp280c | protein_coding |
| ENSMUSG00000036790.5 | 1309.742 | 0.123 | 0.046 | 2.691 | 0.007117785 | 0.046379134 | Slitrk2 | protein_coding |
| ENSMUSG00000026281.15 | 1166.070 | -0.125 | 0.046 | -2.691 | 0.007125334 | 0.046406869 | Dtymk | protein_coding |
| ENSMUSG00000039910.10 | 1138.082 | -0.140 | 0.052 | -2.691 | 0.007134382 | 0.04643725 | Cited2 | protein_coding |
| ENSMUSG00000037720.16 | 4761.924 | 0.077 | 0.029 | 2.690 | 0.007136589 | 0.04643725 | Tmem33 | protein_coding |
| ENSMUSG00000024012.17 | 15839.944 | -0.106 | 0.039 | -2.690 | 0.007140933 | 0.046444076 | Mtch1 | protein_coding |
| ENSMUSG00000068740.13 | 17826.844 | -0.075 | 0.028 | -2.689 | 0.007167096 | 0.046592738 | Celsr2 | protein_coding |
| ENSMUSG00000058748.9 | 390.564 | 0.184 | 0.069 | 2.687 | 0.007206343 | 0.046810091 | Zfp958 | protein_coding |
| ENSMUSG00000019689.4 | 370.163 | -0.170 | 0.063 | -2.687 | 0.007207173 | 0.046810091 | Fmc1 | protein_coding |
| ENSMUSG00000011877.13 | 19549.112 | -0.092 | 0.034 | -2.686 | 0.007233572 | 0.046922564 | Git1 | protein_coding |
| ENSMUSG00000031393.16 | 9061.807 | 0.062 | 0.023 | 2.686 | 0.007234478 | 0.046922564 | Mecp2 | protein_coding |
| ENSMUSG00000032409.15 | 1125.174 | 0.157 | 0.058 | 2.686 | 0.007237827 | 0.046922692 | Atr | protein_coding |
| ENSMUSG00000041548.4 | 597.647 | -0.173 | 0.065 | -2.684 | 0.00728405 | 0.047197763 | Hspb8 | protein_coding |
| ENSMUSG00000053347.14 | 1087.190 | 0.134 | 0.050 | 2.683 | 0.007287066 | 0.047197763 | Zfp943 | protein_coding |
| ENSMUSG00000044067.6 | 2387.410 | 0.124 | 0.046 | 2.683 | 0.007290303 | 0.047197763 | Gpr22 | protein_coding |
| ENSMUSG00000031099.16 | 2363.641 | 0.121 | 0.045 | 2.683 | 0.007306212 | 0.047279043 | Smarca1 | protein_coding |
| ENSMUSG00000037857.16 | 2184.582 | 0.120 | 0.045 | 2.682 | 0.007314549 | 0.047311267 | Nufip2 | protein_coding |
| ENSMUSG00000027287.14 | 962.019 | 0.132 | 0.049 | 2.682 | 0.007322486 | 0.04734088 | Snap23 | protein_coding |
| ENSMUSG00000034973.16 | 4085.498 | 0.084 | 0.031 | 2.681 | 0.007339583 | 0.047429655 | Dopey1 | protein_coding |
| ENSMUSG00000078185.3 | 895.546 | 0.133 | 0.050 | 2.680 | 0.007361616 | 0.047550239 | Chml | protein_coding |

| | | | | | | | | |
|---|---|---|---|---|---|---|---|---|
| ENSMUSG00000002617.14 | 1165.963 | 0.140 | 0.052 | 2.679 | 0.007378464 | 0.047637232 | Zfp40 | protein_coding |
| ENSMUSG00000062619.5 | 58.950 | -0.200 | 0.075 | -2.679 | 0.00737943 | NA | 2310039H08Rik | protein_coding |
| ENSMUSG00000038831.16 | 6307.480 | 0.078 | 0.029 | 2.678 | 0.007400811 | 0.047759632 | Ralgps1 | protein_coding |
| ENSMUSG00000029405.16 | 16300.226 | 0.070 | 0.026 | 2.678 | 0.007408037 | 0.047784383 | G3bp2 | protein_coding |
| ENSMUSG00000033282.14 | 1762.898 | 0.108 | 0.040 | 2.677 | 0.007422698 | 0.047844895 | Rpgrip1l | protein_coding |
| ENSMUSG00000030629.15 | 2268.933 | 0.106 | 0.039 | 2.677 | 0.007424208 | 0.047844895 | Zfand6 | protein_coding |
| ENSMUSG00000028322.11 | 274.590 | -0.194 | 0.072 | -2.677 | 0.007439339 | 0.047920492 | Exosc3 | protein_coding |
| ENSMUSG00000023883.13 | 3234.964 | 0.089 | 0.033 | 2.676 | 0.007451321 | 0.04797575 | Phf10 | protein_coding |
| ENSMUSG00000027201.16 | 4202.662 | 0.086 | 0.032 | 2.676 | 0.007458014 | 0.047996918 | Myef2 | protein_coding |
| ENSMUSG00000002043.17 | 227.930 | -0.188 | 0.070 | -2.675 | 0.007482645 | 0.048133453 | Trappc6a | protein_coding |
| ENSMUSG00000028161.17 | 41244.093 | 0.082 | 0.031 | 2.674 | 0.007487491 | 0.048142655 | Ppp3ca | protein_coding |
| ENSMUSG00000050321.2 | 3482.175 | 0.093 | 0.035 | 2.674 | 0.007499394 | 0.048197199 | Neto1 | protein_coding |
| ENSMUSG00000039270.9 | 6770.498 | 0.086 | 0.032 | 2.674 | 0.007503347 | 0.048200628 | Megf9 | protein_coding |
| ENSMUSG00000047731.17 | 2050.544 | -0.101 | 0.038 | -2.673 | 0.007512855 | 0.048217751 | Wbp1l | protein_coding |
| ENSMUSG00000063785.12 | 966.427 | 0.128 | 0.048 | 2.673 | 0.007517067 | 0.048222826 | Utp14a | protein_coding |
| ENSMUSG00000032410.13 | 2476.364 | 0.091 | 0.034 | 2.672 | 0.007540384 | 0.04832841 | Xrn1 | protein_coding |
| ENSMUSG00000010914.10 | 1946.506 | 0.110 | 0.041 | 2.670 | 0.007574058 | 0.048522167 | Pdhx | protein_coding |
| ENSMUSG00000061118.8 | 1274.726 | -0.130 | 0.049 | -2.670 | 0.007582972 | 0.048557206 | Dnajc30 | protein_coding |
| ENSMUSG00000057421.12 | 3418.129 | 0.091 | 0.034 | 2.669 | 0.007613474 | 0.048730379 | Las1l | protein_coding |
| ENSMUSG00000014932.15 | 574.521 | 0.145 | 0.054 | 2.668 | 0.007636795 | 0.048857461 | Yes1 | protein_coding |
| ENSMUSG00000003099.11 | 5552.623 | -0.088 | 0.033 | -2.667 | 0.007647231 | 0.048889144 | Ppp5c | protein_coding |
| ENSMUSG00000042804.13 | 1206.048 | -0.178 | 0.067 | -2.667 | 0.007649974 | 0.048889144 | Gpr153 | protein_coding |
| ENSMUSG00000044066.14 | 1914.024 | 0.110 | 0.041 | 2.667 | 0.007652154 | 0.048889144 | Cep68 | protein_coding |
| ENSMUSG00000003299.10 | 1900.200 | -0.114 | 0.043 | -2.667 | 0.007662792 | 0.048934928 | Mrpl4 | protein_coding |
| ENSMUSG00000010406.11 | 445.176 | -0.172 | 0.065 | -2.666 | 0.007674969 | 0.048982569 | Mrpl52 | protein_coding |
| ENSMUSG00000042197.13 | 3407.182 | 0.092 | 0.034 | 2.666 | 0.007680858 | 0.048982569 | Zfp451 | protein_coding |
| ENSMUSG00000036437.6 | 1293.833 | 0.140 | 0.052 | 2.666 | 0.007682872 | 0.048982569 | Npy1r | protein_coding |
| ENSMUSG00000036615.6 | 962.322 | -0.142 | 0.053 | -2.666 | 0.007684154 | 0.048982569 | Rfxap | protein_coding |
| ENSMUSG00000018446.8 | 1737.771 | -0.119 | 0.045 | -2.663 | 0.007737277 | 0.049298907 | C1qbp | protein_coding |
| ENSMUSG00000034659.14 | 2439.170 | -0.129 | 0.049 | -2.663 | 0.007750776 | 0.049362599 | Tmem109 | protein_coding |
| ENSMUSG00000036257.15 | 6555.799 | 0.080 | 0.030 | 2.661 | 0.007794153 | 0.049594034 | Pnpla8 | protein_coding |
| ENSMUSG00000024054.13 | 2452.193 | 0.099 | 0.037 | 2.658 | 0.00785059 | 0.0499306 | Smchd1 | protein_coding |
| ENSMUSG00000020034.7 | 1302.962 | 0.122 | 0.046 | 2.658 | 0.007859044 | 0.049961827 | Tcp11l2 | protein_coding |

**Supplementary Table S3.** Gene expression changes in mPFC (FDR 5%) in Plcb1+/- mice compared to WT after cocaine seeking reinstatement

| ids | baseMean | log2FoldChange | lfcSE | stat | pvalue | padj | Gene Name | Gene type |
|---|---|---|---|---|---|---|---|---|
| ENSMUSG00000022769.7 | 906.219 | -0.242 | 0.039 | -6.25 | 4.22E-10 | 9.05E-06 | Sdf2l1 | protein_coding |
| ENSMUSG00000026864.13 | 25343.092 | -0.156 | 0.034 | -4.57 | 4.82E-06 | 0.021 | Hspa5 | protein_coding |
| ENSMUSG00000038393.14 | 2084.497 | -0.162 | 0.034 | -4.75 | 2.03E-06 | 0.021 | Txnip | protein_coding |
| ENSMUSG00000040046.14 | 69.482 | -0.088 | 0.019 | -4.60 | 4.19E-06 | 0.021 | Tph1 | protein_coding |
| ENSMUSG00000021025.8 | 960.708 | -0.178 | 0.039 | -4.63 | 3.68E-06 | 0.021 | Nfkbia | protein_coding |
| ENSMUSG00000027342.14 | 1321.497 | -0.156 | 0.036 | -4.31 | 1.62E-05 | 0.035 | Pcna | protein_coding |
| ENSMUSG00000003814.8 | 21535.466 | -0.128 | 0.029 | -4.34 | 1.40E-05 | 0.035 | Calr | protein_coding |
| ENSMUSG00000005958.14 | 1125.700 | 0.155 | 0.036 | 4.31 | 1.61E-05 | 0.035 | Ephb3 | protein_coding |
| ENSMUSG00000024966.8 | 8739.320 | -0.115 | 0.026 | -4.38 | 1.19E-05 | 0.035 | Stip1 | protein_coding |
| ENSMUSG00000067608.4 | 81.222 | 0.061 | 0.014 | 4.40 | 1.08E-05 | 0.035 | Pcna-ps2 | protein_coding |
| ENSMUSG00000004951.10 | 382.579 | -0.156 | 0.037 | -4.23 | 2.32E-05 | 0.042 | Hspb1 | protein_coding |
| ENSMUSG00000096768.7 | 184.618 | -0.146 | 0.034 | -4.24 | 2.28E-05 | 0.042 | Erdr1 | protein_coding |

**Supplementary Table S4.** Enrichment of GO (Gene Ontology) terms in differentially expressed genes in mPFC

| Term | Genes | Count | % | P-Value | Benjamini |
|---|---|---|---|---|---|
| Type I pneumocyte differentiation | | 6 | 0,3 | 2,2E-4 | 3,3E-2 |
| regulation of alpha-amino-3-hydroxy-5-methyl-4-isoxazole propionate selective glutamate receptor activity | | 10 | 0,5 | 3,0E-5 | 6,9E-3 |
| mRNA splice site selection | | 11 | 0,5 | 2,8E-7 | 2,7E-4 |
| positive regulation of excitatory postsynaptic potential | | 11 | 0,5 | 3,4E-4 | 4,8E-2 |
| neurotransmitter secretion | | 13 | 0,6 | 1,0E-5 | 3,5E-3 |
| dendrite morphogenesis | | 15 | 0,7 | 7,5E-5 | 1,4E-2 |
| long-term synaptic potentiation | | 15 | 0,7 | 3,5E-4 | 4,8E-2 |
| synapse organization | | 16 | 0,7 | 1,9E-6 | 9,0E-4 |
| learning | | 19 | 0,9 | 1,0E-4 | 1,9E-2 |
| response to endoplasmic reticulum stress | | 23 | 1,0 | 6,5E-6 | 2,6E-3 |
| regulation of cell cycle | | 26 | 1,2 | 1,9E-4 | 3,0E-2 |
| axonogenesis | | 28 | 1,3 | 1,1E-5 | 3,4E-3 |
| rhythmic process | | 28 | 1,3 | 2,9E-4 | 4,2E-2 |
| regulation of translation | | 30 | 1,4 | 1,5E-5 | 4,2E-3 |
| peptidyl-serine phosphorylation | | 30 | 1,4 | 9,5E-5 | 1,8E-2 |
| neuron projection development | | 32 | 1,5 | 3,5E-5 | 7,7E-3 |
| actin cytoskeleton organization | | 33 | 1,5 | 2,1E-5 | 5,4E-3 |
| ubiquitin-dependent protein catabolic process | | 35 | 1,6 | 1,9E-5 | 5,1E-3 |
| chemical synaptic transmission | | 35 | 1,6 | 1,9E-4 | 3,0E-2 |
| endocytosis | | 40 | 1,8 | 8,7E-6 | 3,3E-3 |
| cell-cell adhesion | | 40 | 1,8 | 2,5E-5 | 6,1E-3 |
| brain development | | 48 | 2,2 | 8,2E-7 | 5,0E-4 |
| RNA splicing | | 54 | 2,5 | 1,2E-7 | 1,4E-4 |
| positive regulation of apoptotic process | | 58 | 2,6 | 1,5E-4 | 2,5E-2 |
| mRNA processing | | 65 | 3,0 | 3,1E-7 | 2,5E-4 |
| nervous system development | | 85 | 3,9 | 1,2E-11 | 2,9E-8 |
| protein phosphorylation | | 90 | 4,1 | 1,0E-4 | 1,8E-2 |
| apoptotic process | | 91 | 4,1 | 4,3E-5 | 8,7E-3 |
| protein transport | | 98 | 4,5 | 4,6E-6 | 2,0E-3 |
| phosphorylation | | 99 | 4,5 | 1,1E-5 | 3,4E-3 |

| | | | | |
|---|---|---|---|---|
| negative regulation of transcription from RNA polymerase II promoter | 120 | 5,5 | 4,8E-7 | 3,3E-4 |
| positive regulation of transcription from RNA polymerase II promoter | 144 | 6,5 | 4,3E-5 | 9,0E-3 |
| transport | 252 | 11,4 | 1,5E-6 | 8,1E-4 |
| transcription, DNA-templated | 302 | 13,7 | 3,9E-15 | 1,9E-11 |
| regulation of transcription, DNA-templated | 335 | 15,2 | 1,2E-11 | 2,0E-8 |

**Supplementary Table S5.** Enrichment of KEGG (Kyoto Encyclopedia of Genes and Genomes) terms in differentially expressed genes in mPFC

| Terms | Genes | Count | % | P-Value | Benjamini |
|---|---|---|---|---|---|
| Endocytosis | | 56 | 2,5 | 1,2E-7 | 3,2E-5 |
| Protein processing in endoplasmic reticulum | | 40 | 1,8 | 1,2E-7 | 1,6E-5 |
| MAPK signaling pathway | | 52 | 2,4 | 1,9E-7 | 1,7E-5 |
| mTOR signaling pathway | | 20 | 0,9 | 1,3E-6 | 8,8E-5 |
| Ubiquitin mediated proteolysis | | 33 | 1,5 | 3,3E-6 | 1,8E-4 |
| Amphetamine addiction | | 19 | 0,9 | 4,1E-5 | 1,8E-3 |
| Synaptic vesicle cycle | | 18 | 0,8 | 5,0E-5 | 1,9E-3 |
| Nicotine addiction | | 14 | 0,6 | 5,6E-5 | 1,9E-3 |
| Long-term potentiation | | 18 | 0,8 | 1,2E-4 | 3,5E-3 |
| FoxO signaling pathway | | 28 | 1,3 | 1,5E-4 | 4,0E-3 |
| Neurotrophin signaling pathway | | 26 | 1,2 | 2,0E-4 | 4,8E-3 |
| B cell receptor signaling pathway | | 18 | 0,8 | 2,6E-4 | 5,7E-3 |
| Spliceosome | | 27 | 1,2 | 3,3E-4 | 6,7E-3 |
| Estrogen signaling pathway | | 22 | 1,0 | 3,3E-4 | 6,4E-3 |
| Renal cell carcinoma | | 17 | 0,8 | 4,7E-4 | 8,4E-3 |
| Choline metabolism in cancer | | 22 | 1,0 | 5,1E-4 | 8,6E-3 |
| Insulin signaling pathway | | 27 | 1,2 | 7,5E-4 | 1,2E-2 |
| Colorectal cancer | | 16 | 0,7 | 8,7E-4 | 1,3E-2 |
| Glioma | | 16 | 0,7 | 1,0E-3 | 1,4E-2 |
| Chronic myeloid leukemia | | 17 | 0,8 | 1,1E-3 | 1,5E-2 |
| Axon guidance | | 25 | 1,1 | 1,1E-3 | 1,5E-2 |
| ErbB signaling pathway | | 19 | 0,9 | 1,3E-3 | 1,6E-2 |
| VEGF signaling pathway | | 15 | 0,7 | 1,3E-3 | 1,6E-2 |
| Prostate cancer | | 19 | 0,9 | 1,5E-3 | 1,7E-2 |
| Insulin resistance | | 22 | 1,0 | 1,6E-3 | 1,7E-2 |
| Dopaminergic synapse | | 25 | 1,1 | 2,0E-3 | 2,0E-2 |
| Non-alcoholic fatty liver disease (NAFLD) | | 28 | 1,3 | 2,0E-3 | 1,9E-2 |
| HTLV-I infection | | 43 | 2,0 | 2,0E-3 | 1,9E-2 |
| cAMP signaling pathway | | 33 | 1,5 | 2,1E-3 | 1,9E-2 |
| Oxytocin signaling pathway | | 28 | 1,3 | 2,2E-3 | 1,9E-2 |
| Amyotrophic lateral sclerosis (ALS) | | 13 | 0,6 | 2,7E-3 | 2,3E-2 |
| Pancreatic cancer | | 15 | 0,7 | 3,0E-3 | 2,5E-2 |
| Endometrial cancer | | 13 | 0,6 | 3,2E-3 | 2,6E-2 |
| Progesterone-mediated oocyte maturation | | 18 | 0,8 | 3,4E-3 | 2,7E-2 |
| Osteoclast differentiation | | 23 | 1,0 | 4,1E-3 | 3,1E-2 |
| Focal adhesion | | 33 | 1,5 | 4,6E-3 | 3,4E-2 |
| Acute myeloid leukemia | | 13 | 0,6 | 6,1E-3 | 4,3E-2 |
| Signaling pathways regulating pluripotency of stem cells | | 24 | 1,1 | 6,1E-3 | 4,2E-2 |
| Hepatitis B | | 25 | 1,1 | 6,2E-3 | 4,2E-2 |
| Melanoma | | 15 | 0,7 | 6,9E-3 | 4,5E-2 |
| Central carbon metabolism in cancer | | 14 | 0,6 | 7,0E-3 | 4,5E-2 |
| Adipocytokine signaling pathway | | 15 | 0,7 | 7,9E-3 | 4,9E-2 |
| Epstein-Barr virus infection | | 33 | 1,5 | 8,1E-3 | 4,9E-2 |

**Supplementary Table S6.** Correspondence between genes and proteins from the Dopaminergic synapse from KEGG pathways

| Protein in Figure 4 | Gene Symbol | Gene Name | Entrez Gene ID | RNAseq FC |
|---|---|---|---|---|
| Akt | Akt1 | thymoma viral proto-oncogene 1 | 11651 | -1.068 |
| | Akt2 | thymoma viral proto-oncogene 2 | 11652 | -1.088 |
| | Akt3 | thymoma viral proto-oncogene 3 | 23797 | 1.052 |
| Cav2.1/2.2 and Vgcc | Cacna1a | calcium channel, voltage-dependent, P/Q type, alpha 1A subunit | 12286 | -1.093 |
| Cam | Calm2 | calmodulin 2 | 12314 | 1.092 |
| | Calm3 | calmodulin 3 | 12315 | -1.079 |
| Comt | Comt | catechol-O-methyltransferase | 12846 | -1.091 |
| Creb | Atf6b | activating transcription factor 6 beta | 12915 | -1.084 |
| | Creb1 | cAMP responsive element binding protein 1 | 12912 | 1.069 |
| Gi/o | Gnao1 | guanine nucleotide binding protein, alpha O | 14681 | -1.051 |
| | Gng2 | guanine nucleotide binding protein (G protein), gamma 2 | 14702 | 1.104 |
| Ampar | Gria2 | glutamate receptor, ionotropic, AMPA2 (alpha 2) | 14800 | 1.133 |
| | Gria3 | glutamate receptor, ionotropic, AMPA3 (alpha 3) | 53623 | 1.090 |
| | Gria4 | glutamate receptor, ionotropic, AMPA4 (alpha 4) | 14802 | 1.070 |
| Girk | Kcnj3 | potassium inwardly-rectifying channel, subfamily J, member 3 | 16519 | 1.103 |
| Mao | Maoa | monoamine oxidase A | 17161 | 1.115 |
| Mapk | Mapk8 | mitogen-activated protein kinase 8 | 26419 | 1.088 |
| | Mapk9 | mitogen-activated protein kinase 9 | 26420 | 1.065 |
| Pp1 | Ppp1cb | protein phosphatase 1 catalytic subunit beta | 19046 | 1.152 |
| Pp2a | Ppp2r3d | protein phosphatase 2 (formerly 2A), regulatory subunit B'', delta | 19054 | -1.213 |
| | Ppp2r5b | protein phosphatase 2, regulatory subunit B', beta | 225849 | -1.070 |
| Pp2b | Ppp3ca | protein phosphatase 3, catalytic subunit, alpha isoform | 19055 | 1.058 |
| | Ppp3cb | protein phosphatase 3, catalytic subunit, beta isoform | 19056 | 1.134 |
| Pka | Prkacb | protein kinase, cAMP dependent, catalytic, beta | 18749 | 1.043 |
| Vssc | Scn1a | sodium channel, voltage-gated, type I, alpha | 20265 | 1.097 |

FC= Fold-change from RNAseq of mPFC comparing Plcb1+/- and WT mice. In red, genes upregulated. In green, genes downregulated.

**Supplementary Table S6.** Enrichment of transcription factor binding sites (TFBS) in genes differentially expressed in mPFC

| TFBS | Size | Expect | Observed | Ratio | P Value | FDR |
|---|---|---|---|---|---|---|
| GCCATNTTG_YY1_Q6 | 672 | 78.886 | **Size** | 1.724 | 6.20E-11 | 1.38E-08 |
| MAZR_01 | 368 | 43.199 | 87 | 2.0139 | 7.31E-11 | 1.38E-08 |
| YGCGYRCGC_UNKNOWN | 459 | 53.882 | 102 | 1.893 | 7.91E-11 | 1.38E-08 |
| GCANCTGNY_MYOD_Q6 | 1475 | 173.15 | 252 | 1.4554 | 9.15E-11 | 1.38E-08 |
| AP2ALPHA_01 | 380 | 44.608 | 88 | 1.9727 | 1.78E-10 | 2.16E-08 |
| CTGCAGY_UNKNOWN | 1236 | 145.09 | 216 | 1.4887 | 3.31E-10 | 3.34E-08 |
| TGCGCANK_UNKNOWN | 804 | 94.381 | 153 | 1.6211 | 4.41E-10 | 3.81E-08 |
| CACBINDINGPROTEIN_Q6 | 397 | 46.604 | 89 | 1.9097 | 8.20E-10 | 6.20E-08 |
| RCGCANGCGY_NRF1_Q6 | 1263 | 148.26 | 216 | 1.4569 | 2.34E-09 | 1.57E-07 |
| EGR1_01 | 429 | 50.36 | 90 | 1.7871 | 2.07E-08 | 1.25E-06 |
| AP2GAMMA_01 | 382 | 44.843 | 82 | 1.8286 | 3.07E-08 | 1.69E-06 |
| ERR1_Q2 | 414 | 48.599 | 86 | 1.7696 | 6.82E-08 | 3.32E-06 |
| TCF1P_Q6 | 421 | 49.421 | 87 | 1.7604 | 7.32E-08 | 3.32E-06 |
| EGR_Q6 | 396 | 46.486 | 83 | 1.7855 | 7.68E-08 | 3.32E-06 |
| VDR_Q3 | 366 | 42.964 | 78 | 1.8155 | 9.17E-08 | 3.70E-06 |
| TGACCTY_ERR1_Q2 | 1551 | 182.07 | 247 | 1.3566 | 1.11E-07 | 4.20E-06 |
| FAC1_01 | 344 | 40.382 | 73 | 1.8077 | 2.82E-07 | 1.00E-05 |
| RNGTGGGC_UNKNOWN | 1164 | 136.64 | 192 | 1.4051 | 3.09E-07 | 1.04E-05 |
| CCCNNGGGAR_OLF1_01 | 521 | 61.16 | 100 | 1.6351 | 3.47E-07 | 1.11E-05 |
| PAX4_01 | 408 | 47.895 | 82 | 1.7121 | 5.86E-07 | 1.77E-05 |
| WTTGKCTG_UNKNOWN | 846 | 99.311 | 146 | 1.4701 | 6.85E-07 | 1.97E-05 |
| RTAAACA_FREAC2_01 | 1379 | 161.88 | 218 | 1.3467 | 1.2171E-06 | 3.18E-05 |
| AAAYRNCTG_UNKNOWN | 596 | 69.964 | 109 | 1.5579 | 1.2211E-06 | 3.18E-05 |
| LFA1_Q6 | 370 | 43.434 | 75 | 1.7268 | 1.2629E-06 | 3.18E-05 |
| AR_01 | 246 | 28.878 | 55 | 1.9046 | 1.5287E-06 | 3.62E-05 |
| GATA1_01 | 411 | 48.247 | 81 | 1.6789 | 1.5557E-06 | 3.62E-05 |
| PAX4_03 | 452 | 53.06 | 87 | 1.6397 | 1.7888E-06 | 3.93E-05 |
| YY1_Q6 | 406 | 47.66 | 80 | 1.6786 | 1.8168E-06 | 3.93E-05 |
| TCCATTKW_UNKNOWN | 388 | 45.547 | 77 | 1.6906 | 2.1371E-06 | 4.46E-05 |
| ZF5_B | 383 | 44.96 | 76 | 1.6904 | 2.4956E-06 | 5.03E-05 |
| NGFIC_01 | 399 | 46.838 | 78 | 1.6653 | 3.3359E-06 | 6.43E-05 |
| FOXJ2_02 | 373 | 43.786 | 74 | 1.69 | 3.4025E-06 | 6.43E-05 |
| MYOD_01 | 466 | 54.703 | 88 | 1.6087 | 3.5165E-06 | 6.45E-05 |
| TFIII_Q6 | 335 | 39.325 | 68 | 1.7292 | 3.7303E-06 | 6.64E-05 |
| TGCTGAY_UNKNOWN | 952 | 111.75 | 157 | 1.4049 | 4.1129E-06 | 6.98E-05 |
| YY1_02 | 336 | 39.443 | 68 | 1.724 | 4.1526E-06 | 6.98E-05 |
| EGR2_01 | 317 | 37.212 | 65 | 1.7467 | 0.000004276 | 6.99E-05 |
| MTF1_Q4 | 435 | 51.064 | 83 | 1.6254 | 4.3921E-06 | 6.99E-05 |
| TTCYRGAA_UNKNOWN | 523 | 61.395 | 96 | 1.5637 | 4.5215E-06 | 7.01E-05 |
| CATTGTYY_SOX9_B1 | 586 | 68.79 | 105 | 1.5264 | 4.9945E-06 | 7.40E-05 |
| GATTGGY_NFY_Q6_01 | 1839 | 215.88 | 274 | 1.2692 | 6.9387E-06 | 8.90E-05 |
| TGACATY_UNKNOWN | 1034 | 121.38 | 167 | 1.3758 | 7.0622E-06 | 8.90E-05 |
| CDP_02 | 220 | 25.826 | 48 | 1.8586 | 0.000013889 | 1.56E-04 |
| EGR3_01 | 149 | 17.491 | 36 | 2.0582 | 0.000016232 | 1.70E-04 |
| SOX9_B1 | 377 | 44.256 | 72 | 1.6269 | 0.000018087 | 1.77E-04 |
| GTGACGY_E4F1_Q6 | 965 | 113.28 | 155 | 1.3683 | 0.00002078 | 1.96E-04 |

| | | | | | | |
|---|---|---|---|---|---|---|
| TGACCTTG_SF1_Q6 | 391 | 45.899 | 73 | 1.5904 | 0.000034911 | 2.97E-04 |
| KCCGNSWTTT_UNKNOWN | 150 | 17.608 | 35 | 1.9877 | 0.000046306 | 3.74E-04 |
| OCT1_04 | 355 | 41.673 | 67 | 1.6077 | 0.00005124 | 4.08E-04 |
| HSF_Q6 | 356 | 41.791 | 67 | 1.6032 | 0.00005605 | 4.29E-04 |
| HEN1_01 | 317 | 37.212 | 61 | 1.6392 | 0.000061522 | 4.65E-04 |
| AP1_Q4_01 | 446 | 52.356 | 79 | 1.5089 | 0.00010872 | 7.39E-04 |
| NRF2_Q4 | 428 | 50.243 | 76 | 1.5127 | 0.00013453 | 8.84E-04 |
| TGACAGNY_MEIS1_01 | 1337 | 156.95 | 198 | 1.2616 | 0.0002177 | 1.24E-03 |
| CACGTG_MYC_Q2 | 1558 | 182.89 | 226 | 1.2357 | 0.00026579 | 1.45E-03 |
| E2F_Q2 | 288 | 33.808 | 54 | 1.5973 | 0.00031278 | 1.66E-03 |
| RGAANNTTC_HSF1_01 | 767 | 90.038 | 120 | 1.3328 | 0.00052592 | 2.59E-03 |
| OCT_Q6 | 457 | 53.647 | 77 | 1.4353 | 0.00063826 | 2.93E-03 |
| CATRRAGC_UNKNOWN | 239 | 28.056 | 45 | 1.6039 | 0.00086894 | 3.70E-03 |
| E4BP4_01 | 406 | 47.66 | 69 | 1.4478 | 0.0009417 | 3.90E-03 |
| AP1_Q2 | 385 | 45.195 | 66 | 1.4603 | 0.00095563 | 3.93E-03 |
| CDPCR1_01 | 221 | 25.943 | 42 | 1.6189 | 0.001049 | 4.12E-03 |
| ZIC3_01 | 430 | 50.477 | 72 | 1.4264 | 0.0011215 | 4.32E-03 |
| CREB_01 | 422 | 49.538 | 70 | 1.413 | 0.0016763 | 6.05E-03 |
| ER_Q6_01 | 408 | 47.895 | 68 | 1.4198 | 0.0017129 | 6.13E-03 |
| RFX1_01 | 416 | 48.834 | 69 | 1.413 | 0.001805 | 6.39E-03 |
| FOXO1_01 | 385 | 45.195 | 64 | 1.4161 | 0.0024536 | 7.88E-03 |
| E2F_02 | 414 | 48.599 | 68 | 1.3992 | 0.0024883 | 7.88E-03 |
| MYOD_Q6_01 | 457 | 53.647 | 73 | 1.3608 | 0.0036598 | 0.010544 |
| CREB_Q2 | 465 | 54.586 | 74 | 1.3557 | 0.0037971 | 0.010785 |
| SP1_Q6 | 395 | 46.369 | 64 | 1.3802 | 0.0045082 | 0.012341 |
| HIF1_Q5 | 388 | 45.547 | 63 | 1.3832 | 0.0045782 | 0.012421 |
| NRF2_01 | 403 | 47.308 | 65 | 1.374 | 0.0047014 | 0.012586 |
| FOXO3_01 | 367 | 43.082 | 60 | 1.3927 | 0.0047836 | 0.012749 |
| RP58_01 | 353 | 41.438 | 58 | 1.3997 | 0.0049148 | 0.013015 |
| NKX25_02 | 413 | 48.482 | 66 | 1.3613 | 0.0054775 | 0.014162 |
| RORA1_01 | 436 | 51.182 | 68 | 1.3286 | 0.0085232 | 0.019385 |
| NF1_Q6_01 | 449 | 52.708 | 69 | 1.3091 | 0.011251 | 0.024396 |
| AR_Q2 | 188 | 22.069 | 33 | 1.4953 | 0.011569 | 0.024733 |
| E2F_Q3_01 | 376 | 44.138 | 59 | 1.3367 | 0.012024 | 0.025615 |
| HNF3_Q6 | 289 | 33.925 | 47 | 1.3854 | 0.012637 | 0.026639 |
| DR1_Q3 | 371 | 43.551 | 58 | 1.3318 | 0.013637 | 0.028254 |
| TFIIA_Q6 | 431 | 50.595 | 66 | 1.3045 | 0.013943 | 0.02879 |
| ETS_Q4 | 391 | 45.899 | 60 | 1.3072 | 0.017663 | 0.034584 |
| GTTRYCATRR_UNKNOWN | 277 | 32.517 | 44 | 1.3531 | 0.02247 | 0.042088 |
| TAL1ALPHAE47_01 | 404 | 47.425 | 61 | 1.2862 | 0.022922 | 0.04267 |
| RSRFC4_01 | 452 | 53.06 | 67 | 1.2627 | 0.025666 | 0.04663 |
